# Machine learning evaluation of gene expression-based ALS subtypes across brain and blood tissues

**DOI:** 10.64898/2026.06.19.733362

**Authors:** Esteban A. Gomez, Ahmad Al Khleifat, Claire Troakes, Ammar Al-Chalabi, NYGC ALS Consortium, Alfredo Iacoangeli

## Abstract

The clinical and molecular heterogeneity observed in amyotrophic lateral sclerosis (ALS) presents a challenge for diagnosis, prognosis, and treatment. RNA sequencing of post-mortem brain samples from ALS patients has identified several subtypes with distinct molecular signatures. We sought to evaluate these subtypes across diverse tissues and datasets and assess the feasibility of supervised machine learning models for sample classification. Unsupervised clustering and pathway analysis were performed to confirm the presence of ALS subtypes in motor cortex samples. Three machine learning strategies were then used to create models based on post-mortem motor cortex expression data of 112 people with ALS from the London Neurodegenerative Diseases Brain Bank. These models were subsequently improved through feature selection and evaluated in independent cohorts from motor cortex (n = 257, NYGC ALS Consortium) and blood (n = 96, Macquarie University Neurodegenerative Disease Biobank) samples. Multi-class linear discriminant analysis (LDA) models were then used for subtype classification. Clustering of ALS post-mortem motor cortex samples confirmed the presence of three subtypes: neuroinflammation (ALS-Neu), extracellular matrix organisation and muscle contraction (ALS-OxA), and synaptic and neuropeptide signalling (ALS-SNs). Among all machine learning strategies, random forests produced the most accurate and stable models for binary classification (∼93% accuracy across the three subtypes). After feature selection, random forest models were able to classify samples from an independent post-mortem motor cortex cohort in their respective subtypes (AUC of ∼0.98 across the three subtypes). When these models were evaluated in blood using LDA, we found consistent clustering patterns, with samples aligning in the same subtype regions of the post-mortem motor cortex samples, with ALS-SNs being the subtype in which samples were classified with the highest confidence (LDA class probability ∼86%). Moreover, classification for this subtype improved when blood samples were collected closer to death. Our findings support the presence of three gene expression-based ALS subtypes in motor cortex samples and the utility of machine learning strategies for subtype classification. We also observed that the subtypes identified in the brain partially match those in the blood, with samples from the late stages of the disease more likely to be correctly predicted into the ALS-SNs cluster. This suggests a longitudinal effect in subtype identification that requires further investigation.

## Introduction

Amyotrophic lateral sclerosis (ALS) is a neurodegenerative disease causing the progressive loss of both upper and lower motor neurons, leading to muscle weakness, paralysis and ultimately death from respiratory failure ^1^. ALS is characterised by highly heterogeneous clinical and mechanistic landscapes that greatly limit all aspects of ALS research and care, including its diagnosis and the development of treatments ^2–6^.

Machine learning (ML) and high-throughput molecular profiling have been used to identify diverse molecular subtypes of ALS based on gene expression data. Marriott *et al*. ^7^ identified three ALS subtypes using transcriptomic data from post-mortem motor cortex: ALS-Neu, associated with glial activation and neuroinflammation; ALS-OxA, characterised by muscle system and extracellular matrix (ECM) molecular pathways and linked to oxidative stress; and ALS-SNs, associated with neuronal and synaptic signalling-related processes. These results confirmed the subtypes identified in previous studies ^8,9^, specifically glial activation and oxidative stress/ECM organisation clusters, with these studies identifying an additional subtype based on transposable element (TE) data (ALS-TE) across different brain regions, including frontal and motor cortex. In spinal cord samples, various studies have identified diverse subtypes, including clusters of glial activation, oxidative stress, synaptic dysfunction, and neuronal degeneration ^10,11^. In addition, Grima *et al*. ^12^ used whole-blood RNA-seq data to identify 4 new blood-based subgroups that did not directly match brain and spinal cord subtypes. This highlights the potential of subtype identification but also its limitations, with cluster identification differing by tissue and study design, and a lack of validation in independent cohorts or populations. Moreover, because most of these studies used post-mortem samples, the evolution of these subtypes and their signatures throughout disease stages remains unknown and adds an extra layer of complexity that needs to be taken into account if we want to establish robust and meaningful disease molecular subtypes and a new biology-informed disease classification ^13^.

We therefore sought to evaluate the robustness of the current proposed ALS molecular subtypes and to assess the feasibility of developing an ML classifier of disease subtype across tissues, independent samples and disease stages.

## Methods

### Study cohorts

Transcriptomic data were obtained in raw read count matrices from three datasets. The first, used as the training dataset for the machine learning models, consists of 112 post-mortem motor cortex samples from 112 ALS patients from King’s College London and the MRC London Neurodegenerative Diseases Brain Bank (KCL BrainBank) ^14,15^. The second dataset, from the NYGC ALS Consortium (ALS Consortium) ^16^, was downloaded from figshare and the Gene Expression Omnibus (GEO) database (accession code GSE153960) and consists of 257 post-mortem motor cortex tissue samples from 166 ALS patients. This dataset was used for model evaluation, feature selection, and to build the blood classification model. Finally, peripheral blood RNA-seq data from 96 ALS patients (Grima dataset) ^12^ were downloaded from GEO (accession code GSE234297) and used to evaluate the utility of post-mortem motor cortex ML models for ALS subtype classification in blood.

### Data preprocessing and batch correction

Data analyses were performed using R (version 4.4.3) ^17^ and the RStudio environment (version 2026.01.1+403). A gene was considered expressed at a reasonable level if it had 5 counts in at least 10 samples. The R package DeSeq2 (version 1.46.0) ^18^ was used for data normalisation using variance-stabilising transformation (VST) and principal component analysis (PCA). PCs were generated using the top 5000 most variable genes to assess external sources of variability, including batch and technical effects.

Since transcriptomic data were obtained from three different studies and generated using different technologies, a batch effect was expected. Batch effect correction was performed using the “ComBat” function from the R package sva (version 3.54.0) ^19^. Pre-batch correction datasets were harmonised to share the same genes, and samples with zero counts across a single dataset were removed from all datasets. KCL BrainBank was used as the reference batch.

### Unsupervised clustering analysis

Unsupervised clustering of harmonised and combined datasets from KCL BrainBank and ALS Consortium was performed, as reported in previous studies that identified ALS subtypes in the brain ^7,8^. The non-smooth negative factorisation (nsNMF) algorithm from the R package NMF (version 0.28) ^20^ was used to identify clusters using the 5000 most variable expressed genes, with the optimal number of clusters set to three, as is the number of ALS subtypes previously reported in the brain. Relevant genes for cluster separation and sample classification were extracted from these analyses.

To correctly identify the newly found clusters, the associated genes for each cluster were overlapped with those reported by Marriot *et al*. ^7^, and Fisher’s exact test was performed. Moreover, enriched pathway analyses of relevant genes were completed using the R package ClusterProfiler ^21^ for Gene Ontology (GO) enrichment and the R package ReactomePA ^22^ for pathway identification in the Reactome database. Genes from the expression matrix were used as a custom gene background, and *Homo sapiens* was set as the organism’s database. Cluster identity was then corroborated by evaluating enriched pathways in each cluster.

### Supervised machine learning models

The KCL BrainBank dataset was used as the training dataset of the ML models. VST-normalised (after filtering, data harmonisation, and pruning) gene expression of the top 5000 most variable genes was used as features for building the models. Bootstrapping (random sampling with replacement) was used for model validation.

Three ML methodologies were used to create the ALS subtype classifier models: Elastic Net, random forest, and Gradient Boosted Trees (XGBoost). Elastic Net regression is a regularised regression method that combines the constraints of the LASSO and Ridge methods with a penalisation scheme that shrinks coefficients (the contribution of each variable to class classification) to improve the models ^23^. For parameter tuning (⍺ and λ), bootstrapping was used to generate the most stable model, reducing the variance of the prediction error. The R package caret ^24^ (version 7.0.1) was used to create the Elastic Net models.

Random forest works by getting the consensus from weak decision tree classifiers created by randomly separating the explanatory variables ^25^. The R package randomForest ^26^ (version 4.7.1.2), which uses bootstrapping as the test method, was used to build the models. A small loop was created to identify the best *mtry* value, which defines the number of variables randomly sampled as candidates at each split, with *ntree* (number of trees) set to 1,000 to generate stable models.

XGBoost also works with decision trees, but different from random forest, it builds a tree after another by improving the previous one. It’s based on the bias-variance trade-off, where it strikes a balance between a simple yet effective prediction model ^27^. The R package “caret” and xgboost ^28^ (version 3.2.0.1) were used for parameter tuning and model building. Since XGBoost has many parameters, a parameter grid combined with bootstrapping was used to identify the best model settings. The model with the highest accuracy after parameter tuning was selected as the best model.

As all the above ML methodologies are for binary classification and there are three ALS subtypes, ML models for each subtype were created using the “one-vs-the rest” approach, where the multi-class classification is split into one binary classification problem per class.

To determine the optimal number of samples for our models and confirm that sufficient samples were used to generate stable models, we built learning curves using random forest for each ALS subtype. Learning curves were created by increasing the number of available samples and identifying the minimum set of samples for which the model’s accuracy remains high and the variance is small.

Accuracy scores, confusion matrices, and parameter tuning plots were created for all methodologies.

### Model evaluation, improvement and feature selection

Receiver operating characteristic (ROC) curves were constructed to evaluate the models’ prediction accuracy when classifying samples in the evaluation dataset. The area under the curve (AUC), sensitivity, and specificity were used to assess the models’ predictive performance. ROC curves were created using the R package pROC ^29^ (version 1.19.0.1).

Since the random forest achieved the best accuracy during the validation step, the models were improved using this methodology. To identify the most relevant genes for subtype classification, the “importance” function of the R package randomForest was used, which ranks features by relevance based on how much the model’s accuracy decreases when that feature is excluded. A gene was considered relevant if the model’s mean accuracy decreased by more than 2%. Then, two strategies were applied to filter relevant genes for subtype classification: the *delete* strategy used a loop to repeatedly fit models with same parameters, excluding the least importance feature in each iteration and identifying the minimum number of features were the accuracy of the models improved; on the other hand, the *remove* strategy eliminate genes in each interaction and verify the accuracy of the models, keeping only the genes that increase or don’t reduce the accuracy of the models.

After identifying the relevant genes, new models were built using the ALS Consortium dataset, and they were evaluated by calculating the AUC of the ROC curves when classifying the KCL BrainBank samples.

### Machine learning model evaluation in blood samples

For multi-class prediction of ALS subtypes in peripheral blood samples (Grima dataset), linear discrimination analysis (LDA) models were built. LDA solves multi-class classification by separating data through dimensionality reduction. LDA works by identifying a linear combination of features that separates two or more classes so they can be more easily classified ^30^. The R package caret was used to build the models.

Before fitting the model with genes from the feature selection step, the genes were filtered to exclude those expressed in the brain but not in the blood. For that purpose, a list of genes expressed in the brain and blood was obtained from The Human Protein Atlas ^31^, with genes not on the list removed before creating the LDA models. In addition, since LDA is affected by collinearity, weighted gene co-expression network analysis (WGCNA) was performed using the R package WGCNA ^32^ (version 1.74) to establish modules based on similarity in gene expression and identify genes that don’t co-express with genes in other modules. The function “pickSoftThreshold” was used to automatically identify the ideal soft threshold to get a scale of independence of 0.6 (this was done to reduce the strictness of the analysis and identify high average connectivity). The top 10% of genes ranked by connectivity for each module, and genes without co-expression, were selected to reduce collinearity in the LDA models.

The LDA model was trained using the ALS Consortium dataset and evaluated in Grima’s samples. Evaluation performance was defined as the percentage of samples assigned to a specific class with high probability (> 85%), and a classification was considered good if more than 85% of the samples classified to a subtype were predicted with high probability. In addition, and based on the LDA model’s sample classification, three-way differential gene expression (DGE) analysis was performed using the R packages Volcano3D ^33^ (version 2.0.11) and DeSeq2 to identify differentially expressed genes in blood across ALS subtypes. Data normalisation and filtering (as mentioned in the data preprocessing section) were applied before analysis, and the likelihood ratio test was used, considering sex, age, and RNA integrity number (RIN) of the samples as covariates. Volcano 3D plots were generated to identify significant genes, and enrichment analysis was performed as described before. Matching between pathways identified in these analyses and the original unsupervised clustering enrichment analysis was used to assess the similarity of molecular signatures, indirectly evaluating the classification performance of the LDA models.

### Disease progression-associated analysis

Since ML models trained on post-mortem brain samples were used to classify pre-mortem blood samples, we wanted to explore whether prediction performance varied by disease stage. For that purpose, we conducted a Spearman correlation analysis between “Collection Point” (Min-Max scaled to range from 0 to 1) and “Probability of assignment” as predicted by the LDA models on disease samples from the Grima dataset. Additionally, to test the combined effect of disease status and subtype classification on cluster prediction in blood, we employed Beta regression models with the final model *Predicted probability ∼ Collection Point * ALS Subtype + Sex + Age of Collection*, where “Collection Point” was either a continuous (range from 0 to 1) or categorical (early vs late) variable, and “Sex” and “Age of Collection” were set as covariates. Coefficient values, effects and *p* values were extracted from the resulting models. Beta regression models were built using the R package betareg ^34^ (version 3.2.4), and correlation and prediction plots were generated using the R package ggeffects ^35^ (version 2.3.2). Finally, the DGE of the mean of scaled expression values of candidate genes used to build the LDA models was calculated to assess the up- or downregulation of these genes at the early or late stages of the disease.

## Results

### ALS cohorts

To validate ALS subtypes across tissues and explore the utility of supervised ML models for sample classification, we obtained access to three datasets: KLC BrainBank (112 samples) and ALS Consortium (257 samples) for post-mortem motor cortex ALS samples and Grima (96 samples) for peripheral blood ALS samples (Fig. 1). A summary of the demographic and clinical information for the datasets is provided in Table 1. The three datasets present similar demographics, with all three having a male-to-female ratio of ∼1.36 and a similar age of onset of ∼60 years. Disease duration varies between the brain and blood samples, with motor cortex samples having an approximate median disease duration of 37 months, while blood samples presented a longer median disease duration (∼52 months).

**Figure 1.**
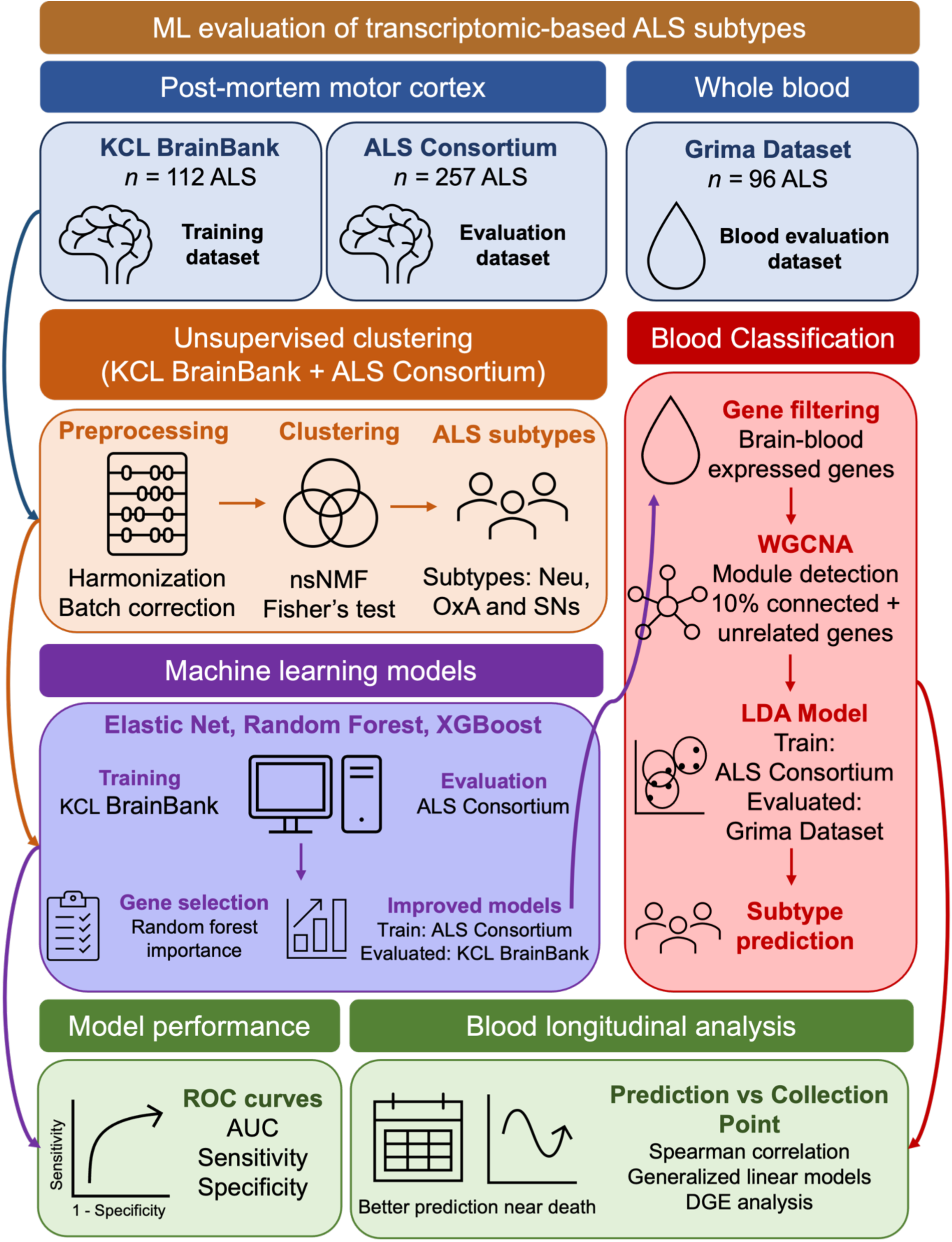
Study design. Experimental design and analysis workflow for evaluation of gene expression-based ALS subtypes across brain and blood tissues.

**Table 1.** Demographics of ALS participants for post-mortem motor cortex and blood cohorts.

| Demographics | KCL BrainBank | ALS Consortium | Grima |
| --- | --- | --- | --- |
| **ALS tissue samples** | 112 (112 patients) | 257 (166 patients) | 96 (96 patientes) |
| **Tissue** | Motor Cortex | Motor Cortex | Peripheral blood |
| **Male (%)** | 65 (58.04%) | 113 (54.3%) | 58 (60.41%) |
| **Age (median ± SD)\*** | 69 ± 12.71 | NA | 66.15 ± 13.75 |
| **Post-mortem delay in hours (median ± SD)** | 27 ± 12.08 | NA | NA |
| **Age of Onset (median ± SD)** | 60.5 ± 13.20 | 61.25 ± 10.08 | 61.95 ± 13.96 |
| **Disease Duration in months (median ± SD)** | 35 ± 18.75 | 39.2 ± 27.71 | 52.25 ± 51.36 |

\* Age refers to the age of death in the KCL BrainBank and the age of blood collection in Grima

### Unsupervised clustering identified consistent ALS subtypes in the KCL BrainBank and ALS Consortium datasets

To validate the presence of similar ALS subtypes in post-mortem motor cortex samples from the KCL BrainBank and ALS Consortium datasets, we first addressed technical variation between the datasets arising from samples generated in different batches and across different sequencing platforms and studies. PCA analysis demonstrated a clear separation between the two datasets (Supplementary Fig. 1A), with ALS Consortium samples clustered together despite being processed in two different batches and using two different sequencing platforms (HiSeq and NovaSeq). We applied ComBat for batch correction, which adjusts the data to match means and variances across batches and removes technical noise using empirical Bayes methods ^36^. PCA after batch correction showed no clear separation of the samples, forming a single, overlapping distribution, indicating effective removal of technical variability (Supplementary Fig. 1B).

After batch correction, we then combined the two datasets (369 post-mortem motor cortex samples in total) and identified ALS subtypes using the nsNMF-based unsupervised clustering of the top 5,000 most variable genes. Here, we found three clusters with distinctive molecular signatures (Fig. 2A). PCA analysis demonstrated clear separation between samples assigned to each cluster, with 20% of the samples assigned to Cluster 1, 28% to Cluster 2 and 52% to Cluster 3 (Fig. 2B and 2C).

**Figure 2.**
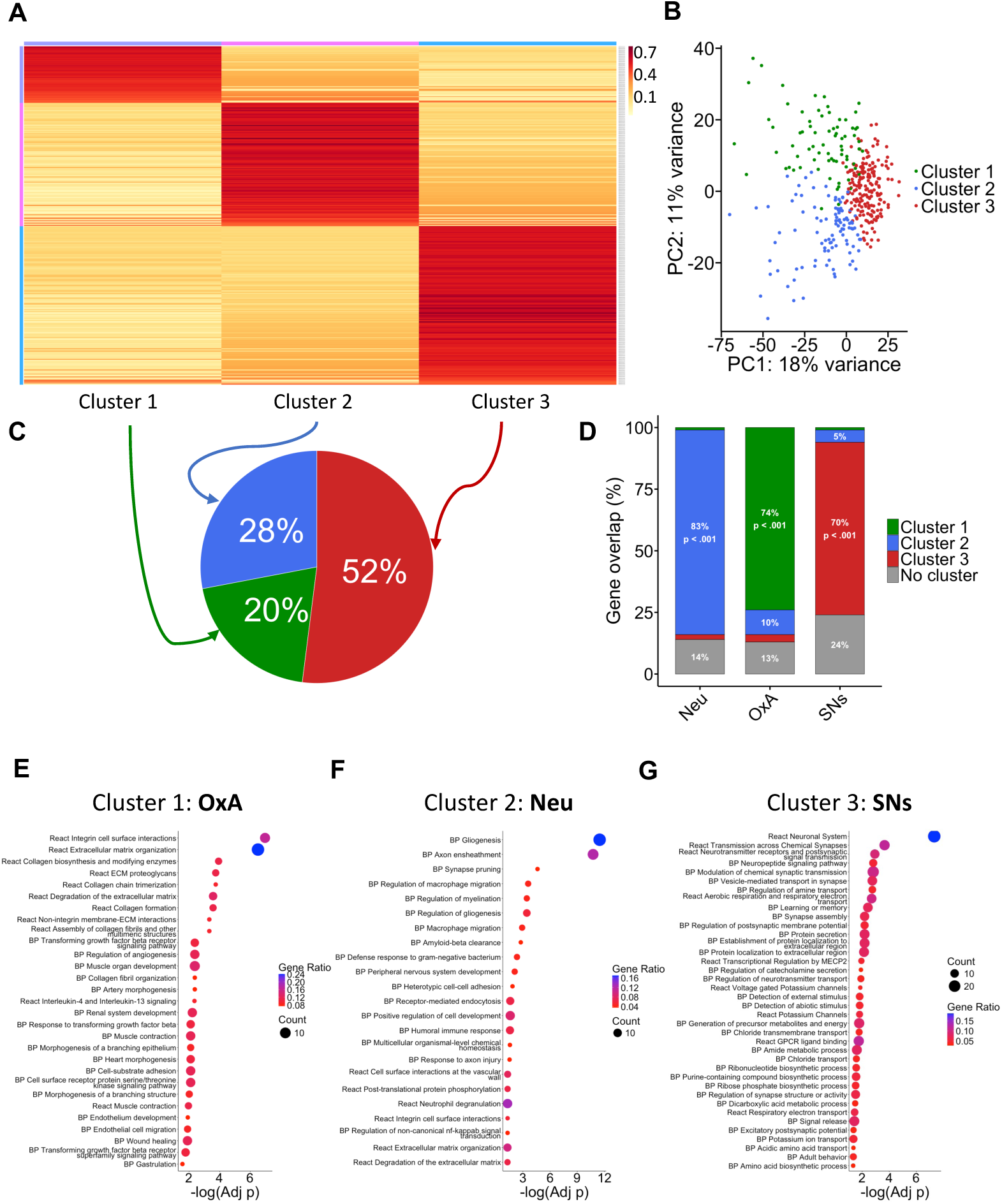
Unsupervised clustering analysis of post-mortem motor cortex transcriptomes identifies previously described ALS subtypes. Transcriptomic data of 369 post-mortem motor cortex samples from the KCL BrainBank and ALS Consortium were combined after batch correction, and nsNMF-based unsupervised clustering was performed on the top 5,000 most variable genes. (A) Heatmap showing 3 clusters based on the top 5,000 most variable genes. (B) PCA plot showing samples by ALS subtype. (C) Distribution of samples for each ALS subtype. (D) Bar plot displaying gene overlap between associated genes for each cluster and relevant genes associated with each ALS subtype (ALS-Neu, ALS-OxA, and ALS-SNs) identified by Marriot *et al.* (p-values for Fisher’s exact test are displayed). (E, F and G) Enrichment pathway analysis (GO:BP terms and Reactome database; p-values for Fisher’s exact test after BH correction are displayed) for the three clusters (ALS-OxA, ALS-Neu and ALS-SNs, respectively).

ALS subtypes have previously been reported using unsupervised clustering and transcriptomics from post-mortem motor cortex samples; to assess whether the clusters identified in our analysis corresponded to those previously reported by Marriot *et al*. ^7^ (ALS-Neu, ALS-OxA and ALS-SNs), we performed hypergeometric enrichment analysis (Fisher’s exact test) comparing the genes associated with each cluster to the subtype-related genes reported in that study (Supplementary Table 1). We selected the results from this study because it was the only one focused on motor cortex samples and didn’t integrate transposable elements as features for clustering. From Cluster 1, 216 genes overlapped with those from ALS-OxA (74%, p < .001), including the muscle contraction-associated genes *MFAP4*, *MRGPRF* and *PLAUR*; Cluster 2 was linked with the ALS-Neu subtype (310 overlapped genes, representing 83% of genes from this subtype; p < .001), characterized by the presence of neuroinflammation-related genes such as *TREM2*, *C3* and *TLR2*. Finally, Cluster 3 has 92 genes that are common with the ALS-SNs subtype (70%, p < .001), including the neuronal and synaptic signalling-associated genes *ADCYAP1*, *DRD5*, *VIP* and *NPY2R* (Fig. 2D).

In addition, we explored the molecular signatures of each cluster by using enrichment pathway analysis (GO and Reactome databases; Supplementary Table 2) and found that Cluster 1 was significantly enriched with extracellular-matrix and muscle associated pathways, such as ECM organization, collagen biosynthesis, muscle contraction and muscle organ development; furthermore, we also identified pathways associated with angiogenesis, epithelial cell development and peptide ligand-binding receptors, as reported for the ALS-OxA subtype ^7,8^ (Fig. 2E). Although we did not detect significant enrichment for oxidative stress-related pathways, the cluster showed strong and statistically significant overlap with the previously reported ALS-OxA subtype, and we therefore retained the ALS-OxA designation to reflect this concordance. Cluster 2, as the ALS-Neu subtype, was enriched with pathways associated with neuroinflammation, such as gliogenesis, regulation of macrophage migration, humoral immune response and neutrophil degradation (Fig. 2F), while Cluster 3 was enriched for neuronal and synaptic pathways (e.g. neuropeptide signalling, regulation of postsynaptic membrane potential, regulation of synapse structure and transmission across chemical synapses) (Fig. 2E), mimicking the ALS-SNs subtype.

Of note, since Eshima *et al*. and Tam *et al.* utilised ALS Consortium data and added TEs for their clustering analysis, identifying a TE-associated subtype (ALS-TE) instead of the ALS-SNs cluster found by Marriot *et al*. ^7–9^ in the KCL BrainBank data, we rerun the nsNMF-based unsupervised clustering in the ALS Consortium dataset motor cortex transcriptomic data to investigate whether the clusters identified in the joint dataset were present also when the ALS Consortium dataset was analysed on its own. Once again, we found three distinct clusters with their own molecular signature (Supplementary Fig. 2A, 2B and 2C). Gene’s overlapping and enrichment analysis confirmed the three ALS-subtypes identified previously by Marriot *et al.*, with genes and pathways associated with neuronal signalling (ALS-SNs), ECM organisation (ALS-OxA), and neuroinflammation (ALS-Neu), with the ALS-OxA subtype displaying some pathways (e.g. central nervous system myelination and nervous system development) shared with the ALS-SNs subtype (Supplementary Fig. 2D). We then explored the distribution of samples in ALS subtypes before and after combining datasets; here, we found that both KCL BrainBank and ALS Consortium samples belonging to the ALS-Neu subtype remain consisting in their classification (Supplementary Fig. 3A and 3B), with ALS-SNs samples also remaining in the same cluster for KCL BrainBank. In the case of the ALS-OxA samples, we observed a more widespread distribution between ALS-OxA and ALS-SNs, similar to what is observed in the ALS Consortium samples, where samples originally classified as ALS-OxA were predicted to be ALS-SNs, in line with the overlapping enriched pathways we saw for ALS-OxA and ALS-SNs. In the case of the ALS-TE (TE-related subtype) samples from the original analysis performed by Eshima *et al*, they spread across all subtypes, with a higher prevalence associated with the ALS-OxA (Supplementary Fig. 3B), which was already described by Eshima *et al*., when they found overlap in disease mechanism between the ALS-TE and ALS-OxA subtypes ^8^. Of Note, Eshima *et al*.’s analysis included not only motor cortex samples but also frontal cortex samples, which may explain the divergence in sample distribution. Finally, we explored whether samples with the *C9orf72* expansion and genetic variants in *SOD1* in both datasets were clustered in the same ALS subtype, finding an even distribution of these mutation carriers across all subtypes (Supplementary Table 3).

Overall, these results show that, after batch correction, motor cortex datasets can be effectively integrated to identify consistent ALS molecular subtypes. The three clusters identified recapitulate subtypes previously reported by Marriott *et al*., with concordant gene signatures and pathway enrichment, supporting their biological relevance. While most samples remained stable across datasets, some overlap between subtypes suggests partial convergence of underlying molecular mechanisms.

### Machine learning classifiers trained on transcriptomic data can predict ALS subtypes

Having found that ALS subtypes remain consistent across post-mortem motor cortex datasets, we next investigated whether gene expression data were predictive of ALS subtypes. For that purpose, we built ML models using the top 5,000 most variable genes from the KCL BrainBank dataset (training dataset with ALS subtype labels obtained from the unsupervised clustering) and three different ML methodologies: Elastic Net, XGBoost and random forest. Bootstrapping was used for model validation, and the ALS Consortium was used as the independent test cohort for evaluation. First, we created learning curves for the different ALS subtypes (“one-vs-the rest” approach) and corroborated that we got enough samples to build stable models, with 60 samples seeming to be the minimum required to have stable models (Supplementary Fig. 4). We performed parameter tuning for all the models (Supplementary Fig. 5) and compared the accuracy scores for all the ML methodologies (Table 2). Overall, the three machine learning methodologies were able to classify the ALS subtypes with high accuracy (∼ 90%), with random forest models performing the best in terms of accuracy (ALS-Neu: 90%, ALS-OxA: 95%, ALS-SNs: 85%), sensitivity (ALS-Neu: 82%, ALS-OxA: 84%, ALS-SNs: 95%) and specificity (ALS-Neu: 93%, ALS-OxA: 98%, ALS-SNs: 95%). Elastic Net and XGBoost also performed well in terms of sensitivity and specificity, with a slight drop in sensitivity across all three subtypes, particularly for ALS-Neu and ALS-OxA (Fig. 3A and 3B).

**Figure 3.**
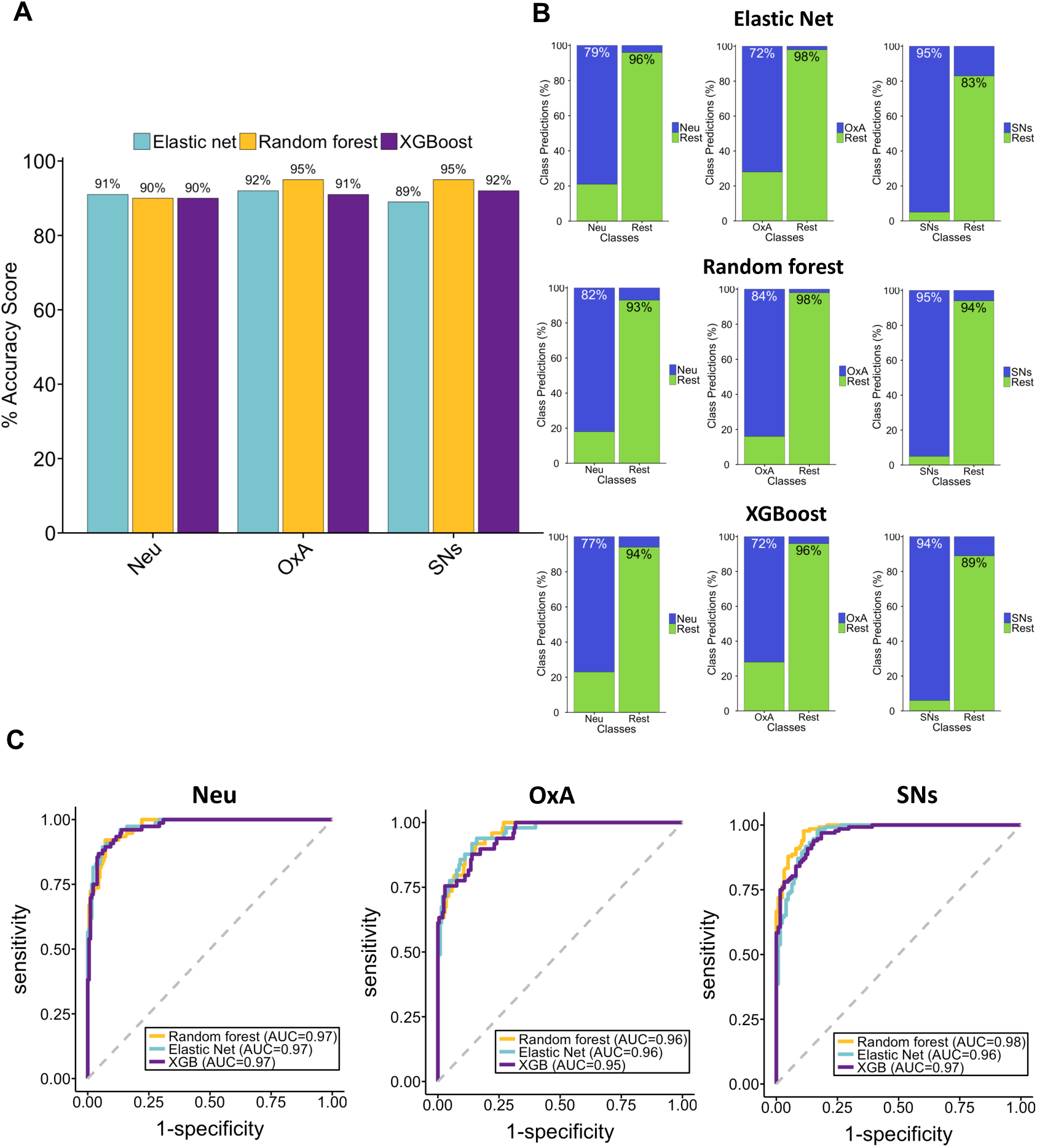
Machine learning classifiers trained on transcriptomic data can predict ALS subtypes. ML models to classify ALS subtypes were created using the top 5,000 most variable genes from 112 post-mortem motor cortex samples from the KCL BrainBank (training dataset). (A) Percentage accuracy score of prediction models based on transcriptomic data for ALS subtype using Elastic net, random forest and XGBoost methodologies. (B) Classification predictions for each class (sensitivity and specificity) of the ML models *(top*: Elastic net, *middle*: random forest, *bottom*: XGBoost) for each ALS subtype *(left*: ALS-Neu, *center*: ALS-OxA, *right*: ALS-SNs). Blue indicates the samples were predicted as the specific subtype, while green indicates those patients don’t belong to that cluster. Percentages indicate true positives and true negatives. (C) ROC curves and AUC values for predictive models evaluated in an independent cohort (ALS Consortium; 257 post-mortem motor cortex samples) for ALS-Neu (*left*), ALS-OxA (*center*) and ALS-SNs (*right*) subtypes.

**Table 2.** Summary of classification models for the prediction of ALS subtypes.

| ML strategies | Training |  |  |  |  |  | Evaluation |  |  |  |  |  |
| --- | --- | --- | --- | --- | --- | --- | --- | --- | --- | --- | --- | --- |
|  | ALS Subtype | Subtype samples | Other samples | Accuracy (%) | Sensitivity | Specificity | Subtype samples | Other samples | Accuracy (%) | AUC | Sensitivity | Specificity |
| Random Forest | Neu | 28 | 84 | 90 | 0.82 | 0.93 | 76 | 181 | 84 | 0.97 | 0.7 | 0.99 |
| Elastic net |  |  |  | 91 | 0.79 | 0.96 |  |  | 83 | 0.97 | 0.67 | 0.98 |
| XGBoost |  |  |  | 90 | 0.77 | 0.94 |  |  | 85 | 0.97 | 0.71 | 0.98 |
| Random Forest | OxA | 25 | 87 | 95 | 0.84 | 0.98 | 49 | 208 | 80 | 0.96 | 0.63 | 0.98 |
| Elastic net |  |  |  | 92 | 0.72 | 0.98 |  |  | 84 | 0.96 | 0.69 | 0.98 |
| XGBoost |  |  |  | 91 | 0.72 | 0.96 |  |  | 81 | 0.95 | 0.63 | 0.98 |
| Random Forest | SNs | 59 | 53 | 95 | 0.95 | 0.94 | 132 | 125 | 91 | 0.98 | 0.93 | 0.9 |
| Elastic net |  |  |  | 89 | 0.95 | 0.83 |  |  | 89 | 0.96 | 0.92 | 0.87 |
| XGBoost |  |  |  | 92 | 0.94 | 0.89 |  |  | 88 | 0.97 | 0.86 | 0.89 |

To validate the robustness of our models and demonstrate that they do not overfit, we tested whether they could classify ALS subtypes in the ALS Consortium dataset. To this end, we employed ROC curves to evaluate the classifier’s diagnostic performance across various decision thresholds. Evaluation of the ROC curves displayed an AUC > 95% for all the models across the three subtypes, with random forest producing the best results (Fig. 3C). Exploring the accuracy of the models in the ALS Consortium dataset, we observed an average accuracy for all models of 85%, with the ALS-SNs models performing the best in terms of accuracy (∼89%) and also sensitivity (∼90%), and with the ALS-Neu and ALS-OxA showing an average sensitivity of 70% and 65%, respectively (Table 2). Thus, these results indicate that gene expression data can predict and classify ALS subtypes from post-mortem motor cortex ALS samples.

### Feature selection identified candidate genes relevant for the classification of ALS subtypes in brain and blood

Since the best predictors are those with the fewest features and the random forest models achieved the best overall performance, we next performed feature selection using the “importance” function in the random forest methodology. Here, and using the ALS Consortium as the training dataset to avoid overfitting, as the candidate genes are identified based on their ability to predict across datasets rather than capturing dataset-specific noise or technical variation from the original dataset, we applied two stepwise approaches: *delete*, which creates ML models progressively eliminating genes until it identifies the minimum number of genes for which the accuracy of the models is best; and *remove*, which builds models eliminating genes that reduce their accuracy (Supplementary Fig. 6A and 6B). Overall, the ALS-Neu classifier benefited from the *remove* strategy, with an ∼7% increase in accuracy (96% for both strategies). In contrast, the accuracy of the ALS-OxA and ALS-SNs models was not affected by the feature selection (Supplementary Fig. 6C and Supplementary Table 4). However, when evaluating these new models in the KCL BrainBank dataset, we observed a general increase in the AUC of the ROC curves (∼99%) that was reflected in an improvement of the sensitivity of all the models (ALS-Neu: 89%, ALS-OxA: 96% and ALS-SNs: 98%) (Supplementary Fig. 6D and Supplementary Table 4).

For the ALS-Neu, 60 candidate genes were identified after feature selection using the *remove* strategy, including the neuroinflammation-related genes *TREM2*, *FCGR1A* and *ALOX5AP*. In contrast, for ALX-OxA, 79 genes were highlighted as relevant to this subtype identification, including *LAMA5*, *ITGA10,* and *ADAMTS10*, which are associated with ECM interactions and muscle biology. Finally, ALS-SNs identified 136 relevant genes, doubling the previous ones, with some of the genes, such as *PTBP1*, *WASF2* and *ATP6V1G2*, associated with synaptic function and neuropeptide signalling (Fig. 4A and Supplementary Table 5).

**Figure 4.**
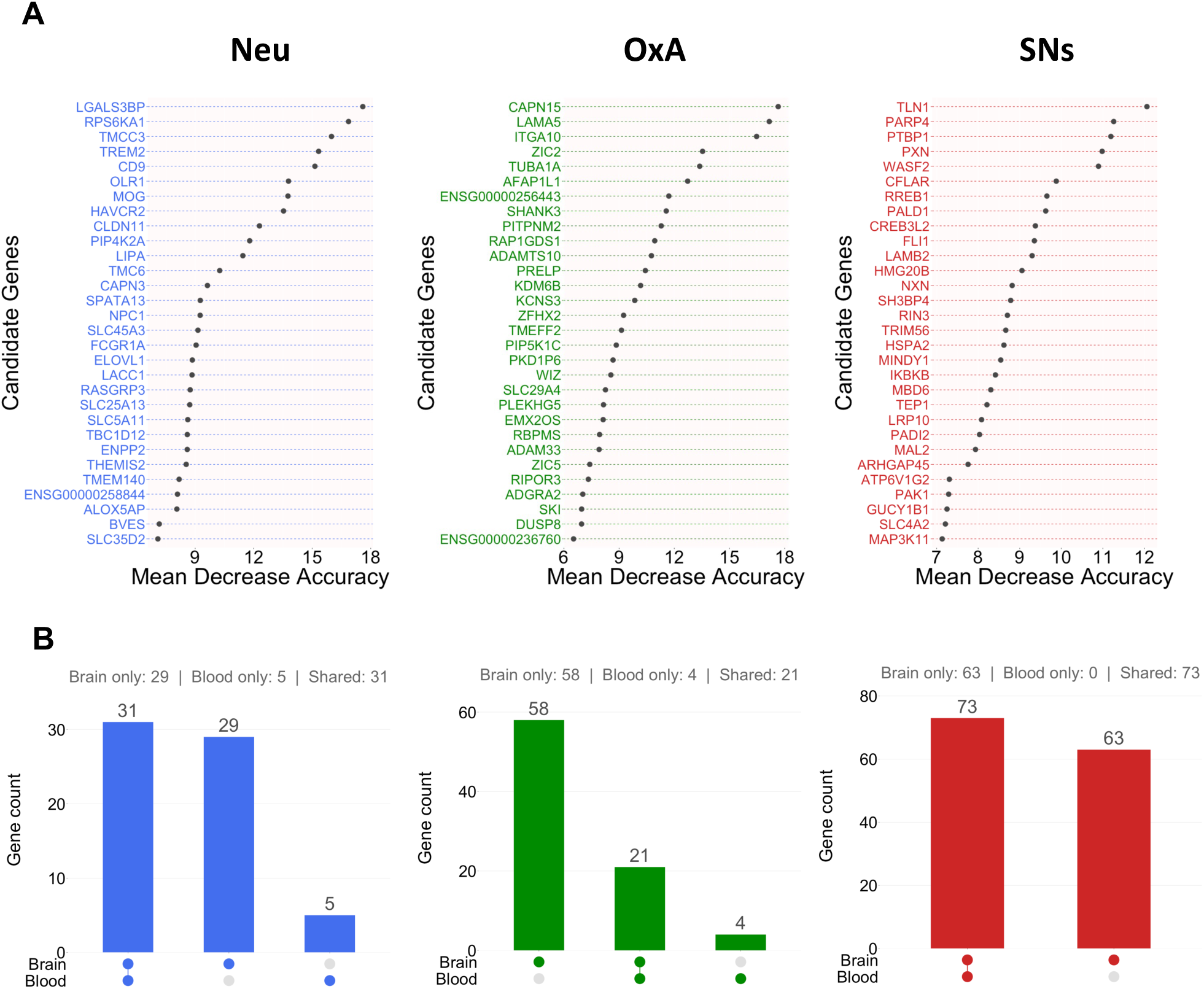
Candidate biomarkers for ALS subtype classification. Feature selection was performed based on the random forest methodology (“importance” function), using the *remove* strategy (building of classifiers by eliminating genes that reduce their accuracy) and the ALS Consortium (257 post-mortem motor cortex samples) as the training dataset. (A) Top 30 most relevant candidate genes for the prediction performance of the ALS-Neu (*left*), ALS-OxA (*center*) and ALS-SNs (*right*) classifiers based on decreasing accuracy. (B) UpSet plots displaying overlapping candidate genes between the “only blood” prediction models and the post-mortem motor cortex-based ML models (*left*: ALS-Neu, *center*: ALS-OxA, *right*: ALS-SNs).

Since we wanted to explore if ML models built using post-mortem motor cortex data were able to classify ALS subtypes from peripheral blood samples, first, we filtered out the candidate genes from the original models that are exclusively expressed in the brain, generating a list of genes expressed in both the brain and blood. Then we performed the *remove* approach for feature selection and built the best models based on these genes (Supplementary Fig. 7A and Supplementary Table 5). For the ALS-Neu model, 36 genes were identified as relevant features for this subtype, with 31 genes overlapping between the brain and blood models and 5 genes, including *SPP1* and *FCGR3A*, associated with neuroinflammation, important only for the blood model. In the ALS-OxA model, a set of 25 genes contributed to subtype classification, with most shared between brain and blood. A smaller subset, including *TSPAN4* and *SNCA*, was specific to the blood model. The ALS-SNs model comprised 73 genes, all of which were also present in the brain and blood model (Fig. 4B, Supplementary Fig. 7A, and Supplementary Table 5). Of note, model improvement and feature selection, while only considering genes that are expressed in blood, didn’t affect the subtype classification of brain samples (Supplementary Fig. 7B and 7C). This demonstrates that genes expressed in brain and blood can be used to create ALS subtype classifiers.

Having identified blood candidate genes for ALS subtype classification, we next explored whether these genes predicted ALS subtypes in blood samples from ALS patients. For that purpose, we first performed WGCNA to identify representative genes correlated with others, as well as genes without co-expression, to avoid collinearity in the multi-class LDA models. Based on this analysis, 53 genes were selected, and a multiclassification model was created using the ALS Consortium dataset as the training set (Supplementary Table 6). LDA projection (LD1 and LD2) revealed a clear separation of the samples into three distinct clusters, corresponding to each subtype (Fig. 5A), indicating that the selected features capture subtype-specific molecular variation. To assess whether this model could classify ALS subtypes from blood samples, we evaluated it on the Grima dataset, which comprises RNA-seq data from peripheral blood of 96 ALS samples. LDA projection of these samples showed consistent clustering patterns, with samples aligning in the same subtype regions of the post-mortem motor cortex, with the ALS-SNs subtype being the predominant cluster. In addition, all subtypes showed a high proportion of samples assigned to their respective clusters with high confidence (>85%), with ALS-SNs exhibiting 86% of samples predicted with high probability, consisting with previous findings by Marriot *et al*.^7^ (Fig. 5B). When using the random forest binary classifiers to predict membership of blood samples in each ALS subtype, we found that the ALS-SNs identified the highest percentage of samples (18%), follow by ALS-OxA (5%) and ALS-Neu (5%), with 72% of the samples not being classified in any subtype (Supplementary Fig. 8). Thus, suggesting some degree of overlap, especially for ALS-SNs, between the subtypes identified in brain and blood.

**Figure 5.**
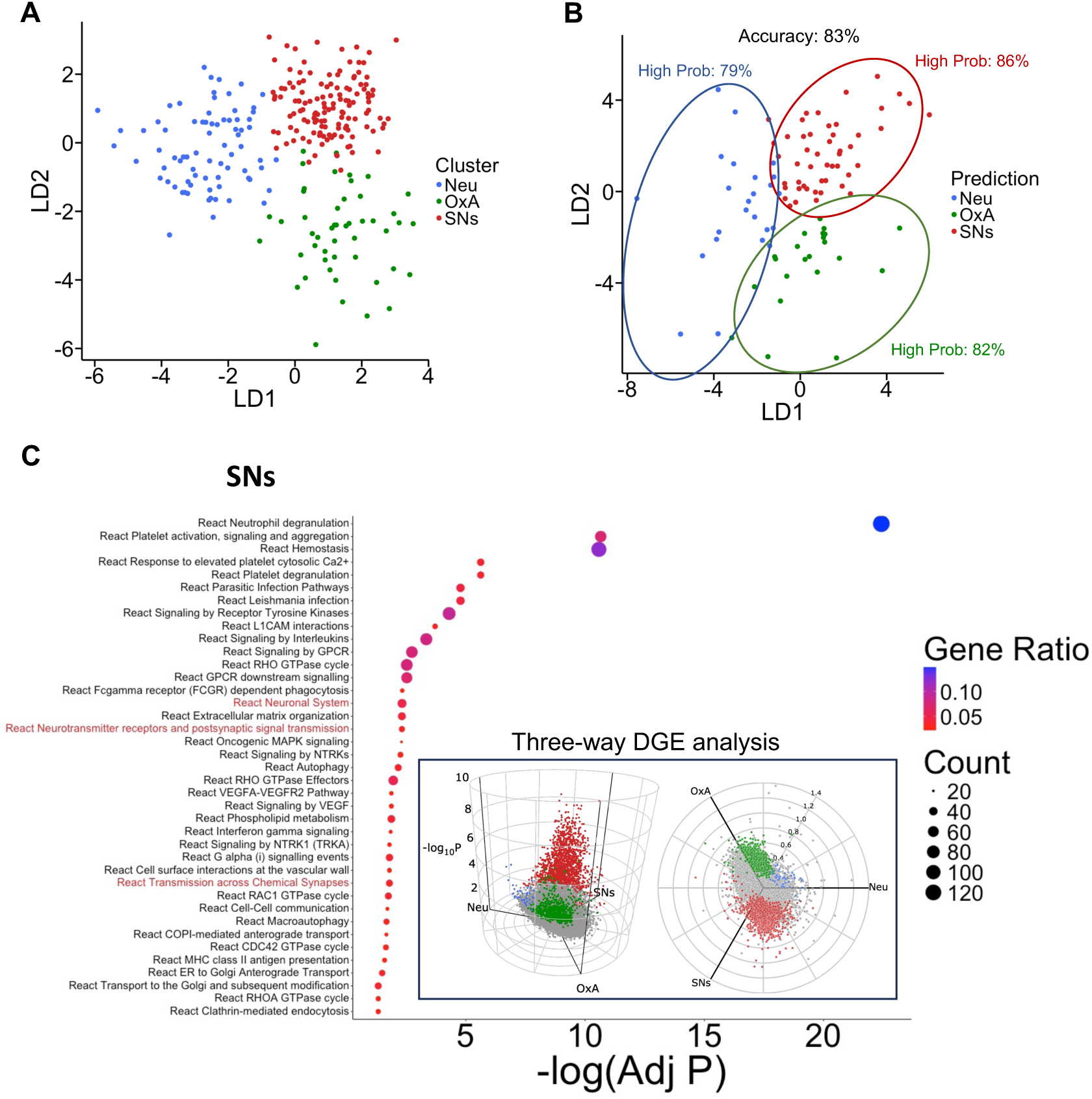
ALS subtype classification of blood samples using a post-mortem motor cortex-based multi-classifier. WGCNA was applied to identify representative genes correlated with others and genes without co-expression and multi-class LDA models were created using ALS Consortium (257 post-mortem motor cortex samples) as the training dataset and evaluated in blood samples (Grima dataset). (A) LDA projection (LD1 and LD2) of post-mortem motor cortex samples into three distinct ALS subtypes. (B) LDA projection displaying the classification of 96 blood samples using the post-mortem motor cortex-based multi-classifier. Accuracy of the model is showing at the top, while percentage of samples classified with high probability (>85%) for each ALS subtype is displayed in colours (*blue*: ALS-Neu, *green*: ALS-OxA, *red*: ALS-SNs). (C) Three-way DGE analysis were performed based on blood sample classification and significantly expressed genes for each ALS subtype were used for enriched pathway analysis. *In the box*: 3D Volcano plot of three-way DGE analysis. Enriched pathway analysis (Reactome database) for the ALS-SNs is shown, and terms highlighted in red indicates pathways previously identified for that subtype in the unsupervised clustering analysis.

Next, to evaluate whether the blood samples were correctly classified into their subtypes, we performed a three-way DGE analysis to identify subtype-specific differentially expressed genes between samples classified as ALS-Neu, ALS-OxA, or ALS-SN by the LDA model (24, 22, and 50 samples, respectively). Here, we identified 41 genes associated with ALS-Neu, 779 genes associated with ALS-OxA, and 1358 genes associated with ALS-SNs (Fig. 5C and Supplementary Table 7). We then performed pathway enrichment analysis to determine whether the resulting molecular signatures matched those identified in the unsupervised clustering analysis. From this analysis, ALS-SNs displayed pathways from the Reactome database associated with neuronal processes, such as Neuronal System, Neurotransmitter receptors and postsynaptic signal transmission, and Transmission across Chemical synapses. In addition, other pathways relevant to inflammation were present, such as neutrophil degranulation and MHC II antigen presentation. Thus, these results support the presence of an ALS subtype in blood that resembles the ALS-SNs subtype identified in brain, particularly through the enrichment of neuronal and synaptic pathways, while also capturing additional immune-related processes that may reflect systemic components of the disease (Fig. 5C and Supplementary Table 8). In contrast, samples classified as ALS-OxA in blood were primarily enriched for mitochondrial and RNA processing pathways, including Mitochondrial translation termination and mRNA splicing (Supplementary Fig. 9 and Supplementary Table 8), suggesting that this subtype may reflect distinct molecular processes in blood that are not fully captured in post-mortem motor cortex. ALS-Neu associated genes were not significantly enriched for any pathway, most likely due to the limited number of differentially expressed genes identified for this subtype. Overall, these findings demonstrate that, although blood samples maintain clustering patterns consistent with the brain samples as shown by LDA, ML models trained on brain-derived data can identify overlapping ALS subtypes in blood only in a limited proportion of samples, with the ALS-SNs subtype being the most frequently identifiable and exhibiting the most consistent pathways across tissues, supporting a shared subtype signature.

### Subtype associated with synaptic and neuropeptide signalling is more likely to be predicted in blood in close-to-death samples

Having found comparable molecular signatures of ALS subtypes in post-mortem brain samples and in blood samples, specifically for the ALS-SNs subtype, we next investigated whether blood samples collected near death were more likely to be classified into these subtypes and whether there was a longitudinal effect on the model’s ability to classify samples correctly. For that purpose, we employed Beta regression models to evaluate the effect of blood collection point on the LDA model’s predicted probability. Here, we found that the effect of disease progression differed by subtype. ALS-SNs subtype showed a higher probability of being classified as this subtype when samples were collected close to death, whereas ALS-Neu probability decreases with progression. These effects were observed for ALS-SNs and ALS-Neu either when the collection point was considered as a range (continuous range from 0 to 1; odds ratio = 23.52, p = 0.004, and odds ratio = 0.07, p = 0.003; respectively) (Fig. 6A and Supplementary Table 9) or as a categorical variable (early vs late; odds ratio = 4.22, p = 0.02, and odds ratio = 0.33, p = 0.03; respectively) (Fig. 6B and Supplementary Table 9). Spearman correlation analysis corroborates a trend of a reduction in probability of assignment for the ALS-Neu (*R =* −0.42, p = 0.058) (Fig. 6C). In addition, DGE of the mean of scaled expression values of ALS-OxA-associated genes used to build the LDA model demonstrated a downregulation in the gene expression of these genes in the late stage of the disease (Fig. 6D). This suggests a reduction in the discriminative signal used by the LDA model, leading to poorer classification for this subtype. Thus, these results indicate that model performance is driven by subtype-specific molecular signatures across tissues and changes over the course of the disease. In particular, ALS-SNs show increased predictability in blood at later stages, supporting the presence of shared biomarkers between brain and blood samples. In contrast, ALS-Neu and ALS-OxA display reduced detectability, likely due to weaker or diminishing molecular signals in blood, which contributes to the overall decline in classification performance. Together, these findings highlight a longitudinal effect on subtype identification, in which the ability to detect specific molecular signatures in blood depends on disease stage and subtype biology.

**Figure 6.**
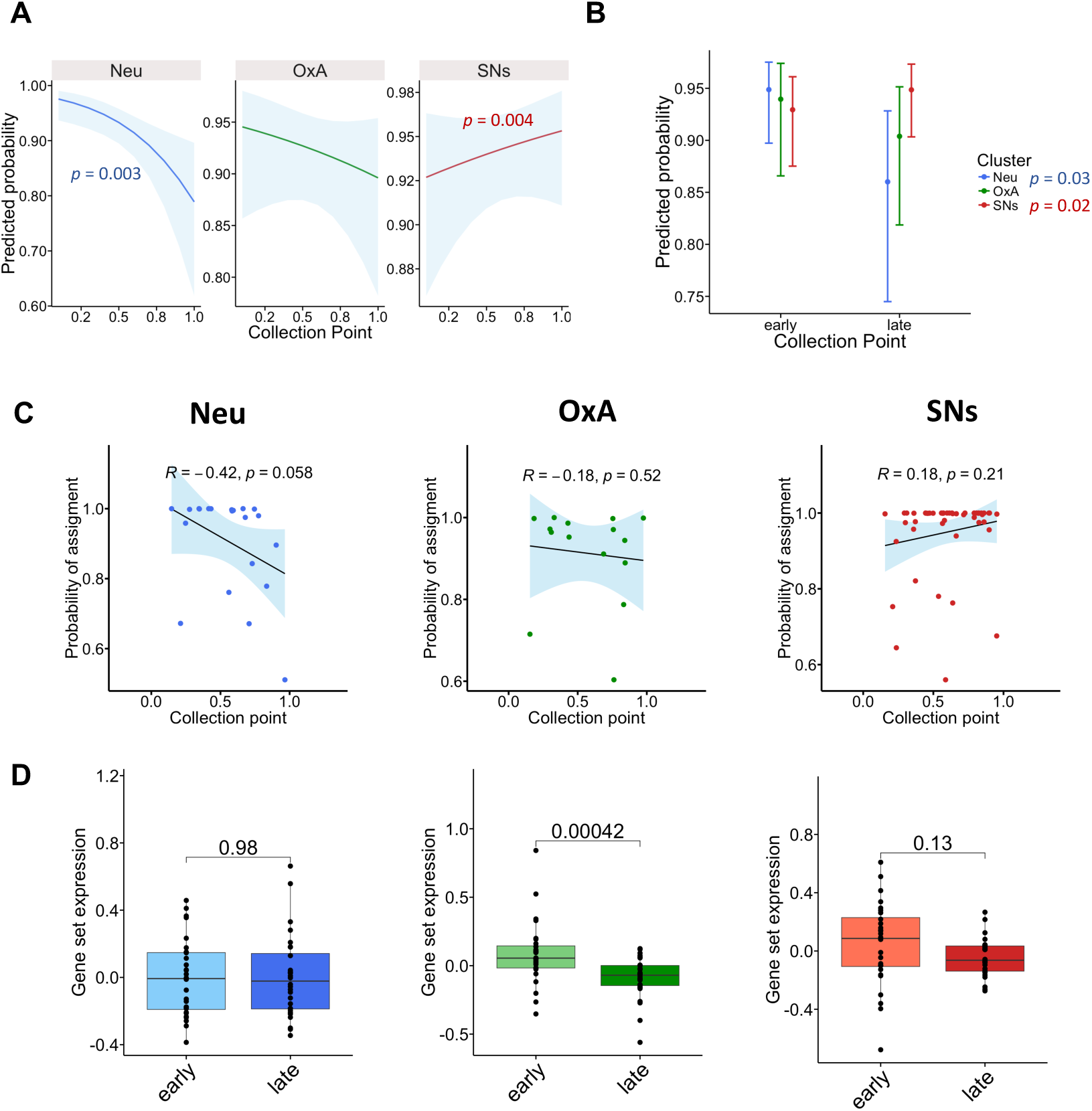
Classification of blood samples from the ALS-SNs subtype more precise when collected near death. A post-mortem motor cortex-based multi-class LDA model was used to classify blood samples in ALS subtypes, and the predicted probability of each sample to belong to a subtype was evaluated using Beta regression models (*Predicted probability ∼ Collection Point * ALS Subtype + Sex + Age of Collection*). (A) Predicted probability for each ALS subtype when the collection point is considered a continuous range [0, 1]. Significant p-values for the effect of disease progression in ALS-Neu (blue) and ALS-SNs (red) are displayed. (B) Predicted probability for each ALS subtype when the collection point is considered a categorical variable (early vs late status). Significant p-values for the effect of disease progression in ALS-Neu (blue) and ALS-SNs are displayed (red). (C) Spearman correlation analysis comparing the collection point (range) against the LDA probability of assignment. Light blue background indicates a confidence interval of 95%. *R* and p-values of the correlation are displayed. (D). DGE of the mean scaled expression values of the features (53 candidate genes) used to build the LDA model in early vs late blood samples. For A, C, and D, *left*: ALS-Neu, *center*: ALS-OxA, *right*: ALS-SNs.

## Discussion

In this study, we demonstrate that ALS molecular subtypes derived from post-mortem motor cortex transcriptomic data can be robustly reproduced across independent cohorts. The three subtypes, ALS-Neu, ALS-OxA and ALS-SNs, were consistently recovered following batch correction and combined unsupervised clustering analysis, with consistency in terms of representative genes and molecular signatures. These findings support the robustness of the previously described gene expression-based ALS subtypes in the brain and highlight the importance of standardized methodology when defining disease subtypes.

ALS-Neu demonstrated to be the most consistent subtype out of the three, presenting the highest number of overlapping genes and biological pathways when compared with previously reported clustering analysis ^7^ (Fig. 2); moreover, when comparing the assignment of samples with previous results from Marriot *et al*. and Eshima, *et al*., more than 90% of the samples remained in the same cluster for the KCL BrainBank dataset, while 70% of samples were conserved in the ALS Consortium (Supplementary Fig 3), regardless the fact that Eshima, *et al*., added TE data in the original analysis that led the clustering to identify an ALS-TE subtype that is not present in our results ^8^. The role of neuroinflammation has been greatly described in ALS, with microglia driving the inflammation and astrocytes gaining toxic properties, although this varies depending on disease status ^37,38^. Our results suggest that this inflammatory signature is not only a hallmark of ALS pathology but also one of the most reproducible molecular signals across cohorts.

We also identified a subtype enriched for ECM and muscle organisation processes (Fig 2). These pathways were previously reported ^7,8^ for the ALS-OxA subtype, although this cluster was mainly described by its oxidative stress signatures. The ECM and skeletal muscle pathways are deeply involved in disease progression, often acting as active drivers of degeneration. Aberrant ECM remodelling leads to tissue fibrosis and stiffness, while increased atrophic signalling and mitochondrial dysfunction have been observed in muscle-related processes, with some of these disturbances driven by oxidative stress ^39,40^. This, together with the observed gene overlap (Fig 2), supports the presence of a reproducible molecular cluster across analyses, even when oxidative stress signals were absent. The ALS-SNs subtype has been previously reported ^7,11^ and is associated with neurotransmission, postsynaptic and neuropeptide signalling. In the ALS Consortium dataset, where this subtype wasn’t reported before, samples assigned to this subtype were previously clustered in the ALS-OxA subtype, indicating a possible overlap between molecular signatures in these two clusters (Supplementary Fig. 3).

Several mutations have been associated with the development of ALS, including the *C9orf72* expansion and genetic variants in *SOD1* ^41^, and although the pathogenesis in patients carrying these mutations is thought that these mutations involve similar molecular mechanisms, the clinical presentation and disease progression are highly heterogeneous ^42–44^. In our datasets, only a small number of samples carried these mutations, with these samples distributed across the different subtypes (Supplementary Table 3). This further supports the heterogeneous nature of these genetic forms of ALS and suggests that the subtype differences identified in our analyses are not primarily driven by these mutations.

ML classifiers confirmed that these subtypes are not only biologically meaningful but also predictable (Fig. 3). Supervised ML models of transcriptomic data have been used in the past to classify ALS patients from controls, and predict progression and survival ^45,46^. In addition, clinical data have been used to predict ALS subtypes ^6,47^. In our analysis, random forest models achieved high accuracy and reproducibility across datasets, with feature selection identifying strong subtype-specific genes. Notably, genes such as *TREM2*, *ALOX5AP* and *FCGR3A* for ALS-Neu, *LAMA5*, *ITGA10* and *ADAMTS10* for ALS-OxA, and *GRM1*, *GABRA1* and *SYT4* for ALS-SNs capture key aspects of the underlying biology, including neuroinflammation, ECM remodelling and synaptic signalling, respectively (Fig. 4); with some of these genes being previously associated with ALS, such as *TREM2* ^48^, link with ALS progression; and *GRM1*, with knockout mouse models for this gene showing an increase of survival ^49^.

The fact that many of these genes are present both in the brain and blood supports their potential as biomarkers of the disease. When extending the post-mortem motor cortex classifiers to peripheral blood, we observed a partial overlap between brain and blood molecular signatures. In particular, the ALS-SNs subtype showed strong consistency, with enrichment of synaptic pathways in blood mirroring those observed in the motor cortex (Fig. 5). Grima *et al*. ^12^ identified four ALS subtypes in blood, most of which were associated with immune response and RNA processing pathways. This may reflect, at least in part, differences in the clustering and enrichment strategies used between studies. Applying comparable methodologies could help resolve subtype structure more consistently and may reveal signatures that more closely align between brain and blood. ALS-OxA and ALS-Neu showed weaker overlap, suggesting that some processes related to ECM and neuroinflammation may be less detectable in peripheral blood or manifest differently outside the central nervous system ^50^. Nonetheless, the ability of the models to identify overlapping subtypes and associated signatures across tissues supports the presence of shared molecular mechanisms between the brain and blood and highlights the potential utility of peripheral biomarkers and ML classifiers for ALS patient stratification.

A key finding of this study is the presence of a longitudinal effect on blood-based subtype prediction. We showed that the probability of correctly classifying ALS-SNs samples increases in samples collected near death (Fig. 6). This likely shows blood and brain signatures converging late in the disease, possibly because of more systemic involvement or stronger neuronal dysfunction signals. On the other hand, ALS-Neu and ALS-OxA subtypes exhibited reduced predictability over time (Fig. 6), which may be driven by a decline in candidate gene expression in blood or by a disparity in immune responses between tissues. These findings highlight the dynamic and independent nature of ALS molecular signatures across disease development and suggest that disease stage is a critical factor influencing our ability to detect subtypes that requires further investigation.

Several limitations of this study should be considered. The study relies on a limited number of available datasets, particularly for blood samples, which may constrain the generalizability of the findings. In addition, these analyses focused on the validation of three previously reported subtypes; however, some of the results indicate some level of overlapping between ALS-OxA and ALS-SNs; clustering analysis considering only two subtypes can explore the similarities between these samples and their molecular signatures. The integration of additional independent cohorts, including longitudinal datasets from both brain and blood, would greatly enhance the robustness of subtype identification and improve model performance. Furthermore, expanding analyses to include diverse populations, earlier disease stages and additional omics layers could provide a more comprehensive view of ALS heterogeneity. Overcoming these limitations is crucial for converting subtype-based stratification into practical clinical tools.

## Acknowledgements

We gratefully acknowledge the ALS Consortium for providing access to the motor cortex transcriptomic dataset and the investigators of the Grima blood RNA-seq cohort for making their data publicly available. We also thank King’s College London and the MRC London Neurodegenerative Diseases Brain Bank for access to post-mortem brain tissue samples and associated data. We would like to thank Professor Michael Benatar from the University of Miami for the invaluable scientific discussions. We are especially grateful to all patients and their families whose contributions made this research possible.

## References

1. Longinetti, E. & Fang, F. Epidemiology of amyotrophic lateral sclerosis: an update of recent literature. Curr. Opin. Neurol. 32, 771–776 (2019).

2. Brown, R. H. & Al-Chalabi, A. Amyotrophic Lateral Sclerosis. N. Engl. J. Med. 377, 162–172 (2017).

3. Couratier, P., Lautrette, G., Luna, J. A. & Corcia, P. Phenotypic variability in amyotrophic lateral sclerosis. Rev. Neurol. (Paris) 177, 536–543 (2021).

4. Jammal, J. K., Gomez, E. A., Al-Chalabi, A. & Iacoangeli, A. Classification of ALS molecular subtypes: a literature review on machine learning applications and their clinical value. BMC Med. 24, 183 (2026).

5. Theunissen, F. et al. Entering the era of precision medicine to treat amyotrophic lateral sclerosis. Mol. Neurodegener. 20, 111 (2025).

6. Spargo, T. P. et al. Unsupervised machine-learning identifies clinically distinct subtypes of ALS that reflect different genetic architectures and biological mechanisms. 2023.06.12.23291304 Preprint at 10.1101/2023.06.12.23291304 (2023).

7. Marriott, H. et al. Unsupervised machine learning identifies distinct ALS molecular subtypes in post-mortem motor cortex and blood expression data. Acta Neuropathol. Commun. 11, 208 (2023).

8. Eshima, J. et al. Molecular subtypes of ALS are associated with differences in patient prognosis. Nat. Commun. 14, 95 (2023).

9. Tam, O. H. et al. Postmortem Cortex Samples Identify Distinct Molecular Subtypes of ALS: Retrotransposon Activation, Oxidative Stress, and Activated Glia. Cell Rep. 29, 1164–1177.e5 (2019).

10. O’Neill, K. et al. ALS molecular subtypes are a combination of cellular and pathological features learned by deep multiomics classifiers. Cell Rep. 44, 115402 (2025).

11. Pasternack, N., Paulsen, O. & Nath, A. Machine learning predicts distinct biotypes of amyotrophic lateral sclerosis. Eur. J. Hum. Genet. 33, 1290–1299 (2025).

12. Grima, N. et al. RNA sequencing of peripheral blood in amyotrophic lateral sclerosis reveals distinct molecular subtypes: Considerations for biomarker discovery. Neuropathol. Appl. Neurobiol. 49, e12943 (2023).

13. Dergai, O. et al. Skeletal Muscle Biomarkers of Amyotrophic Lateral Sclerosis: A Large-Scale, Multi-Cohort Proteomic Study. Ann. Neurol. 99, 393–407 (2026).

14. Iacoangeli, A. et al. SCFD1 expression quantitative trait loci in amyotrophic lateral sclerosis are differentially expressed. Brain Commun. 3, fcab236 (2021).

15. Jones, A. R. et al. A HML6 endogenous retrovirus on chromosome 3 is upregulated in amyotrophic lateral sclerosis motor cortex. Sci. Rep. 11, 14283 (2021).

16. Conlon, E. G. et al. Unexpected similarities between C9ORF72 and sporadic forms of ALS/FTD suggest a common disease mechanism. eLife 7, e37754 (2018).

17. R Core Team. R: A Language and Environment for Statistical Computing. (2025).

18. Love, M. I., Huber, W. & Anders, S. Moderated estimation of fold change and dispersion for RNA-seq data with DESeq2. Genome Biol. 15, 550 (2014).

19. Leek, J. T., Johnson, W. E., Parker, H. S., Jaffe, A. E. & Storey, J. D. The sva package for removing batch effects and other unwanted variation in high-throughput experiments. Bioinformatics 28, 882–883 (2012).

20. Gaujoux, R. & Seoighe, C. A flexible R package for nonnegative matrix factorization. BMC Bioinformatics 11, 367 (2010).

21. Yu, G., Wang, L.-G., Han, Y. & He, Q.-Y. clusterProfiler: an R Package for Comparing Biological Themes Among Gene Clusters. OMICS J. Integr. Biol. 16, 284–287 (2012).

22. Yu, G. & He, Q.-Y. ReactomePA: an R/Bioconductor package for reactome pathway analysis and visualization. Mol. Biosyst. 12, 477–479 (2016).

23. Zou, H. & Hastie, T. Regularization and Variable Selection Via the Elastic Net. J. R. Stat. Soc. Ser. B Stat. Methodol. 67, 301–320 (2005).

24. Kuhn, M. Building Predictive Models in R Using the caret Package. J. Stat. Softw. 28, 1–26 (2008).

25. Denisko, D. & Hoffman, M. M. Classification and interaction in random forests. Proc. Natl. Acad. Sci. U. S. A. 115, 1690–1692 (2018).

26. Liaw, A. & Wiener, M. Classification and Regression by randomForest.

27. Rizkallah, L. W. Enhancing the performance of gradient boosting trees on regression problems. J. Big Data 12, 35 (2025).

28. Chen, T. & Guestrin, C. XGBoost: A Scalable Tree Boosting System. In Proceedings of the 22nd ACM SIGKDD International Conference on Knowledge Discovery and Data Mining 785–794 (Association for Computing Machinery, New York, NY, USA, 2016). doi:10.1145/2939672.2939785.

29. Robin, X. et al. pROC: an open-source package for R and S+ to analyze and compare ROC curves. BMC Bioinformatics 12, 77 (2011).

30. Gardner-Lubbe, S. Linear discriminant analysis for multiple functional data analysis. J. Appl. Stat. 48, 1917 (2020).

31. Thul, P. J. & Lindskog, C. The human protein atlas: A spatial map of the human proteome. Protein Sci. Publ. Protein Soc. 27, 233–244 (2018).

32. Langfelder, P. & Horvath, S. WGCNA: an R package for weighted correlation network analysis. BMC Bioinformatics 9, 559 (2008).

33. Lewis, M. J. et al. Molecular Portraits of Early Rheumatoid Arthritis Identify Clinical and Treatment Response Phenotypes. Cell Rep. 28, 2455–2470.e5 (2019).

34. Cribari-Neto, F. & Zeileis, A. Beta Regression in R. J. Stat. Softw. 34, 1–24 (2010).

35. Lüdecke, D. ggeffects: Tidy Data Frames of Marginal Effects from Regression Models. J. Open Source Softw. 3, 772 (2018).

36. Zhang, Y., Parmigiani, G. & Johnson, W. E. ComBat-seq: batch effect adjustment for RNA-seq count data. NAR Genomics Bioinforma. 2, lqaa078 (2020).

37. Komine, O. & Yamanaka, K. Neuroinflammation in motor neuron disease. Nagoya J. Med. Sci. 77, 537–549 (2015).

38. Murdock, B. J. et al. Increased ratio of circulating neutrophils to monocytes in amyotrophic lateral sclerosis. Neurol. Neuroimmunol. Neuroinflammation 3, e242 (2016).

39. Apolloni, S., Tortoriello, S., Milani, M. & Rossi, S. Extracellular Matrix Remodeling in Motor Neuron Diseases. Int. J. Mol. Sci. 26, 11376 (2025).

40. Pikatza-Menoio, O. et al. The Skeletal Muscle Emerges as a New Disease Target in Amyotrophic Lateral Sclerosis. J. Pers. Med. 11, 671 (2021).

41. Nijs, M. & Van Damme, P. The genetics of amyotrophic lateral sclerosis. Curr. Opin. Neurol. 37, 560–569 (2024).

42. Hao, Z., Wang, R., Ren, H. & Wang, G. Role of the C9ORF72 Gene in the Pathogenesis of Amyotrophic Lateral Sclerosis and Frontotemporal Dementia. Neurosci. Bull. 36, 1057–1070 (2020).

43. Opie-Martin, S. et al. The SOD1-mediated ALS phenotype shows a decoupling between age of symptom onset and disease duration. Nat. Commun. 13, 6901 (2022).

44. Caldi Gomes, L., et al. Multiomic ALS signatures highlight subclusters and sex differences suggesting the MAPK pathway as therapeutic target. Nat. Commun. 15, 4893 (2024).

45. Bain, C., Ramamoorthy, D. & Fraenkel, E. Using Supervised Machine Learning Methods to Create a Gene-Based ALS Predictor from Postmortem Transcriptomics Data. (2023).

46. Zhao, Y. et al. Gene expression signatures from whole blood predict amyotrophic lateral sclerosis case status and survival. Nat. Commun. 16, 9631 (2025).

47. Faghri, F. et al. Identifying and predicting amyotrophic lateral sclerosis subgroups: a population-based machine learning study. Lancet Digit. Health 4, e359–e369 (2022).

48. Xie, M., Zhao, S., Bosco, D. B., Nguyen, A. & Wu, L.-J. Microglial TREM2 in amyotrophic lateral sclerosis. Dev. Neurobiol. 82, 125–137 (2022).

49. Milanese, M. et al. Knocking down metabotropic glutamate receptor 1 improves survival and disease progression in the SOD1(G93A) mouse model of amyotrophic lateral sclerosis. Neurobiol. Dis. 64, 48–59 (2014).

50. Cao, M. C. et al. Serum biomarkers of neuroinflammation and blood-brain barrier leakage in amyotrophic lateral sclerosis. BMC Neurol. 22, 216 (2022).

