## Supplemental information for "Machine learning evaluation of gene expression-based ALS subtypes across brain and blood tissues"

**This file includes:**

- Supplemental Figures
- Supplemental Tables
- Supplemental NYGC ALS Consortium authors

**Supplemental Figures**

*Supplemental Figure 1*

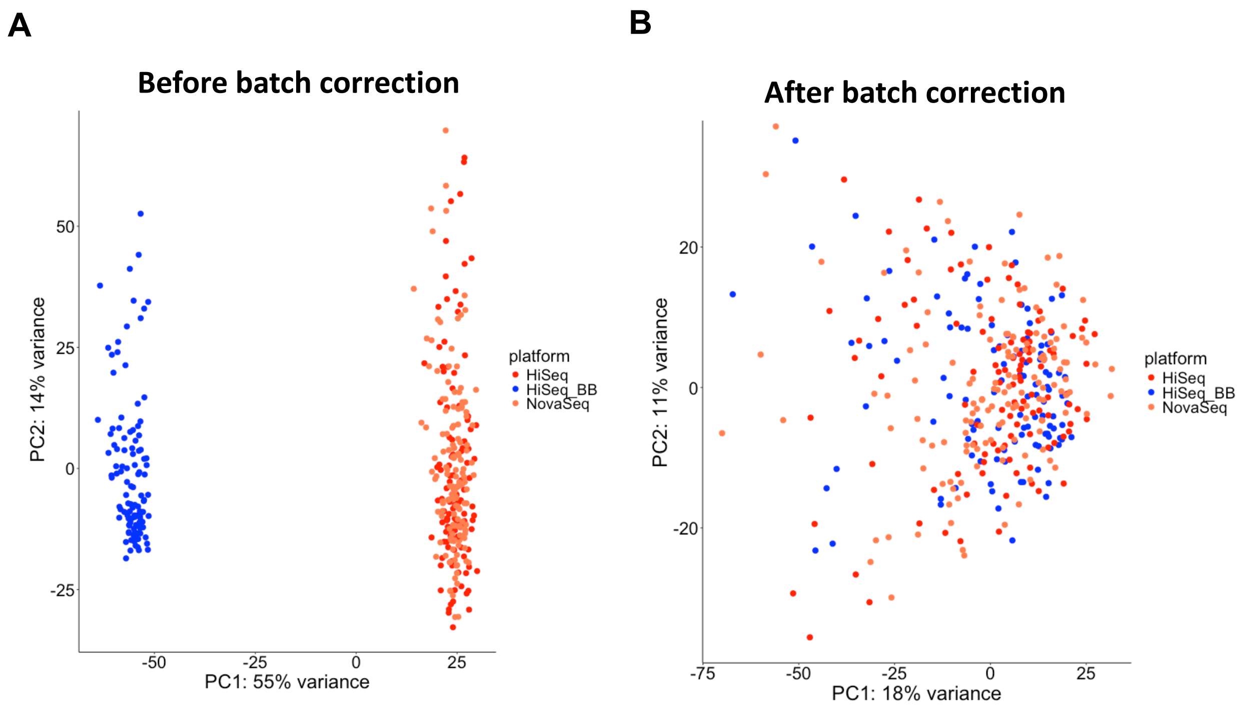

**Suppl Figure 1. ComBat can performed batch correction between datasets.** (A) PCA analysis before batch correction of samples from KCL BrainBank (HiSeq_BB) and ALS Consortium (HiSeq and NovaSeq). (B) PCA analysis after batch correction for KCL BrainBank and ALS Consortium.

*Supplemental Figure 2*

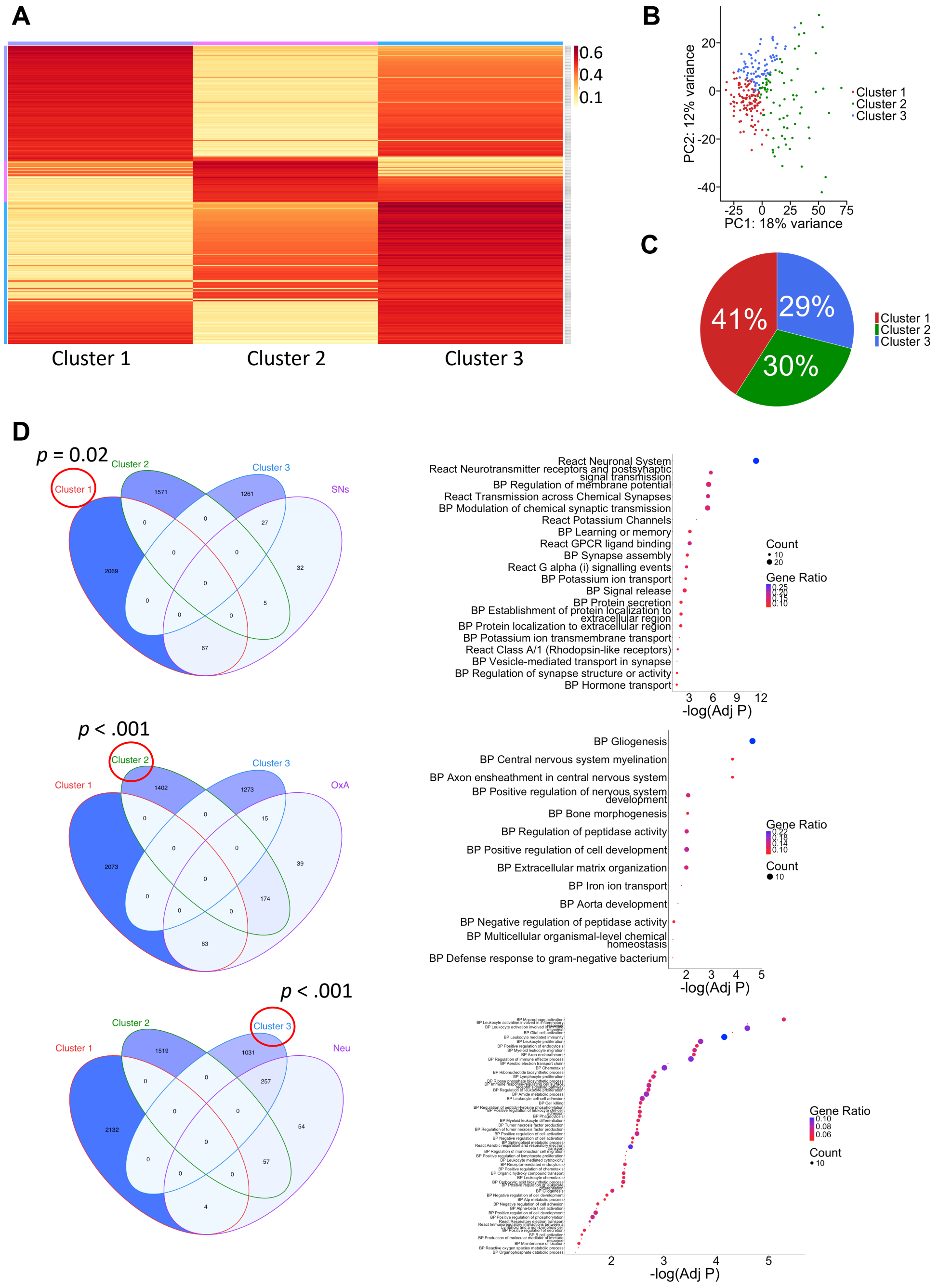

**Suppl Figure 2. Unsupervised clustering analysis of post-mortem motor cortex transcriptomes from the ALS Consortium dataset.** Batch corrected-transcriptomic data of 257 post-mortem motor cortex samples from the ALS Consortium were used to perform nsNMF-based unsupervised clustering on the top 5,000 most variable genes. (A) Heatmap showing 3 clusters based on the top 5,000 most variable genes. (B) PCA plot showing samples by ALS subtype. (C) Distribution of samples for each ALS subtype. (D) Venn Diagrams (*left;* p-values for Fisher's exact test are displayed) and enrichment pathway analysis (GO:BP terms and Reactome database; *right*) for the three ALS subtypes (*top:* ALS-SNs, *middle:* ALS-OxA and *bottom:* ALS-Neu; p-values for Fisher's exact test after BH correction are displayed).

*Supplemental Figure 3*

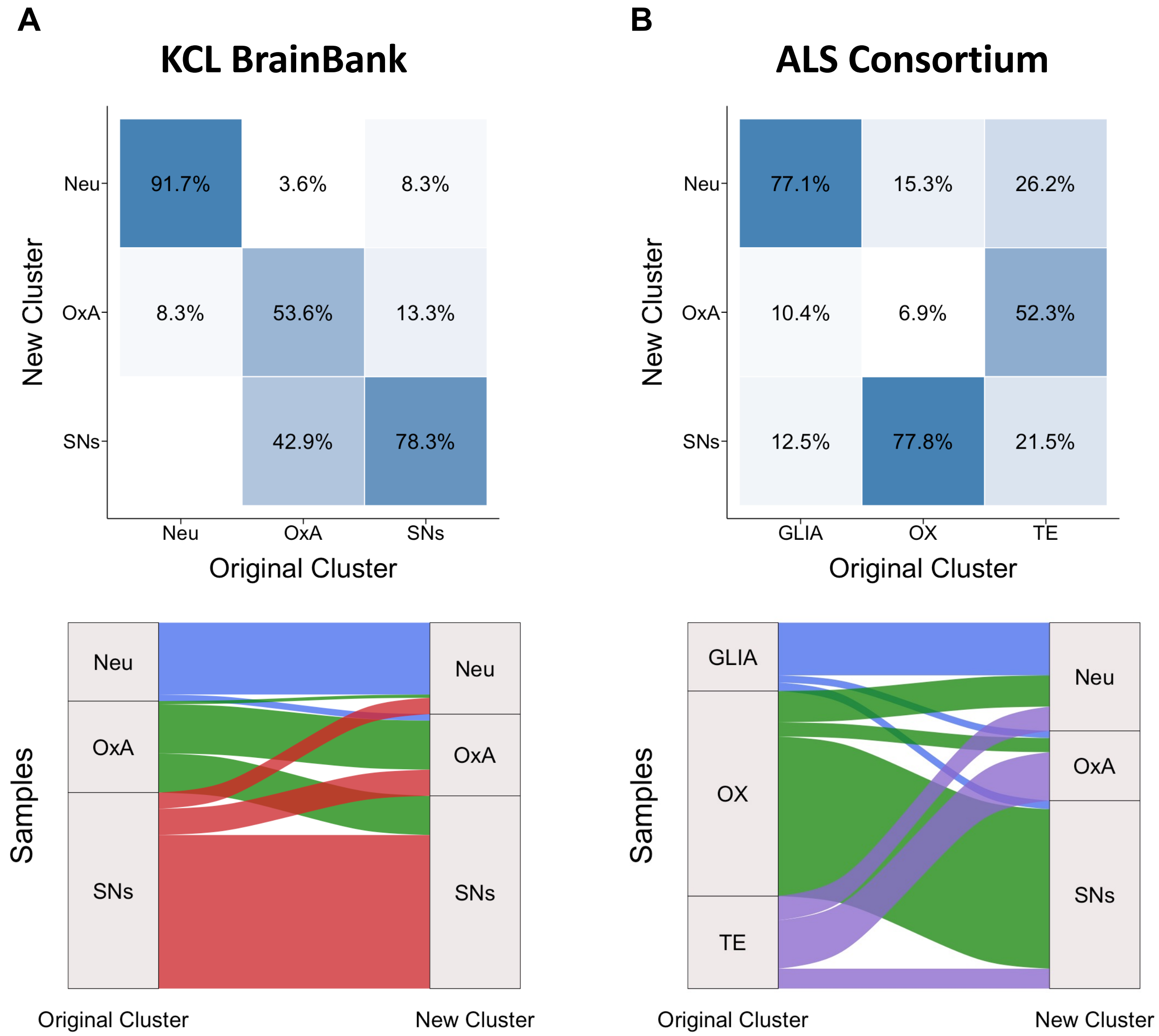

**Suppl Figure 3. Distribution of samples across ALS subtypes before and after combined unsupervised clustering.** Transcriptomic data of 369 post-mortem motor cortex samples from the KCL BrainBank and ALS Consortium were combined after batch correction, and nsNMF-based unsupervised clustering was performed on the top 5,000 most variable genes. Distribution of samples from KCL Brain Bank (A) and ALS Consortium (B) before and after the new unsupervised clustering. *Top*: Heatmap showing the percentage of overlapping samples between the new and original clusters. The x-axis refers to the original cluster (A: ALS-Neu, ALS-OxA and ALS-SNs; B: GLIA, OX and TE), and the y-axis to the new one (ALS-Neu, ALS-OxA and ALS-SNs). *Bottom*: Sankey plot depicting the movement of the samples from the original clusters (A: ALS-Neu, ALS-OxA and ALS-SNs; B: GLIA, OX and TE) to the new ones (ALS-Neu, ALS-OxA and ALS-SNs).

*Supplemental Figure 4*

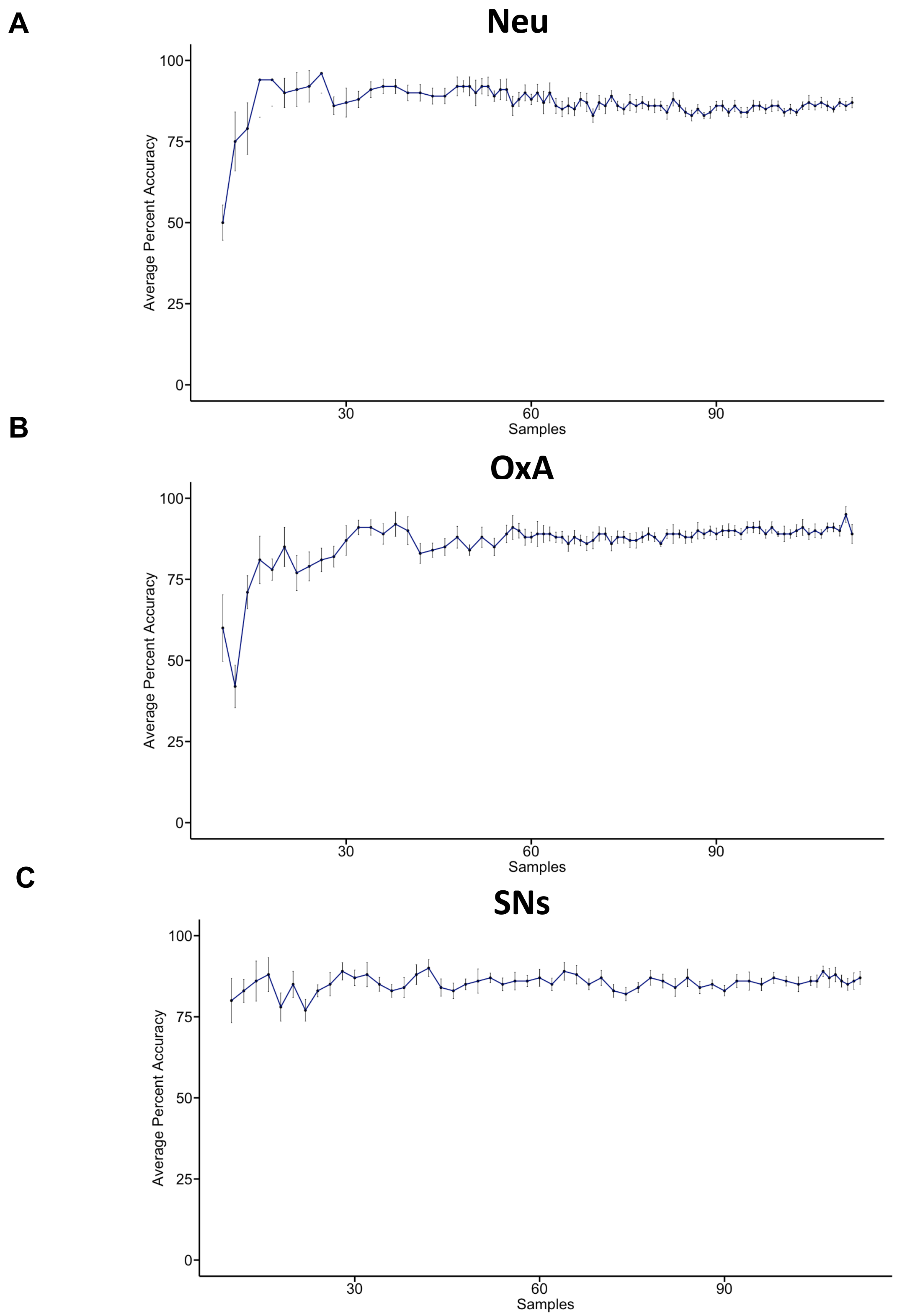

**Suppl Figure 4. Learning curves of machine learning models for the prediction of ALS subtypes.** Learning curves were created by adding one case and one non-case sample at the time (“one-vs-the rest” approach) using the random forest methodology for (A) ALS-Neu, (B) ALS-OxA and (C) ALS-SNs.

*Supplemental Figure 5*

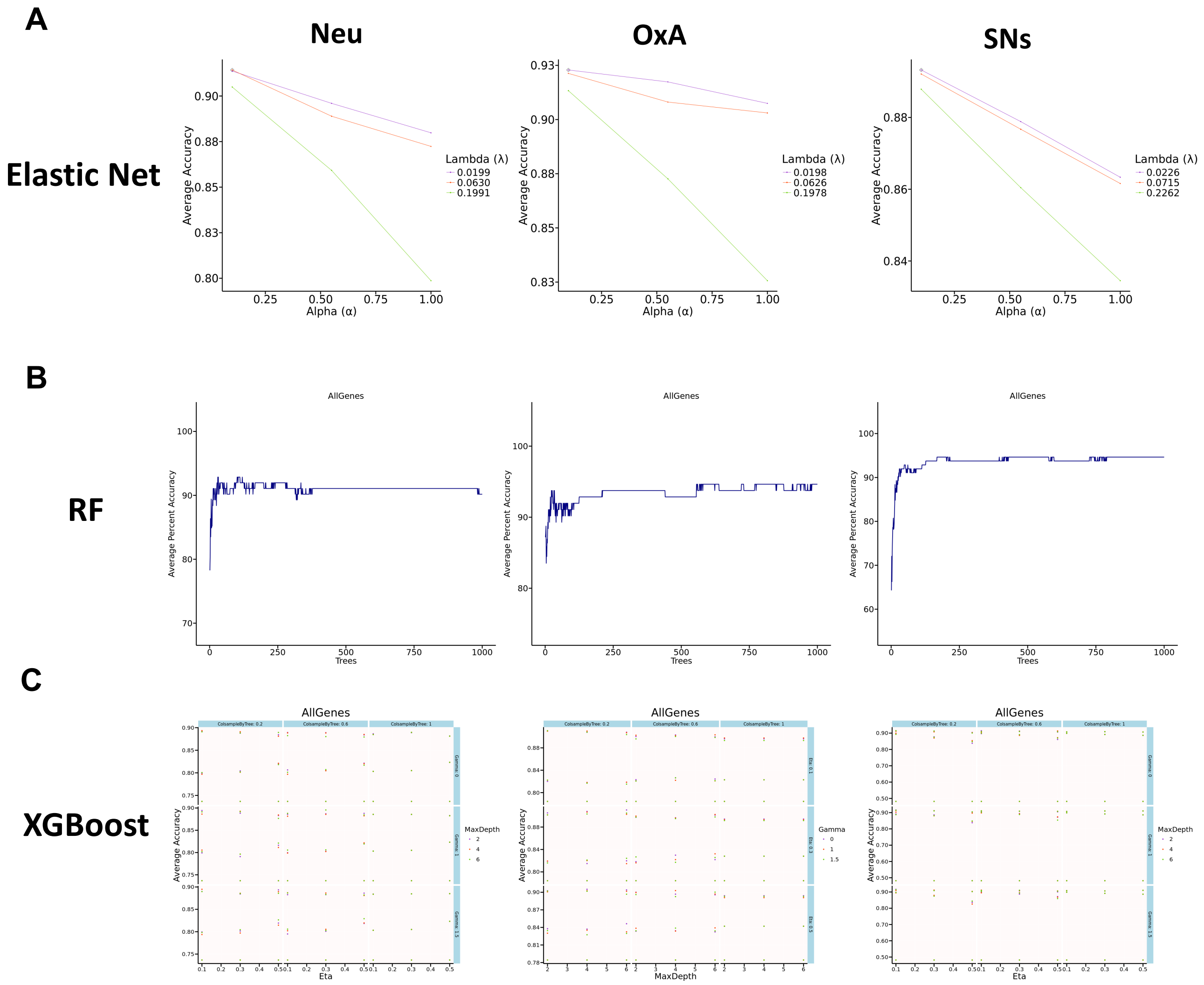

**Suppl Figure 5. Parameter tunning for ML for ALS subtype classification.** Parameter tuning for the Elastic Net (A), random forest (RF; B) and XGBoost (C) models. *Left*: ALS-Neu, *center*: ALS-OxA and *right*: ALS-SNs.

*Supplemental Figure 6*

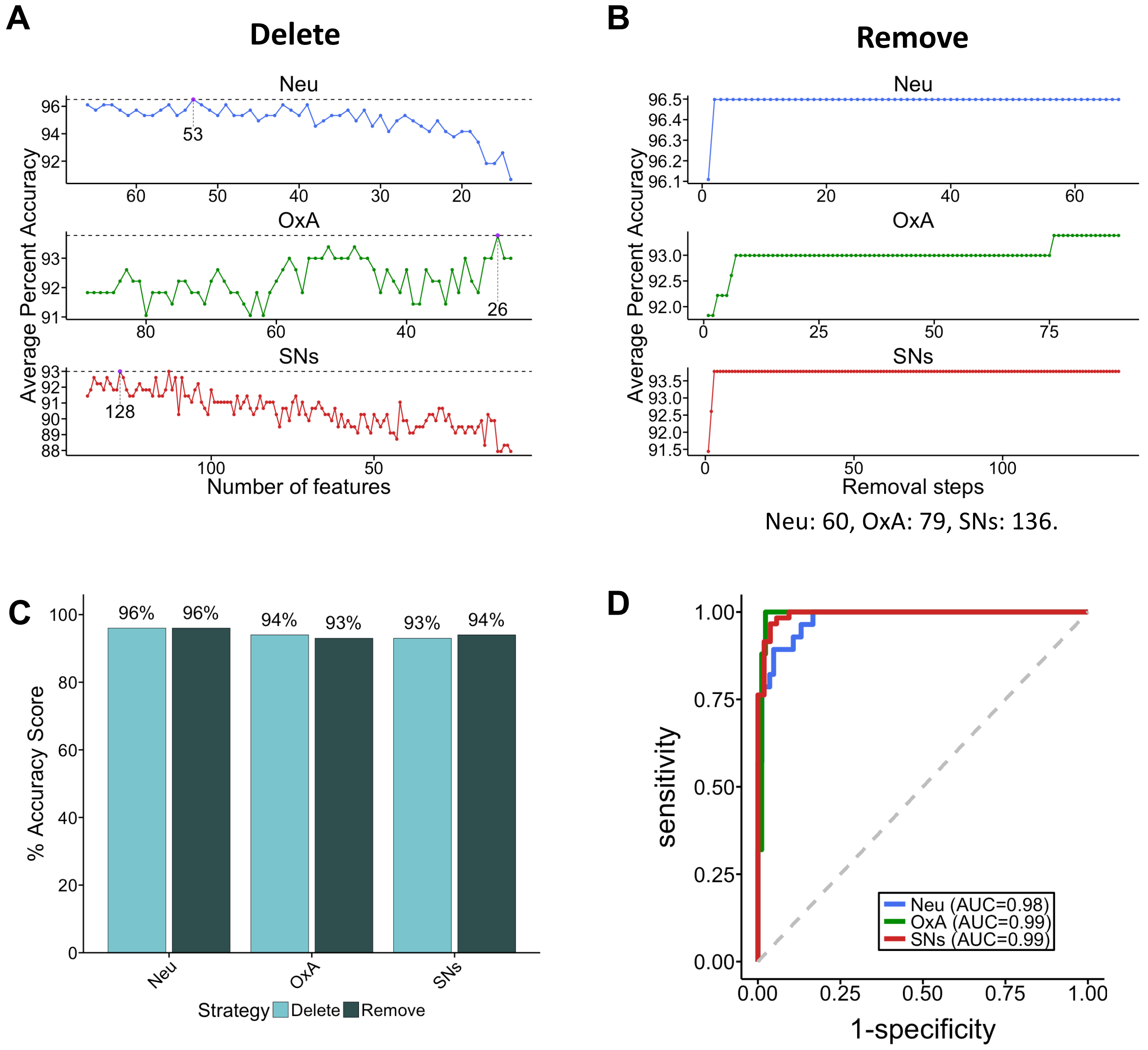

**Suppl Figure 6. Machine learning model improvement strategies.** Two strategies were used for improvement of the classifiers using ALS Consortium as training dataset: *Delete* (A), which creates ML models progressively eliminating genes until identifies the minimum number of genes with high accuracy, and *remove* (B), which builds models eliminating genes that reduce their accuracy *(Bottom*: final number of genes for each ALS subtypes). (C) Percent accuracy score of improved prediction models based on the *delete* and *remove* strategies. (D) ROC curves and AUC values for predictive models when evaluated in an independent cohort (KCL BrainBank).

*Supplemental Figure 7*

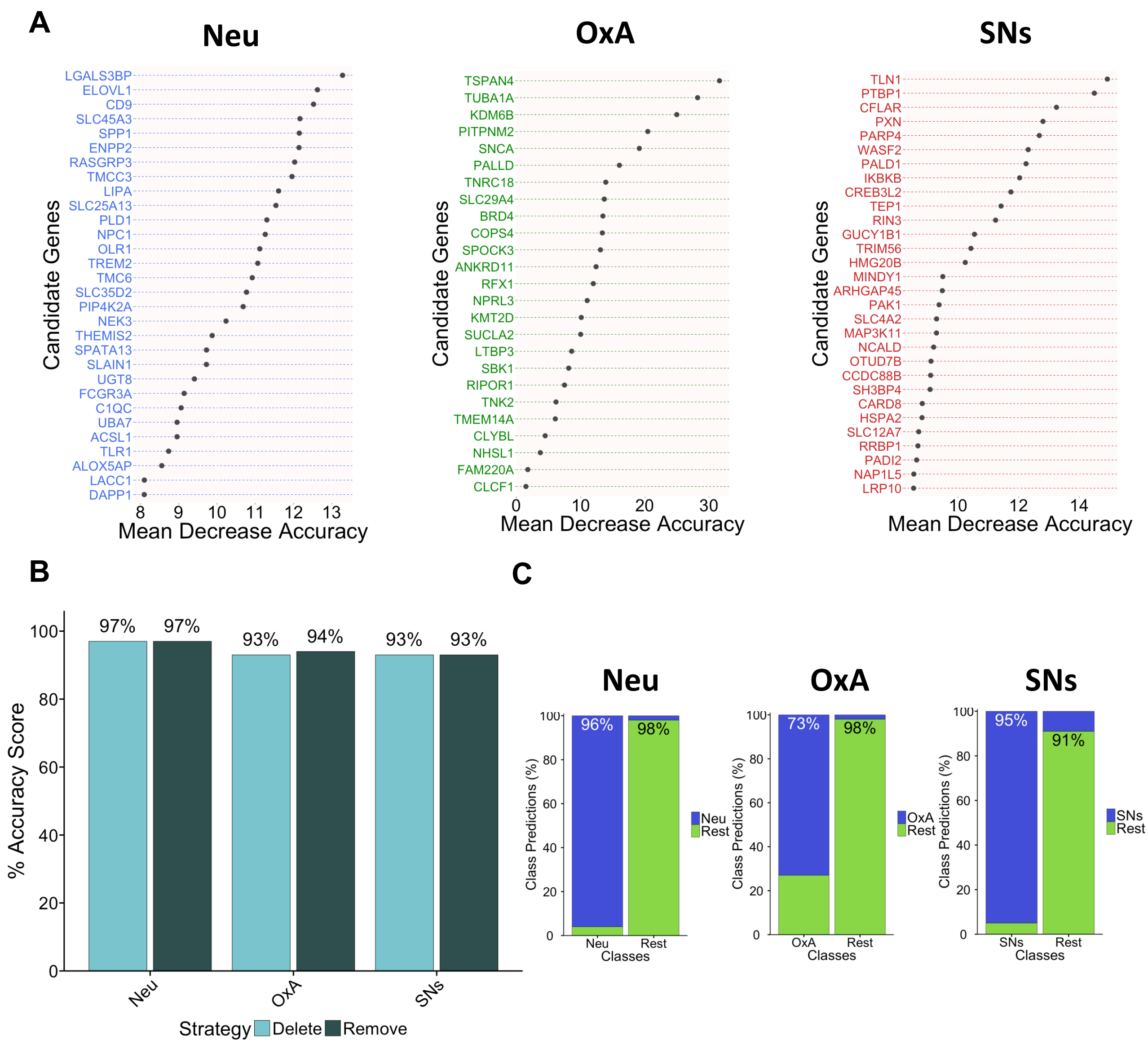

**Suppl Figure 7. Prediction accuracy of the models is not affected when feature selection is focused in genes expressed in blood.** Feature selection was performed based in the random forest methodology ("importance" function), focusing only on candidate genes expressed in blood, using the *remove* and *delete* strategies and the ALS Consortium (257 post-mortem motor cortex samples) as the training dataset. (A) Top 30 most relevant candidate genes expressed in blood for the prediction performance of the ALS-Neu (*left*), ALS-OxA (*center*) and ALS-SNs (*right*) classifiers based on decreasing accuracy of the *remove* models. (B) Percent accuracy score of improved prediction models based on the *delete* and *remove* strategies focusing on genes expressed in blood. (C) Classification predictions for each class (sensitivity and specificity) of the *remove* model using blood biomarkers (*left*: ALS-Neu, *center*: ALS-OxA, *right*: ALS-SNs*)*. Blue indicates the samples were predicted as the specific subtype while green indicates those patients don't belong to that cluster. Percentages indicate true positives and true negatives.

*Supplemental Figure 8*

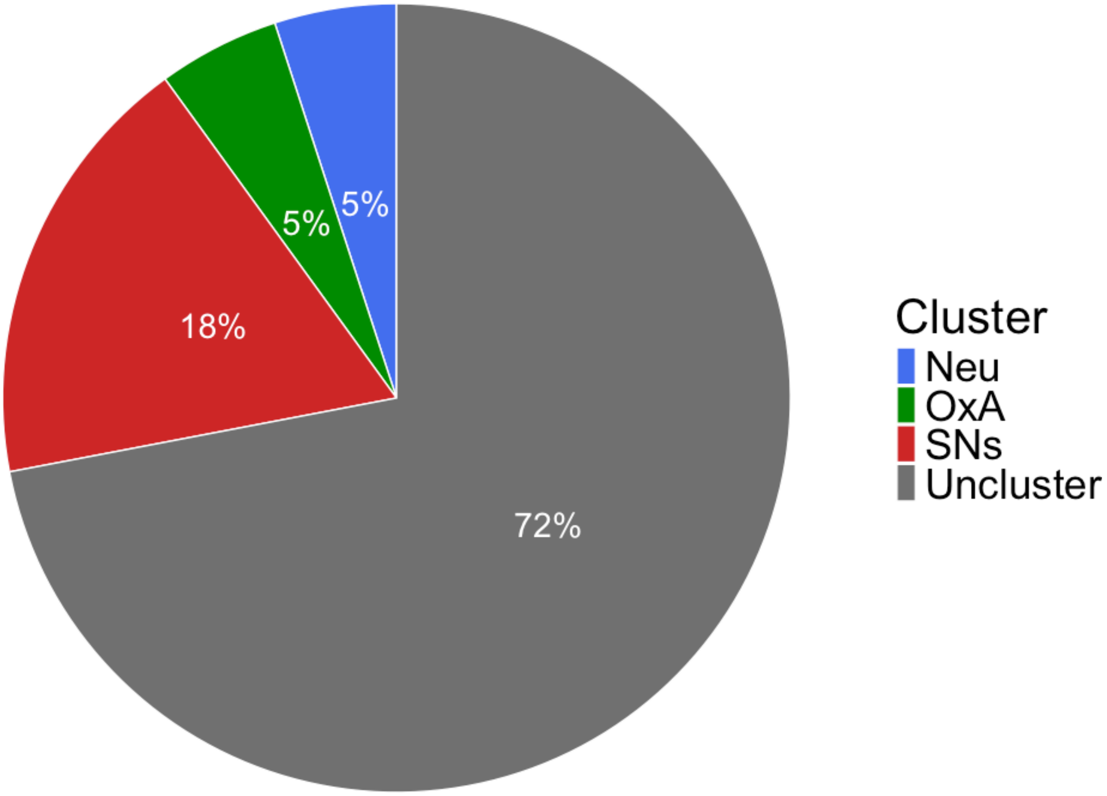

**Suppl Figure 8. Random forest binary classifiers can classify 18% of the blood samples in the ALS-SNs subtype.** Binary classifiers trained using post-mortem motor cortex samples (ALS Consortium as the training dataset) were evaluated in blood samples (Grima dataset). Pie chart shows the distribution of samples for each ALS subtype when applying the random forest models for each subtype. Uncluster samples refers to samples that weren't classified in neither of the ALS subtypes.

*Supplemental Figure 9*

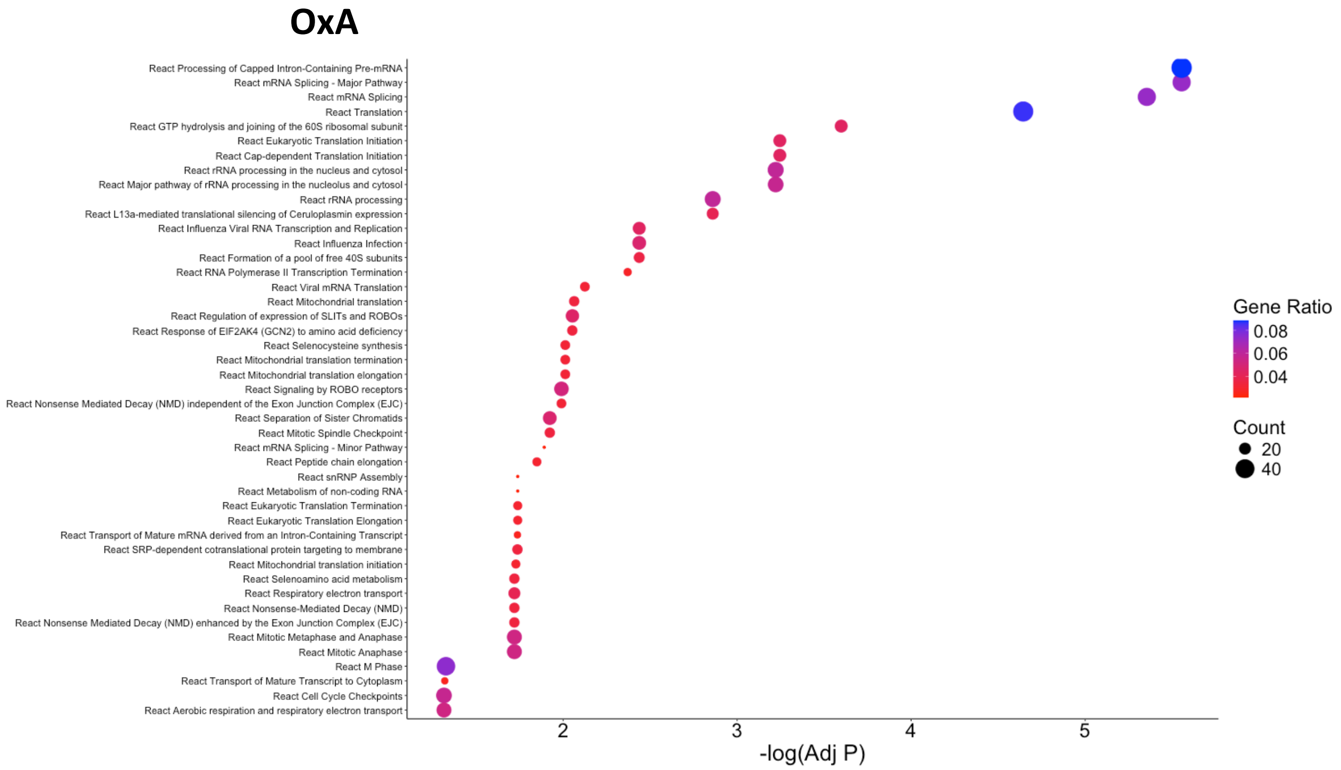

**Suppl Figure 9. Enriched pathway analysis of ALS-OxA after DGE analysis of LDA samples.** WGCNA was applied to identify representative genes correlated with others and genes without co-expression and multi-class LDA models were created using ALS Consortium (257 post-mortem motor cortex samples) as the training dataset and evaluated in blood samples (Grima dataset). Three-way DGE analysis were performed based on blood sample classification and significantly expressed genes for each ALS subtype were used for enriched pathway analysis. Enriched pathway analysis (Reactome database) for the ALS-OxA is shown.

### Supplemental Tables

**Suppl Table 1.** Associated genes for cluster identification.

| Ensembl ID | Gene Symbol | Cluster | Reference ALS-Subtype |
|---|---|---|---|
| ENSG00000000938 | FGR | Cluster 1 | ALS-OxA |
| ENSG00000000971 | CFH | Cluster 1 | ALS-OxA |
| ENSG00000003436 | TFPI | Cluster 1 | ALS-OxA |
| ENSG00000011422 | PLAUR | Cluster 1 | ALS-OxA |
| ENSG00000013588 | GPRC5A | Cluster 1 | ALS-OxA |
| ENSG00000015520 | NPC1L1 | Cluster 1 | ALS-OxA |
| ENSG00000047617 | ANO2 | Cluster 1 | ALS-OxA |
| ENSG00000054598 | FOXC1 | Cluster 1 | ALS-OxA |
| ENSG00000060138 | YBX3 | Cluster 1 | ALS-OxA |
| ENSG00000073792 | IGF2BP2 | Cluster 1 | ALS-OxA |
| ENSG00000074047 | GLI2 | Cluster 1 | ALS-OxA |
| ENSG00000074219 | TEAD2 | Cluster 1 | ALS-OxA |
| ENSG00000077238 | IL4R | Cluster 1 | ALS-OxA |
| ENSG00000088882 | CPXM1 | Cluster 1 | ALS-OxA |
| ENSG00000090339 | ICAM1 | Cluster 1 | ALS-OxA |
| ENSG00000094963 | FMO2 | Cluster 1 | ALS-OxA |
| ENSG00000096696 | DSP | Cluster 1 | ALS-OxA |
| ENSG00000099994 | SUSD2 | Cluster 1 | ALS-OxA |
| ENSG00000100336 | APOL4 | Cluster 1 | ALS-OxA |
| ENSG00000100504 | PYGL | Cluster 1 | ALS-OxA |
| ENSG00000101187 | SLCO4A1 | Cluster 1 | ALS-OxA |
| ENSG00000101335 | MYL9 | Cluster 1 | ALS-OxA |
| ENSG00000105641 | SLC5A5 | Cluster 1 | ALS-OxA |
| ENSG00000106211 | HSPB1 | Cluster 1 | ALS-OxA |
| ENSG00000106366 | SERPINE1 | Cluster 1 | ALS-OxA |
| ENSG00000106624 | AEBP1 | Cluster 1 | ALS-OxA |
| ENSG00000107438 | PDLIM1 | Cluster 1 | ALS-OxA |
| ENSG00000107796 | ACTA2 | Cluster 1 | ALS-OxA |
| ENSG00000108821 | COL1A1 | Cluster 1 | ALS-OxA |
| ENSG00000108823 | SGCA | Cluster 1 | ALS-OxA |
| ENSG00000109758 | HGFAC | Cluster 1 | ALS-OxA |
| ENSG00000110719 | TCIRG1 | Cluster 1 | ALS-OxA |
| ENSG00000110852 | CLEC2B | Cluster 1 | ALS-OxA |
| ENSG00000111057 | KRT18 | Cluster 1 | ALS-OxA |
| ENSG00000111341 | MGP | Cluster 1 | ALS-OxA |
| ENSG00000112175 | BMP5 | Cluster 1 | ALS-OxA |
| ENSG00000112214 | FHL5 | Cluster 1 | ALS-OxA |
| ENSG00000112299 | VNN1 | Cluster 1 | ALS-OxA |
| ENSG00000112303 | VNN2 | Cluster 1 | ALS-OxA |
| ENSG00000112837 | TBX18 | Cluster 1 | ALS-OxA |
| ENSG00000112936 | C7 | Cluster 1 | ALS-OxA |
| ENSG00000115590 | IL1R2 | Cluster 1 | ALS-OxA |
| ENSG00000115594 | IL1R1 | Cluster 1 | ALS-OxA |
| ENSG00000115602 | IL1RL1 | Cluster 1 | ALS-OxA |
| ENSG00000115604 | IL18R1 | Cluster 1 | ALS-OxA |
| ENSG00000115607 | IL18RAP | Cluster 1 | ALS-OxA |
| ENSG00000115648 | MLPH | Cluster 1 | ALS-OxA |
| ENSG00000117318 | ID3 | Cluster 1 | ALS-OxA |
| ENSG00000117595 | IRF6 | Cluster 1 | ALS-OxA |
| ENSG00000118271 | TTR | Cluster 1 | ALS-OxA |
| ENSG00000118503 | TNFAIP3 | Cluster 1 | ALS-OxA |
| ENSG00000118729 | CASQ2 | Cluster 1 | ALS-OxA |
| ENSG00000120708 | TGFBI | Cluster 1 | ALS-OxA |
| ENSG00000122176 | FMOD | Cluster 1 | ALS-OxA |
| ENSG00000122862 | SRGN | Cluster 1 | ALS-OxA |
| ENSG00000123374 | CDK2 | Cluster 1 | ALS-OxA |
| ENSG00000124107 | SLPI | Cluster 1 | ALS-OxA |
| ENSG00000124212 | PTGIS | Cluster 1 | ALS-OxA |
| ENSG00000124253 | PCK1 | Cluster 1 | ALS-OxA |
| ENSG00000124762 | CDKN1A | Cluster 1 | ALS-OxA |
| ENSG00000125144 | MT1G | Cluster 1 | ALS-OxA |
| ENSG00000125430 | HS3ST3B1 | Cluster 1 | ALS-OxA |
| ENSG00000125733 | TRIP10 | Cluster 1 | ALS-OxA |

| | | | |
|---|---|---|---|
| ENSG00000125810 | CD93 | Cluster 1 | ALS-OxA |
| ENSG00000126778 | SIX1 | Cluster 1 | ALS-OxA |
| ENSG00000128274 | A4GALT | Cluster 1 | ALS-OxA |
| ENSG00000128917 | DLL4 | Cluster 1 | ALS-OxA |
| ENSG00000129654 | FOXJ1 | Cluster 1 | ALS-OxA |
| ENSG00000130176 | CNN1 | Cluster 1 | ALS-OxA |
| ENSG00000130600 | H19 | Cluster 1 | ALS-OxA |
| ENSG00000130635 | COL5A1 | Cluster 1 | ALS-OxA |
| ENSG00000131471 | AOC3 | Cluster 1 | ALS-OxA |
| ENSG00000132357 | CARD6 | Cluster 1 | ALS-OxA |
| ENSG00000133392 | MYH11 | Cluster 1 | ALS-OxA |
| ENSG00000135245 | HILPDA | Cluster 1 | ALS-OxA |
| ENSG00000135842 | NIBAN1 | Cluster 1 | ALS-OxA |
| ENSG00000137507 | LRRC32 | Cluster 1 | ALS-OxA |
| ENSG00000137801 | THBS1 | Cluster 1 | ALS-OxA |
| ENSG00000137834 | SMAD6 | Cluster 1 | ALS-OxA |
| ENSG00000138061 | CYP1B1 | Cluster 1 | ALS-OxA |
| ENSG00000138722 | MMRN1 | Cluster 1 | ALS-OxA |
| ENSG00000140682 | TGFB1I1 | Cluster 1 | ALS-OxA |
| ENSG00000140839 | CLEC18B | Cluster 1 | ALS-OxA |
| ENSG00000141052 | MYOCD | Cluster 1 | ALS-OxA |
| ENSG00000141574 | SECTM1 | Cluster 1 | ALS-OxA |
| ENSG00000142089 | IFITM3 | Cluster 1 | ALS-OxA |
| ENSG00000142102 | PGGHG | Cluster 1 | ALS-OxA |
| ENSG00000142173 | COL6A2 | Cluster 1 | ALS-OxA |
| ENSG00000142798 | HSPG2 | Cluster 1 | ALS-OxA |
| ENSG00000142973 | CYP4B1 | Cluster 1 | ALS-OxA |
| ENSG00000143546 | S100A8 | Cluster 1 | ALS-OxA |
| ENSG00000143867 | OSR1 | Cluster 1 | ALS-OxA |
| ENSG00000144810 | COL8A1 | Cluster 1 | ALS-OxA |
| ENSG00000144837 | PLA1A | Cluster 1 | ALS-OxA |
| ENSG00000145623 | OSMR | Cluster 1 | ALS-OxA |
| ENSG00000145779 | TNFAIP8 | Cluster 1 | ALS-OxA |
| ENSG00000148604 | RGR | Cluster 1 | ALS-OxA |
| ENSG00000149257 | SERPINH1 | Cluster 1 | ALS-OxA |
| ENSG00000149452 | SLC22A8 | Cluster 1 | ALS-OxA |
| ENSG00000149573 | MPZL2 | Cluster 1 | ALS-OxA |
| ENSG00000149591 | TAGLN | Cluster 1 | ALS-OxA |
| ENSG00000150048 | CLEC1A | Cluster 1 | ALS-OxA |
| ENSG00000151929 | BAG3 | Cluster 1 | ALS-OxA |
| ENSG00000152049 | KCNE4 | Cluster 1 | ALS-OxA |
| ENSG00000152779 | SLC16A12 | Cluster 1 | ALS-OxA |
| ENSG00000153404 | PLEKHG4B | Cluster 1 | ALS-OxA |
| ENSG00000159167 | STC1 | Cluster 1 | ALS-OxA |
| ENSG00000159403 | C1R | Cluster 1 | ALS-OxA |
| ENSG00000160183 | TMPRSS3 | Cluster 1 | ALS-OxA |
| ENSG00000161638 | ITGA5 | Cluster 1 | ALS-OxA |
| ENSG00000162383 | SLC1A7 | Cluster 1 | ALS-OxA |
| ENSG00000162458 | FBLIM1 | Cluster 1 | ALS-OxA |
| ENSG00000162747 | FCGR3B | Cluster 1 | ALS-OxA |
| ENSG00000162998 | FRZB | Cluster 1 | ALS-OxA |
| ENSG00000163017 | ACTG2 | Cluster 1 | ALS-OxA |
| ENSG00000163220 | S100A9 | Cluster 1 | ALS-OxA |
| ENSG00000163395 | IGFN1 | Cluster 1 | ALS-OxA |
| ENSG00000163431 | LMOD1 | Cluster 1 | ALS-OxA |
| ENSG00000163464 | CXCR1 | Cluster 1 | ALS-OxA |
| ENSG00000163638 | ADAMTS9 | Cluster 1 | ALS-OxA |
| ENSG00000163736 | PPBP | Cluster 1 | ALS-OxA |
| ENSG00000163739 | CXCL1 | Cluster 1 | ALS-OxA |
| ENSG00000163817 | SLC6A20 | Cluster 1 | ALS-OxA |
| ENSG00000164294 | GPX8 | Cluster 1 | ALS-OxA |
| ENSG00000164692 | COL1A2 | Cluster 1 | ALS-OxA |
| ENSG00000164707 | SLC13A4 | Cluster 1 | ALS-OxA |
| ENSG00000164761 | TNFRSF11B | Cluster 1 | ALS-OxA |
| ENSG00000164867 | NOS3 | Cluster 1 | ALS-OxA |
| ENSG00000165474 | GJB2 | Cluster 1 | ALS-OxA |
| ENSG00000165507 | DEPP1 | Cluster 1 | ALS-OxA |
| ENSG00000166482 | MFAP4 | Cluster 1 | ALS-OxA |
| ENSG00000166523 | CLEC4E | Cluster 1 | ALS-OxA |
| ENSG00000166592 | RRAD | Cluster 1 | ALS-OxA |
| ENSG00000167244 | IGF2 | Cluster 1 | ALS-OxA |

| | | | |
|---|---|---|---|
| ENSG00000167772 | ANGPTL4 | Cluster 1 | ALS-OxA |
| ENSG00000168404 | MLKL | Cluster 1 | ALS-OxA |
| ENSG00000168427 | KLHL30 | Cluster 1 | ALS-OxA |
| ENSG00000168542 | COL3A1 | Cluster 1 | ALS-OxA |
| ENSG00000169429 | CXCL8 | Cluster 1 | ALS-OxA |
| ENSG00000169894 | MUC3A | Cluster 1 | ALS-OxA |
| ENSG00000169908 | TM4SF1 | Cluster 1 | ALS-OxA |
| ENSG00000170345 | FOS | Cluster 1 | ALS-OxA |
| ENSG00000170577 | SIX2 | Cluster 1 | ALS-OxA |
| ENSG00000170801 | HTRA3 | Cluster 1 | ALS-OxA |
| ENSG00000171049 | FPR2 | Cluster 1 | ALS-OxA |
| ENSG00000171345 | KRT19 | Cluster 1 | ALS-OxA |
| ENSG00000172935 | MRGPRF | Cluster 1 | ALS-OxA |
| ENSG00000173110 | HSPA6 | Cluster 1 | ALS-OxA |
| ENSG00000173421 | IHO1 | Cluster 1 | ALS-OxA |
| ENSG00000173530 | TNFRSF10D | Cluster 1 | ALS-OxA |
| ENSG00000173597 | SULT1B1 | Cluster 1 | ALS-OxA |
| ENSG00000173641 | HSPB7 | Cluster 1 | ALS-OxA |
| ENSG00000173918 | C1QTNF1 | Cluster 1 | ALS-OxA |
| ENSG00000174226 | SNX31 | Cluster 1 | ALS-OxA |
| ENSG00000174348 | PODN | Cluster 1 | ALS-OxA |
| ENSG00000175084 | DES | Cluster 1 | ALS-OxA |
| ENSG00000176692 | FOXC2 | Cluster 1 | ALS-OxA |
| ENSG00000177464 | GPR4 | Cluster 1 | ALS-OxA |
| ENSG00000177575 | CD163 | Cluster 1 | ALS-OxA |
| ENSG00000179023 | KLHDC7A | Cluster 1 | ALS-OxA |
| ENSG00000183508 | TENT5C | Cluster 1 | ALS-OxA |
| ENSG00000183888 | SRARP | Cluster 1 | ALS-OxA |
| ENSG00000184557 | SOCS3 | Cluster 1 | ALS-OxA |
| ENSG00000184811 | TRARG1 | Cluster 1 | ALS-OxA |
| ENSG00000185201 | IFITM2 | Cluster 1 | ALS-OxA |
| ENSG00000185499 | MUC1 | Cluster 1 | ALS-OxA |
| ENSG00000185561 | TLCD2 | Cluster 1 | ALS-OxA |
| ENSG00000185585 | OLFML2A | Cluster 1 | ALS-OxA |
| ENSG00000186431 | FCAR | Cluster 1 | ALS-OxA |
| ENSG00000186564 | FOXD2 | Cluster 1 | ALS-OxA |
| ENSG00000187479 | C11orf96 | Cluster 1 | ALS-OxA |
| ENSG00000187498 | COL4A1 | Cluster 1 | ALS-OxA |
| ENSG00000187513 | GJA4 | Cluster 1 | ALS-OxA |
| ENSG00000187922 | LCN10 | Cluster 1 | ALS-OxA |
| ENSG00000187955 | COL14A1 | Cluster 1 | ALS-OxA |
| ENSG00000188511 | MIR3667HG | Cluster 1 | ALS-OxA |
| ENSG00000188536 | HBA2 | Cluster 1 | ALS-OxA |
| ENSG00000196154 | S100A4 | Cluster 1 | ALS-OxA |
| ENSG00000196616 | ADH1B | Cluster 1 | ALS-OxA |
| ENSG00000196954 | CASP4 | Cluster 1 | ALS-OxA |
| ENSG00000197405 | C5AR1 | Cluster 1 | ALS-OxA |
| ENSG00000197901 | SLC22A6 | Cluster 1 | ALS-OxA |
| ENSG00000198467 | TPM2 | Cluster 1 | ALS-OxA |
| ENSG00000198848 | CES1 | Cluster 1 | ALS-OxA |
| ENSG00000198959 | TGM2 | Cluster 1 | ALS-OxA |
| ENSG00000204389 | HSPA1A | Cluster 1 | ALS-OxA |
| ENSG00000205038 | PKHD1L1 | Cluster 1 | ALS-OxA |
| ENSG00000205358 | MT1H | Cluster 1 | ALS-OxA |
| ENSG00000205364 | MT1M | Cluster 1 | ALS-OxA |
| ENSG00000205592 | MUC19 | Cluster 1 | ALS-OxA |
| ENSG00000206172 | HBA1 | Cluster 1 | ALS-OxA |
| ENSG00000206538 | VGLL3 | Cluster 1 | ALS-OxA |
| ENSG00000221869 | CEBPD | Cluster 1 | ALS-OxA |
| ENSG00000233117 | LINC00702 | Cluster 1 | ALS-OxA |
| ENSG00000235750 | KIAA0040 | Cluster 1 | ALS-OxA |
| ENSG00000236908 | LINC02827 | Cluster 1 | ALS-OxA |
| ENSG00000237424 | FOXD2-AS1 | Cluster 1 | ALS-OxA |
| ENSG00000241644 | INMT | Cluster 1 | ALS-OxA |
| ENSG00000244734 | HBB | Cluster 1 | ALS-OxA |
| ENSG00000249669 | CARMN | Cluster 1 | ALS-OxA |
| ENSG00000250863 | Lnc-KCTD8-1 | Cluster 1 | ALS-OxA |
| ENSG00000253123 | LOC100507403 | Cluster 1 | ALS-OxA |
| ENSG00000253686 | LINC01484 | Cluster 1 | ALS-OxA |
| ENSG00000256443 | Lnc-CPSF7-3 | Cluster 1 | ALS-OxA |
| ENSG00000256955 | Lnc-MMP17-1 | Cluster 1 | ALS-OxA |

| | | | |
|---|---|---|---|
| ENSG00000258647 | LINC00930 | Cluster 1 | ALS-OxA |
| ENSG00000260337 | Lnc-FAM174B-1 | Cluster 1 | ALS-OxA |
| ENSG00000265107 | GJA5 | Cluster 1 | ALS-OxA |
| ENSG00000267065 | LINC02080 | Cluster 1 | ALS-OxA |
| ENSG00000268388 | FENDRR | Cluster 1 | ALS-OxA |
| ENSG00000269113 | TRABD2B | Cluster 1 | ALS-OxA |
| ENSG00000270640 | Lnc-BABAM2-2 | Cluster 1 | ALS-OxA |
| ENSG00000272512 | Lnc-HES4-2 | Cluster 1 | ALS-OxA |
| ENSG00000279320 | None | Cluster 1 | ALS-OxA |
| ENSG00000279821 | None | Cluster 1 | ALS-OxA |
| ENSG00000281327 | LINC01338 | Cluster 1 | ALS-OxA |
| ENSG00000005844 | ITGAL | Cluster 2 | ALS-Neu |
| ENSG00000006042 | TMEM98 | Cluster 2 | ALS-Neu |
| ENSG00000006747 | SCIN | Cluster 2 | ALS-Neu |
| ENSG00000011426 | ANLN | Cluster 2 | ALS-Neu |
| ENSG00000011600 | TYROBP | Cluster 2 | ALS-Neu |
| ENSG00000012124 | CD22 | Cluster 2 | ALS-Neu |
| ENSG00000012223 | LTF | Cluster 2 | ALS-Neu |
| ENSG00000012779 | ALOX5 | Cluster 2 | ALS-Neu |
| ENSG00000013297 | CLDN11 | Cluster 2 | ALS-Neu |
| ENSG00000014257 | ACP3 | Cluster 2 | ALS-Neu |
| ENSG00000016602 | CLCA4 | Cluster 2 | ALS-Neu |
| ENSG00000018280 | SLC11A1 | Cluster 2 | ALS-Neu |
| ENSG00000019582 | CD74 | Cluster 2 | ALS-Neu |
| ENSG00000025708 | TYMP | Cluster 2 | ALS-Neu |
| ENSG00000026508 | CD44 | Cluster 2 | ALS-Neu |
| ENSG00000038945 | MSR1 | Cluster 2 | ALS-Neu |
| ENSG00000041982 | TNC | Cluster 2 | ALS-Neu |
| ENSG00000042493 | CAPG | Cluster 2 | ALS-Neu |
| ENSG00000051523 | CYBA | Cluster 2 | ALS-Neu |
| ENSG00000064300 | NGFR | Cluster 2 | ALS-Neu |
| ENSG00000064886 | CHI3L2 | Cluster 2 | ALS-Neu |
| ENSG00000066294 | CD84 | Cluster 2 | ALS-Neu |
| ENSG00000066336 | SPI1 | Cluster 2 | ALS-Neu |
| ENSG00000070190 | DAPP1 | Cluster 2 | ALS-Neu |
| ENSG00000071991 | CDH19 | Cluster 2 | ALS-Neu |
| ENSG00000073737 | DHRS9 | Cluster 2 | ALS-Neu |
| ENSG00000077420 | APBB1IP | Cluster 2 | ALS-Neu |
| ENSG00000080031 | PTPRH | Cluster 2 | ALS-Neu |
| ENSG00000081237 | PTPRC | Cluster 2 | ALS-Neu |
| ENSG00000081479 | LRP2 | Cluster 2 | ALS-Neu |
| ENSG00000082074 | FYB1 | Cluster 2 | ALS-Neu |
| ENSG00000086205 | FOLH1 | Cluster 2 | ALS-Neu |
| ENSG00000086730 | LAT2 | Cluster 2 | ALS-Neu |
| ENSG00000090104 | RGS1 | Cluster 2 | ALS-Neu |
| ENSG00000090382 | LYZ | Cluster 2 | ALS-Neu |
| ENSG00000095970 | TREM2 | Cluster 2 | ALS-Neu |
| ENSG00000100292 | HMOX1 | Cluster 2 | ALS-Neu |
| ENSG00000100365 | NCF4 | Cluster 2 | ALS-Neu |
| ENSG00000100368 | CSF2RB | Cluster 2 | ALS-Neu |
| ENSG00000101049 | SGK2 | Cluster 2 | ALS-Neu |
| ENSG00000101336 | HCK | Cluster 2 | ALS-Neu |
| ENSG00000101670 | LIPG | Cluster 2 | ALS-Neu |
| ENSG00000103089 | FA2H | Cluster 2 | ALS-Neu |
| ENSG00000104972 | LILRB1 | Cluster 2 | ALS-Neu |
| ENSG00000104974 | LILRA1 | Cluster 2 | ALS-Neu |
| ENSG00000105122 | RASAL3 | Cluster 2 | ALS-Neu |
| ENSG00000105281 | SLC1A5 | Cluster 2 | ALS-Neu |
| ENSG00000105366 | SIGLEC8 | Cluster 2 | ALS-Neu |
| ENSG00000105383 | CD33 | Cluster 2 | ALS-Neu |
| ENSG00000105695 | MAG | Cluster 2 | ALS-Neu |
| ENSG00000105697 | HAMP | Cluster 2 | ALS-Neu |
| ENSG00000105967 | TFEC | Cluster 2 | ALS-Neu |
| ENSG00000107099 | DOCK8 | Cluster 2 | ALS-Neu |
| ENSG00000108691 | CCL2 | Cluster 2 | ALS-Neu |
| ENSG00000108798 | ABI3 | Cluster 2 | ALS-Neu |
| ENSG00000110077 | MS4A6A | Cluster 2 | ALS-Neu |
| ENSG00000110079 | MS4A4A | Cluster 2 | ALS-Neu |
| ENSG00000110934 | BIN2 | Cluster 2 | ALS-Neu |
| ENSG00000112799 | LY86 | Cluster 2 | ALS-Neu |
| ENSG00000113396 | SLC27A6 | Cluster 2 | ALS-Neu |

| | | | |
|---|---|---|---|
| ENSG00000114013 | CD86 | Cluster 2 | ALS-Neu |
| ENSG00000115523 | GNLY | Cluster 2 | ALS-Neu |
| ENSG00000115956 | PLEK | Cluster 2 | ALS-Neu |
| ENSG00000116774 | OLFML3 | Cluster 2 | ALS-Neu |
| ENSG00000117228 | GBP1 | Cluster 2 | ALS-Neu |
| ENSG00000118640 | VAMP8 | Cluster 2 | ALS-Neu |
| ENSG00000118785 | SPP1 | Cluster 2 | ALS-Neu |
| ENSG00000119535 | CSF3R | Cluster 2 | ALS-Neu |
| ENSG00000121933 | TMIGD3 | Cluster 2 | ALS-Neu |
| ENSG00000122367 | LDB3 | Cluster 2 | ALS-Neu |
| ENSG00000122585 | NPY | Cluster 2 | ALS-Neu |
| ENSG00000125730 | C3 | Cluster 2 | ALS-Neu |
| ENSG00000126266 | FFAR1 | Cluster 2 | ALS-Neu |
| ENSG00000126860 | EVI2A | Cluster 2 | ALS-Neu |
| ENSG00000127324 | TSPAN8 | Cluster 2 | ALS-Neu |
| ENSG00000127412 | TRPV5 | Cluster 2 | ALS-Neu |
| ENSG00000127507 | ADGRE2 | Cluster 2 | ALS-Neu |
| ENSG00000128262 | POM121L9P | Cluster 2 | ALS-Neu |
| ENSG00000128283 | CDC42EP1 | Cluster 2 | ALS-Neu |
| ENSG00000128340 | RAC2 | Cluster 2 | ALS-Neu |
| ENSG00000128645 | HOXD1 | Cluster 2 | ALS-Neu |
| ENSG00000128652 | HOXD3 | Cluster 2 | ALS-Neu |
| ENSG00000129450 | SIGLEC9 | Cluster 2 | ALS-Neu |
| ENSG00000129465 | RIPK3 | Cluster 2 | ALS-Neu |
| ENSG00000129667 | RHBDF2 | Cluster 2 | ALS-Neu |
| ENSG00000129757 | CDKN1C | Cluster 2 | ALS-Neu |
| ENSG00000130300 | PLVAP | Cluster 2 | ALS-Neu |
| ENSG00000130592 | LSP1 | Cluster 2 | ALS-Neu |
| ENSG00000131401 | NAPSB | Cluster 2 | ALS-Neu |
| ENSG00000131747 | TOP2A | Cluster 2 | ALS-Neu |
| ENSG00000132702 | HAPLN2 | Cluster 2 | ALS-Neu |
| ENSG00000132854 | KANK4 | Cluster 2 | ALS-Neu |
| ENSG00000132965 | ALOX5AP | Cluster 2 | ALS-Neu |
| ENSG00000133048 | CHI3L1 | Cluster 2 | ALS-Neu |
| ENSG00000133321 | PLAAT4 | Cluster 2 | ALS-Neu |
| ENSG00000134061 | CD180 | Cluster 2 | ALS-Neu |
| ENSG00000134516 | DOCK2 | Cluster 2 | ALS-Neu |
| ENSG00000134817 | APLNR | Cluster 2 | ALS-Neu |
| ENSG00000136167 | LCP1 | Cluster 2 | ALS-Neu |
| ENSG00000136235 | GPNMB | Cluster 2 | ALS-Neu |
| ENSG00000136286 | MYO1G | Cluster 2 | ALS-Neu |
| ENSG00000136867 | SLC31A2 | Cluster 2 | ALS-Neu |
| ENSG00000136960 | ENPP2 | Cluster 2 | ALS-Neu |
| ENSG00000137462 | TLR2 | Cluster 2 | ALS-Neu |
| ENSG00000137752 | CASP1 | Cluster 2 | ALS-Neu |
| ENSG00000137869 | CYP19A1 | Cluster 2 | ALS-Neu |
| ENSG00000138316 | ADAMTS14 | Cluster 2 | ALS-Neu |
| ENSG00000138964 | PARVG | Cluster 2 | ALS-Neu |
| ENSG00000139292 | LGR5 | Cluster 2 | ALS-Neu |
| ENSG00000139629 | GALNT6 | Cluster 2 | ALS-Neu |
| ENSG00000140030 | GPR65 | Cluster 2 | ALS-Neu |
| ENSG00000140379 | BCL2A1 | Cluster 2 | ALS-Neu |
| ENSG00000140873 | ADAMTS18 | Cluster 2 | ALS-Neu |
| ENSG00000140968 | IRF8 | Cluster 2 | ALS-Neu |
| ENSG00000141526 | SLC16A3 | Cluster 2 | ALS-Neu |
| ENSG00000142512 | SIGLEC10 | Cluster 2 | ALS-Neu |
| ENSG00000142583 | SLC2A5 | Cluster 2 | ALS-Neu |
| ENSG00000143226 | FCGR2A | Cluster 2 | ALS-Neu |
| ENSG00000145287 | PLAC8 | Cluster 2 | ALS-Neu |
| ENSG00000146192 | FGD2 | Cluster 2 | ALS-Neu |
| ENSG00000148773 | MKI67 | Cluster 2 | ALS-Neu |
| ENSG00000148826 | NKX6-2 | Cluster 2 | ALS-Neu |
| ENSG00000148908 | RGS10 | Cluster 2 | ALS-Neu |
| ENSG00000150337 | FCGR1A | Cluster 2 | ALS-Neu |
| ENSG00000150656 | CNDP1 | Cluster 2 | ALS-Neu |
| ENSG00000152689 | RASGRP3 | Cluster 2 | ALS-Neu |
| ENSG00000152766 | ANKRD22 | Cluster 2 | ALS-Neu |
| ENSG00000152804 | HHEX | Cluster 2 | ALS-Neu |
| ENSG00000153551 | CMTM7 | Cluster 2 | ALS-Neu |
| ENSG00000154642 | C21orf91 | Cluster 2 | ALS-Neu |
| ENSG00000154864 | PIEZO2 | Cluster 2 | ALS-Neu |

| | | | |
|---|---|---|---|
| ENSG00000155307 | SAMSN1 | Cluster 2 | ALS-Neu |
| ENSG00000155629 | PIK3AP1 | Cluster 2 | ALS-Neu |
| ENSG00000155926 | SLA | Cluster 2 | ALS-Neu |
| ENSG00000157227 | MMP14 | Cluster 2 | ALS-Neu |
| ENSG00000158714 | SLAMF8 | Cluster 2 | ALS-Neu |
| ENSG00000158715 | SLC45A3 | Cluster 2 | ALS-Neu |
| ENSG00000158859 | ADAMTS4 | Cluster 2 | ALS-Neu |
| ENSG00000158865 | SLC5A11 | Cluster 2 | ALS-Neu |
| ENSG00000158869 | FCER1G | Cluster 2 | ALS-Neu |
| ENSG00000159189 | C1QC | Cluster 2 | ALS-Neu |
| ENSG00000159216 | RUNX1 | Cluster 2 | ALS-Neu |
| ENSG00000159399 | HK2 | Cluster 2 | ALS-Neu |
| ENSG00000160223 | ICOSLG | Cluster 2 | ALS-Neu |
| ENSG00000160255 | ITGB2 | Cluster 2 | ALS-Neu |
| ENSG00000160791 | CCR5 | Cluster 2 | ALS-Neu |
| ENSG00000161896 | IP6K3 | Cluster 2 | ALS-Neu |
| ENSG00000162444 | RBP7 | Cluster 2 | ALS-Neu |
| ENSG00000162511 | LAPTM5 | Cluster 2 | ALS-Neu |
| ENSG00000162949 | CAPN13 | Cluster 2 | ALS-Neu |
| ENSG00000163191 | S100A11 | Cluster 2 | ALS-Neu |
| ENSG00000163563 | MNDA | Cluster 2 | ALS-Neu |
| ENSG00000163565 | IFI16 | Cluster 2 | ALS-Neu |
| ENSG00000163694 | RBM47 | Cluster 2 | ALS-Neu |
| ENSG00000163823 | CCR1 | Cluster 2 | ALS-Neu |
| ENSG00000164342 | TLR3 | Cluster 2 | ALS-Neu |
| ENSG00000165025 | SYK | Cluster 2 | ALS-Neu |
| ENSG00000166091 | CMTM5 | Cluster 2 | ALS-Neu |
| ENSG00000166473 | PKD1L2 | Cluster 2 | ALS-Neu |
| ENSG00000166923 | GREM1 | Cluster 2 | ALS-Neu |
| ENSG00000166927 | MS4A7 | Cluster 2 | ALS-Neu |
| ENSG00000167208 | SNX20 | Cluster 2 | ALS-Neu |
| ENSG00000167588 | GPD1 | Cluster 2 | ALS-Neu |
| ENSG00000167613 | LAIR1 | Cluster 2 | ALS-Neu |
| ENSG00000167641 | PPP1R14A | Cluster 2 | ALS-Neu |
| ENSG00000167755 | KLK6 | Cluster 2 | ALS-Neu |
| ENSG00000167851 | CD300A | Cluster 2 | ALS-Neu |
| ENSG00000168329 | CX3CR1 | Cluster 2 | ALS-Neu |
| ENSG00000169403 | PTAFR | Cluster 2 | ALS-Neu |
| ENSG00000169413 | RNASE6 | Cluster 2 | ALS-Neu |
| ENSG00000169896 | ITGAM | Cluster 2 | ALS-Neu |
| ENSG00000170458 | CD14 | Cluster 2 | ALS-Neu |
| ENSG00000170775 | GPR37 | Cluster 2 | ALS-Neu |
| ENSG00000171051 | FPR1 | Cluster 2 | ALS-Neu |
| ENSG00000171631 | P2RY6 | Cluster 2 | ALS-Neu |
| ENSG00000171840 | NINJ2 | Cluster 2 | ALS-Neu |
| ENSG00000171860 | C3AR1 | Cluster 2 | ALS-Neu |
| ENSG00000172005 | MAL | Cluster 2 | ALS-Neu |
| ENSG00000172243 | CLEC7A | Cluster 2 | ALS-Neu |
| ENSG00000172548 | NIPAL4 | Cluster 2 | ALS-Neu |
| ENSG00000172578 | KLHL6 | Cluster 2 | ALS-Neu |
| ENSG00000173253 | DMRT2 | Cluster 2 | ALS-Neu |
| ENSG00000173369 | C1QB | Cluster 2 | ALS-Neu |
| ENSG00000173372 | C1QA | Cluster 2 | ALS-Neu |
| ENSG00000173391 | OLR1 | Cluster 2 | ALS-Neu |
| ENSG00000173988 | LRRC63 | Cluster 2 | ALS-Neu |
| ENSG00000174123 | TLR10 | Cluster 2 | ALS-Neu |
| ENSG00000174125 | TLR1 | Cluster 2 | ALS-Neu |
| ENSG00000174607 | UGT8 | Cluster 2 | ALS-Neu |
| ENSG00000174837 | ADGRE1 | Cluster 2 | ALS-Neu |
| ENSG00000175489 | LRRC25 | Cluster 2 | ALS-Neu |
| ENSG00000175567 | UCP2 | Cluster 2 | ALS-Neu |
| ENSG00000175785 | PRIMA1 | Cluster 2 | ALS-Neu |
| ENSG00000176381 | PRR18 | Cluster 2 | ALS-Neu |
| ENSG00000178082 | TWF1P1 | Cluster 2 | ALS-Neu |
| ENSG00000179178 | TMEM125 | Cluster 2 | ALS-Neu |
| ENSG00000179420 | OR6W1P | Cluster 2 | ALS-Neu |
| ENSG00000179468 | OR9A2 | Cluster 2 | ALS-Neu |
| ENSG00000180353 | HCLS1 | Cluster 2 | ALS-Neu |
| ENSG00000180739 | S1PR5 | Cluster 2 | ALS-Neu |
| ENSG00000180929 | GPR62 | Cluster 2 | ALS-Neu |
| ENSG00000183160 | TMEM119 | Cluster 2 | ALS-Neu |

| | | | |
|---|---|---|---|
| ENSG00000183196 | CHST6 | Cluster 2 | ALS-Neu |
| ENSG00000183760 | ACP7 | Cluster 2 | ALS-Neu |
| ENSG00000184357 | H1-5 | Cluster 2 | ALS-Neu |
| ENSG00000184574 | LPAR5 | Cluster 2 | ALS-Neu |
| ENSG00000184730 | APOBR | Cluster 2 | ALS-Neu |
| ENSG00000185811 | IKZF1 | Cluster 2 | ALS-Neu |
| ENSG00000185905 | C16orf54 | Cluster 2 | ALS-Neu |
| ENSG00000186417 | GLDN | Cluster 2 | ALS-Neu |
| ENSG00000187554 | TLR5 | Cluster 2 | ALS-Neu |
| ENSG00000187908 | DMBT1 | Cluster 2 | ALS-Neu |
| ENSG00000187912 | CLEC17A | Cluster 2 | ALS-Neu |
| ENSG00000188488 | SERPINA5 | Cluster 2 | ALS-Neu |
| ENSG00000196126 | HLA-DRB1 | Cluster 2 | ALS-Neu |
| ENSG00000196136 | SERPINA3 | Cluster 2 | ALS-Neu |
| ENSG00000197249 | SERPINA1 | Cluster 2 | ALS-Neu |
| ENSG00000197430 | OPALIN | Cluster 2 | ALS-Neu |
| ENSG00000197471 | SPN | Cluster 2 | ALS-Neu |
| ENSG00000197747 | S100A10 | Cluster 2 | ALS-Neu |
| ENSG00000197993 | KEL | Cluster 2 | ALS-Neu |
| ENSG00000198019 | FCGR1B | Cluster 2 | ALS-Neu |
| ENSG00000198502 | HLA-DRB5 | Cluster 2 | ALS-Neu |
| ENSG00000198774 | RASSF9 | Cluster 2 | ALS-Neu |
| ENSG00000198835 | GJC2 | Cluster 2 | ALS-Neu |
| ENSG00000203710 | CR1 | Cluster 2 | ALS-Neu |
| ENSG00000203747 | FCGR3A | Cluster 2 | ALS-Neu |
| ENSG00000204161 | TMEM273 | Cluster 2 | ALS-Neu |
| ENSG00000204252 | HLA-DOA | Cluster 2 | ALS-Neu |
| ENSG00000204278 | TMEM235 | Cluster 2 | ALS-Neu |
| ENSG00000204287 | HLA-DRA | Cluster 2 | ALS-Neu |
| ENSG00000204472 | AIF1 | Cluster 2 | ALS-Neu |
| ENSG00000204482 | LST1 | Cluster 2 | ALS-Neu |
| ENSG00000204655 | MOG | Cluster 2 | ALS-Neu |
| ENSG00000205116 | TMEM88B | Cluster 2 | ALS-Neu |
| ENSG00000205403 | CFI | Cluster 2 | ALS-Neu |
| ENSG00000207955 | MIR219A2HG | Cluster 2 | ALS-Neu |
| ENSG00000213186 | TRIM59 | Cluster 2 | ALS-Neu |
| ENSG00000214688 | C10orf105 | Cluster 2 | ALS-Neu |
| ENSG00000221887 | HMSD | Cluster 2 | ALS-Neu |
| ENSG00000224189 | HAGLR | Cluster 2 | ALS-Neu |
| ENSG00000224389 | C4B | Cluster 2 | ALS-Neu |
| ENSG00000224397 | PELATON | Cluster 2 | ALS-Neu |
| ENSG00000225217 | HSPA7 | Cluster 2 | ALS-Neu |
| ENSG00000226496 | LINC00323 | Cluster 2 | ALS-Neu |
| ENSG00000226994 | Lnc-RASGRP3-5 | Cluster 2 | ALS-Neu |
| ENSG00000227544 | LINC03013 | Cluster 2 | ALS-Neu |
| ENSG00000228058 | LINC01736 | Cluster 2 | ALS-Neu |
| ENSG00000228392 | None | Cluster 2 | ALS-Neu |
| ENSG00000228408 | Lnc-TAB2-1 | Cluster 2 | ALS-Neu |
| ENSG00000228789 | HCG22 | Cluster 2 | ALS-Neu |
| ENSG00000228793 | LOC100507336 | Cluster 2 | ALS-Neu |
| ENSG00000231131 | LNCAROD | Cluster 2 | ALS-Neu |
| ENSG00000231389 | HLA-DPA1 | Cluster 2 | ALS-Neu |
| ENSG00000231427 | LINC01445 | Cluster 2 | ALS-Neu |
| ENSG00000232310 | LINC03002 | Cluster 2 | ALS-Neu |
| ENSG00000232474 | NCKAP5-IT1 | Cluster 2 | ALS-Neu |
| ENSG00000232504 | ST3GAL5-AS1 | Cluster 2 | ALS-Neu |
| ENSG00000232560 | LINC01549 | Cluster 2 | ALS-Neu |
| ENSG00000233670 | PIRT | Cluster 2 | ALS-Neu |
| ENSG00000235568 | NFAM1 | Cluster 2 | ALS-Neu |
| ENSG00000236700 | LINC01010 | Cluster 2 | ALS-Neu |
| ENSG00000237166 | LINC01792 | Cluster 2 | ALS-Neu |
| ENSG00000239961 | LILRA4 | Cluster 2 | ALS-Neu |
| ENSG00000239998 | LILRA2 | Cluster 2 | ALS-Neu |
| ENSG00000240534 | RPL34P17 | Cluster 2 | ALS-Neu |
| ENSG00000240583 | AQP1 | Cluster 2 | ALS-Neu |
| ENSG00000241345 | LOC105375483 | Cluster 2 | ALS-Neu |
| ENSG00000242574 | HLA-DMB | Cluster 2 | ALS-Neu |
| ENSG00000244062 | ACSL3 Pseudogene | Cluster 2 | ALS-Neu |
| ENSG00000244682 | FCGR2C | Cluster 2 | ALS-Neu |
| ENSG00000244731 | C4A | Cluster 2 | ALS-Neu |
| ENSG00000244738 | Lnc-CLDN11-2 | Cluster 2 | ALS-Neu |

| | | | |
|---|---|---|---|
| ENSG00000248208 | WDR45P1 | Cluster 2 | ALS-Neu |
| ENSG00000250072 | SH3TC2-DT | Cluster 2 | ALS-Neu |
| ENSG00000250198 | LINC02199 | Cluster 2 | ALS-Neu |
| ENSG00000250771 | TLR2 Pseudogene | Cluster 2 | ALS-Neu |
| ENSG00000251138 | LINC02882 | Cluster 2 | ALS-Neu |
| ENSG00000251429 | AIDAP2 | Cluster 2 | ALS-Neu |
| ENSG00000251442 | LINC01094 | Cluster 2 | ALS-Neu |
| ENSG00000253877 | LINC01608 | Cluster 2 | ALS-Neu |
| ENSG00000254081 | LINC01299 | Cluster 2 | ALS-Neu |
| ENSG00000254415 | SIGLEC14 | Cluster 2 | ALS-Neu |
| ENSG00000257585 | LINC00609 | Cluster 2 | ALS-Neu |
| ENSG00000258227 | CLEC5A | Cluster 2 | ALS-Neu |
| ENSG00000258342 | LINC00609 | Cluster 2 | ALS-Neu |
| ENSG00000261121 | LINC02473 | Cluster 2 | ALS-Neu |
| ENSG00000261710 | Lnc-ADGRB1-3 | Cluster 2 | ALS-Neu |
| ENSG00000261795 | Lnc-PDGFA-4 | Cluster 2 | ALS-Neu |
| ENSG00000261997 | None | Cluster 2 | ALS-Neu |
| ENSG00000262188 | LINC01978 | Cluster 2 | ALS-Neu |
| ENSG00000265531 | FCGR1CP | Cluster 2 | ALS-Neu |
| ENSG00000267287 | Lnc-NFATC1-1 | Cluster 2 | ALS-Neu |
| ENSG00000267432 | DNAH17-AS1 | Cluster 2 | ALS-Neu |
| ENSG00000268170 | Lnc-FAM131B-2 | Cluster 2 | ALS-Neu |
| ENSG00000268758 | ADGRE4P | Cluster 2 | ALS-Neu |
| ENSG00000269553 | Lnc-MAG-2 | Cluster 2 | ALS-Neu |
| ENSG00000271605 | MILR1 | Cluster 2 | ALS-Neu |
| ENSG00000272808 | GCAWKR | Cluster 2 | ALS-Neu |
| ENSG00000273259 | Thioesterase | Cluster 2 | ALS-Neu |
| ENSG00000275395 | FCGBP | Cluster 2 | ALS-Neu |
| ENSG00000276980 | Lnc-GPR108-3 | Cluster 2 | ALS-Neu |
| ENSG00000277117 | LOC102723996 | Cluster 2 | ALS-Neu |
| ENSG00000277494 | GPIHBP1 | Cluster 2 | ALS-Neu |
| ENSG00000279419 | None | Cluster 2 | ALS-Neu |
| ENSG00000282608 | ADORA3 | Cluster 2 | ALS-Neu |
| ENSG00000283431 | Novel Zinc Finger Protein Pseudogene | Cluster 2 | ALS-Neu |
| ENSG00000283462 | LOC101930276 | Cluster 2 | ALS-Neu |
| ENSG00000283608 | Lnc-SUMO4-6 | Cluster 2 | ALS-Neu |
| ENSG00000285269 | SLC35D2-HSD17B3 readthrough | Cluster 2 | ALS-Neu |
| ENSG00000006128 | TAC1 | Cluster 3 | ALS-SNs |
| ENSG00000100362 | PVALB | Cluster 3 | ALS-SNs |
| ENSG00000104938 | CLEC4M | Cluster 3 | ALS-SNs |
| ENSG00000106236 | NPTX2 | Cluster 3 | ALS-SNs |
| ENSG00000107165 | TYRP1 | Cluster 3 | ALS-SNs |
| ENSG00000111863 | ADTRP | Cluster 3 | ALS-SNs |
| ENSG00000117594 | HSD11B1 | Cluster 3 | ALS-SNs |
| ENSG00000120875 | DUSP4 | Cluster 3 | ALS-SNs |
| ENSG00000122012 | SV2C | Cluster 3 | ALS-SNs |
| ENSG00000122025 | FLT3 | Cluster 3 | ALS-SNs |
| ENSG00000126545 | CSN1S1 | Cluster 3 | ALS-SNs |
| ENSG00000128564 | VGF | Cluster 3 | ALS-SNs |
| ENSG00000130701 | RBBP8NL | Cluster 3 | ALS-SNs |
| ENSG00000131885 | KRT17P1 | Cluster 3 | ALS-SNs |
| ENSG00000132744 | ACY3 | Cluster 3 | ALS-SNs |
| ENSG00000138678 | GPAT3 | Cluster 3 | ALS-SNs |
| ENSG00000141433 | ADCYAP1 | Cluster 3 | ALS-SNs |
| ENSG00000143858 | SYT2 | Cluster 3 | ALS-SNs |
| ENSG00000144227 | NXPH2 | Cluster 3 | ALS-SNs |
| ENSG00000145708 | CRHBP | Cluster 3 | ALS-SNs |
| ENSG00000146469 | VIP | Cluster 3 | ALS-SNs |
| ENSG00000147571 | CRH | Cluster 3 | ALS-SNs |
| ENSG00000150175 | FRMPD2B | Cluster 3 | ALS-SNs |
| ENSG00000151790 | TDO2 | Cluster 3 | ALS-SNs |
| ENSG00000152595 | MEPE | Cluster 3 | ALS-SNs |
| ENSG00000156076 | WIF1 | Cluster 3 | ALS-SNs |
| ENSG00000157005 | SST | Cluster 3 | ALS-SNs |
| ENSG00000159248 | GJD2 | Cluster 3 | ALS-SNs |
| ENSG00000160221 | GATD3A | Cluster 3 | ALS-SNs |
| ENSG00000164326 | CARTPT | Cluster 3 | ALS-SNs |
| ENSG00000164600 | NEUROD6 | Cluster 3 | ALS-SNs |
| ENSG00000165899 | OTOGL | Cluster 3 | ALS-SNs |
| ENSG00000169035 | KLK7 | Cluster 3 | ALS-SNs |
| ENSG00000169676 | DRD5 | Cluster 3 | ALS-SNs |

| | | | |
|---|---|---|---|
| ENSG00000170231 | FABP6 | Cluster 3 | ALS-SNs |
| ENSG00000170290 | SLN | Cluster 3 | ALS-SNs |
| ENSG00000174403 | MIR1-1HG-AS1 | Cluster 3 | ALS-SNs |
| ENSG00000174576 | NPAS4 | Cluster 3 | ALS-SNs |
| ENSG00000176697 | BDNF | Cluster 3 | ALS-SNs |
| ENSG00000179477 | ALOX12B | Cluster 3 | ALS-SNs |
| ENSG00000179520 | SLC17A8 | Cluster 3 | ALS-SNs |
| ENSG00000182632 | CCNYL2 | Cluster 3 | ALS-SNs |
| ENSG00000183090 | FREM3 | Cluster 3 | ALS-SNs |
| ENSG00000185149 | NPY2R | Cluster 3 | ALS-SNs |
| ENSG00000186081 | KRT5 | Cluster 3 | ALS-SNs |
| ENSG00000188729 | OSTN | Cluster 3 | ALS-SNs |
| ENSG00000200959 | SNORA74A | Cluster 3 | ALS-SNs |
| ENSG00000205279 | CTXN3 | Cluster 3 | ALS-SNs |
| ENSG00000205634 | LINC00898 | Cluster 3 | ALS-SNs |
| ENSG00000223812 | PYDC2-AS1 | Cluster 3 | ALS-SNs |
| ENSG00000223930 | Lnc-ZMAT3-3 | Cluster 3 | ALS-SNs |
| ENSG00000224404 | Lnc-LARGE1-1 | Cluster 3 | ALS-SNs |
| ENSG00000226405 | HSD17B12 Pseudogene | Cluster 3 | ALS-SNs |
| ENSG00000227582 | ADGRF5P1 | Cluster 3 | ALS-SNs |
| ENSG00000228999 | LINC01830 | Cluster 3 | ALS-SNs |
| ENSG00000229618 | Lnc-ARL4A-54 | Cluster 3 | ALS-SNs |
| ENSG00000231824 | AKAIN1 | Cluster 3 | ALS-SNs |
| ENSG00000233123 | LINC01007 | Cluster 3 | ALS-SNs |
| ENSG00000236714 | LINC01844 | Cluster 3 | ALS-SNs |
| ENSG00000237390 | Lnc-RHBG-1 | Cluster 3 | ALS-SNs |
| ENSG00000239672 | NME1 | Cluster 3 | ALS-SNs |
| ENSG00000241945 | PWP2 | Cluster 3 | ALS-SNs |
| ENSG00000242159 | ABCF2P1 | Cluster 3 | ALS-SNs |
| ENSG00000246363 | LINC02458 | Cluster 3 | ALS-SNs |
| ENSG00000248837 | LOC105374524 | Cluster 3 | ALS-SNs |
| ENSG00000250634 | LINC01182 | Cluster 3 | ALS-SNs |
| ENSG00000254101 | LINC02055 | Cluster 3 | ALS-SNs |
| ENSG00000254363 | LOC101929719 | Cluster 3 | ALS-SNs |
| ENSG00000254510 | Lnc-NPAS4-1 | Cluster 3 | ALS-SNs |
| ENSG00000254561 | None | Cluster 3 | ALS-SNs |
| ENSG00000255087 | LOC101929473 | Cluster 3 | ALS-SNs |
| ENSG00000256193 | LINC00507 | Cluster 3 | ALS-SNs |
| ENSG00000259520 | SLC28A2-AS1 | Cluster 3 | ALS-SNs |
| ENSG00000260658 | Lnc-CDH8-10 | Cluster 3 | ALS-SNs |
| ENSG00000261325 | LINC02192 | Cluster 3 | ALS-SNs |
| ENSG00000261502 | Lnc-CDH8-10 | Cluster 3 | ALS-SNs |
| ENSG00000261738 | MIR3976HG | Cluster 3 | ALS-SNs |
| ENSG00000261757 | miR-1255 | Cluster 3 | ALS-SNs |
| ENSG00000265179 | Lnc-YES1-8 | Cluster 3 | ALS-SNs |
| ENSG00000267623 | Cyclin Y-Like Pseudogene | Cluster 3 | ALS-SNs |
| ENSG00000268297 | CLEC4GP1 | Cluster 3 | ALS-SNs |
| ENSG00000271538 | LINC02427 | Cluster 3 | ALS-SNs |
| ENSG00000276462 | LINC03025 | Cluster 3 | ALS-SNs |
| ENSG00000277918 | RNVU1-28 | Cluster 3 | ALS-SNs |
| ENSG00000279875 | None | Cluster 3 | ALS-SNs |
| ENSG00000280776 | LINC01202 | Cluster 3 | ALS-SNs |
| ENSG00000285561 | None | Cluster 3 | ALS-SNs |
| ENSG00000285578 | Lnc-DUSP22-2 | Cluster 3 | ALS-SNs |
| ENSG00000285634 | LOC105376121 | Cluster 3 | ALS-SNs |
| ENSG00000285735 | LINC02717 | Cluster 3 | ALS-SNs |
| ENSG00000286016 | None | Cluster 3 | ALS-SNs |
| ENSG00000286109 | Lnc-KCNK1-1 | Cluster 3 | ALS-SNs |

### **Suppl Table 2.** Enriched pathway analysis for each ALS subtype.

| Cluster | Reference ALS-Subtype | Database | Term ID | Enriched Pathway | Gene Ratio | Adj P value | Count |
|---|---|---|---|---|---|---|---|
| Cluster 1 | ALS-OxA | GO:BP | GO:0030199 | BP Collagen fibril organization | 0.075 | 0.004 | 5 |
| Cluster 1 | ALS-OxA | GO:BP | GO:0048844 | BP Artery morphogenesis | 0.075 | 0.004 | 5 |
| Cluster 1 | ALS-OxA | GO:BP | GO:0007517 | BP Muscle organ development | 0.134 | 0.004 | 9 |
| Cluster 1 | ALS-OxA | GO:BP | GO:0045765 | BP Regulation of angiogenesis | 0.119 | 0.004 | 8 |
| Cluster 1 | ALS-OxA | GO:BP | GO:0007179 | BP Transforming growth factor beta receptor signaling pathway | 0.104 | 0.004 | 7 |

| | | | | | | | |
|---|---|---|---|---|---|---|---|
| Cluster 1 | ALS-OxA | GO:BP | GO:0003413 | BP Chondrocyte differentiation involved in endochondral bone morphogenesis | 0.045 | 0.004 | 3 |
| Cluster 1 | ALS-OxA | GO:BP | GO:0072001 | BP Renal system development | 0.119 | 0.006 | 8 |
| Cluster 1 | ALS-OxA | GO:BP | GO:0006936 | BP Muscle contraction | 0.119 | 0.008 | 8 |
| Cluster 1 | ALS-OxA | GO:BP | GO:0031589 | BP Cell-substrate adhesion | 0.119 | 0.008 | 8 |
| Cluster 1 | ALS-OxA | GO:BP | GO:0003007 | BP Heart morphogenesis | 0.104 | 0.008 | 7 |
| Cluster 1 | ALS-OxA | GO:BP | GO:0071559 | BP Response to transforming growth factor beta | 0.104 | 0.008 | 7 |
| Cluster 1 | ALS-OxA | GO:BP | GO:0061138 | BP Morphogenesis of a branching epithelium | 0.090 | 0.008 | 6 |
| Cluster 1 | ALS-OxA | GO:BP | GO:0007178 | BP Cell surface receptor protein serine/threonine kinase signaling pathway | 0.119 | 0.008 | 8 |
| Cluster 1 | ALS-OxA | GO:BP | GO:0003179 | BP Heart valve morphogenesis | 0.060 | 0.008 | 4 |
| Cluster 1 | ALS-OxA | GO:BP | GO:0003229 | BP Ventricular cardiac muscle tissue development | 0.060 | 0.008 | 4 |
| Cluster 1 | ALS-OxA | GO:BP | GO:0001763 | BP Morphogenesis of a branching structure | 0.090 | 0.010 | 6 |
| Cluster 1 | ALS-OxA | GO:BP | GO:0003158 | BP Endothelium development | 0.075 | 0.012 | 5 |
| Cluster 1 | ALS-OxA | GO:BP | GO:0043542 | BP Endothelial cell migration | 0.090 | 0.013 | 6 |
| Cluster 1 | ALS-OxA | GO:BP | GO:0002825 | BP Regulation of t-helper 1 type immune response | 0.045 | 0.013 | 3 |
| Cluster 1 | ALS-OxA | GO:BP | GO:0042060 | BP Wound healing | 0.119 | 0.013 | 8 |
| Cluster 1 | ALS-OxA | GO:BP | GO:0002063 | BP Chondrocyte development | 0.045 | 0.013 | 3 |
| Cluster 1 | ALS-OxA | GO:BP | GO:0001569 | BP Branching involved in blood vessel morphogenesis | 0.045 | 0.013 | 3 |
| Cluster 1 | ALS-OxA | GO:BP | GO:0141091 | BP Transforming growth factor beta receptor superfamily signaling pathway | 0.104 | 0.017 | 7 |
| Cluster 1 | ALS-OxA | GO:BP | GO:0034109 | BP Homotypic cell-cell adhesion | 0.060 | 0.022 | 4 |
| Cluster 1 | ALS-OxA | GO:BP | GO:0009408 | BP Response to heat | 0.060 | 0.024 | 4 |
| Cluster 1 | ALS-OxA | GO:BP | GO:0035272 | BP Exocrine system development | 0.045 | 0.027 | 3 |
| Cluster 1 | ALS-OxA | GO:BP | GO:0055010 | BP Ventricular cardiac muscle tissue morphogenesis | 0.045 | 0.027 | 3 |
| Cluster 1 | ALS-OxA | GO:BP | GO:0042088 | BP T-helper 1 type immune response | 0.045 | 0.027 | 3 |
| Cluster 1 | ALS-OxA | GO:BP | GO:0007369 | BP Gastrulation | 0.075 | 0.027 | 5 |
| Cluster 1 | ALS-OxA | GO:BP | GO:0045446 | BP Endothelial cell differentiation | 0.060 | 0.030 | 4 |
| Cluster 1 | ALS-OxA | GO:BP | GO:1902414 | BP Protein localization to cell junction | 0.060 | 0.035 | 4 |
| Cluster 1 | ALS-OxA | GO:BP | GO:1902895 | BP Positive regulation of mirna transcription | 0.045 | 0.036 | 3 |
| Cluster 1 | ALS-OxA | GO:BP | GO:0048010 | BP Vascular endothelial growth factor receptor signaling pathway | 0.045 | 0.037 | 3 |
| Cluster 1 | ALS-OxA | GO:BP | GO:0060390 | BP Regulation of smad protein signal transduction | 0.045 | 0.046 | 3 |
| Cluster 1 | ALS-OxA | GO:BP | GO:0035924 | BP Cellular response to vascular endothelial growth factor stimulus | 0.045 | 0.046 | 3 |
| Cluster 1 | ALS-OxA | GO:BP | GO:2000630 | BP Positive regulation of mirna metabolic process | 0.045 | 0.046 | 3 |
| Cluster 1 | ALS-OxA | GO:BP | GO:0030168 | BP Platelet activation | 0.060 | 0.046 | 4 |
| Cluster 1 | ALS-OxA | GO:BP | GO:0006986 | BP Response to unfolded protein | 0.060 | 0.046 | 4 |
| Cluster 1 | ALS-OxA | GO:CC | GO:0062023 | CC Collagen-containing extracellular matrix | 0.212 | 0.000 | 14 |
| Cluster 1 | ALS-OxA | GO:CC | GO:0009897 | CC External side of plasma membrane | 0.152 | 0.000 | 10 |
| Cluster 1 | ALS-OxA | GO:CC | GO:0005581 | CC Collagen trimer | 0.076 | 0.000 | 5 |
| Cluster 1 | ALS-OxA | GO:CC | GO:0043292 | CC Contractile muscle fiber | 0.106 | 0.001 | 7 |
| Cluster 1 | ALS-OxA | GO:CC | GO:0005925 | CC Focal adhesion | 0.106 | 0.011 | 7 |
| Cluster 1 | ALS-OxA | GO:CC | GO:0005788 | CC Endoplasmic reticulum lumen | 0.091 | 0.011 | 6 |
| Cluster 1 | ALS-OxA | GO:CC | GO:0001725 | CC Stress fiber | 0.045 | 0.033 | 3 |
| Cluster 1 | ALS-OxA | GO:MF | GO:0005201 | MF Extracellular matrix structural constituent | 0.136 | 0.000 | 9 |
| Cluster 1 | ALS-OxA | GO:MF | GO:0005518 | MF Collagen binding | 0.076 | 0.000 | 5 |
| Cluster 1 | ALS-OxA | GO:MF | GO:0048407 | MF Platelet-derived growth factor binding | 0.045 | 0.001 | 3 |
| Cluster 1 | ALS-OxA | GO:MF | GO:0008307 | MF Structural constituent of muscle | 0.061 | 0.001 | 4 |
| Cluster 1 | ALS-OxA | GO:MF | GO:0005178 | MF Integrin binding | 0.091 | 0.001 | 6 |
| Cluster 1 | ALS-OxA | GO:MF | GO:0140416 | MF Transcription regulator inhibitor activity | 0.045 | 0.008 | 3 |
| Cluster 1 | ALS-OxA | GO:MF | GO:0098631 | MF Cell adhesion mediator activity | 0.045 | 0.041 | 3 |
| Cluster 1 | ALS-OxA | GO:MF | GO:0097110 | MF Scaffold protein binding | 0.045 | 0.041 | 3 |
| Cluster 1 | ALS-OxA | Reactome | R-HSA-216083 | React Integrin cell surface interactions | 0.167 | 0.000 | 9 |
| Cluster 1 | ALS-OxA | Reactome | R-HSA-1474244 | React Extracellular matrix organization | 0.241 | 0.000 | 13 |
| Cluster 1 | ALS-OxA | Reactome | R-HSA-1650814 | React Collagen biosynthesis and modifying enzymes | 0.111 | 0.000 | 6 |
| Cluster 1 | ALS-OxA | Reactome | R-HSA-3000178 | React ECM proteoglycans | 0.111 | 0.000 | 6 |
| Cluster 1 | ALS-OxA | Reactome | R-HSA-8948216 | React Collagen chain trimerization | 0.093 | 0.000 | 5 |
| Cluster 1 | ALS-OxA | Reactome | R-HSA-1474228 | React Degradation of the extracellular matrix | 0.130 | 0.000 | 7 |

| | | | | | | | |
|---|---|---|---|---|---|---|---|
| Cluster 1 | ALS-OxA | Reactome | R-HSA-1474290 | React Collagen formation | 0.111 | 0.000 | 6 |
| Cluster 1 | ALS-OxA | Reactome | R-HSA-3000171 | React Non-integrin membrane-ECM interactions | 0.093 | 0.000 | 5 |
| Cluster 1 | ALS-OxA | Reactome | R-HSA-2022090 | React Assembly of collagen fibrils and other multimeric structures | 0.093 | 0.000 | 5 |
| Cluster 1 | ALS-OxA | Reactome | R-HSA-8874081 | React MET activates PTK2 signaling | 0.074 | 0.001 | 4 |
| Cluster 1 | ALS-OxA | Reactome | R-HSA-8875878 | React MET promotes cell motility | 0.074 | 0.002 | 4 |
| Cluster 1 | ALS-OxA | Reactome | R-HSA-75892 | React Platelet Adhesion to exposed collagen | 0.056 | 0.002 | 3 |
| Cluster 1 | ALS-OxA | Reactome | R-HSA-2173782 | React Binding and Uptake of Ligands by Scavenger Receptors | 0.074 | 0.002 | 4 |
| Cluster 1 | ALS-OxA | Reactome | R-HSA-445355 | React Smooth Muscle Contraction | 0.074 | 0.002 | 4 |
| Cluster 1 | ALS-OxA | Reactome | R-HSA-6785807 | React Interleukin-4 and Interleukin-13 signaling | 0.093 | 0.004 | 5 |
| Cluster 1 | ALS-OxA | Reactome | R-HSA-1442490 | React Collagen degradation | 0.074 | 0.005 | 4 |
| Cluster 1 | ALS-OxA | Reactome | R-HSA-3000157 | React Laminin interactions | 0.056 | 0.009 | 3 |
| Cluster 1 | ALS-OxA | Reactome | R-HSA-397014 | React Muscle contraction | 0.111 | 0.011 | 6 |
| Cluster 1 | ALS-OxA | Reactome | R-HSA-6806834 | React Signaling by MET | 0.074 | 0.012 | 4 |
| Cluster 1 | ALS-OxA | Reactome | R-HSA-419037 | React NCAM1 interactions | 0.056 | 0.021 | 3 |
| Cluster 1 | ALS-OxA | Reactome | R-HSA-186797 | React Signaling by PDGF | 0.056 | 0.045 | 3 |
| Cluster 2 | ALS-Neu | GO:BP | GO:0042063 | BP Gliogenesis | 0.170 | 0.000 | 24 |
| Cluster 2 | ALS-Neu | GO:BP | GO:0008366 | BP Axon ensheathment | 0.121 | 0.000 | 17 |
| Cluster 2 | ALS-Neu | GO:BP | GO:0098883 | BP Synapse pruning | 0.035 | 0.000 | 5 |
| Cluster 2 | ALS-Neu | GO:BP | GO:1905521 | BP Regulation of macrophage migration | 0.043 | 0.000 | 6 |
| Cluster 2 | ALS-Neu | GO:BP | GO:1903977 | BP Positive regulation of glial cell migration | 0.028 | 0.000 | 4 |
| Cluster 2 | ALS-Neu | GO:BP | GO:0031641 | BP Regulation of myelination | 0.043 | 0.000 | 6 |
| Cluster 2 | ALS-Neu | GO:BP | GO:0014013 | BP Regulation of gliogenesis | 0.057 | 0.000 | 8 |
| Cluster 2 | ALS-Neu | GO:BP | GO:1905517 | BP Macrophage migration | 0.043 | 0.001 | 6 |
| Cluster 2 | ALS-Neu | GO:BP | GO:0097242 | BP Amyloid-beta clearance | 0.035 | 0.002 | 5 |
| Cluster 2 | ALS-Neu | GO:BP | GO:0032288 | BP Myelin assembly | 0.028 | 0.004 | 4 |
| Cluster 2 | ALS-Neu | GO:BP | GO:0050829 | BP Defense response to gram-negative bacterium | 0.043 | 0.005 | 6 |
| Cluster 2 | ALS-Neu | GO:BP | GO:0007422 | BP Peripheral nervous system development | 0.043 | 0.008 | 6 |
| Cluster 2 | ALS-Neu | GO:BP | GO:0034113 | BP Heterotypic cell-cell adhesion | 0.035 | 0.014 | 5 |
| Cluster 2 | ALS-Neu | GO:BP | GO:0031646 | BP Positive regulation of nervous system process | 0.028 | 0.016 | 4 |
| Cluster 2 | ALS-Neu | GO:BP | GO:0034381 | BP Plasma lipoprotein particle clearance | 0.028 | 0.025 | 4 |
| Cluster 2 | ALS-Neu | GO:BP | GO:0036005 | BP Response to macrophage colony-stimulating factor | 0.021 | 0.025 | 3 |
| Cluster 2 | ALS-Neu | GO:BP | GO:0043217 | BP Myelin maintenance | 0.021 | 0.025 | 3 |
| Cluster 2 | ALS-Neu | GO:BP | GO:0006958 | BP Complement activation, classical pathway | 0.028 | 0.025 | 4 |
| Cluster 2 | ALS-Neu | GO:BP | GO:0006898 | BP Receptor-mediated endocytosis | 0.064 | 0.025 | 9 |
| Cluster 2 | ALS-Neu | GO:BP | GO:0010720 | BP Positive regulation of cell development | 0.085 | 0.025 | 12 |
| Cluster 2 | ALS-Neu | GO:BP | GO:0006959 | BP Humoral immune response | 0.064 | 0.027 | 9 |
| Cluster 2 | ALS-Neu | GO:BP | GO:0140962 | BP Multicellular organismal-level chemical homeostasis | 0.035 | 0.027 | 5 |
| Cluster 2 | ALS-Neu | GO:BP | GO:0048678 | BP Response to axon injury | 0.035 | 0.030 | 5 |
| Cluster 2 | ALS-Neu | GO:BP | GO:0006911 | BP Phagocytosis, engulfment | 0.028 | 0.036 | 4 |
| Cluster 2 | ALS-Neu | GO:BP | GO:0010878 | BP Cholesterol storage | 0.021 | 0.045 | 3 |
| Cluster 2 | ALS-Neu | GO:BP | GO:0048260 | BP Positive regulation of receptor-mediated endocytosis | 0.028 | 0.048 | 4 |
| Cluster 2 | ALS-Neu | GO:BP | GO:0006826 | BP Iron ion transport | 0.028 | 0.049 | 4 |
| Cluster 2 | ALS-Neu | GO:BP | GO:1901222 | BP Regulation of non-canonical nf-kappab signal transduction | 0.035 | 0.049 | 5 |
| Cluster 2 | ALS-Neu | GO:CC | GO:0070821 | CC Tertiary granule membrane | 0.048 | 0.000 | 7 |
| Cluster 2 | ALS-Neu | GO:CC | GO:0016324 | CC Apical plasma membrane | 0.088 | 0.003 | 13 |
| Cluster 2 | ALS-Neu | GO:CC | GO:0043209 | CC Myelin sheath | 0.034 | 0.003 | 5 |
| Cluster 2 | ALS-Neu | GO:CC | GO:0045177 | CC Apical part of cell | 0.088 | 0.005 | 13 |
| Cluster 2 | ALS-Neu | GO:CC | GO:0072562 | CC Blood microparticle | 0.048 | 0.005 | 7 |
| Cluster 2 | ALS-Neu | GO:CC | GO:0044298 | CC Cell body membrane | 0.027 | 0.005 | 4 |
| Cluster 2 | ALS-Neu | GO:CC | GO:0030139 | CC Endocytic vesicle | 0.061 | 0.040 | 9 |
| Cluster 2 | ALS-Neu | GO:CC | GO:0062023 | CC Collagen-containing extracellular matrix | 0.068 | 0.040 | 10 |
| Cluster 2 | ALS-Neu | GO:CC | GO:0044853 | CC Plasma membrane raft | 0.034 | 0.040 | 5 |

| | | | | | | | |
|---|---|---|---|---|---|---|---|
| Cluster 2 | ALS-Neu | GO:CC | GO:0045121 | CC Membrane raft | 0.054 | 0.040 | 8 |
| Cluster 2 | ALS-Neu | GO:CC | GO:0045178 | CC Basal part of cell | 0.054 | 0.042 | 8 |
| Cluster 2 | ALS-Neu | GO:MF | GO:0038024 | MF Cargo receptor activity | 0.043 | 0.003 | 6 |
| Cluster 2 | ALS-Neu | GO:MF | GO:0008381 | MF Mechanosensitive monoatomic ion channel activity | 0.029 | 0.003 | 4 |
| Cluster 2 | ALS-Neu | GO:MF | GO:0019911 | MF Structural constituent of myelin sheath | 0.021 | 0.010 | 3 |
| Cluster 2 | ALS-Neu | GO:MF | GO:0015144 | MF Carbohydrate transmembrane transporter activity | 0.029 | 0.025 | 4 |
| Cluster 2 | ALS-Neu | GO:MF | GO:0005178 | MF Integrin binding | 0.050 | 0.025 | 7 |
| Cluster 2 | ALS-Neu | Reactome | R-HSA-202733 | React Cell surface interactions at the vascular wall | 0.075 | 0.048 | 7 |
| Cluster 2 | ALS-Neu | Reactome | R-HSA-8957275 | React Post-translational protein phosphorylation | 0.065 | 0.049 | 6 |
| Cluster 2 | ALS-Neu | Reactome | R-HSA-6798695 | React Neutrophil degranulation | 0.129 | 0.049 | 12 |
| Cluster 2 | ALS-Neu | Reactome | R-HSA-381426 | React Regulation of Insulin-like Growth Factor (IGF) transport and uptake by Insulin-like Growth Factor Binding Proteins (IGFBPs) | 0.065 | 0.049 | 6 |
| Cluster 2 | ALS-Neu | Reactome | R-HSA-216083 | React Integrin cell surface interactions | 0.054 | 0.049 | 5 |
| Cluster 2 | ALS-Neu | Reactome | R-HSA-166663 | React Initial triggering of complement | 0.032 | 0.049 | 3 |
| Cluster 2 | ALS-Neu | Reactome | R-HSA-1474244 | React Extracellular matrix organization | 0.097 | 0.050 | 9 |
| Cluster 2 | ALS-Neu | Reactome | R-HSA-1474228 | React Degradation of the extracellular matrix | 0.065 | 0.050 | 6 |
| Cluster 3 | ALS-SNs | GO:BP | GO:0050804 | BP Modulation of chemical synaptic transmission | 0.106 | 0.002 | 19 |
| Cluster 3 | ALS-SNs | GO:BP | GO:0007218 | BP Neuropeptide signaling pathway | 0.050 | 0.002 | 9 |
| Cluster 3 | ALS-SNs | GO:BP | GO:0051952 | BP Regulation of amine transport | 0.044 | 0.002 | 8 |
| Cluster 3 | ALS-SNs | GO:BP | GO:0099003 | BP Vesicle-mediated transport in synapse | 0.072 | 0.002 | 13 |
| Cluster 3 | ALS-SNs | GO:BP | GO:0007611 | BP Learning or memory | 0.072 | 0.004 | 13 |
| Cluster 3 | ALS-SNs | GO:BP | GO:0009306 | BP Protein secretion | 0.078 | 0.007 | 14 |
| Cluster 3 | ALS-SNs | GO:BP | GO:0035592 | BP Establishment of protein localization to extracellular region | 0.078 | 0.007 | 14 |
| Cluster 3 | ALS-SNs | GO:BP | GO:0007416 | BP Synapse assembly | 0.067 | 0.007 | 12 |
| Cluster 3 | ALS-SNs | GO:BP | GO:0060078 | BP Regulation of postsynaptic membrane potential | 0.050 | 0.007 | 9 |
| Cluster 3 | ALS-SNs | GO:BP | GO:0071692 | BP Protein localization to extracellular region | 0.078 | 0.007 | 14 |
| Cluster 3 | ALS-SNs | GO:BP | GO:0050433 | BP Regulation of catecholamine secretion | 0.028 | 0.012 | 5 |
| Cluster 3 | ALS-SNs | GO:BP | GO:0051588 | BP Regulation of neurotransmitter transport | 0.039 | 0.013 | 7 |
| Cluster 3 | ALS-SNs | GO:BP | GO:0009581 | BP Detection of external stimulus | 0.044 | 0.015 | 8 |
| Cluster 3 | ALS-SNs | GO:BP | GO:0009065 | BP Glutamine family amino acid catabolic process | 0.022 | 0.015 | 4 |
| Cluster 3 | ALS-SNs | GO:BP | GO:0009582 | BP Detection of abiotic stimulus | 0.044 | 0.015 | 8 |
| Cluster 3 | ALS-SNs | GO:BP | GO:1902476 | BP Chloride transmembrane transport | 0.039 | 0.017 | 7 |
| Cluster 3 | ALS-SNs | GO:BP | GO:0006091 | BP Generation of precursor metabolites and energy | 0.083 | 0.017 | 15 |
| Cluster 3 | ALS-SNs | GO:BP | GO:0043603 | BP Amide metabolic process | 0.078 | 0.025 | 14 |
| Cluster 3 | ALS-SNs | GO:BP | GO:0006821 | BP Chloride transport | 0.039 | 0.027 | 7 |
| Cluster 3 | ALS-SNs | GO:BP | GO:0009260 | BP Ribonucleotide biosynthetic process | 0.044 | 0.028 | 8 |
| Cluster 3 | ALS-SNs | GO:BP | GO:0006536 | BP Glutamate metabolic process | 0.022 | 0.028 | 4 |
| Cluster 3 | ALS-SNs | GO:BP | GO:0072522 | BP Purine-containing compound biosynthetic process | 0.056 | 0.028 | 10 |
| Cluster 3 | ALS-SNs | GO:BP | GO:0046390 | BP Ribose phosphate biosynthetic process | 0.044 | 0.031 | 8 |
| Cluster 3 | ALS-SNs | GO:BP | GO:0050803 | BP Regulation of synapse structure or activity | 0.061 | 0.031 | 11 |
| Cluster 3 | ALS-SNs | GO:BP | GO:0043648 | BP Dicarboxylic acid metabolic process | 0.033 | 0.033 | 6 |
| Cluster 3 | ALS-SNs | GO:BP | GO:0043649 | BP Dicarboxylic acid catabolic process | 0.017 | 0.035 | 3 |
| Cluster 3 | ALS-SNs | GO:BP | GO:0023061 | BP Signal release | 0.078 | 0.038 | 14 |
| Cluster 3 | ALS-SNs | GO:BP | GO:0060079 | BP Excitatory postsynaptic potential | 0.033 | 0.044 | 6 |
| Cluster 3 | ALS-SNs | GO:BP | GO:0015800 | BP Acidic amino acid transport | 0.028 | 0.044 | 5 |
| Cluster 3 | ALS-SNs | GO:BP | GO:0006813 | BP Potassium ion transport | 0.050 | 0.044 | 9 |
| Cluster 3 | ALS-SNs | GO:BP | GO:0030534 | BP Adult behavior | 0.039 | 0.045 | 7 |
| Cluster 3 | ALS-SNs | GO:BP | GO:0008652 | BP Amino acid biosynthetic process | 0.028 | 0.046 | 5 |
| Cluster 3 | ALS-SNs | GO:BP | GO:0031340 | BP Positive regulation of vesicle fusion | 0.017 | 0.049 | 3 |
| Cluster 3 | ALS-SNs | GO:CC | GO:0030672 | CC Synaptic vesicle membrane | 0.095 | 0.000 | 18 |
| Cluster 3 | ALS-SNs | GO:CC | GO:1902495 | CC Transmembrane transporter complex | 0.126 | 0.000 | 24 |
| Cluster 3 | ALS-SNs | GO:CC | GO:0098982 | CC Gaba-ergic synapse | 0.058 | 0.000 | 11 |
| Cluster 3 | ALS-SNs | GO:CC | GO:0043025 | CC Neuronal cell body | 0.116 | 0.000 | 22 |
| Cluster 3 | ALS-SNs | GO:CC | GO:0043679 | CC Axon terminus | 0.058 | 0.000 | 11 |
| Cluster 3 | ALS-SNs | GO:CC | GO:0044306 | CC Neuron projection terminus | 0.058 | 0.000 | 11 |
| Cluster 3 | ALS-SNs | GO:CC | GO:0150034 | CC Distal axon | 0.079 | 0.000 | 15 |

| | | | | | | | |
|---|---|---|---|---|---|---|---|
| Cluster 3 | ALS-SNs | GO:CC | GO:0097060 | CC Synaptic membrane | 0.089 | 0.000 | 17 |
| Cluster 3 | ALS-SNs | GO:CC | GO:0098803 | CC Respiratory chain complex | 0.042 | 0.000 | 8 |
| Cluster 3 | ALS-SNs | GO:CC | GO:0031045 | CC Dense core granule | 0.026 | 0.001 | 5 |
| Cluster 3 | ALS-SNs | GO:CC | GO:0005883 | CC Neurofilament | 0.016 | 0.003 | 3 |
| Cluster 3 | ALS-SNs | GO:CC | GO:0032589 | CC Neuron projection membrane | 0.026 | 0.007 | 5 |
| Cluster 3 | ALS-SNs | GO:CC | GO:0008180 | CC Cop9 signalosome | 0.021 | 0.007 | 4 |
| Cluster 3 | ALS-SNs | GO:CC | GO:0045259 | CC Proton-transporting atp synthase complex | 0.016 | 0.014 | 3 |
| Cluster 3 | ALS-SNs | GO:CC | GO:0097449 | CC Astrocyte projection | 0.016 | 0.016 | 3 |
| Cluster 3 | ALS-SNs | GO:CC | GO:0098686 | CC Hippocampal mossy fiber to ca3 synapse | 0.021 | 0.018 | 4 |
| Cluster 3 | ALS-SNs | GO:CC | GO:0019898 | CC Extrinsic component of membrane | 0.037 | 0.018 | 7 |
| Cluster 3 | ALS-SNs | GO:CC | GO:0031312 | CC Extrinsic component of organelle membrane | 0.016 | 0.025 | 3 |
| Cluster 3 | ALS-SNs | GO:CC | GO:0043083 | CC Synaptic cleft | 0.016 | 0.025 | 3 |
| Cluster 3 | ALS-SNs | GO:CC | GO:0008076 | CC Voltage-gated potassium channel complex | 0.026 | 0.026 | 5 |
| Cluster 3 | ALS-SNs | GO:CC | GO:0031256 | CC Leading edge membrane | 0.037 | 0.032 | 7 |
| Cluster 3 | ALS-SNs | GO:CC | GO:0032809 | CC Neuronal cell body membrane | 0.016 | 0.032 | 3 |
| Cluster 3 | ALS-SNs | GO:CC | GO:0005771 | CC Multivesicular body | 0.021 | 0.041 | 4 |
| Cluster 3 | ALS-SNs | GO:CC | GO:0070069 | CC Cytochrome complex | 0.016 | 0.048 | 3 |
| Cluster 3 | ALS-SNs | GO:MF | GO:0005253 | MF Monoatomic anion channel activity | 0.044 | 0.004 | 8 |
| Cluster 3 | ALS-SNs | GO:MF | GO:0048018 | MF Receptor ligand activity | 0.088 | 0.005 | 16 |
| Cluster 3 | ALS-SNs | GO:MF | GO:0030546 | MF Signaling receptor activator activity | 0.088 | 0.005 | 16 |
| Cluster 3 | ALS-SNs | GO:MF | GO:0160041 | MF Neuropeptide activity | 0.022 | 0.009 | 4 |
| Cluster 3 | ALS-SNs | GO:MF | GO:0015267 | MF Channel activity | 0.082 | 0.009 | 15 |
| Cluster 3 | ALS-SNs | GO:MF | GO:0008528 | MF G protein-coupled peptide receptor activity | 0.038 | 0.010 | 7 |
| Cluster 3 | ALS-SNs | GO:MF | GO:0001653 | MF Peptide receptor activity | 0.038 | 0.013 | 7 |
| Cluster 3 | ALS-SNs | GO:MF | GO:0030594 | MF Neurotransmitter receptor activity | 0.033 | 0.014 | 6 |
| Cluster 3 | ALS-SNs | GO:MF | GO:0031489 | MF Myosin v binding | 0.016 | 0.017 | 3 |
| Cluster 3 | ALS-SNs | GO:MF | GO:0031681 | MF G-protein beta-subunit binding | 0.016 | 0.024 | 3 |
| Cluster 3 | ALS-SNs | GO:MF | GO:0061891 | MF Calcium ion sensor activity | 0.016 | 0.026 | 3 |
| Cluster 3 | ALS-SNs | GO:MF | GO:0140784 | MF Metal ion sensor activity | 0.016 | 0.028 | 3 |
| Cluster 3 | ALS-SNs | GO:MF | GO:0005251 | MF Delayed rectifier potassium channel activity | 0.016 | 0.032 | 3 |
| Cluster 3 | ALS-SNs | GO:MF | GO:0008483 | MF Transaminase activity | 0.016 | 0.038 | 3 |
| Cluster 3 | ALS-SNs | GO:MF | GO:0099529 | MF Neurotransmitter receptor activity involved in regulation of postsynaptic membrane potential | 0.022 | 0.039 | 4 |
| Cluster 3 | ALS-SNs | GO:MF | GO:0015078 | MF Proton transmembrane transporter activity | 0.038 | 0.039 | 7 |
| Cluster 3 | ALS-SNs | GO:MF | GO:0030170 | MF Pyridoxal phosphate binding | 0.022 | 0.039 | 4 |
| Cluster 3 | ALS-SNs | GO:MF | GO:0022853 | MF Active monoatomic ion transmembrane transporter activity | 0.049 | 0.039 | 9 |
| Cluster 3 | ALS-SNs | GO:MF | GO:0042562 | MF Hormone binding | 0.027 | 0.039 | 5 |
| Cluster 3 | ALS-SNs | Reactome | R-HSA-112316 | React Neuronal System | 0.188 | 0.000 | 24 |
| Cluster 3 | ALS-SNs | Reactome | R-HSA-112315 | React Transmission across Chemical Synapses | 0.117 | 0.000 | 15 |
| Cluster 3 | ALS-SNs | Reactome | R-HSA-112314 | React Neurotransmitter receptors and postsynaptic signal transmission | 0.094 | 0.001 | 12 |
| Cluster 3 | ALS-SNs | Reactome | R-HSA-1428517 | React Aerobic respiration and respiratory electron transport | 0.102 | 0.002 | 13 |
| Cluster 3 | ALS-SNs | Reactome | R-HSA-8986944 | React Transcriptional Regulation by MECP2 | 0.047 | 0.011 | 6 |
| Cluster 3 | ALS-SNs | Reactome | R-HSA-1296072 | React Voltage gated Potassium channels | 0.039 | 0.014 | 5 |
| Cluster 3 | ALS-SNs | Reactome | R-HSA-168799 | React Neurotoxicity of clostridium toxins | 0.023 | 0.016 | 3 |
| Cluster 3 | ALS-SNs | Reactome | R-HSA-1296071 | React Potassium Channels | 0.055 | 0.016 | 7 |
| Cluster 3 | ALS-SNs | Reactome | R-HSA-500792 | React GPCR ligand binding | 0.117 | 0.018 | 15 |
| Cluster 3 | ALS-SNs | Reactome | R-HSA-611105 | React Respiratory electron transport | 0.063 | 0.036 | 8 |

**Suppl Table 3.** Distribution of samples with the *C9orf72* expansion and genetic variants in *SOD1* across ALS subtypes.

| Genetic variation | ALS subtype | Number of samples | Percentage (%) |
|---|---|---|---|
| C9orf72 | ALS-Neu | 21 | 0.42 |
| C9orf72 | ALS-OxA | 11 | 0.22 |
| C9orf72 | ALS-SNs | 18 | 0.36 |
| SOD1 | ALS-Neu | 2 | 0.29 |
| SOD1 | ALS-OxA | 1 | 0.14 |
| SOD1 | ALS-SNs | 3 | 0.50 |

**Suppl Table 4.** Summary of classification models for the prediction of ALS subtypes after model improvement and feature selection.

| Training | | | | | | | Evaluation | | | | | |
|---|---|---|---|---|---|---|---|---|---|---|---|---|
| ALS Subtype | Subtype samples | Other samples | # Features | Accuracy (%) | Sensitivity | Specificity | Subtype samples | Other samples | Accuracy (%) | AUC | Sensitivity | Specificity |
| Neu | 76 | 181 | 60 | 96 | 0.95 | 0.97 | 28 | 84 | 92 | 0.98 | 0.89 | 0.95 |
| OxA | 49 | 208 | 79 | 93 | 0.73 | 0.98 | 25 | 87 | 97 | 0.99 | 0.96 | 0.98 |
| SNs | 132 | 125 | 136 | 94 | 0.96 | 0.91 | 59 | 53 | 96 | 0.99 | 0.98 | 0.94 |

**Suppl Table 5.** Candidate genes for brain-based classifiers and blood based-classifiers.

| Brain models | | | Blood models | | |
|---|---|---|---|---|---|
| Symbol | Mean Decrease Accuracy | Cluster | Symbol | Mean Decrease Accuracy | Cluster |
| LGALS3BP | 17.592 | Neu | LGALS3BP | 13.281 | Neu |
| RPS6KA1 | 16.861 | Neu | ELOVL1 | 12.624 | Neu |
| TMCC3 | 15.987 | Neu | CD9 | 12.523 | Neu |
| TREM2 | 15.327 | Neu | SLC45A3 | 12.167 | Neu |
| CD9 | 15.137 | Neu | SPP1 | 12.150 | Neu |
| OLR1 | 13.783 | Neu | ENPP2 | 12.142 | Neu |
| MOG | 13.755 | Neu | RASGRP3 | 12.029 | Neu |
| HAVCR2 | 13.530 | Neu | TMCC3 | 11.959 | Neu |
| CLDN11 | 12.296 | Neu | LIPA | 11.608 | Neu |
| PIP4K2A | 11.793 | Neu | SLC25A13 | 11.539 | Neu |
| LIPA | 11.444 | Neu | PLD1 | 11.301 | Neu |
| TMC6 | 10.257 | Neu | NPC1 | 11.260 | Neu |
| CAPN3 | 9.629 | Neu | OLR1 | 11.116 | Neu |
| SPATA13 | 9.262 | Neu | TREM2 | 11.065 | Neu |
| NPC1 | 9.255 | Neu | TMC6 | 10.920 | Neu |
| SLC45A3 | 9.143 | Neu | SLC35D2 | 10.771 | Neu |
| FCGR1A | 9.048 | Neu | PIP4K2A | 10.683 | Neu |
| ELOVL1 | 8.854 | Neu | NEK3 | 10.234 | Neu |
| LACC1 | 8.840 | Neu | THEMIS2 | 9.873 | Neu |
| RASGRP3 | 8.745 | Neu | SPATA13 | 9.725 | Neu |
| SLC25A13 | 8.722 | Neu | SLAIN1 | 9.722 | Neu |
| SLC5A11 | 8.622 | Neu | UGT8 | 9.409 | Neu |
| TBC1D12 | 8.596 | Neu | FCGR3A | 9.140 | Neu |
| ENPP2 | 8.594 | Neu | C1QC | 9.061 | Neu |
| THEMIS2 | 8.541 | Neu | UBA7 | 8.955 | Neu |
| TMEM140 | 8.176 | Neu | ACSL1 | 8.955 | Neu |
| ENSG00000258844 | 8.093 | Neu | TLR1 | 8.734 | Neu |
| ALOX5AP | 8.062 | Neu | ALOX5AP | 8.555 | Neu |
| BVES | 7.160 | Neu | LACC1 | 8.100 | Neu |
| SLC35D2 | 7.089 | Neu | DAPP1 | 8.097 | Neu |
| C1QC | 6.841 | Neu | SERPINA1 | 8.040 | Neu |
| OR9A2 | 6.630 | Neu | ZNF708 | 7.660 | Neu |
| UBA7 | 6.618 | Neu | REEP3 | 7.419 | Neu |

| | | | | | |
|---|---|---|---|---|---|
| TGFBR1 | 6.527 | Neu | TGFBR1 | 7.366 | Neu |
| FCER1G | 6.324 | Neu | ADGRE2 | 6.924 | Neu |
| SELENOP | 6.296 | Neu | KCNG1 | 3.205 | Neu |
| NKX6-2 | 6.089 | Neu | TSPAN4 | 31.638 | OxA |
| PLD1 | 5.972 | Neu | TUBA1A | 28.229 | OxA |
| SERPINA1 | 5.712 | Neu | KDM6B | 24.981 | OxA |
| NEK3 | 5.656 | Neu | PITPNM2 | 20.463 | OxA |
| FGF1 | 5.528 | Neu | SNCA | 19.144 | OxA |
| TLR1 | 5.506 | Neu | PALLD | 16.041 | OxA |
| CFTR | 5.384 | Neu | TNRC18 | 13.912 | OxA |
| CLDND1 | 5.276 | Neu | SLC29A4 | 13.685 | OxA |
| SLAIN1 | 5.262 | Neu | BRD4 | 13.468 | OxA |
| LINC00639 | 5.202 | Neu | COPS4 | 13.378 | OxA |
| SILC1 | 5.157 | Neu | SPOCK3 | 13.082 | OxA |
| ADGRE2 | 4.977 | Neu | ANKRD11 | 12.407 | OxA |
| CDH19 | 4.323 | Neu | RFX1 | 11.975 | OxA |
| SLC44A1 | 4.261 | Neu | NPRL3 | 11.015 | OxA |
| SIGLEC22P | 3.664 | Neu | KMT2D | 10.096 | OxA |
| REEP3 | 3.497 | Neu | SUCLA2 | 9.998 | OxA |
| LRRC63 | 3.326 | Neu | LTBP3 | 8.599 | OxA |
| ZNF708 | 2.873 | Neu | SBK1 | 8.133 | OxA |
| C21orf91 | 2.514 | Neu | RIPOR1 | 7.477 | OxA |
| LIMS1 | 2.368 | Neu | TNK2 | 6.158 | OxA |
| PLPPR3 | 1.999 | Neu | TMEM14A | 6.047 | OxA |
| KCNG1 | 1.916 | Neu | CLYBL | 4.475 | OxA |
| S100A3 | 0.858 | Neu | NHSL1 | 3.712 | OxA |
| PMP2 | -0.162 | Neu | FAM220A | 1.761 | OxA |
| CAPN15 | 17.648 | OxA | CLCF1 | 1.468 | OxA |
| LAMA5 | 17.161 | OxA | TLN1 | 14.925 | SNs |
| ITGA10 | 16.473 | OxA | PTBP1 | 14.496 | SNs |
| ZIC2 | 13.536 | OxA | CFLAR | 13.248 | SNs |
| TUBA1A | 13.381 | OxA | PXN | 12.802 | SNs |
| AFAP1L1 | 12.726 | OxA | PARP4 | 12.681 | SNs |
| ENSG00000256443 | 11.703 | OxA | WASF2 | 12.314 | SNs |
| SHANK3 | 11.562 | OxA | PALD1 | 12.242 | SNs |
| PITPNM2 | 11.289 | OxA | IKBKB | 12.032 | SNs |
| RAP1GDS1 | 10.937 | OxA | CREB3L2 | 11.743 | SNs |
| ADAMTS10 | 10.758 | OxA | TEP1 | 11.419 | SNs |
| PRELP | 10.428 | OxA | RIN3 | 11.239 | SNs |
| KDM6B | 10.171 | OxA | GUCY1B1 | 10.543 | SNs |
| KCNS3 | 9.851 | OxA | TRIM56 | 10.426 | SNs |
| ZFHX2 | 9.244 | OxA | HMG20B | 10.239 | SNs |
| TMEFF2 | 9.130 | OxA | MINDY1 | 9.497 | SNs |
| PIP5K1C | 8.847 | OxA | ARHGAP45 | 9.482 | SNs |
| PKD1P6 | 8.672 | OxA | PAK1 | 9.374 | SNs |
| WIZ | 8.553 | OxA | SLC4A2 | 9.293 | SNs |
| SLC29A4 | 8.261 | OxA | MAP3K11 | 9.290 | SNs |
| PLEKHG5 | 8.152 | OxA | NCALD | 9.197 | SNs |
| EMX2OS | 8.133 | OxA | OTUD7B | 9.108 | SNs |
| RBPMS | 7.940 | OxA | CCDC88B | 9.095 | SNs |
| ADAM33 | 7.914 | OxA | SH3BP4 | 9.079 | SNs |
| ZIC5 | 7.406 | OxA | CARD8 | 8.821 | SNs |
| RIPOR3 | 7.328 | OxA | HSPA2 | 8.813 | SNs |
| ADGRA2 | 7.028 | OxA | SLC12A7 | 8.703 | SNs |
| SKI | 6.963 | OxA | RRBP1 | 8.678 | SNs |
| DUSP8 | 6.960 | OxA | PADI2 | 8.635 | SNs |
| ENSG00000236760 | 6.525 | OxA | NAP1L5 | 8.542 | SNs |
| ENSG00000285106 | 5.758 | OxA | LRP10 | 8.534 | SNs |
| EVC2 | 5.658 | OxA | CSF1 | 8.212 | SNs |
| ANKRD11 | 5.620 | OxA | CAMK4 | 8.116 | SNs |
| QRICH2 | 5.618 | OxA | CLASRP | 8.078 | SNs |
| SPOCK3 | 5.520 | OxA | SCARB1 | 7.878 | SNs |
| LHX2 | 5.434 | OxA | UQCRH | 7.845 | SNs |
| RABGEF1P1 | 5.266 | OxA | TAFA1 | 7.844 | SNs |
| COPS4 | 5.144 | OxA | SFT2D2 | 7.759 | SNs |
| SPEG | 4.878 | OxA | MPST | 7.696 | SNs |
| FBXL18 | 4.836 | OxA | FTL | 7.229 | SNs |
| CNTFR | 4.802 | OxA | PLK2 | 7.074 | SNs |
| BRD4 | 4.778 | OxA | CTDSP1 | 6.955 | SNs |
| ZNF594-DT | 4.653 | OxA | PLIN3 | 6.950 | SNs |
| MAST1 | 4.646 | OxA | IL17RA | 6.898 | SNs |

| | | |
|---|---|---|
| CNTNAP4 | 4.562 | OxA |
| RFX1 | 4.275 | OxA |
| ENSG00000283529 | 4.163 | OxA |
| DOCK6 | 4.112 | OxA |
| TNRC18 | 3.963 | OxA |
| ELFN2 | 3.764 | OxA |
| ENSG00000278864 | 3.756 | OxA |
| SETD1B | 3.626 | OxA |
| LRFN1 | 3.573 | OxA |
| SUCLA2 | 3.398 | OxA |
| KMT2D | 3.287 | OxA |
| GRIK5 | 3.262 | OxA |
| WNK2 | 3.230 | OxA |
| ZFPM1 | 3.197 | OxA |
| FOXP4 | 3.101 | OxA |
| CLYBL | 3.016 | OxA |
| TNK2 | 2.989 | OxA |
| ADGRB1 | 2.946 | OxA |
| RIPOR1 | 2.900 | OxA |
| NPRL3 | 2.721 | OxA |
| NFATC4 | 2.703 | OxA |
| SAMD14 | 2.677 | OxA |
| SBK1 | 2.557 | OxA |
| ZSWIM9 | 2.382 | OxA |
| ZNF316 | 2.007 | OxA |
| ENSG00000256542 | 1.244 | OxA |
| TMEM14A | 1.042 | OxA |
| LTBP3 | 0.991 | OxA |
| NHSL1 | 0.876 | OxA |
| LMTK3 | 0.872 | OxA |
| PCDHGA12 | 0.428 | OxA |
| COL18A1-AS1 | 0.160 | OxA |
| CLCF1 | -0.307 | OxA |
| ENSG00000282160 | -1.085 | OxA |
| CIMAP1D | -1.925 | OxA |
| TLN1 | 12.061 | SNs |
| PARP4 | 11.261 | SNs |
| PTBP1 | 11.193 | SNs |
| PXN | 10.983 | SNs |
| WASF2 | 10.894 | SNs |
| CFLAR | 9.879 | SNs |
| RREB1 | 9.654 | SNs |
| PALD1 | 9.625 | SNs |
| CREB3L2 | 9.377 | SNs |
| FLI1 | 9.352 | SNs |
| LAMB2 | 9.298 | SNs |
| HMG20B | 9.054 | SNs |
| NXN | 8.818 | SNs |
| SH3BP4 | 8.782 | SNs |
| RIN3 | 8.703 | SNs |
| TRIM56 | 8.664 | SNs |
| HSPA2 | 8.619 | SNs |
| MINDY1 | 8.543 | SNs |
| IKBKB | 8.415 | SNs |
| MBD6 | 8.304 | SNs |
| TEP1 | 8.211 | SNs |
| LRP10 | 8.082 | SNs |
| PADI2 | 8.033 | SNs |
| MAL2 | 7.936 | SNs |
| ARHGAP45 | 7.756 | SNs |
| ATP6V1G2 | 7.306 | SNs |
| PAK1 | 7.289 | SNs |
| GUCY1B1 | 7.250 | SNs |
| SLC4A2 | 7.211 | SNs |
| MAP3K11 | 7.134 | SNs |
| RRBP1 | 7.053 | SNs |
| CARD8 | 7.039 | SNs |
| NOTCH4 | 6.995 | SNs |
| HTR2A | 6.986 | SNs |
| DDR1 | 6.959 | SNs |
| CSF1 | 6.874 | SNs |
| FAM81A | 6.887 | SNs |
| RGS3 | 6.749 | SNs |
| MDH1 | 6.748 | SNs |
| SYNE3 | 6.573 | SNs |
| BOK | 6.294 | SNs |
| PLEKHM1 | 6.257 | SNs |
| ARRDC2 | 6.159 | SNs |
| SLC39A10 | 6.055 | SNs |
| SNX10 | 6.014 | SNs |
| ASAH2B | 5.905 | SNs |
| ERBIN | 5.883 | SNs |
| ANXA5 | 5.766 | SNs |
| MPC2 | 5.714 | SNs |
| REST | 5.640 | SNs |
| ZBTB7B | 5.444 | SNs |
| CHD7 | 5.403 | SNs |
| ABHD4 | 5.022 | SNs |
| CEP295NL | 5.019 | SNs |
| ANXA9 | 4.855 | SNs |
| NME1 | 4.810 | SNs |
| ZFP36L2 | 4.681 | SNs |
| TTC38 | 4.493 | SNs |
| ATP5MK | 4.371 | SNs |
| CLIC4 | 4.291 | SNs |
| BCAT1 | 4.062 | SNs |
| TMBIM1 | 3.487 | SNs |
| CHDH | 3.356 | SNs |
| GPRC5B | 3.352 | SNs |
| EPHX1 | 1.655 | SNs |
| DNAAF8 | 0.007 | SNs |

| | | |
|---|---|---|
| ARAP1 | 6.800 | SNs |
| ZFHX3 | 6.754 | SNs |
| CAMK4 | 6.747 | SNs |
| SFT2D2 | 6.675 | SNs |
| OTUD7B | 6.631 | SNs |
| SYT4 | 6.605 | SNs |
| SVOP | 6.303 | SNs |
| RAB3C | 6.243 | SNs |
| NCALD | 6.243 | SNs |
| PLIN3 | 6.080 | SNs |
| TANC1 | 6.066 | SNs |
| SYNE3 | 6.055 | SNs |
| MPST | 6.012 | SNs |
| PLPPR4 | 5.966 | SNs |
| CDH8 | 5.913 | SNs |
| RGS4 | 5.789 | SNs |
| CCDC88B | 5.750 | SNs |
| ATOH8 | 5.711 | SNs |
| ROBO2 | 5.639 | SNs |
| UQCRH | 5.621 | SNs |
| TAFA1 | 5.589 | SNs |
| PHLDB1 | 5.586 | SNs |
| SLC6A9 | 5.570 | SNs |
| NAP1L5 | 5.542 | SNs |
| OLFM3 | 5.513 | SNs |
| GABRA1 | 5.488 | SNs |
| ANXA5 | 5.477 | SNs |
| CTDSP1 | 5.415 | SNs |
| FTL | 5.398 | SNs |
| PLEKHM1 | 5.392 | SNs |
| CA10 | 5.362 | SNs |
| VWC2 | 5.345 | SNs |
| CLASRP | 5.328 | SNs |
| IL17RA | 5.326 | SNs |
| ZCCHC24 | 5.304 | SNs |
| SALL1 | 5.238 | SNs |
| SCARB1 | 5.182 | SNs |
| ARRDC2 | 5.116 | SNs |
| SLC12A7 | 5.101 | SNs |
| OAT | 5.071 | SNs |
| NDUFA5 | 5.066 | SNs |
| SERTM1 | 4.993 | SNs |
| RGS3 | 4.990 | SNs |
| ST6GALNAC5 | 4.952 | SNs |
| PLK2 | 4.937 | SNs |
| MTUS1 | 4.933 | SNs |
| MIRLET7BHG | 4.933 | SNs |
| ZFP36L2 | 4.872 | SNs |
| MAMDC4 | 4.863 | SNs |
| GPR62 | 4.821 | SNs |
| REST | 4.812 | SNs |
| NSG2 | 4.728 | SNs |
| FAM81A | 4.707 | SNs |
| EPHA5 | 4.693 | SNs |
| ASAH2B | 4.688 | SNs |
| SLC39A10 | 4.602 | SNs |
| RFK | 4.582 | SNs |
| PLXNB1 | 4.539 | SNs |
| PLEKHB1 | 4.419 | SNs |
| MDH1 | 4.261 | SNs |
| GDA | 4.218 | SNs |
| BOK | 4.209 | SNs |
| CHD7 | 4.181 | SNs |
| PACSIN3 | 4.065 | SNs |
| ERBIN | 4.058 | SNs |
| SLC2A13 | 4.055 | SNs |
| TTC38 | 3.946 | SNs |
| KCNB2 | 3.852 | SNs |
| NME1 | 3.849 | SNs |
| GPRC5B | 3.847 | SNs |
| MPC2 | 3.815 | SNs |

| | | |
|---|---|---|
| HPCA | 3.808 | SNs |
| LIMD1 | 3.800 | SNs |
| CYP2U1-AS1 | 3.781 | SNs |
| DOCK1 | 3.669 | SNs |
| ZBTB7B | 3.656 | SNs |
| CLIC4 | 3.581 | SNs |
| SNX10 | 3.581 | SNs |
| GAP43 | 3.477 | SNs |
| FAM107A | 3.397 | SNs |
| ABHD4 | 3.382 | SNs |
| CEP295NL | 3.361 | SNs |
| NXPH1 | 3.310 | SNs |
| ANXA9 | 3.281 | SNs |
| CECR2 | 3.261 | SNs |
| C1QL3 | 3.230 | SNs |
| TMBIM1 | 3.093 | SNs |
| KIF1C | 2.914 | SNs |
| ATP5MK | 2.880 | SNs |
| BCAT1 | 2.724 | SNs |
| GRM1 | 2.576 | SNs |
| MTCL2 | 2.546 | SNs |
| CHDH | 2.454 | SNs |
| SYNPR | 2.435 | SNs |
| ADAMTS3 | 2.267 | SNs |
| ZFHX4 | 1.637 | SNs |
| FCGR1CP | 1.615 | SNs |
| EPHX1 | 1.430 | SNs |
| EPHA3 | 1.163 | SNs |
| DNAAF8 | -2.049 | SNs |

**Suppl Table 6.** Candidate genes used as features for the build of the multi-class LDA model.

| Gene ID | WGCNA Module | Symbol |
|---|---|---|
| ENSG00000057704 | turquoise | TMCC3 |
| ENSG00000141524 | turquoise | TMC6 |
| ENSG00000143409 | turquoise | MINDY1 |
| ENSG00000158715 | turquoise | SLC45A3 |
| ENSG00000264522 | turquoise | OTUD7B |
| ENSG00000016391 | grey | CHDH |
| ENSG00000026559 | grey | KCNG1 |
| ENSG00000039523 | grey | RIPOR1 |
| ENSG00000061938 | grey | TNK2 |
| ENSG00000070190 | grey | DAPP1 |
| ENSG00000090975 | grey | PITPNM2 |
| ENSG00000095970 | grey | TREM2 |
| ENSG00000103148 | grey | NPRL3 |
| ENSG00000105483 | grey | CARD8 |
| ENSG00000106799 | grey | TGFBR1 |
| ENSG00000107719 | grey | PALD1 |
| ENSG00000113504 | grey | SLC12A7 |
| ENSG00000118785 | grey | SPP1 |
| ENSG00000125246 | grey | CLYBL |
| ENSG00000127507 | grey | ADGRE2 |
| ENSG00000129116 | grey | PALLD |
| ENSG00000132005 | grey | RFX1 |
| ENSG00000132510 | grey | KDM6B |
| ENSG00000132965 | grey | ALOX5AP |
| ENSG00000135540 | grey | NHSL1 |
| ENSG00000135926 | grey | TMBIM1 |
| ENSG00000141867 | grey | BRD4 |
| ENSG00000143412 | grey | ANXA9 |
| ENSG00000143819 | grey | EPHX1 |
| ENSG00000152518 | grey | ZFP36L2 |
| ENSG00000159189 | grey | C1QC |
| ENSG00000164111 | grey | ANXA5 |
| ENSG00000164638 | grey | SLC29A4 |
| ENSG00000166246 | grey | DNAAF8 |
| ENSG00000167522 | grey | ANKRD11 |

| | | |
|---|---|---|
| ENSG00000167548 | grey | KMT2D |
| ENSG00000167552 | grey | TUBA1A |
| ENSG00000168071 | grey | CCDC88B |
| ENSG00000173391 | grey | OLR1 |
| ENSG00000174125 | grey | TLR1 |
| ENSG00000175505 | grey | CLCF1 |
| ENSG00000177663 | grey | IL17RA |
| ENSG00000178397 | grey | FAM220A |
| ENSG00000180448 | grey | ARHGAP45 |
| ENSG00000182095 | grey | TNRC18 |
| ENSG00000182179 | grey | UBA7 |
| ENSG00000188322 | grey | SBK1 |
| ENSG00000197249 | grey | SERPINA1 |
| ENSG00000203747 | grey | FCGR3A |
| ENSG00000214063 | grey | TSPAN4 |
| ENSG00000014641 | blue | MDH1 |
| ENSG00000061918 | blue | GUCY1B1 |
| ENSG00000177432 | blue | NAP1L5 |

**Suppl Table 7.** Three-way differential gene expression results of ALS subtype-predicted blood samples.

| Gene Symbol | Cluster | Z score | Adj p value |
|---|---|---|---|
| ACYP2 | Neu | 2.39384199 | 0.00403792 |
| DCTD | Neu | 2.24594979 | 0.0056761 |
| NR2F6 | Neu | 3.743842807 | 0.00018037 |
| FH | Neu | 2.286862056 | 0.0051658 |
| GPX7 | Neu | 3.19830369 | 0.00063343 |
| HOXB4 | Neu | 2.111972715 | 0.00772729 |
| HNRNPM | Neu | 3.028674708 | 0.00093611 |
| NME2 | Neu | 2.917110484 | 0.00121029 |
| SLC22A5 | Neu | 3.161803699 | 0.00068896 |
| SOX12 | Neu | 1.98958641 | 0.01024268 |
| XBP1 | Neu | 1.990553764 | 0.01021989 |
| ZNF132 | Neu | 2.314412177 | 0.00484828 |
| ARHGEF7 | Neu | 2.322141092 | 0.00476276 |
| GDF11 | Neu | 2.530748366 | 0.00294613 |
| MRPS30 | Neu | 2.250743008 | 0.0056138 |
| PPP1R13B | Neu | 2.038759803 | 0.00914619 |
| IGHV1-46 | Neu | 2.851667267 | 0.00140713 |
| TRBV18 | Neu | 2.05422859 | 0.00882615 |
| TRAV26-1 | Neu | 2.729251484 | 0.0018653 |
| IGLV4-69 | Neu | 2.783422132 | 0.00164656 |
| IGLV4-60 | Neu | 2.783999226 | 0.00164437 |
| IGKV2-30 | Neu | 2.453501066 | 0.00351965 |
| NOP53 | Neu | 2.073448786 | 0.00844406 |
| PLEKHA8P1 | Neu | 2.324838705 | 0.00473327 |
| FKBP11 | Neu | 3.209889695 | 0.00061675 |
| ULK4 | Neu | 1.987835598 | 0.01028406 |
| STRBP | Neu | 2.08901161 | 0.00814683 |
| AVEN | Neu | 2.671669 | 0.00212976 |
| ZNF286A | Neu | 2.365784679 | 0.0043074 |
| PALB2 | Neu | 2.233372415 | 0.00584289 |
| GOLGA2P10 | Neu | 2.260424965 | 0.00549003 |
| STARD5 | Neu | 4.028756264 | 9.36E-05 |
| ITM2C | Neu | 2.165808479 | 0.0068264 |
| CDKN2AIPNL | Neu | 2.331231318 | 0.00466411 |
| KTI12 | Neu | 3.445042209 | 0.00035889 |
| SELENOM | Neu | 2.372916097 | 0.00423725 |
| ZFPM1 | Neu | 2.099524495 | 0.00795198 |
| KANK3 | Neu | 2.289062647 | 0.0051397 |
| C2orf74-DT | Neu | 2.293577102 | 0.00508655 |
| MZT2A | Neu | 2.018112811 | 0.00959151 |
| TPT1-AS1 | Neu | 2.122221824 | 0.00754707 |
| ACP1 | OxA | 2.396672587 | 0.00401169 |
| ACTL6A | OxA | 2.518827199 | 0.00302812 |
| ADD3 | OxA | 2.504987126 | 0.00312617 |
| ANK3 | OxA | 2.369525574 | 0.00427046 |
| BIRC3 | OxA | 3.882446101 | 0.00013109 |

| | | | |
|---|---|---|---|
| FASLG | OxA | 4.048391567 | 8.95E-05 |
| ARL1 | OxA | 3.965968885 | 0.00010815 |
| ASNS | OxA | 2.346160304 | 0.0045065 |
| ATM | OxA | 3.560496159 | 0.00027511 |
| ATP5F1C | OxA | 1.98281362 | 0.01040367 |
| ATP5PB | OxA | 2.916369022 | 0.00121236 |
| ATP5PF | OxA | 3.504013597 | 0.00031332 |
| ATP5PO | OxA | 3.289948657 | 0.00051292 |
| ATR | OxA | 3.452997426 | 0.00035237 |
| BCL2 | OxA | 2.835406611 | 0.00146081 |
| C4BPA | OxA | 2.221080819 | 0.00601062 |
| CAMLG | OxA | 2.089973955 | 0.00812879 |
| CASP6 | OxA | 3.448471012 | 0.00035606 |
| CASP10 | OxA | 2.291356052 | 0.00511263 |
| CCNC | OxA | 2.490561967 | 0.00323175 |
| CCT6A | OxA | 2.904568641 | 0.00124575 |
| MS4A1 | OxA | 2.369774372 | 0.00426801 |
| CD47 | OxA | 2.325821875 | 0.00472257 |
| CD48 | OxA | 2.800795599 | 0.00158199 |
| CD69 | OxA | 2.023581089 | 0.0094715 |
| LRBA | OxA | 2.670950312 | 0.00213329 |
| CDC5L | OxA | 2.704145399 | 0.00197631 |
| SEPTIN7 | OxA | 2.162652033 | 0.00687619 |
| CDC27 | OxA | 2.183266645 | 0.00655743 |
| CD52 | OxA | 2.213196748 | 0.00612073 |
| CENPC | OxA | 2.694894888 | 0.00201885 |
| CLNS1A | OxA | 2.315212756 | 0.00483935 |
| COPB1 | OxA | 2.240833381 | 0.00574337 |
| COX6C | OxA | 5.499427678 | 3.17E-06 |
| COX7C | OxA | 3.473120588 | 0.00033642 |
| COX11 | OxA | 1.995306379 | 0.01010866 |
| CSE1L | OxA | 2.729374182 | 0.00186477 |
| CTLA4 | OxA | 2.359310559 | 0.00437209 |
| DBI | OxA | 2.657123047 | 0.0022023 |
| DDX1 | OxA | 4.364209221 | 4.32E-05 |
| RCAN1 | OxA | 2.192922918 | 0.00641323 |
| GPR183 | OxA | 3.187965559 | 0.00064869 |
| S1PR1 | OxA | 2.118994159 | 0.00760337 |
| PHC1 | OxA | 2.422354914 | 0.00378133 |
| EIF2S1 | OxA | 2.667433807 | 0.00215063 |
| EIF4A2 | OxA | 3.317685152 | 0.00048119 |
| EIF5 | OxA | 2.476612634 | 0.00333724 |
| ELK4 | OxA | 4.064210015 | 8.63E-05 |
| EPHX2 | OxA | 2.46520537 | 0.00342606 |
| FANCF | OxA | 2.722761096 | 0.00189338 |
| FDX1 | OxA | 2.90571186 | 0.00124248 |
| FER | OxA | 3.215941866 | 0.00060822 |
| FKBP3 | OxA | 5.585319079 | 2.60E-06 |
| FOXO1 | OxA | 2.597789604 | 0.0025247 |
| KDSR | OxA | 2.934004558 | 0.00116411 |
| GART | OxA | 2.447549395 | 0.00356821 |
| GBP3 | OxA | 3.501497013 | 0.00031514 |
| GLS | OxA | 2.078826972 | 0.00834013 |
| GPR18 | OxA | 2.488952909 | 0.00324375 |
| GRSF1 | OxA | 3.120424846 | 0.00075784 |
| GTF2F2 | OxA | 2.480097358 | 0.00331057 |
| HELLS | OxA | 2.085094763 | 0.00822063 |
| HMGB1 | OxA | 2.287031374 | 0.00516379 |
| HNRNPA2B1 | OxA | 2.835272077 | 0.00146126 |
| HNRNPH1 | OxA | 2.684653954 | 0.00206703 |
| HNRNPH3 | OxA | 2.017135989 | 0.00961311 |
| HSP90AA1 | OxA | 6.108823256 | 7.78E-07 |
| HSPD1 | OxA | 2.788336017 | 0.00162804 |
| IL6ST | OxA | 2.879282568 | 0.00132044 |
| EIF3E | OxA | 4.632334669 | 2.33E-05 |
| ITGA6 | OxA | 2.737987885 | 0.00182815 |
| ITGA4 | OxA | 2.829295185 | 0.00148151 |
| ITGB1 | OxA | 2.564201473 | 0.00272771 |
| ITPR2 | OxA | 2.570928627 | 0.00268579 |
| KCNA3 | OxA | 3.444635766 | 0.00035922 |
| KIF2A | OxA | 3.427328936 | 0.00037383 |

| | | | |
|---|---|---|---|
| KLRB1 | OxA | 2.727795047 | 0.00187157 |
| KTN1 | OxA | 3.626686111 | 0.00023622 |
| LMAN1 | OxA | 2.190480679 | 0.0064494 |
| LMO7 | OxA | 2.680492697 | 0.00208693 |
| LNPEP | OxA | 2.728118326 | 0.00187017 |
| MAT2A | OxA | 3.392443015 | 0.0004051 |
| DNAJB9 | OxA | 2.563240336 | 0.00273376 |
| MDH1 | OxA | 4.031974871 | 9.29E-05 |
| MAP3K4 | OxA | 2.907634386 | 0.00123699 |
| KMT2A | OxA | 3.088912926 | 0.00081487 |
| MLLT3 | OxA | 1.9830352 | 0.01039836 |
| NR3C2 | OxA | 3.021591113 | 0.0009515 |
| MSH2 | OxA | 4.614250646 | 2.43E-05 |
| MTR | OxA | 4.990143571 | 1.02E-05 |
| TRIM37 | OxA | 2.239088911 | 0.00576648 |
| NDUFA4 | OxA | 2.839243601 | 0.00144796 |
| NDUFA5 | OxA | 2.097280247 | 0.00799318 |
| NDUFA6 | OxA | 2.58742519 | 0.00258568 |
| NDUFB5 | OxA | 2.008162443 | 0.00981381 |
| NDUFS4 | OxA | 3.016289534 | 0.00096319 |
| NDUFS5 | OxA | 4.335513588 | 4.62E-05 |
| NFX1 | OxA | 2.636641644 | 0.00230865 |
| NUCB2 | OxA | 3.450946358 | 0.00035404 |
| NUP88 | OxA | 2.738053138 | 0.00182788 |
| ORC4 | OxA | 2.63055503 | 0.00234123 |
| PAM | OxA | 2.146979649 | 0.00712886 |
| PCM1 | OxA | 2.801625092 | 0.00157897 |
| PFDN4 | OxA | 3.80774015 | 0.00015569 |
| PFDN5 | OxA | 2.295864874 | 0.00505982 |
| PIGC | OxA | 2.188778099 | 0.00647473 |
| PIGF | OxA | 3.082513323 | 0.00082696 |
| PIGH | OxA | 1.986950829 | 0.01030503 |
| PIK3R1 | OxA | 2.449281447 | 0.00355401 |
| PLCG1 | OxA | 2.210463473 | 0.00615937 |
| PLRG1 | OxA | 3.020639497 | 0.00095359 |
| PPA1 | OxA | 4.815067092 | 1.53E-05 |
| PPP1CC | OxA | 2.911277742 | 0.00122665 |
| PPP1R8 | OxA | 2.589426101 | 0.00257379 |
| PPP2R1B | OxA | 2.130207277 | 0.00740957 |
| PPP2R5C | OxA | 2.010396833 | 0.00976345 |
| PRKACB | OxA | 3.908644717 | 0.00012341 |
| PKIA | OxA | 2.999458635 | 0.00100125 |
| PRKCA | OxA | 2.235354954 | 0.00581628 |
| EIF2AK2 | OxA | 2.326275903 | 0.00471763 |
| PSMA2 | OxA | 2.190509595 | 0.00644897 |
| PSMA3 | OxA | 2.60802919 | 0.00246587 |
| PSMA4 | OxA | 2.889631423 | 0.00128934 |
| PSMA6 | OxA | 2.712309462 | 0.0019395 |
| PSMC6 | OxA | 2.355314061 | 0.00441251 |
| PTGER2 | OxA | 4.157660154 | 6.96E-05 |
| PTS | OxA | 2.438117312 | 0.00364655 |
| PEX2 | OxA | 2.256442255 | 0.00554061 |
| RABGGTB | OxA | 2.817334708 | 0.00152288 |
| RAD51C | OxA | 2.958013782 | 0.0011015 |
| RANBP2 | OxA | 2.225223473 | 0.00595356 |
| RAP1GDS1 | OxA | 2.647652503 | 0.00225085 |
| RARS1 | OxA | 3.567189835 | 0.0002709 |
| RASGRF2 | OxA | 2.809847877 | 0.00154936 |
| RBBP4 | OxA | 2.592285897 | 0.0025569 |
| RBL1 | OxA | 2.780401761 | 0.00165805 |
| RECQL | OxA | 3.178249106 | 0.00066336 |
| RFC1 | OxA | 2.147572447 | 0.00711914 |
| RORA | OxA | 2.399742604 | 0.00398343 |
| RPL7 | OxA | 4.973166605 | 1.06E-05 |
| RPL11 | OxA | 5.289126621 | 5.14E-06 |
| RPL17 | OxA | 5.916453562 | 1.21E-06 |
| RPL22 | OxA | 4.631089351 | 2.34E-05 |
| RPL26 | OxA | 5.121178628 | 7.57E-06 |
| RPL27 | OxA | 3.919859158 | 0.00012027 |
| RPL32 | OxA | 2.830255109 | 0.00147824 |
| RPL34 | OxA | 4.92610358 | 1.19E-05 |

| | | | |
|---|---|---|---|
| RPL37 | OxA | 3.400452129 | 0.00039769 |
| RPL41 | OxA | 2.267135052 | 0.00540586 |
| RPS10 | OxA | 3.316660713 | 0.00048232 |
| RPS24 | OxA | 4.383256523 | 4.14E-05 |
| RRM1 | OxA | 2.723300214 | 0.00189104 |
| SATB1 | OxA | 2.709590617 | 0.00195168 |
| MSMO1 | OxA | 2.339596881 | 0.00457513 |
| SET | OxA | 2.450023928 | 0.00354794 |
| SFPQ | OxA | 2.325526015 | 0.00472579 |
| SRSF1 | OxA | 2.102866089 | 0.00789103 |
| SRSF2 | OxA | 2.023136434 | 0.00948121 |
| SRSF3 | OxA | 2.123313126 | 0.00752813 |
| SRSF6 | OxA | 2.741905006 | 0.00181174 |
| SRSF7 | OxA | 3.215019018 | 0.00060951 |
| TRA2B | OxA | 4.548770298 | 2.83E-05 |
| SKP2 | OxA | 2.109922873 | 0.00776385 |
| SLC5A3 | OxA | 2.168210954 | 0.00678874 |
| SNRPA1 | OxA | 2.281154717 | 0.00523414 |
| SNRPB2 | OxA | 1.989386134 | 0.0102474 |
| SNRPD1 | OxA | 3.277089206 | 0.00052834 |
| SNRPD2 | OxA | 2.671600481 | 0.0021301 |
| SNRPE | OxA | 2.176972291 | 0.00665316 |
| SNRPF | OxA | 2.210296347 | 0.00616174 |
| SNRPG | OxA | 2.024853975 | 0.00944378 |
| SNX2 | OxA | 2.025284015 | 0.00943444 |
| SPTBN1 | OxA | 2.523582515 | 0.00299514 |
| SQLE | OxA | 2.410415912 | 0.00388673 |
| SRP14 | OxA | 2.104639022 | 0.00785889 |
| SRP72 | OxA | 4.077738816 | 8.36E-05 |
| TAF9 | OxA | 4.168841896 | 6.78E-05 |
| TAF11 | OxA | 3.725678369 | 0.00018807 |
| MAP3K7 | OxA | 2.441987624 | 0.0036142 |
| TARBP1 | OxA | 3.0616783 | 0.0008676 |
| TARS1 | OxA | 2.647843252 | 0.00224987 |
| TBCA | OxA | 2.047696057 | 0.00895992 |
| TCEA1 | OxA | 1.98097436 | 0.01044782 |
| TFAM | OxA | 2.34394058 | 0.0045296 |
| TFRC | OxA | 3.016082766 | 0.00096365 |
| TGFBR3 | OxA | 3.017968104 | 0.00095947 |
| TIAM1 | OxA | 2.070127066 | 0.00850889 |
| TMPO | OxA | 2.965337515 | 0.00108308 |
| TOP2B | OxA | 3.898257405 | 0.0001264 |
| TPP2 | OxA | 2.428317319 | 0.00372978 |
| HSP90B1 | OxA | 2.529570327 | 0.00295413 |
| TTC3 | OxA | 3.579705364 | 0.00026321 |
| TXK | OxA | 2.840654346 | 0.00144326 |
| UBE2G2 | OxA | 2.085938169 | 0.00820468 |
| UBE2N | OxA | 2.792975633 | 0.00161074 |
| SUMO1 | OxA | 2.223997981 | 0.00597038 |
| UBP1 | OxA | 3.241938005 | 0.00057288 |
| UGP2 | OxA | 2.698938423 | 0.00200015 |
| UQCRB | OxA | 3.217199182 | 0.00060646 |
| UQCRH | OxA | 2.327692919 | 0.00470226 |
| XPO1 | OxA | 2.197211413 | 0.00635022 |
| ZNF23 | OxA | 3.091118664 | 0.00081074 |
| ZBTB25 | OxA | 2.924942776 | 0.00118866 |
| ZNF75A | OxA | 2.899254807 | 0.00126109 |
| ZNF84 | OxA | 4.125401026 | 7.49E-05 |
| ZNF146 | OxA | 3.785533326 | 0.00016386 |
| ZNF202 | OxA | 2.534077547 | 0.00292363 |
| ZNF204P | OxA | 3.004744074 | 0.00098914 |
| ZNF224 | OxA | 2.692088434 | 0.00203194 |
| ZNF226 | OxA | 4.228955425 | 5.90E-05 |
| CCDC6 | OxA | 2.68505952 | 0.0020651 |
| MKKS | OxA | 3.629044551 | 0.00023494 |
| NRIP1 | OxA | 3.835044228 | 0.0001462 |
| CMAHP | OxA | 2.434863965 | 0.00367397 |
| DYRK2 | OxA | 3.606714008 | 0.00024734 |
| SMARCA5 | OxA | 2.868142087 | 0.00135475 |
| HAT1 | OxA | 2.694096658 | 0.00202257 |
| API5 | OxA | 4.226312543 | 5.94E-05 |

| | | | |
|---|---|---|---|
| AGPS | OxA | 2.732656022 | 0.00185073 |
| CGGBP1 | OxA | 2.182417054 | 0.00657027 |
| PSMG1 | OxA | 2.280905878 | 0.00523714 |
| EIF3J | OxA | 2.218438591 | 0.0060473 |
| CDC23 | OxA | 2.520734974 | 0.00301485 |
| B4GALT4 | OxA | 2.264437954 | 0.00543954 |
| EED | OxA | 2.166314594 | 0.00681845 |
| SUCLA2 | OxA | 2.228602529 | 0.00590741 |
| INPP4B | OxA | 4.096368085 | 8.01E-05 |
| SOCS2 | OxA | 3.061359296 | 0.00086824 |
| FUBP1 | OxA | 2.419441288 | 0.00380679 |
| CDC16 | OxA | 3.903480824 | 0.00012489 |
| PRP4K | OxA | 2.74056125 | 0.00181735 |
| RAB29 | OxA | 2.595550297 | 0.00253776 |
| SLC7A6 | OxA | 3.601870622 | 0.00025011 |
| USP14 | OxA | 2.336295262 | 0.00461004 |
| SMC3 | OxA | 2.528322152 | 0.00296263 |
| RABEP1 | OxA | 3.293680569 | 0.00050853 |
| EBAG9 | OxA | 2.918458405 | 0.00120654 |
| BUB3 | OxA | 3.778977556 | 0.00016635 |
| SLC16A7 | OxA | 2.365641608 | 0.00430882 |
| NOLC1 | OxA | 2.3402097 | 0.00456868 |
| NOG | OxA | 2.34820177 | 0.00448537 |
| SRSF11 | OxA | 2.5666363 | 0.00271246 |
| PTER | OxA | 2.565749534 | 0.00271801 |
| TRIP11 | OxA | 2.643311501 | 0.00227347 |
| HMGN3 | OxA | 4.476927286 | 3.33E-05 |
| GTF3C4 | OxA | 2.875926841 | 0.00133068 |
| RPL23 | OxA | 5.392430545 | 4.05E-06 |
| PPIG | OxA | 2.526207038 | 0.0029771 |
| OTOF | OxA | 2.435775629 | 0.00366627 |
| ATG5 | OxA | 2.091575364 | 0.00809887 |
| ATP6V1G1 | OxA | 2.136317981 | 0.00730604 |
| CHD1L | OxA | 2.903665091 | 0.00124835 |
| MINPP1 | OxA | 2.679494585 | 0.00209173 |
| CLOCK | OxA | 2.343163236 | 0.00453771 |
| KIF20B | OxA | 2.659711123 | 0.00218922 |
| RNF7 | OxA | 2.157616469 | 0.00695638 |
| SKIC3 | OxA | 3.172264791 | 0.00067257 |
| DZIP3 | OxA | 3.076429114 | 0.00083863 |
| EIF5B | OxA | 3.450129265 | 0.00035471 |
| CEP57 | OxA | 2.69616098 | 0.00201298 |
| MATR3 | OxA | 3.567909119 | 0.00027045 |
| TOMM20 | OxA | 5.317169511 | 4.82E-06 |
| C2CD5 | OxA | 3.225871724 | 0.00059447 |
| LRIG2 | OxA | 2.899178917 | 0.00126131 |
| TOMM70 | OxA | 3.797906801 | 0.00015926 |
| TBC1D4 | OxA | 2.374142972 | 0.00422529 |
| SUPT7L | OxA | 2.090668555 | 0.0081158 |
| PUM3 | OxA | 2.5728706 | 0.0026738 |
| EXOG | OxA | 2.547359933 | 0.00283557 |
| NR1D2 | OxA | 2.691573768 | 0.00203435 |
| RBX1 | OxA | 2.014561929 | 0.00967026 |
| HNRNPDL | OxA | 3.722283063 | 0.00018955 |
| DMTF1 | OxA | 1.99841607 | 0.01003654 |
| PIGK | OxA | 2.568037529 | 0.00270372 |
| RAD50 | OxA | 3.712138619 | 0.00019403 |
| FEM1B | OxA | 2.273821363 | 0.00532327 |
| RASGRP1 | OxA | 2.585041807 | 0.00259991 |
| LRPPRC | OxA | 3.374682941 | 0.000422 |
| RBM12 | OxA | 3.593556651 | 0.00025494 |
| YAF2 | OxA | 2.427152497 | 0.00373979 |
| CEBPZ | OxA | 2.850578883 | 0.00141066 |
| LPAR6 | OxA | 3.818685268 | 0.00015182 |
| ZNF256 | OxA | 2.264249179 | 0.0054419 |
| MPHOSPH9 | OxA | 2.265220257 | 0.00542975 |
| MPHOSPH10 | OxA | 3.950239341 | 0.00011214 |
| USPL1 | OxA | 2.158434931 | 0.00694329 |
| PSMD14 | OxA | 3.239706095 | 0.00057583 |
| HNRNPR | OxA | 2.749238159 | 0.0017814 |
| MRPS31 | OxA | 2.048972871 | 0.00893361 |

| | | | |
|---|---|---|---|
| BET1 | OxA | 2.007329196 | 0.00983266 |
| SNAPC5 | OxA | 1.999357581 | 0.0100148 |
| LANCL1 | OxA | 2.950866249 | 0.00111978 |
| TRIM22 | OxA | 2.325672791 | 0.00472419 |
| HMGB1P5 | OxA | 2.052810094 | 0.00885503 |
| BTN3A3 | OxA | 2.601590469 | 0.0025027 |
| ARIH2 | OxA | 1.981521472 | 0.01043467 |
| EMG1 | OxA | 2.685190427 | 0.00206447 |
| EIF3M | OxA | 2.327143482 | 0.00470822 |
| RPP30 | OxA | 2.450609338 | 0.00354316 |
| SLC35A1 | OxA | 2.111671639 | 0.00773265 |
| IFI44 | OxA | 2.604719555 | 0.00248474 |
| SMC2 | OxA | 1.991210924 | 0.01020444 |
| USP16 | OxA | 3.495504311 | 0.00031952 |
| ATP5MG | OxA | 2.226308503 | 0.0059387 |
| CCT8 | OxA | 2.45656314 | 0.00349492 |
| HBS1L | OxA | 2.209570233 | 0.00617205 |
| ZMYND11 | OxA | 2.75941378 | 0.00174015 |
| ZNF271P | OxA | 2.731457651 | 0.00185585 |
| ZNF234 | OxA | 2.044685556 | 0.00902224 |
| MTHFD2 | OxA | 3.721087007 | 0.00019007 |
| WDR3 | OxA | 2.201993091 | 0.00628068 |
| MALT1 | OxA | 3.296678942 | 0.00050503 |
| SUGT1 | OxA | 3.6245926 | 0.00023736 |
| TCERG1 | OxA | 3.517649874 | 0.00030363 |
| SUB1 | OxA | 2.853345066 | 0.0014017 |
| C11orf58 | OxA | 2.67619797 | 0.00210767 |
| SF3A3 | OxA | 1.988726144 | 0.01026299 |
| YWHAQ | OxA | 2.068140568 | 0.0085479 |
| TMED10 | OxA | 2.324733622 | 0.00473442 |
| COPS5 | OxA | 2.346978352 | 0.00449802 |
| METAP2 | OxA | 3.575549262 | 0.00026574 |
| NUDT21 | OxA | 2.98847446 | 0.00102689 |
| CNTRL | OxA | 3.578972614 | 0.00026365 |
| KRR1 | OxA | 3.891537336 | 0.00012837 |
| PSIP1 | OxA | 2.698980927 | 0.00199995 |
| RPL35 | OxA | 2.587299851 | 0.00258643 |
| DUSP12 | OxA | 2.728320766 | 0.0018693 |
| KLF12 | OxA | 3.526097083 | 0.00029779 |
| VPS45 | OxA | 2.406374083 | 0.00392307 |
| MGAT4A | OxA | 3.70084804 | 0.00019914 |
| CBX3 | OxA | 2.324445647 | 0.00473756 |
| EXOSC8 | OxA | 3.821368582 | 0.00015088 |
| MTF2 | OxA | 2.968525004 | 0.00107516 |
| ZNF507 | OxA | 2.109367231 | 0.00777379 |
| AAK1 | OxA | 2.638005987 | 0.00230141 |
| FASTKD2 | OxA | 2.328243658 | 0.00469631 |
| KHDC4 | OxA | 2.044699812 | 0.00902195 |
| ZBTB1 | OxA | 2.469250242 | 0.0033943 |
| RUFY3 | OxA | 1.998844984 | 0.01002663 |
| NCBP2 | OxA | 3.220762709 | 0.0006015 |
| GOLGA8A | OxA | 2.693467606 | 0.0020255 |
| RBM34 | OxA | 2.856878398 | 0.00139034 |
| DOP1A | OxA | 2.632882389 | 0.00232872 |
| TNIK | OxA | 3.691853042 | 0.0002033 |
| CLASP2 | OxA | 2.161689437 | 0.00689145 |
| DCUN1D4 | OxA | 2.144880449 | 0.00716341 |
| WDR43 | OxA | 3.417142006 | 0.0003827 |
| MPRIP | OxA | 2.344138227 | 0.00452753 |
| CEP68 | OxA | 2.07435991 | 0.00842636 |
| MDN1 | OxA | 3.281963441 | 0.00052244 |
| ARL6IP1 | OxA | 2.346220777 | 0.00450588 |
| VPS13A | OxA | 2.702417819 | 0.00198419 |
| ANKRD12 | OxA | 2.442906184 | 0.00360657 |
| DDHD2 | OxA | 2.577687929 | 0.00264431 |
| TSPYL4 | OxA | 3.683622108 | 0.00020719 |
| TASOR | OxA | 2.28520646 | 0.00518553 |
| NEMP1 | OxA | 2.651026036 | 0.00223344 |
| SUN1 | OxA | 2.48155603 | 0.00329947 |
| USP24 | OxA | 3.207481247 | 0.00062018 |
| ICE1 | OxA | 2.480707129 | 0.00330592 |

| | | | |
|---|---|---|---|
| ECPAS | OxA | 2.907032608 | 0.0012387 |
| PPWD1 | OxA | 2.623573043 | 0.00237918 |
| EXOSC2 | OxA | 2.321735567 | 0.00476721 |
| SIRT5 | OxA | 3.59686768 | 0.00025301 |
| MTREX | OxA | 3.193988102 | 0.00063975 |
| ATP6V0A2 | OxA | 3.17383126 | 0.00067014 |
| KLHDC2 | OxA | 2.814324332 | 0.00153347 |
| CD2AP | OxA | 2.44457446 | 0.00359274 |
| SNHG1 | OxA | 3.160529052 | 0.00069099 |
| TNFAIP8 | OxA | 3.571272077 | 0.00026837 |
| ZNF345 | OxA | 3.008218335 | 0.00098125 |
| NEPRO | OxA | 3.495101009 | 0.00031982 |
| POT1 | OxA | 2.375565626 | 0.00421148 |
| THUMPD3 | OxA | 2.452151153 | 0.0035306 |
| ATL3 | OxA | 2.542315565 | 0.0028687 |
| NGDN | OxA | 2.232357339 | 0.00585656 |
| SS18L1 | OxA | 3.371810201 | 0.00042481 |
| EDRF1 | OxA | 3.317823427 | 0.00048103 |
| FGFR1OP2 | OxA | 2.879138842 | 0.00132087 |
| SERBP1 | OxA | 2.20042453 | 0.00630341 |
| ZNF337 | OxA | 3.219223724 | 0.00060364 |
| GIMAP2 | OxA | 6.930147988 | 1.17E-07 |
| KLHL3 | OxA | 2.539277454 | 0.00288883 |
| BLOC1S6 | OxA | 2.154734612 | 0.0070027 |
| FBXO3 | OxA | 2.729660029 | 0.00186355 |
| HIBCH | OxA | 2.733989108 | 0.00184506 |
| GNL3 | OxA | 3.45513859 | 0.00035064 |
| RANBP6 | OxA | 2.331784033 | 0.00465818 |
| CHORDC1 | OxA | 3.683302786 | 0.00020735 |
| DNAJC2 | OxA | 3.546712057 | 0.00028398 |
| UTP25 | OxA | 2.978495223 | 0.00105076 |
| SNX5 | OxA | 2.578480479 | 0.00263949 |
| SESN1 | OxA | 3.305472678 | 0.00049491 |
| ERLEC1 | OxA | 2.278550064 | 0.00526563 |
| PDCD4 | OxA | 2.894934949 | 0.00127369 |
| LSM1 | OxA | 2.151822447 | 0.00704981 |
| LSM3 | OxA | 2.690721677 | 0.00203835 |
| TNRC6A | OxA | 2.158143282 | 0.00694795 |
| MAT2B | OxA | 2.008283182 | 0.00981108 |
| EML4 | OxA | 2.325386325 | 0.00472731 |
| TRIB2 | OxA | 2.984546919 | 0.00103622 |
| MRPL42 | OxA | 3.124551446 | 0.00075067 |
| MRPL13 | OxA | 2.90108046 | 0.0012558 |
| THYN1 | OxA | 2.787954323 | 0.00162947 |
| MRPL15 | OxA | 2.402088487 | 0.00396197 |
| DROSHA | OxA | 2.220387359 | 0.00602022 |
| CLEC2D | OxA | 3.064035075 | 0.00086291 |
| OLA1 | OxA | 2.960443903 | 0.00109536 |
| ICOS | OxA | 3.755462561 | 0.00017561 |
| CNOT7 | OxA | 2.878316365 | 0.00132338 |
| GNL2 | OxA | 2.037386567 | 0.00917516 |
| ANAPC4 | OxA | 2.661052268 | 0.00218247 |
| DNTTIP2 | OxA | 2.972676608 | 0.00106494 |
| ARHGEF3 | OxA | 4.068072851 | 8.55E-05 |
| AK3 | OxA | 2.334899051 | 0.00462489 |
| MTERF3 | OxA | 2.84121735 | 0.00144139 |
| RRP15 | OxA | 2.959531424 | 0.00109766 |
| HDDC2 | OxA | 2.450761385 | 0.00354192 |
| VPS36 | OxA | 2.437438342 | 0.00365226 |
| TVP23B | OxA | 3.018673276 | 0.00095791 |
| MEMO1 | OxA | 2.501777867 | 0.00314936 |
| SEPSECS | OxA | 2.281811697 | 0.00522623 |
| UTP18 | OxA | 2.240197795 | 0.00575178 |
| RMDN1 | OxA | 2.275696473 | 0.00530034 |
| SBDS | OxA | 3.486908204 | 0.00032591 |
| PHF11 | OxA | 2.472592584 | 0.00336827 |
| CRBN | OxA | 2.486829459 | 0.00325965 |
| SS18L2 | OxA | 2.875327156 | 0.00133252 |
| DDX47 | OxA | 2.948679616 | 0.00112543 |
| COX16 | OxA | 4.259045345 | 5.51E-05 |
| GCNT4 | OxA | 2.177966717 | 0.00663794 |

| | | | |
|---|---|---|---|
| PLAC8 | OxA | 2.411018181 | 0.00388134 |
| AMZ2 | OxA | 2.123910612 | 0.00751778 |
| KLRF1 | OxA | 3.749173671 | 0.00017817 |
| HOOK1 | OxA | 2.116357804 | 0.00764966 |
| TMA7 | OxA | 2.132231773 | 0.00737511 |
| COMMD10 | OxA | 3.930484565 | 0.00011736 |
| ANAPC7 | OxA | 1.991623741 | 0.01019474 |
| PCYOX1 | OxA | 2.32182767 | 0.0047662 |
| HACD3 | OxA | 3.151228668 | 0.00070595 |
| TRIAP1 | OxA | 2.760396921 | 0.00173621 |
| CWC15 | OxA | 2.762004346 | 0.0017298 |
| CHMP5 | OxA | 2.512542001 | 0.00307226 |
| TMEM14C | OxA | 2.908539712 | 0.00123441 |
| VTA1 | OxA | 2.033446147 | 0.00925878 |
| PPHLN1 | OxA | 2.456910065 | 0.00349213 |
| IL23A | OxA | 2.518750439 | 0.00302865 |
| RTRAF | OxA | 2.821285612 | 0.00150909 |
| PPIL1 | OxA | 3.278356907 | 0.0005268 |
| MRPS23 | OxA | 2.139546461 | 0.00725193 |
| MRPS33 | OxA | 5.176214766 | 6.66E-06 |
| MPC1 | OxA | 2.174879243 | 0.0066853 |
| IFT25 | OxA | 2.841470578 | 0.00144055 |
| LSM8 | OxA | 2.952921759 | 0.0011145 |
| CMPK1 | OxA | 2.662109007 | 0.00217716 |
| LUC7L3 | OxA | 2.170872469 | 0.00674726 |
| ERAP1 | OxA | 2.218221392 | 0.00605032 |
| DNAJC10 | OxA | 2.83707524 | 0.00145521 |
| GAR1 | OxA | 2.461830127 | 0.00345279 |
| RETREG1 | OxA | 4.302729006 | 4.98E-05 |
| ETAA1 | OxA | 2.730109666 | 0.00186162 |
| MRPL50 | OxA | 2.524434789 | 0.00298927 |
| SHLD2 | OxA | 2.315870649 | 0.00483203 |
| TOMM7 | OxA | 3.55492514 | 0.00027866 |
| RRN3 | OxA | 1.990828128 | 0.01021344 |
| DCAF16 | OxA | 3.175482791 | 0.0006676 |
| TTC19 | OxA | 3.495611319 | 0.00031944 |
| OCIAD1 | OxA | 3.055624635 | 0.00087978 |
| COMMD8 | OxA | 2.477079654 | 0.00333365 |
| HPF1 | OxA | 6.201591784 | 6.29E-07 |
| TTC12 | OxA | 2.741740048 | 0.00181242 |
| NMRK1 | OxA | 2.201472882 | 0.00628821 |
| ZNF770 | OxA | 2.10925874 | 0.00777573 |
| MCUB | OxA | 2.684522415 | 0.00206765 |
| PRPF39 | OxA | 3.144612424 | 0.00071678 |
| HEATR3 | OxA | 2.891264971 | 0.0012845 |
| USP47 | OxA | 3.240920149 | 0.00057422 |
| FKBP14 | OxA | 2.729620745 | 0.00186371 |
| NOL8 | OxA | 3.464178827 | 0.00034342 |
| PTCD3 | OxA | 3.943529451 | 0.00011389 |
| GABPB1-IT1 | OxA | 2.117060868 | 0.00763729 |
| ARGLU1 | OxA | 2.37817945 | 0.00418621 |
| RALGPS2 | OxA | 2.415528505 | 0.00384124 |
| PRPF38B | OxA | 2.42392465 | 0.00376769 |
| LARP1B | OxA | 2.958362689 | 0.00110062 |
| SDAD1 | OxA | 1.992531622 | 0.01017345 |
| SHQ1 | OxA | 2.014785356 | 0.00966528 |
| RIC8B | OxA | 2.973799275 | 0.00106219 |
| SMU1 | OxA | 2.210282901 | 0.00616193 |
| UBA6 | OxA | 2.369546163 | 0.00427026 |
| CCDC25 | OxA | 3.34392865 | 0.00045297 |
| ELP2 | OxA | 4.013092326 | 9.70E-05 |
| TYW1 | OxA | 2.541696973 | 0.00287278 |
| TBC1D19 | OxA | 2.538131398 | 0.00289647 |
| BRIX1 | OxA | 3.047368395 | 0.00089667 |
| SYNJ2BP | OxA | 3.377833857 | 0.00041895 |
| KIF21A | OxA | 2.111740602 | 0.00773142 |
| ANKRD10 | OxA | 2.044793366 | 0.00902 |
| DOCK10 | OxA | 3.311141062 | 0.00048849 |
| TTC27 | OxA | 2.173717723 | 0.0067032 |
| PNRC2 | OxA | 2.306591884 | 0.00493637 |
| SLF2 | OxA | 2.116023239 | 0.00765556 |

| | | | |
|---|---|---|---|
| NUP133 | OxA | 2.430361 | 0.00371227 |
| CCAR1 | OxA | 4.788257086 | 1.63E-05 |
| NGLY1 | OxA | 2.075675927 | 0.00840087 |
| ZNF83 | OxA | 4.591488717 | 2.56E-05 |
| EXOC2 | OxA | 3.219038799 | 0.00060389 |
| BDP1 | OxA | 2.657619241 | 0.00219979 |
| CAND1 | OxA | 4.123174184 | 7.53E-05 |
| ZC3H15 | OxA | 2.611461967 | 0.00244646 |
| ECHDC1 | OxA | 3.773676621 | 0.00016839 |
| TMEM126B | OxA | 2.249469299 | 0.00563029 |
| ZNF253 | OxA | 2.509735993 | 0.00309217 |
| BCCIP | OxA | 2.909062085 | 0.00123293 |
| CLDND1 | OxA | 2.618651428 | 0.00240629 |
| CFAP298 | OxA | 1.985754872 | 0.01033344 |
| BDH2 | OxA | 2.107265108 | 0.00781151 |
| SMARCAD1 | OxA | 3.352805181 | 0.00044381 |
| MCCC1 | OxA | 3.07360922 | 0.00084409 |
| CMC2 | OxA | 2.297512532 | 0.00504066 |
| NIT2 | OxA | 2.543848718 | 0.00285859 |
| POGLUT1 | OxA | 2.584779103 | 0.00260148 |
| ADPRM | OxA | 5.016830144 | 9.62E-06 |
| TULP4 | OxA | 2.345947168 | 0.00450872 |
| PCNP | OxA | 2.137432086 | 0.00728732 |
| GOPC | OxA | 2.77278274 | 0.0016874 |
| NUP107 | OxA | 2.93215231 | 0.00116909 |
| MRPL47 | OxA | 2.085655302 | 0.00821003 |
| PHTF2 | OxA | 2.055832218 | 0.00879362 |
| DENND11 | OxA | 2.029417752 | 0.00934506 |
| ODF2L | OxA | 2.783590916 | 0.00164592 |
| COG6 | OxA | 2.042454163 | 0.00906872 |
| TMEM181 | OxA | 3.486603182 | 0.00032613 |
| ZFP14 | OxA | 3.331411917 | 0.00046622 |
| GPAM | OxA | 2.540336681 | 0.0028818 |
| TNRC6C | OxA | 2.055223539 | 0.00880595 |
| METTL14 | OxA | 2.634603498 | 0.00231951 |
| ANKRA2 | OxA | 2.112478886 | 0.00771829 |
| GATAD1 | OxA | 2.645764916 | 0.00226066 |
| ENOPH1 | OxA | 2.826477205 | 0.00149116 |
| CREBZF | OxA | 2.273505985 | 0.00532714 |
| OSTC | OxA | 2.831288 | 0.00147473 |
| SINHCAF | OxA | 2.702479556 | 0.0019839 |
| MRPS35 | OxA | 3.262540555 | 0.00054634 |
| NIF3L1 | OxA | 2.373942937 | 0.00422724 |
| CCDC90B | OxA | 3.356498843 | 0.00044005 |
| LDAH | OxA | 2.434029234 | 0.00368104 |
| RINT1 | OxA | 3.649502007 | 0.00022413 |
| GAS5 | OxA | 4.718943202 | 1.91E-05 |
| FAM111A | OxA | 4.347374639 | 4.49E-05 |
| NAPB | OxA | 2.364283083 | 0.00432232 |
| RBM26 | OxA | 2.86110526 | 0.00137688 |
| MOAP1 | OxA | 4.246394978 | 5.67E-05 |
| SLC39A8 | OxA | 2.534750836 | 0.0029191 |
| C17orf75 | OxA | 2.339464934 | 0.00457652 |
| NOC3L | OxA | 2.373039057 | 0.00423605 |
| ZMAT3 | OxA | 3.447479088 | 0.00035688 |
| ACTR6 | OxA | 3.830620895 | 0.0001477 |
| MRPS25 | OxA | 3.450663567 | 0.00035427 |
| AIDA | OxA | 3.655357677 | 0.00022113 |
| MRPS9 | OxA | 3.136580713 | 0.00073016 |
| MRPS6 | OxA | 2.393801094 | 0.0040383 |
| MRPL32 | OxA | 3.233055713 | 0.00058472 |
| MRPL1 | OxA | 2.71949835 | 0.00190766 |
| CAPRIN2 | OxA | 2.011103261 | 0.00974758 |
| ZBTB10 | OxA | 3.007337337 | 0.00098325 |
| MIS12 | OxA | 2.654439901 | 0.00221595 |
| DDX50 | OxA | 3.450999417 | 0.000354 |
| C2orf49 | OxA | 2.261637209 | 0.00547473 |
| ZNF426 | OxA | 2.239729264 | 0.00575799 |
| TAF1D | OxA | 3.151459968 | 0.00070557 |
| GNPTAB | OxA | 3.189621721 | 0.00064622 |
| HAUS3 | OxA | 3.016967832 | 0.00096168 |

| | | | |
|---|---|---|---|
| TTC13 | OxA | 2.909879504 | 0.00123061 |
| OBI1 | OxA | 2.113238054 | 0.00770481 |
| RIC3 | OxA | 3.271438997 | 0.00053526 |
| NAA16 | OxA | 3.096935621 | 0.00079995 |
| RNASEH2B | OxA | 2.98917984 | 0.00102523 |
| SMC6 | OxA | 3.139279364 | 0.00072564 |
| CCDC82 | OxA | 2.704376513 | 0.00197526 |
| CAMKMT | OxA | 2.269077783 | 0.00538173 |
| ZC3H14 | OxA | 2.016838493 | 0.0096197 |
| ERMP1 | OxA | 4.326596764 | 4.71E-05 |
| TRAPPC13 | OxA | 2.045775026 | 0.00899964 |
| PHC3 | OxA | 2.232869074 | 0.00584966 |
| RPF1 | OxA | 2.133640615 | 0.00735122 |
| SIKE1 | OxA | 2.408474053 | 0.00390415 |
| THOC7 | OxA | 2.39355768 | 0.00404057 |
| NAA15 | OxA | 2.781282061 | 0.00165469 |
| IFT74 | OxA | 4.002745318 | 9.94E-05 |
| SKIC8 | OxA | 2.722435165 | 0.00189481 |
| THUMPD2 | OxA | 2.939750092 | 0.00114881 |
| CYB5B | OxA | 2.214724096 | 0.00609924 |
| POLR1HASP | OxA | 2.354814832 | 0.00441759 |
| SLC38A1 | OxA | 3.461718085 | 0.00034537 |
| TMX1 | OxA | 2.074132749 | 0.00843077 |
| ANP32E | OxA | 2.775109152 | 0.00167838 |
| RNF170 | OxA | 2.023297693 | 0.00947769 |
| TMEM14B | OxA | 2.829411505 | 0.00148111 |
| SLIRP | OxA | 2.171528986 | 0.00673707 |
| PUS7L | OxA | 2.260152829 | 0.00549348 |
| C19orf12 | OxA | 2.009333813 | 0.00978737 |
| ARMC10 | OxA | 2.283917264 | 0.00520095 |
| SNX25 | OxA | 3.484172589 | 0.00032796 |
| TATDN1 | OxA | 4.858989274 | 1.38E-05 |
| WDR75 | OxA | 3.363171731 | 0.00043334 |
| ZCCHC7 | OxA | 2.351483449 | 0.0044516 |
| LRRC8C | OxA | 2.147992945 | 0.00711225 |
| MED10 | OxA | 2.397261077 | 0.00400626 |
| DCUN1D5 | OxA | 2.67028831 | 0.00213654 |
| POLR3GL | OxA | 2.718315396 | 0.00191287 |
| RPAIN | OxA | 2.425840595 | 0.00375111 |
| UTP23 | OxA | 2.864605937 | 0.00136582 |
| UQCC2 | OxA | 2.28855136 | 0.00514575 |
| SARNP | OxA | 2.116889857 | 0.0076403 |
| NIFK | OxA | 2.109754391 | 0.00776686 |
| ZNF527 | OxA | 2.861535787 | 0.00137551 |
| ZC3H8 | OxA | 2.140001592 | 0.00724433 |
| ZNF559 | OxA | 3.454460824 | 0.00035119 |
| MAK16 | OxA | 2.444700612 | 0.00359169 |
| DPY30 | OxA | 2.246241053 | 0.0056723 |
| PLEKHA8 | OxA | 2.117946288 | 0.00762173 |
| TRIM52 | OxA | 5.165799124 | 6.83E-06 |
| ATAD1 | OxA | 3.336745407 | 0.00046053 |
| FAM136A | OxA | 2.832413454 | 0.00147091 |
| RBM17 | OxA | 2.031763737 | 0.00929472 |
| GFM1 | OxA | 2.905379229 | 0.00124343 |
| CCDC65 | OxA | 2.332502178 | 0.00465048 |
| ZNF766 | OxA | 4.582191243 | 2.62E-05 |
| TMEM41A | OxA | 2.698281329 | 0.00200317 |
| CEP95 | OxA | 2.984391727 | 0.00103659 |
| PRMT9 | OxA | 2.666066862 | 0.00215741 |
| DMAC1 | OxA | 3.234648388 | 0.00058257 |
| BOD1 | OxA | 3.067208188 | 0.00085663 |
| RLIG1 | OxA | 2.269786794 | 0.00537296 |
| SLFN11 | OxA | 2.129970624 | 0.0074136 |
| NXPE3 | OxA | 2.555794412 | 0.00278103 |
| CHMP7 | OxA | 2.016509014 | 0.009627 |
| FDXACB1 | OxA | 2.158500103 | 0.00694224 |
| NDUFAF2 | OxA | 2.016763806 | 0.00962135 |
| OXNAD1 | OxA | 3.083098972 | 0.00082585 |
| ZNF585B | OxA | 2.263955304 | 0.00544559 |
| METTL18 | OxA | 2.913133504 | 0.00122142 |
| MRRF | OxA | 2.645847254 | 0.00226023 |

| | | | |
|---|---|---|---|
| UBE2Q2 | OxA | 3.76866294 | 0.00017035 |
| TBC1D31 | OxA | 2.780895226 | 0.00165617 |
| ZNF101 | OxA | 2.610374918 | 0.00245259 |
| ORMDL1 | OxA | 2.514264206 | 0.0030601 |
| TP53RK | OxA | 2.699505294 | 0.00199754 |
| HELQ | OxA | 2.358685986 | 0.00437839 |
| HENMT1 | OxA | 2.316153162 | 0.00482888 |
| SMYD4 | OxA | 2.302651522 | 0.00498137 |
| TMEM123 | OxA | 2.353265412 | 0.00443338 |
| HAUS1 | OxA | 3.440043854 | 0.00036304 |
| OMA1 | OxA | 2.76079676 | 0.00173462 |
| UHRF2 | OxA | 3.247050589 | 0.00056617 |
| PABIR1 | OxA | 2.060228119 | 0.00870506 |
| SSX2IP | OxA | 2.55035506 | 0.00281608 |
| FRA10AC1 | OxA | 3.28489577 | 0.00051892 |
| CYP2R1 | OxA | 2.993309435 | 0.00101552 |
| DNAJC24 | OxA | 2.054893176 | 0.00881266 |
| ARL14EP | OxA | 3.54972122 | 0.00028202 |
| AEBP2 | OxA | 3.17567079 | 0.00066731 |
| TRAPPC6B | OxA | 2.247954775 | 0.00564996 |
| TC2N | OxA | 4.096118297 | 8.01E-05 |
| TARS3 | OxA | 3.11217977 | 0.00077236 |
| METTL23 | OxA | 3.138802961 | 0.00072644 |
| ZSWIM7 | OxA | 2.577281713 | 0.00264678 |
| ZNF816 | OxA | 3.443235397 | 0.00036038 |
| TYW3 | OxA | 3.392939258 | 0.00040463 |
| UHMK1 | OxA | 2.111676403 | 0.00773257 |
| FITM2 | OxA | 2.766421365 | 0.0017123 |
| TSHZ2 | OxA | 2.133630538 | 0.00735139 |
| MITD1 | OxA | 2.271754669 | 0.00534866 |
| SLC66A3 | OxA | 2.814691395 | 0.00153218 |
| JMY | OxA | 3.187810087 | 0.00064892 |
| GRPEL2 | OxA | 2.369964168 | 0.00426615 |
| LSM11 | OxA | 2.64390359 | 0.00227037 |
| NUDCD2 | OxA | 2.746771492 | 0.00179155 |
| PM20D2 | OxA | 2.07009974 | 0.00850943 |
| NDUFAF6 | OxA | 2.276882912 | 0.00528588 |
| SESN3 | OxA | 2.719752772 | 0.00190655 |
| POGLUT3 | OxA | 2.068766266 | 0.00853559 |
| ZNF664 | OxA | 2.027449552 | 0.00938751 |
| ZSCAN29 | OxA | 2.773508049 | 0.00168458 |
| ZFP90 | OxA | 3.249657969 | 0.00056278 |
| LINC02210 | OxA | 2.014122717 | 0.00968004 |
| ZNF582 | OxA | 2.609427772 | 0.00245795 |
| ZNF570 | OxA | 2.232052203 | 0.00586068 |
| PDIK1L | OxA | 2.296409295 | 0.00505348 |
| FAM117B | OxA | 2.794279377 | 0.00160591 |
| TTC14 | OxA | 4.167424735 | 6.80E-05 |
| BTLA | OxA | 2.038096003 | 0.00916018 |
| PPM1K | OxA | 2.627357926 | 0.00235853 |
| CEP120 | OxA | 3.580394702 | 0.00026279 |
| TTC39B | OxA | 3.452616403 | 0.00035268 |
| SLFN5 | OxA | 4.359916894 | 4.37E-05 |
| ZNF600 | OxA | 2.683598292 | 0.00207206 |
| ZNF564 | OxA | 2.008646041 | 0.00980289 |
| ZNF567 | OxA | 2.179196702 | 0.00661917 |
| ZNF383 | OxA | 3.557400229 | 0.00027708 |
| DENND1B | OxA | 2.054802605 | 0.00881449 |
| CNST | OxA | 1.984081984 | 0.01037333 |
| MIER3 | OxA | 2.065346948 | 0.00860306 |
| ZNF92 | OxA | 2.065797975 | 0.00859413 |
| GIMAP7 | OxA | 5.882575154 | 1.31E-06 |
| COMMD6 | OxA | 4.213495927 | 6.12E-05 |
| ASXL1 | OxA | 2.252525767 | 0.0055908 |
| METTL15 | OxA | 2.158084008 | 0.0069489 |
| UBR1 | OxA | 2.574223349 | 0.00266549 |
| RPL22L1 | OxA | 2.020985688 | 0.00952828 |
| STT3B | OxA | 2.729948145 | 0.00186231 |
| PRIMPOL | OxA | 2.072233795 | 0.00846771 |
| DDIAS | OxA | 2.007457289 | 0.00982976 |
| HNRNPA3 | OxA | 2.862599712 | 0.00137215 |

| | | | |
|---|---|---|---|
| NSUN6 | OxA | 2.522511075 | 0.00300254 |
| MICU2 | OxA | 2.047375834 | 0.00896653 |
| OARD1 | OxA | 2.191753199 | 0.00643053 |
| POLR1F | OxA | 2.045300636 | 0.00900947 |
| EEF1A1P6 | OxA | 2.521819177 | 0.00300733 |
| CNOT6L | OxA | 2.065916884 | 0.00859178 |
| GPATCH11 | OxA | 2.770776905 | 0.00169521 |
| SVIP | OxA | 2.270065161 | 0.00536951 |
| SNHG10 | OxA | 2.876389967 | 0.00132926 |
| GDPD1 | OxA | 2.631788731 | 0.00233459 |
| ZNF615 | OxA | 2.660259451 | 0.00218646 |
| LINC00663 | OxA | 2.335496276 | 0.00461853 |
| ZNF621 | OxA | 2.64867594 | 0.00224556 |
| RABL3 | OxA | 2.598188119 | 0.00252239 |
| GPRIN3 | OxA | 3.153526122 | 0.00070222 |
| ARL10 | OxA | 2.466495697 | 0.00341589 |
| SREK1IP1 | OxA | 2.514422775 | 0.00305898 |
| ZNF789 | OxA | 2.548910744 | 0.00282546 |
| DPY19L4 | OxA | 2.60686095 | 0.00247252 |
| ZNF181 | OxA | 2.954625558 | 0.00111013 |
| PRORSD1P | OxA | 2.469167916 | 0.00339494 |
| ZNF879 | OxA | 2.37793223 | 0.00418859 |
| MTX3 | OxA | 2.363660338 | 0.00432852 |
| SKA2 | OxA | 2.511422348 | 0.00308019 |
| SFT2D2 | OxA | 2.131207968 | 0.00739251 |
| ANKRD36 | OxA | 2.430119579 | 0.00371433 |
| C6orf120 | OxA | 2.087601452 | 0.00817332 |
| THEMIS | OxA | 2.479290202 | 0.00331673 |
| LINC01550 | OxA | 2.30616065 | 0.00494128 |
| DIPK1A | OxA | 2.22277766 | 0.00598718 |
| COA6 | OxA | 2.815169478 | 0.00153049 |
| BOLA3 | OxA | 2.87304597 | 0.00133953 |
| RPL27AP6 | OxA | 2.314489575 | 0.00484742 |
| RBM12B | OxA | 3.227598471 | 0.00059211 |
| KNOP1 | OxA | 2.08244052 | 0.00827103 |
| C5orf63 | OxA | 3.189526103 | 0.00064636 |
| RBIS | OxA | 3.248451024 | 0.00056435 |
| TOMM5 | OxA | 4.566132019 | 2.72E-05 |
| PLPP6 | OxA | 2.521927757 | 0.00300658 |
| ZNF506 | OxA | 2.099681715 | 0.00794911 |
| GIMAP6 | OxA | 2.170581224 | 0.00675179 |
| COA5 | OxA | 2.857581252 | 0.00138809 |
| ATXN7L3B | OxA | 3.220896638 | 0.00060132 |
| LEF1-AS1 | OxA | 2.229749441 | 0.00589183 |
| ZFP62 | OxA | 4.54756564 | 2.83E-05 |
| RPL23AP53 | OxA | 2.563731501 | 0.00273067 |
| SCARNA17 | OxA | 2.675583287 | 0.00211065 |
| RPL36AP37 | OxA | 5.326542478 | 4.71E-06 |
| RPL21P16 | OxA | 2.008301169 | 0.00981067 |
| RRN3P1 | OxA | 2.939664316 | 0.00114904 |
| ERCC6L2-AS1 | OxA | 2.350138562 | 0.00446541 |
| SMIM10L1 | OxA | 2.975267877 | 0.0010586 |
| TIMM23 | OxA | 2.226400704 | 0.00593744 |
| THAP9-AS1 | OxA | 2.515290894 | 0.00305288 |
| LINC00674 | OxA | 2.209780138 | 0.00616907 |
| SNHG21 | OxA | 2.070213746 | 0.00850719 |
| IQCH-AS1 | OxA | 2.650106693 | 0.00223817 |
| CHASERR | OxA | 2.14758497 | 0.00711894 |
| SNHG19 | OxA | 2.180694841 | 0.00659637 |
| SDCBP2-AS1 | OxA | 2.153835671 | 0.00701721 |
| CAPN10-DT | OxA | 2.728825788 | 0.00186713 |
| FAM111A-DT | OxA | 2.067619001 | 0.00855817 |
| LINC02648 | OxA | 2.178111385 | 0.00663573 |
| CSRP1-AS1 | OxA | 2.456309851 | 0.00349696 |
| ABR | SNs | 5.017569576 | 9.60E-06 |
| ACAA1 | SNs | 2.44282697 | 0.00360722 |
| ACTA2 | SNs | 1.984484829 | 0.01036371 |
| ACTB | SNs | 5.521938141 | 3.01E-06 |
| ACTG1 | SNs | 3.516106939 | 0.00030471 |
| ACTN4 | SNs | 4.078341729 | 8.35E-05 |
| ACTN1 | SNs | 4.960371643 | 1.10E-05 |

| | | | |
|---|---|---|---|
| ADAM8 | SNs | 4.752922201 | 1.77E-05 |
| ADAR | SNs | 2.488337973 | 0.00324834 |
| ADCY6 | SNs | 2.171801348 | 0.00673285 |
| ADD1 | SNs | 3.192855775 | 0.00064142 |
| GRK2 | SNs | 4.003381425 | 9.92E-05 |
| AP2A1 | SNs | 2.812066584 | 0.00154146 |
| AP1B1 | SNs | 2.787441162 | 0.00163139 |
| AGER | SNs | 3.419339836 | 0.00038077 |
| AKT1 | SNs | 7.677521277 | 2.10E-08 |
| AKT2 | SNs | 3.433767949 | 0.00036833 |
| ALAS1 | SNs | 2.524024876 | 0.00299209 |
| ALDH2 | SNs | 3.218566554 | 0.00060455 |
| ALDH3B1 | SNs | 5.471857337 | 3.37E-06 |
| ALDOA | SNs | 4.630994823 | 2.34E-05 |
| ALOX12 | SNs | 3.731765301 | 0.00018545 |
| ALOX5 | SNs | 6.008686715 | 9.80E-07 |
| ALOX5AP | SNs | 2.433847723 | 0.00368258 |
| ALPL | SNs | 2.965903965 | 0.00108167 |
| AMPD2 | SNs | 4.647000137 | 2.25E-05 |
| ANPEP | SNs | 6.059337077 | 8.72E-07 |
| ANXA11 | SNs | 5.977235433 | 1.05E-06 |
| APLP2 | SNs | 3.200878016 | 0.00062968 |
| ARF1 | SNs | 6.802086911 | 1.58E-07 |
| ARF3 | SNs | 4.625476029 | 2.37E-05 |
| ARF5 | SNs | 4.708358913 | 1.96E-05 |
| RHOA | SNs | 3.604463997 | 0.00024862 |
| RHOG | SNs | 5.700204124 | 1.99E-06 |
| ARHGAP1 | SNs | 4.93368719 | 1.16E-05 |
| ARHGDIB | SNs | 3.135396516 | 0.00073216 |
| ARRB1 | SNs | 2.929406347 | 0.0011765 |
| ARRB2 | SNs | 5.557830123 | 2.77E-06 |
| ARSA | SNs | 3.294606597 | 0.00050745 |
| ASGR1 | SNs | 2.932442393 | 0.00116831 |
| ASGR2 | SNs | 3.712243589 | 0.00019398 |
| ASL | SNs | 2.734288475 | 0.00184379 |
| RERE | SNs | 3.050722051 | 0.00088977 |
| ATP2A3 | SNs | 3.007041828 | 0.00098392 |
| ATP6V0B | SNs | 5.101604813 | 7.91E-06 |
| ATP6V0A1 | SNs | 6.779499395 | 1.66E-07 |
| AZU1 | SNs | 2.644728715 | 0.00226606 |
| BAX | SNs | 2.441056361 | 0.00362196 |
| BCL3 | SNs | 5.623872022 | 2.38E-06 |
| TSPO | SNs | 3.701450962 | 0.00019886 |
| SERPING1 | SNs | 2.656085984 | 0.00220757 |
| C5AR1 | SNs | 3.659924084 | 0.00021881 |
| FMNL1 | SNs | 5.834319087 | 1.46E-06 |
| CA4 | SNs | 2.667630176 | 0.00214966 |
| CALM3 | SNs | 3.28648198 | 0.00051703 |
| CAMK2G | SNs | 3.339439695 | 0.00045768 |
| CAPG | SNs | 2.596002678 | 0.00253511 |
| CAPN1 | SNs | 3.064672316 | 0.00086164 |
| CAPNS1 | SNs | 2.346885041 | 0.00449899 |
| CAPZB | SNs | 4.89269594 | 1.28E-05 |
| CARS1 | SNs | 2.410656278 | 0.00388458 |
| CASP9 | SNs | 3.62978887 | 0.00023454 |
| CCND3 | SNs | 2.637437933 | 0.00230442 |
| CD14 | SNs | 5.159611663 | 6.92E-06 |
| TNFRSF8 | SNs | 3.643080155 | 0.00022747 |
| CD33 | SNs | 2.10958922 | 0.00776982 |
| SCARB1 | SNs | 2.759297159 | 0.00174062 |
| CD37 | SNs | 2.583223252 | 0.00261082 |
| CD63 | SNs | 3.862014446 | 0.0001374 |
| CD68 | SNs | 4.123368873 | 7.53E-05 |
| ADGRE5 | SNs | 3.427105038 | 0.00037402 |
| CDA | SNs | 4.630740859 | 2.34E-05 |
| CDK11B | SNs | 2.251929198 | 0.00559849 |
| CDKN1A | SNs | 3.835652112 | 0.000146 |
| CDKN2D | SNs | 4.743126196 | 1.81E-05 |
| CEBPA | SNs | 2.513901097 | 0.00306266 |
| CFL1 | SNs | 4.685029957 | 2.07E-05 |
| CEACAM3 | SNs | 4.480515369 | 3.31E-05 |

| | | | |
|---|---|---|---|
| CEACAM4 | SNs | 3.900508124 | 0.00012575 |
| CHRNA2 | SNs | 2.987785408 | 0.00102852 |
| AP2M1 | SNs | 2.174325239 | 0.00669383 |
| CLU | SNs | 3.984520379 | 0.00010363 |
| CLK3 | SNs | 4.931540217 | 1.17E-05 |
| TPP1 | SNs | 2.860583077 | 0.00137853 |
| CLN3 | SNs | 3.225998214 | 0.00059429 |
| CLPTM1 | SNs | 4.027320831 | 9.39E-05 |
| CCR1 | SNs | 2.171568354 | 0.00673646 |
| LTB4R | SNs | 2.722581504 | 0.00189417 |
| CNN2 | SNs | 4.600955774 | 2.51E-05 |
| COL7A1 | SNs | 2.363639456 | 0.00432873 |
| COMT | SNs | 2.585286779 | 0.00259844 |
| ATF6B | SNs | 4.458616451 | 3.48E-05 |
| CSF1R | SNs | 2.697538132 | 0.0020066 |
| CSF3R | SNs | 6.044520663 | 9.03E-07 |
| CSK | SNs | 3.662155319 | 0.00021769 |
| CSNK1D | SNs | 4.997820114 | 1.01E-05 |
| CSNK2B | SNs | 3.071319762 | 0.00084856 |
| CST3 | SNs | 3.577681116 | 0.00026443 |
| CSTB | SNs | 2.143484147 | 0.00718647 |
| CTSD | SNs | 5.177890095 | 6.64E-06 |
| CTSZ | SNs | 2.609715013 | 0.00245632 |
| CUX1 | SNs | 3.192395121 | 0.0006421 |
| CYBA | SNs | 5.134536312 | 7.34E-06 |
| DAP | SNs | 2.022665545 | 0.00949149 |
| DAPK3 | SNs | 2.490359299 | 0.00323326 |
| DBN1 | SNs | 2.364395156 | 0.0043212 |
| DCTN1 | SNs | 2.721320153 | 0.00189968 |
| DHX8 | SNs | 3.645883768 | 0.000226 |
| CYB5R3 | SNs | 4.297755908 | 5.04E-05 |
| DIAPH1 | SNs | 3.500046065 | 0.00031619 |
| DLG4 | SNs | 3.054818466 | 0.00088142 |
| DNM1 | SNs | 1.987329856 | 0.01029604 |
| DNM2 | SNs | 4.393487613 | 4.04E-05 |
| DOCK2 | SNs | 2.206614705 | 0.0062142 |
| DOK1 | SNs | 4.448093188 | 3.56E-05 |
| ARID3A | SNs | 3.204883606 | 0.0006239 |
| DUSP1 | SNs | 2.105650679 | 0.0078406 |
| DUSP3 | SNs | 3.503876087 | 0.00031342 |
| DVL3 | SNs | 3.155467855 | 0.00069909 |
| E2F4 | SNs | 2.90552207 | 0.00124302 |
| ECE1 | SNs | 5.137749854 | 7.28E-06 |
| PHC2 | SNs | 4.791113736 | 1.62E-05 |
| EIF4EBP1 | SNs | 2.79139934 | 0.00161659 |
| MARK2 | SNs | 4.124609889 | 7.51E-05 |
| EMP3 | SNs | 2.015741035 | 0.00964404 |
| ADGRE1 | SNs | 2.566207661 | 0.00271514 |
| CTTN | SNs | 3.567519698 | 0.0002707 |
| ENO1 | SNs | 2.689359661 | 0.00204475 |
| EPHB4 | SNs | 2.799504458 | 0.0015867 |
| ERF | SNs | 3.890778966 | 0.00012859 |
| ETV6 | SNs | 2.265939736 | 0.00542076 |
| EWSR1 | SNs | 2.510627427 | 0.00308583 |
| EXTL3 | SNs | 3.44383376 | 0.00035989 |
| F12 | SNs | 2.367520652 | 0.00429022 |
| FAH | SNs | 2.048443428 | 0.00894451 |
| PTK2B | SNs | 4.121605295 | 7.56E-05 |
| FBP1 | SNs | 2.27179721 | 0.00534814 |
| FCER1G | SNs | 2.237441631 | 0.0057884 |
| FCGRT | SNs | 5.818715344 | 1.52E-06 |
| FCN1 | SNs | 4.913812183 | 1.22E-05 |
| FES | SNs | 3.88966267 | 0.00012893 |
| FGR | SNs | 4.453077968 | 3.52E-05 |
| FHL3 | SNs | 2.050082882 | 0.00891081 |
| FKBP1A | SNs | 3.237325948 | 0.00057899 |
| FLII | SNs | 3.552291305 | 0.00028036 |
| FLOT2 | SNs | 6.001573264 | 9.96E-07 |
| FOLR3 | SNs | 2.424511099 | 0.00376261 |
| FOS | SNs | 3.397091483 | 0.00040078 |
| FOSB | SNs | 2.149293609 | 0.00709098 |

| | | | |
|---|---|---|---|
| FOSL2 | SNs | 3.649593248 | 0.00022408 |
| FPR1 | SNs | 4.235924148 | 5.81E-05 |
| FTH1 | SNs | 2.794371185 | 0.00160557 |
| FUT7 | SNs | 3.08355251 | 0.00082499 |
| FZD2 | SNs | 1.985078291 | 0.01034956 |
| GAA | SNs | 3.094666808 | 0.00080414 |
| GAK | SNs | 2.16249502 | 0.00687868 |
| GALK1 | SNs | 3.627800171 | 0.00023561 |
| GALNS | SNs | 3.236870056 | 0.0005796 |
| GALNT2 | SNs | 3.69179072 | 0.00020333 |
| GAPDH | SNs | 2.964243908 | 0.00108582 |
| GBA1 | SNs | 5.142817085 | 7.20E-06 |
| NR6A1 | SNs | 2.760805367 | 0.00173458 |
| GGT1 | SNs | 2.106714575 | 0.00782142 |
| GLB1 | SNs | 1.9949477 | 0.01011701 |
| GLUL | SNs | 3.413131177 | 0.00038625 |
| GNA15 | SNs | 4.346405227 | 4.50E-05 |
| GNAI2 | SNs | 7.628818854 | 2.35E-08 |
| GNAZ | SNs | 2.968465679 | 0.00107531 |
| GNB1 | SNs | 5.446788607 | 3.57E-06 |
| GNB2 | SNs | 6.479928617 | 3.31E-07 |
| GNG5 | SNs | 3.646877085 | 0.00022549 |
| GP1BA | SNs | 4.198385008 | 6.33E-05 |
| GP9 | SNs | 4.500183591 | 3.16E-05 |
| GPM6A | SNs | 4.098555558 | 7.97E-05 |
| FFAR2 | SNs | 2.914733655 | 0.00121693 |
| GRK6 | SNs | 4.30385935 | 4.97E-05 |
| MKNK2 | SNs | 4.547496646 | 2.83E-05 |
| GPS2 | SNs | 2.402066795 | 0.00396217 |
| GRB2 | SNs | 4.751606552 | 1.77E-05 |
| GRN | SNs | 6.551021948 | 2.81E-07 |
| GRINA | SNs | 2.819040149 | 0.00151691 |
| GSK3A | SNs | 3.565122918 | 0.00027219 |
| GSN | SNs | 4.093706696 | 8.06E-05 |
| GSTP1 | SNs | 2.353593258 | 0.00443003 |
| GSTZ1 | SNs | 2.375459564 | 0.00421251 |
| H1-2 | SNs | 2.633590394 | 0.00232493 |
| H3-3A | SNs | 2.723353151 | 0.00189081 |
| H3-3B | SNs | 2.061450068 | 0.0086806 |
| HCK | SNs | 5.76407092 | 1.72E-06 |
| HCLS1 | SNs | 4.497347379 | 3.18E-05 |
| HDLBP | SNs | 2.728727041 | 0.00186755 |
| HK1 | SNs | 2.343422616 | 0.004535 |
| HK3 | SNs | 2.969808913 | 0.00107199 |
| ZBTB48 | SNs | 2.308194054 | 0.0049182 |
| HLA-B | SNs | 3.605917653 | 0.00024779 |
| HLA-C | SNs | 3.200909828 | 0.00062964 |
| HLA-DRB1 | SNs | 2.871114243 | 0.00134551 |
| HLA-E | SNs | 4.117654416 | 7.63E-05 |
| HLX | SNs | 5.663762206 | 2.17E-06 |
| HMOX1 | SNs | 2.742056768 | 0.0018111 |
| HPCAL1 | SNs | 4.060724114 | 8.70E-05 |
| HPS1 | SNs | 2.494493401 | 0.00320263 |
| HRH2 | SNs | 5.897531513 | 1.27E-06 |
| HSPA1A | SNs | 4.858481069 | 1.39E-05 |
| HSPA6 | SNs | 3.745046643 | 0.00017987 |
| NDST1 | SNs | 3.786610959 | 0.00016345 |
| ICAM3 | SNs | 5.022614132 | 9.49E-06 |
| SP110 | SNs | 2.108701458 | 0.00778572 |
| IFIT3 | SNs | 2.181518832 | 0.00658387 |
| IL1RN | SNs | 4.645442576 | 2.26E-05 |
| IL4R | SNs | 2.083061295 | 0.00825921 |
| CXCR1 | SNs | 4.355086906 | 4.41E-05 |
| CXCR2 | SNs | 3.328133024 | 0.00046975 |
| IL10RB | SNs | 2.823233498 | 0.00150233 |
| IL16 | SNs | 2.339105754 | 0.0045803 |
| ILK | SNs | 4.233532454 | 5.84E-05 |
| IMPA2 | SNs | 4.779579705 | 1.66E-05 |
| IMPDH1 | SNs | 7.201021009 | 6.29E-08 |
| INPP5D | SNs | 2.773453744 | 0.00168479 |
| INPPL1 | SNs | 4.001929155 | 9.96E-05 |

| | | | |
|---|---|---|---|
| IRF1 | SNs | 3.390031122 | 0.00040735 |
| IRF7 | SNs | 2.128612596 | 0.00743682 |
| ITGA2B | SNs | 4.278844846 | 5.26E-05 |
| ITGA5 | SNs | 4.483223978 | 3.29E-05 |
| ITGAM | SNs | 4.731944072 | 1.85E-05 |
| ITGAX | SNs | 3.308071939 | 0.00049196 |
| ITGB2 | SNs | 3.632217278 | 0.00023323 |
| ITGB3 | SNs | 2.0407415 | 0.00910455 |
| ITGB4 | SNs | 2.117709954 | 0.00762588 |
| ITGB5 | SNs | 4.454320815 | 3.51E-05 |
| ITPK1 | SNs | 5.972162491 | 1.07E-06 |
| JUNB | SNs | 2.407322169 | 0.00391451 |
| CD82 | SNs | 2.936752057 | 0.00115677 |
| KCNQ1 | SNs | 4.34164631 | 4.55E-05 |
| KIF3C | SNs | 2.74413527 | 0.00180246 |
| LAMP1 | SNs | 2.311308147 | 0.00488306 |
| LASP1 | SNs | 4.964623155 | 1.08E-05 |
| LGALS9 | SNs | 5.87855678 | 1.32E-06 |
| LIMK2 | SNs | 3.534035128 | 0.00029239 |
| LMNA | SNs | 2.277320938 | 0.00528055 |
| LRCH4 | SNs | 3.302436282 | 0.00049838 |
| LRP3 | SNs | 3.649023269 | 0.00022438 |
| LRPAP1 | SNs | 3.149658527 | 0.0007085 |
| LSP1 | SNs | 6.397108318 | 4.01E-07 |
| LTBR | SNs | 7.531235871 | 2.94E-08 |
| MAN2B1 | SNs | 2.711406273 | 0.00194354 |
| MAP1A | SNs | 2.859773868 | 0.0013811 |
| MAX | SNs | 2.145829927 | 0.00714776 |
| MAZ | SNs | 2.586254693 | 0.00259266 |
| MEF2D | SNs | 3.503628775 | 0.0003136 |
| MEFV | SNs | 3.171845844 | 0.00067322 |
| MAP3K3 | SNs | 4.382272167 | 4.15E-05 |
| RAB8A | SNs | 3.319958331 | 0.00047868 |
| MGAT1 | SNs | 4.05779372 | 8.75E-05 |
| MAP3K11 | SNs | 3.16786527 | 0.00067941 |
| MLLT1 | SNs | 4.76274153 | 1.73E-05 |
| MMP9 | SNs | 4.427330276 | 3.74E-05 |
| MMP19 | SNs | 3.594566817 | 0.00025435 |
| MNT | SNs | 2.511316147 | 0.00308094 |
| MPST | SNs | 3.213852715 | 0.00061115 |
| MSRA | SNs | 3.093298227 | 0.00080668 |
| MTF1 | SNs | 2.029948335 | 0.00933365 |
| MYO1F | SNs | 6.056083383 | 8.79E-07 |
| MTX1 | SNs | 5.919380014 | 1.20E-06 |
| MX2 | SNs | 3.315365317 | 0.00048377 |
| MYBL2 | SNs | 1.993877297 | 0.01014198 |
| MYBPC3 | SNs | 5.153855153 | 7.02E-06 |
| MYD88 | SNs | 4.382866221 | 4.14E-05 |
| MYH9 | SNs | 2.612547927 | 0.00244035 |
| MYO7B | SNs | 2.158159848 | 0.00694769 |
| MYO9B | SNs | 4.788783171 | 1.63E-05 |
| NAGA | SNs | 2.434816575 | 0.00367437 |
| NCF2 | SNs | 2.757620038 | 0.00174735 |
| NCF4 | SNs | 5.145870621 | 7.15E-06 |
| NDUFS2 | SNs | 2.71133335 | 0.00194387 |
| NEU1 | SNs | 3.50802044 | 0.00031044 |
| NFE2 | SNs | 5.336639212 | 4.61E-06 |
| NFIC | SNs | 3.202921152 | 0.00062673 |
| NFKB2 | SNs | 2.869623455 | 0.00135013 |
| NFKBIB | SNs | 2.416359327 | 0.0038339 |
| NFYC | SNs | 5.224865933 | 5.96E-06 |
| NINJ1 | SNs | 5.607840276 | 2.47E-06 |
| NINJ2 | SNs | 2.239236031 | 0.00576453 |
| CNOT3 | SNs | 4.833807622 | 1.47E-05 |
| CCN3 | SNs | 3.190583266 | 0.00064479 |
| NRGN | SNs | 4.815905024 | 1.53E-05 |
| NUCB1 | SNs | 2.517982964 | 0.00303401 |
| OAS1 | SNs | 2.324674072 | 0.00473506 |
| OAS2 | SNs | 2.241196233 | 0.00573857 |
| OAZ2 | SNs | 2.103029364 | 0.00788807 |
| OPRL1 | SNs | 3.337949059 | 0.00045925 |

| | | | |
|---|---|---|---|
| SLC22A18 | SNs | 2.831838889 | 0.00147286 |
| OTX1 | SNs | 2.701449184 | 0.00198862 |
| P4HB | SNs | 2.986101249 | 0.00103252 |
| FURIN | SNs | 5.944630696 | 1.14E-06 |
| PCSK6 | SNs | 2.872221237 | 0.00134208 |
| PAK1 | SNs | 2.160173409 | 0.00691555 |
| PBX2 | SNs | 2.876958262 | 0.00132752 |
| PCBP1 | SNs | 5.445046608 | 3.59E-06 |
| CHMP1A | SNs | 4.119987463 | 7.59E-05 |
| PECAM1 | SNs | 2.287811874 | 0.00515452 |
| PEX6 | SNs | 2.032599198 | 0.00927686 |
| PF4 | SNs | 4.268335014 | 5.39E-05 |
| PFKFB3 | SNs | 2.546974427 | 0.00283809 |
| PFKFB4 | SNs | 5.578864782 | 2.64E-06 |
| PFN1 | SNs | 3.467975297 | 0.00034043 |
| PGD | SNs | 6.553322643 | 2.80E-07 |
| PGM1 | SNs | 2.003459586 | 0.00992066 |
| PHKG2 | SNs | 2.904281424 | 0.00124658 |
| SERPINA1 | SNs | 6.19118921 | 6.44E-07 |
| PIK3CD | SNs | 5.643938927 | 2.27E-06 |
| PI4KB | SNs | 3.617122341 | 0.00024148 |
| PITPNA | SNs | 2.770236196 | 0.00169732 |
| PKM | SNs | 3.156883313 | 0.00069681 |
| PLAUR | SNs | 4.81608712 | 1.53E-05 |
| PLCB2 | SNs | 3.927328623 | 0.00011821 |
| PLCG2 | SNs | 3.468944324 | 0.00033967 |
| PLD2 | SNs | 2.483380436 | 0.00328564 |
| PLOD1 | SNs | 4.04880488 | 8.94E-05 |
| PLXNA2 | SNs | 2.804768996 | 0.00156758 |
| PMM2 | SNs | 3.19807697 | 0.00063376 |
| SEPTIN5 | SNs | 2.138984728 | 0.00726131 |
| SEPTIN4 | SNs | 2.971693828 | 0.00106735 |
| POLR2A | SNs | 3.044443671 | 0.00090273 |
| POLR2E | SNs | 4.258325039 | 5.52E-05 |
| POR | SNs | 4.431525024 | 3.70E-05 |
| CTSA | SNs | 7.735971093 | 1.84E-08 |
| PPP1CA | SNs | 2.634133775 | 0.00232202 |
| PPP1R10 | SNs | 3.619381766 | 0.00024023 |
| PPP2R1A | SNs | 2.305143823 | 0.00495286 |
| PPP4C | SNs | 4.965372672 | 1.08E-05 |
| PRCC | SNs | 2.333648964 | 0.00463822 |
| PRKACA | SNs | 4.918768775 | 1.21E-05 |
| PRKCD | SNs | 6.027114305 | 9.39E-07 |
| PRKCSH | SNs | 2.776395354 | 0.00167342 |
| MAPK3 | SNs | 5.213681033 | 6.11E-06 |
| MAPK13 | SNs | 2.060868903 | 0.00869223 |
| MAP2K2 | SNs | 2.832720017 | 0.00146987 |
| MAP2K3 | SNs | 3.709451486 | 0.00019523 |
| MAP2K7 | SNs | 2.720138583 | 0.00190485 |
| PSAP | SNs | 5.17847605 | 6.63E-06 |
| PSMB3 | SNs | 4.246125087 | 5.67E-05 |
| PTAFR | SNs | 6.412776468 | 3.87E-07 |
| PTGIR | SNs | 2.510007054 | 0.00309025 |
| PTGS1 | SNs | 3.968528313 | 0.00010752 |
| QSOX1 | SNs | 2.955678684 | 0.00110744 |
| PTPN6 | SNs | 3.858998183 | 0.00013836 |
| PTPRF | SNs | 3.910881013 | 0.00012278 |
| PVR | SNs | 2.277136466 | 0.00528279 |
| NECTIN1 | SNs | 4.7546471 | 1.76E-05 |
| PXN | SNs | 6.667233229 | 2.15E-07 |
| PYGL | SNs | 2.30924434 | 0.00490632 |
| RGL2 | SNs | 2.566870142 | 0.002711 |
| MAP4K2 | SNs | 2.582392512 | 0.00261582 |
| RAB5C | SNs | 5.177914831 | 6.64E-06 |
| RAC1 | SNs | 4.160641294 | 6.91E-05 |
| RAC2 | SNs | 3.911446102 | 0.00012262 |
| RAF1 | SNs | 2.474304058 | 0.00335503 |
| RALB | SNs | 2.106793077 | 0.00782 |
| RARA | SNs | 6.749809757 | 1.78E-07 |
| RELB | SNs | 4.587906721 | 2.58E-05 |
| UPF1 | SNs | 3.269574568 | 0.00053756 |

| | | | |
|---|---|---|---|
| DPF2 | SNs | 5.199669664 | 6.31E-06 |
| RFX1 | SNs | 2.655444196 | 0.00221083 |
| RFX2 | SNs | 2.96951813 | 0.00107271 |
| RNASE2 | SNs | 2.679994923 | 0.00208932 |
| RNPEP | SNs | 2.134677726 | 0.00733369 |
| RPS6KA1 | SNs | 6.053933803 | 8.83E-07 |
| RPS6KB2 | SNs | 2.070194851 | 0.00850756 |
| RRBP1 | SNs | 2.197973708 | 0.00633908 |
| RTN2 | SNs | 3.807523122 | 0.00015577 |
| RXRA | SNs | 4.708157998 | 1.96E-05 |
| S100A11 | SNs | 6.243036241 | 5.71E-07 |
| SAFB | SNs | 2.35280259 | 0.0044381 |
| CLEC11A | SNs | 3.770304534 | 0.00016971 |
| SCN1B | SNs | 3.961633286 | 0.00010924 |
| SEC14L1 | SNs | 3.758063354 | 0.00017456 |
| SECTM1 | SNs | 5.687177702 | 2.06E-06 |
| SELP | SNs | 2.595441011 | 0.00253839 |
| SELPLG | SNs | 6.550604017 | 2.81E-07 |
| SGTA | SNs | 3.097352991 | 0.00079918 |
| SH3BP2 | SNs | 4.483978216 | 3.28E-05 |
| SH3GL1 | SNs | 3.906887773 | 0.00012391 |
| ST3GAL2 | SNs | 4.432065423 | 3.70E-05 |
| ST3GAL4 | SNs | 3.862407204 | 0.00013728 |
| SIPA1 | SNs | 3.699657482 | 0.00019968 |
| SKIC2 | SNs | 2.338115304 | 0.00459076 |
| SLC2A3 | SNs | 2.504971212 | 0.00312629 |
| SLC6A6 | SNs | 2.403906717 | 0.00394542 |
| SLC9A1 | SNs | 5.44444299 | 3.59E-06 |
| SLC11A1 | SNs | 3.21412088 | 0.00061077 |
| SLC19A1 | SNs | 5.455485209 | 3.50E-06 |
| SLC25A1 | SNs | 2.200232438 | 0.0063062 |
| SMARCD2 | SNs | 2.157843219 | 0.00695275 |
| SMARCD3 | SNs | 4.4672936 | 3.41E-05 |
| SNRPB | SNs | 2.05616262 | 0.00878693 |
| SP1 | SNs | 2.212760208 | 0.00612689 |
| SP2 | SNs | 2.122652306 | 0.00753959 |
| SPARC | SNs | 2.363585094 | 0.00432927 |
| SPI1 | SNs | 7.237619426 | 5.79E-08 |
| SPINT1 | SNs | 4.07615855 | 8.39E-05 |
| SPP1 | SNs | 3.850196176 | 0.00014119 |
| SRF | SNs | 3.210869315 | 0.00061536 |
| SRPRA | SNs | 3.67933398 | 0.00020925 |
| TRIM21 | SNs | 3.561564207 | 0.00027443 |
| STAT3 | SNs | 4.006435173 | 9.85E-05 |
| STAT5A | SNs | 3.441402399 | 0.00036191 |
| STAT6 | SNs | 5.353365161 | 4.43E-06 |
| STK10 | SNs | 5.157578027 | 6.96E-06 |
| AURKC | SNs | 2.125754221 | 0.00748593 |
| SULT1A2 | SNs | 3.629194934 | 0.00023486 |
| STX4 | SNs | 2.072327103 | 0.0084659 |
| STX5 | SNs | 3.940467953 | 0.00011469 |
| STXBP2 | SNs | 5.743189387 | 1.81E-06 |
| SULT1A1 | SNs | 4.192413321 | 6.42E-05 |
| SUPT6H | SNs | 2.267923433 | 0.00539606 |
| SURF4 | SNs | 3.394129388 | 0.00040353 |
| SYK | SNs | 3.998732605 | 0.00010029 |
| TAF10 | SNs | 2.938695856 | 0.00115161 |
| TALDO1 | SNs | 6.181038003 | 6.59E-07 |
| TAPBP | SNs | 4.010041723 | 9.77E-05 |
| TBXA2R | SNs | 2.169742639 | 0.00676484 |
| TBXAS1 | SNs | 4.389459359 | 4.08E-05 |
| TCN2 | SNs | 3.231977663 | 0.00058617 |
| TGFB1 | SNs | 3.601918149 | 0.00025008 |
| TGM3 | SNs | 4.373701933 | 4.23E-05 |
| THBS1 | SNs | 3.085583614 | 0.00082114 |
| THBS3 | SNs | 2.349649767 | 0.00447044 |
| TIMP2 | SNs | 6.393465907 | 4.04E-07 |
| TKT | SNs | 5.973876409 | 1.06E-06 |
| TLE3 | SNs | 6.416401564 | 3.83E-07 |
| TLN1 | SNs | 3.852178927 | 0.00014055 |
| CLDN5 | SNs | 2.745645805 | 0.0017962 |

| | | | |
|---|---|---|---|
| TNFAIP2 | SNs | 3.485983165 | 0.0003266 |
| TNFRSF1A | SNs | 5.176089667 | 6.67E-06 |
| TNFRSF1B | SNs | 5.26269176 | 5.46E-06 |
| TNNI2 | SNs | 3.943732425 | 0.00011383 |
| TOP3A | SNs | 3.498402385 | 0.00031739 |
| TPD52L2 | SNs | 4.330631097 | 4.67E-05 |
| TPI1 | SNs | 1.979076107 | 0.01049359 |
| TPSAB1 | SNs | 1.990342453 | 0.01022486 |
| TRPC2 | SNs | 2.113281367 | 0.00770404 |
| TRPM2 | SNs | 2.999583296 | 0.00100096 |
| TSC2 | SNs | 2.286815658 | 0.00516636 |
| TST | SNs | 4.03558129 | 9.21E-05 |
| TUBA4A | SNs | 4.508065912 | 3.10E-05 |
| TUFM | SNs | 1.992903887 | 0.01016474 |
| TYK2 | SNs | 4.582904385 | 2.61E-05 |
| TYROBP | SNs | 4.966486624 | 1.08E-05 |
| UBE2L3 | SNs | 2.255124772 | 0.00555745 |
| USP4 | SNs | 2.155079812 | 0.00699713 |
| NR1H2 | SNs | 2.790955697 | 0.00161825 |
| UPK3A | SNs | 3.296047171 | 0.00050577 |
| USF1 | SNs | 3.907630874 | 0.0001237 |
| USF2 | SNs | 2.916483906 | 0.00121204 |
| VASP | SNs | 7.87577941 | 1.33E-08 |
| VAV1 | SNs | 2.958301823 | 0.00110077 |
| VCP | SNs | 2.092032975 | 0.00809034 |
| VDR | SNs | 2.007458722 | 0.00982972 |
| BEST1 | SNs | 3.577583493 | 0.00026449 |
| VWF | SNs | 3.656420953 | 0.00022059 |
| WIPF1 | SNs | 2.756051481 | 0.00175367 |
| CLIP2 | SNs | 2.909853405 | 0.00123068 |
| LAT2 | SNs | 3.775706512 | 0.00016761 |
| XRCC1 | SNs | 2.699214075 | 0.00199888 |
| ZFP36 | SNs | 3.152174515 | 0.00070441 |
| TRIM25 | SNs | 2.493237042 | 0.00321191 |
| ZNF213 | SNs | 2.250265911 | 0.00561997 |
| ZYX | SNs | 6.037279146 | 9.18E-07 |
| LAPTM5 | SNs | 4.30524756 | 4.95E-05 |
| TUBA1A | SNs | 5.454008104 | 3.52E-06 |
| MAPKAPK3 | SNs | 4.547464983 | 2.83E-05 |
| RAB7A | SNs | 2.199735851 | 0.00631341 |
| PRRC2A | SNs | 2.985592802 | 0.00103373 |
| ABHD16A | SNs | 2.339304514 | 0.00457821 |
| LST1 | SNs | 3.6084452 | 0.00024635 |
| TFEB | SNs | 5.775901243 | 1.68E-06 |
| NUP214 | SNs | 3.75539622 | 0.00017563 |
| MLF2 | SNs | 5.818546792 | 1.52E-06 |
| ELL | SNs | 3.647986557 | 0.00022491 |
| DGCR6 | SNs | 2.143296129 | 0.00718959 |
| DYSF | SNs | 5.678872402 | 2.09E-06 |
| AXIN1 | SNs | 2.154987192 | 0.00699863 |
| BAP1 | SNs | 2.190775379 | 0.00644503 |
| H2AC18 | SNs | 3.259174943 | 0.00055059 |
| SLC25A11 | SNs | 3.390224135 | 0.00040717 |
| TAGLN2 | SNs | 4.91660963 | 1.21E-05 |
| ULK1 | SNs | 4.786499275 | 1.63E-05 |
| TPST2 | SNs | 2.272843929 | 0.00533527 |
| GAS7 | SNs | 3.486298332 | 0.00032636 |
| FAM193A | SNs | 2.705138358 | 0.00197179 |
| EIF4EBP3 | SNs | 2.143293274 | 0.00718963 |
| STX10 | SNs | 5.206843048 | 6.21E-06 |
| MARCO | SNs | 2.790175254 | 0.00162116 |
| DGAT1 | SNs | 3.769732018 | 0.00016993 |
| S1PR4 | SNs | 3.327449501 | 0.00047049 |
| ABCC3 | SNs | 2.087997537 | 0.00816587 |
| TNFSF13 | SNs | 5.068909684 | 8.53E-06 |
| ADAM15 | SNs | 3.716809484 | 0.00019195 |
| NAPA | SNs | 2.762464292 | 0.00172797 |
| SIGLEC5 | SNs | 2.705195754 | 0.00197153 |
| TNFRSF10C | SNs | 6.354865796 | 4.42E-07 |
| NRP1 | SNs | 2.125850349 | 0.00748427 |
| TMEM11 | SNs | 3.413036332 | 0.00038633 |

| | | | |
|---|---|---|---|
| HDAC3 | SNs | 3.135763231 | 0.00073154 |
| HCAR3 | SNs | 2.131795325 | 0.00738252 |
| SQSTM1 | SNs | 4.72462257 | 1.89E-05 |
| MTMR3 | SNs | 2.025982081 | 0.00941928 |
| AP1M1 | SNs | 2.323180374 | 0.00475138 |
| AP3D1 | SNs | 2.068989512 | 0.00853121 |
| PLOD3 | SNs | 2.372642964 | 0.00423991 |
| PGLYRP1 | SNs | 2.806438244 | 0.00156157 |
| MPZL1 | SNs | 2.304333493 | 0.00496211 |
| SOCS3 | SNs | 2.991942351 | 0.00101873 |
| UBE2M | SNs | 4.334669701 | 4.63E-05 |
| PSTPIP1 | SNs | 4.428810083 | 3.73E-05 |
| SLC7A7 | SNs | 2.07699423 | 0.0083754 |
| CLDN9 | SNs | 3.956404543 | 0.00011056 |
| SART1 | SNs | 2.588052862 | 0.00258195 |
| UNC119 | SNs | 5.786419523 | 1.64E-06 |
| ATP6V0D1 | SNs | 5.976006916 | 1.06E-06 |
| SLC16A5 | SNs | 5.01621506 | 9.63E-06 |
| SLC16A3 | SNs | 4.947987743 | 1.13E-05 |
| PDLIM1 | SNs | 3.307296483 | 0.00049284 |
| SYNGR2 | SNs | 2.555885438 | 0.00278045 |
| HGS | SNs | 2.93192719 | 0.0011697 |
| DYRK1B | SNs | 3.239298893 | 0.00057637 |
| CTDP1 | SNs | 4.483713682 | 3.28E-05 |
| LPAR2 | SNs | 3.776265976 | 0.00016739 |
| ARHGEF2 | SNs | 2.55022878 | 0.0028169 |
| DEDD | SNs | 2.236841375 | 0.0057964 |
| RAB11B | SNs | 3.299628805 | 0.00050162 |
| PDLIM7 | SNs | 6.875188351 | 1.33E-07 |
| MAPKAPK2 | SNs | 4.903598295 | 1.25E-05 |
| CYTH2 | SNs | 3.537161498 | 0.00029029 |
| ATP6V1F | SNs | 2.095171303 | 0.00803209 |
| NHERF1 | SNs | 2.953288116 | 0.00111356 |
| TJP2 | SNs | 2.964901198 | 0.00108417 |
| KCNK6 | SNs | 3.576288013 | 0.00026528 |
| HOMER3 | SNs | 3.770263483 | 0.00016972 |
| THEMIS2 | SNs | 4.262202583 | 5.47E-05 |
| PGS1 | SNs | 2.199960175 | 0.00631015 |
| FXR2 | SNs | 2.513855421 | 0.00306298 |
| LITAF | SNs | 3.232978451 | 0.00058482 |
| TP53I11 | SNs | 4.362367276 | 4.34E-05 |
| RAB3D | SNs | 3.114700154 | 0.00076789 |
| GTPBP1 | SNs | 3.720015771 | 0.00019054 |
| PITPNM1 | SNs | 2.346097639 | 0.00450715 |
| VPS9D1 | SNs | 2.224192469 | 0.00596771 |
| RIN1 | SNs | 4.041281559 | 9.09E-05 |
| AATK | SNs | 4.198281567 | 6.33E-05 |
| ISG15 | SNs | 2.002143053 | 0.00995078 |
| ZNF592 | SNs | 5.174567661 | 6.69E-06 |
| PPM1F | SNs | 2.961776004 | 0.001092 |
| SLC25A44 | SNs | 4.937022757 | 1.16E-05 |
| KIAA0040 | SNs | 2.861483793 | 0.00137568 |
| FAM53B | SNs | 2.027078474 | 0.00939554 |
| N4BP1 | SNs | 4.854755887 | 1.40E-05 |
| PPP6R2 | SNs | 2.283875133 | 0.00520146 |
| DHX34 | SNs | 5.247101335 | 5.66E-06 |
| ZNF646 | SNs | 2.223173763 | 0.00598172 |
| TBKBP1 | SNs | 2.859276955 | 0.00138268 |
| KMT2B | SNs | 3.447074979 | 0.00035721 |
| KIAA0513 | SNs | 5.671407679 | 2.13E-06 |
| ATG13 | SNs | 4.603530261 | 2.49E-05 |
| IST1 | SNs | 3.199170886 | 0.00063216 |
| IP6K1 | SNs | 5.556225577 | 2.78E-06 |
| RNF40 | SNs | 4.873641497 | 1.34E-05 |
| CTIF | SNs | 2.771130882 | 0.00169383 |
| SPATA2 | SNs | 3.377801468 | 0.00041899 |
| ARHGEF11 | SNs | 3.186635007 | 0.00065068 |
| PLEKHM1 | SNs | 5.209657784 | 6.17E-06 |
| ELMO1 | SNs | 2.81054046 | 0.00154689 |
| GAB2 | SNs | 4.567342924 | 2.71E-05 |
| RUSC2 | SNs | 2.153373254 | 0.00702468 |

| | | | |
|---|---|---|---|
| SETDB1 | SNs | 2.398747852 | 0.00399257 |
| TRANK1 | SNs | 2.11363839 | 0.00769771 |
| OSBPL2 | SNs | 3.668791954 | 0.00021439 |
| TECPR2 | SNs | 4.925292714 | 1.19E-05 |
| KLHL21 | SNs | 4.535401148 | 2.91E-05 |
| ARHGAP25 | SNs | 2.710594407 | 0.00194718 |
| WDR1 | SNs | 6.304895011 | 4.96E-07 |
| MVP | SNs | 3.581036301 | 0.0002624 |
| DGCR2 | SNs | 5.539144646 | 2.89E-06 |
| SCO2 | SNs | 2.793729364 | 0.00160794 |
| ACOT8 | SNs | 2.238211856 | 0.00577814 |
| SRA1 | SNs | 2.818808315 | 0.00151772 |
| HDAC5 | SNs | 4.882963043 | 1.31E-05 |
| BCL2L11 | SNs | 2.124607663 | 0.00750572 |
| FRAT1 | SNs | 2.59009866 | 0.00256981 |
| MED16 | SNs | 2.736524577 | 0.00183432 |
| TOM1 | SNs | 5.840505783 | 1.44E-06 |
| SH2D3C | SNs | 3.716652522 | 0.00019202 |
| AP1M2 | SNs | 2.352712998 | 0.00443902 |
| SCAMP2 | SNs | 3.938450992 | 0.00011523 |
| KCNK7 | SNs | 3.741377849 | 0.00018139 |
| ARPC4 | SNs | 3.869879971 | 0.00013493 |
| ARPC1B | SNs | 3.66997551 | 0.00021381 |
| PPIF | SNs | 5.48311388 | 3.29E-06 |
| CTDSP2 | SNs | 2.764131837 | 0.00172135 |
| ACTR1A | SNs | 3.440428241 | 0.00036272 |
| LPCAT3 | SNs | 5.628956409 | 2.35E-06 |
| PSME3 | SNs | 2.331234239 | 0.00466408 |
| FLOT1 | SNs | 5.678825978 | 2.09E-06 |
| CTDSPL | SNs | 3.243642148 | 0.00057063 |
| PLIN3 | SNs | 6.086550539 | 8.19E-07 |
| MFSD10 | SNs | 2.34036603 | 0.00456703 |
| SF3B4 | SNs | 3.263446754 | 0.0005452 |
| CDK2AP2 | SNs | 2.437704532 | 0.00365002 |
| RGS19 | SNs | 3.048393489 | 0.00089455 |
| LILRB2 | SNs | 3.712117222 | 0.00019404 |
| BCKDK | SNs | 6.546939839 | 2.84E-07 |
| KATNB1 | SNs | 2.543598519 | 0.00286023 |
| APBB3 | SNs | 2.102655054 | 0.00789487 |
| TCIRG1 | SNs | 3.717440831 | 0.00019167 |
| TNIP1 | SNs | 2.244425607 | 0.00569606 |
| SIRPB1 | SNs | 5.753169322 | 1.77E-06 |
| IRAG1 | SNs | 2.560775729 | 0.00274931 |
| HMG20B | SNs | 2.517743692 | 0.00303568 |
| MICU1 | SNs | 3.775549472 | 0.00016767 |
| TUBA1B | SNs | 3.84836544 | 0.00014179 |
| IRF9 | SNs | 2.759210391 | 0.00174096 |
| TUBB4B | SNs | 4.026253372 | 9.41E-05 |
| NDRG1 | SNs | 2.998370918 | 0.00100376 |
| MYL9 | SNs | 2.662403719 | 0.00217569 |
| BASP1 | SNs | 3.040334489 | 0.00091131 |
| CD2BP2 | SNs | 3.396906341 | 0.00040095 |
| UBAC1 | SNs | 3.601336408 | 0.00025042 |
| CDIPT | SNs | 4.811094974 | 1.54E-05 |
| CDC42EP2 | SNs | 4.934211376 | 1.16E-05 |
| IFI30 | SNs | 4.47016018 | 3.39E-05 |
| ZER1 | SNs | 2.498519661 | 0.00317308 |
| TACC3 | SNs | 5.050382172 | 8.90E-06 |
| TADA3 | SNs | 2.029494413 | 0.00934341 |
| NXF1 | SNs | 2.41338706 | 0.00386023 |
| CRTAP | SNs | 2.366744359 | 0.00429789 |
| SEMA4D | SNs | 2.328819774 | 0.00469008 |
| SEMA4B | SNs | 4.844099456 | 1.43E-05 |
| KAT5 | SNs | 2.286050516 | 0.00517547 |
| ATG7 | SNs | 5.500730051 | 3.16E-06 |
| TM9SF1 | SNs | 3.152128096 | 0.00070449 |
| ARPC1A | SNs | 2.949401745 | 0.00112357 |
| AGPAT1 | SNs | 3.503400884 | 0.00031376 |
| AGPAT2 | SNs | 3.745264102 | 0.00017978 |
| MRPL28 | SNs | 3.353702304 | 0.00044289 |
| IFITM2 | SNs | 6.120621099 | 7.57E-07 |

| | | | |
|---|---|---|---|
| SH2B2 | SNs | 4.086256754 | 8.20E-05 |
| ST6GALNAC2 | SNs | 2.302413826 | 0.00498409 |
| RBCK1 | SNs | 3.786041785 | 0.00016367 |
| ARID3B | SNs | 4.213219715 | 6.12E-05 |
| GAS2L1 | SNs | 4.934342562 | 1.16E-05 |
| RGS14 | SNs | 5.238081664 | 5.78E-06 |
| CAMKK2 | SNs | 4.46496191 | 3.43E-05 |
| SPINT2 | SNs | 2.392224765 | 0.00405299 |
| TNFSF13B | SNs | 2.233431709 | 0.00584209 |
| CNPY3 | SNs | 5.195458983 | 6.38E-06 |
| KIF1C | SNs | 2.222137512 | 0.00599601 |
| STARD10 | SNs | 2.815612275 | 0.00152893 |
| ARID5A | SNs | 3.053952481 | 0.00088318 |
| TSPAN9 | SNs | 2.678032171 | 0.00209878 |
| USP19 | SNs | 3.587094806 | 0.00025876 |
| CD300C | SNs | 2.506667885 | 0.0031141 |
| TRAFD1 | SNs | 3.302929519 | 0.00049782 |
| PNPLA6 | SNs | 2.094830397 | 0.0080384 |
| EHD1 | SNs | 3.205903462 | 0.00062244 |
| KDELR1 | SNs | 2.535444605 | 0.00291444 |
| STARD3 | SNs | 2.272205093 | 0.00534312 |
| PDIA5 | SNs | 2.067008052 | 0.00857022 |
| OS9 | SNs | 6.155312685 | 6.99E-07 |
| LMAN2 | SNs | 3.399323504 | 0.00039873 |
| CKAP4 | SNs | 3.1358344 | 0.00073142 |
| SLC27A3 | SNs | 2.529784204 | 0.00295268 |
| LILRB4 | SNs | 2.997791786 | 0.0010051 |
| RAB35 | SNs | 3.803887729 | 0.00015708 |
| LILRA1 | SNs | 4.875341876 | 1.33E-05 |
| LILRB3 | SNs | 5.63713703 | 2.31E-06 |
| LILRA2 | SNs | 7.688438577 | 2.05E-08 |
| ADAP1 | SNs | 3.049403782 | 0.00089248 |
| ADRM1 | SNs | 2.775971808 | 0.00167505 |
| OGFR | SNs | 3.290314648 | 0.00051249 |
| WWP2 | SNs | 5.323079253 | 4.75E-06 |
| TMEM115 | SNs | 3.546623584 | 0.00028404 |
| TRIOBP | SNs | 3.959257125 | 0.00010984 |
| HNRNPUL1 | SNs | 2.558127296 | 0.00276613 |
| CLASRP | SNs | 2.027231101 | 0.00939223 |
| CDC42EP1 | SNs | 2.964129074 | 0.0010861 |
| CORO1A | SNs | 4.692883543 | 2.03E-05 |
| PTP4A3 | SNs | 2.343732199 | 0.00453177 |
| BAZ2A | SNs | 2.022694488 | 0.00949086 |
| RNF24 | SNs | 3.833601965 | 0.00014669 |
| PMF1 | SNs | 3.004871036 | 0.00098885 |
| PACSIN2 | SNs | 3.33531446 | 0.00046205 |
| NRM | SNs | 2.792272775 | 0.00161334 |
| TREX1 | SNs | 2.400299289 | 0.00397833 |
| MGAT4B | SNs | 2.18265716 | 0.00656663 |
| PNKP | SNs | 2.593878376 | 0.00254754 |
| LYPLA2 | SNs | 3.863700464 | 0.00013687 |
| CD300A | SNs | 2.12791111 | 0.00744884 |
| COPE | SNs | 3.885506579 | 0.00013016 |
| EXOC3 | SNs | 2.78830703 | 0.00162814 |
| GABARAP | SNs | 4.196718674 | 6.36E-05 |
| MGLL | SNs | 2.401764053 | 0.00396493 |
| TWF2 | SNs | 4.262463042 | 5.46E-05 |
| CASC3 | SNs | 2.264390986 | 0.00544013 |
| COPG1 | SNs | 2.568627343 | 0.00270006 |
| DLGAP4 | SNs | 2.278038997 | 0.00527183 |
| NLRP1 | SNs | 1.986303186 | 0.01032041 |
| PPP6R1 | SNs | 2.89658984 | 0.00126885 |
| MLXIP | SNs | 2.736965278 | 0.00183246 |
| MON1B | SNs | 3.5169909 | 0.00030409 |
| ABLIM3 | SNs | 2.366898617 | 0.00429637 |
| RPH3A | SNs | 2.154300441 | 0.0070097 |
| DENND3 | SNs | 3.88913586 | 0.00012908 |
| SBNO2 | SNs | 2.902624954 | 0.00125134 |
| RALY | SNs | 2.229933294 | 0.00588934 |
| MAPRE3 | SNs | 2.219798309 | 0.00602839 |
| SIRT2 | SNs | 2.929848445 | 0.00117531 |

| | | | |
|---|---|---|---|
| SCAP | SNs | 2.639560372 | 0.00229319 |
| TCF25 | SNs | 2.484296629 | 0.00327871 |
| ACIN1 | SNs | 2.868922866 | 0.00135231 |
| KDM4B | SNs | 3.70875006 | 0.00019555 |
| MAST3 | SNs | 2.73872162 | 0.00182507 |
| WDTC1 | SNs | 5.287659859 | 5.16E-06 |
| ZSWIM8 | SNs | 3.43340193 | 0.00036864 |
| CAMTA2 | SNs | 4.196705428 | 6.36E-05 |
| PLXND1 | SNs | 2.316499001 | 0.00482504 |
| ATG2A | SNs | 2.274516248 | 0.00531476 |
| KDM6B | SNs | 4.119589983 | 7.59E-05 |
| ZC3H3 | SNs | 3.48732428 | 0.00032559 |
| FCHO1 | SNs | 4.051648644 | 8.88E-05 |
| CIC | SNs | 2.597504716 | 0.00252636 |
| GGA3 | SNs | 2.565259914 | 0.00272107 |
| STAB1 | SNs | 2.944703405 | 0.00113579 |
| ATG4B | SNs | 2.695292935 | 0.00201701 |
| ACSBG1 | SNs | 2.79637131 | 0.00159819 |
| PLEKHM2 | SNs | 3.563784084 | 0.00027303 |
| JMJD6 | SNs | 2.258056883 | 0.00552005 |
| XPO6 | SNs | 5.301299935 | 5.00E-06 |
| NBEAL2 | SNs | 3.074506728 | 0.00084235 |
| RRP12 | SNs | 3.266097204 | 0.00054188 |
| KAZN | SNs | 2.307526786 | 0.00492576 |
| IQCE | SNs | 2.100553557 | 0.00793316 |
| MGRN1 | SNs | 4.001872692 | 9.96E-05 |
| FKBP15 | SNs | 2.107705277 | 0.0078036 |
| SIN3B | SNs | 2.380141239 | 0.00416734 |
| KIAA0930 | SNs | 2.43455157 | 0.00367662 |
| SLC9A8 | SNs | 3.710572096 | 0.00019473 |
| VPS39 | SNs | 3.026794223 | 0.00094017 |
| SRGAP2 | SNs | 2.845903091 | 0.00142593 |
| SMG5 | SNs | 2.151348769 | 0.00705751 |
| MAU2 | SNs | 3.345047469 | 0.00045181 |
| NCSTN | SNs | 8.21754866 | 6.06E-09 |
| PIP5K1C | SNs | 3.140293206 | 0.00072395 |
| CTDNEP1 | SNs | 2.678034236 | 0.00209877 |
| FRAT2 | SNs | 3.755859317 | 0.00017544 |
| COTL1 | SNs | 7.080500251 | 8.31E-08 |
| CBX7 | SNs | 2.173284543 | 0.00670989 |
| HAAO | SNs | 3.384809956 | 0.00041228 |
| KCTD2 | SNs | 3.355167933 | 0.0004414 |
| CABIN1 | SNs | 3.41321398 | 0.00038618 |
| SRRM2 | SNs | 2.121638751 | 0.00755721 |
| PIK3R5 | SNs | 2.326931605 | 0.00471052 |
| PSD4 | SNs | 2.749159938 | 0.00178172 |
| WBP2 | SNs | 6.20106684 | 6.29E-07 |
| DDAH2 | SNs | 2.887152819 | 0.00129672 |
| PADI4 | SNs | 2.705215296 | 0.00197145 |
| CARHSP1 | SNs | 2.566179093 | 0.00271532 |
| DAPK2 | SNs | 3.286216363 | 0.00051735 |
| SH3BP1 | SNs | 2.432572525 | 0.00369341 |
| PPP1R15A | SNs | 8.376709989 | 4.20E-09 |
| PLD3 | SNs | 3.012162247 | 0.00097238 |
| PISD | SNs | 3.422968584 | 0.0003776 |
| IL17RA | SNs | 4.316737193 | 4.82E-05 |
| TFIP11 | SNs | 3.003991504 | 0.00099085 |
| TBC1D22A | SNs | 4.046990943 | 8.97E-05 |
| PGLS | SNs | 3.263766954 | 0.00054479 |
| BRI3 | SNs | 4.769498528 | 1.70E-05 |
| TMEM184B | SNs | 3.925428839 | 0.00011873 |
| YIPF3 | SNs | 5.250743506 | 5.61E-06 |
| BRMS1 | SNs | 3.301413773 | 0.00049956 |
| PRKD2 | SNs | 2.371584697 | 0.00425026 |
| ZDHHC5 | SNs | 2.556885382 | 0.00277405 |
| ZNF385A | SNs | 4.19624354 | 6.36E-05 |
| PNKD | SNs | 3.949769232 | 0.00011226 |
| DHRS7B | SNs | 2.934511358 | 0.00116276 |
| TBC1D10B | SNs | 5.783279457 | 1.65E-06 |
| RNF167 | SNs | 4.552336366 | 2.80E-05 |
| LRP10 | SNs | 5.70114478 | 1.99E-06 |

| | | | |
|---|---|---|---|
| PLEKHG3 | SNs | 3.118006905 | 0.00076207 |
| SIPA1L1 | SNs | 2.558561339 | 0.00276337 |
| POLDIP2 | SNs | 3.048028919 | 0.00089531 |
| GGA1 | SNs | 2.453189104 | 0.00352217 |
| CCDC9 | SNs | 2.802980936 | 0.00157405 |
| SZRD1 | SNs | 2.550733921 | 0.00281362 |
| WIPI2 | SNs | 3.662316182 | 0.00021761 |
| CCDC69 | SNs | 2.81291649 | 0.00153845 |
| TRPC4AP | SNs | 4.520290873 | 3.02E-05 |
| MTG2 | SNs | 5.417506347 | 3.82E-06 |
| INTS1 | SNs | 2.023918951 | 0.00946414 |
| TINF2 | SNs | 2.186539096 | 0.0065082 |
| GBGT1 | SNs | 2.840222138 | 0.0014447 |
| PTPN18 | SNs | 3.847533227 | 0.00014206 |
| NARF | SNs | 2.802743687 | 0.00157491 |
| AKAP8L | SNs | 2.068644614 | 0.00853798 |
| USP21 | SNs | 2.024630765 | 0.00944864 |
| SIGLEC7 | SNs | 3.508922507 | 0.0003098 |
| LAMP3 | SNs | 2.122242324 | 0.00754671 |
| CYTH4 | SNs | 6.176262417 | 6.66E-07 |
| CIDEB | SNs | 4.593740631 | 2.55E-05 |
| SLC39A1 | SNs | 3.292979388 | 0.00050936 |
| SIGLEC9 | SNs | 5.010559181 | 9.76E-06 |
| C5AR2 | SNs | 3.254811106 | 0.00055615 |
| GPR162 | SNs | 3.036344604 | 0.00091972 |
| CHMP2A | SNs | 5.231710536 | 5.87E-06 |
| SMPDL3B | SNs | 4.196068299 | 6.37E-05 |
| TOR2A | SNs | 3.952309904 | 0.00011161 |
| SLCO3A1 | SNs | 3.731880612 | 0.0001854 |
| NKIRAS2 | SNs | 3.906955826 | 0.00012389 |
| DBNL | SNs | 5.107190982 | 7.81E-06 |
| SLC43A3 | SNs | 2.014129922 | 0.00967988 |
| PYCARD | SNs | 3.740078241 | 0.00018194 |
| SAP30BP | SNs | 3.621447999 | 0.00023908 |
| PARVB | SNs | 4.902254786 | 1.25E-05 |
| CYP2S1 | SNs | 2.383563437 | 0.00413463 |
| UBN1 | SNs | 4.423731645 | 3.77E-05 |
| STRN4 | SNs | 5.273890585 | 5.32E-06 |
| SNX11 | SNs | 2.826967202 | 0.00148947 |
| EPN1 | SNs | 3.156075001 | 0.00069811 |
| OSGIN1 | SNs | 3.207815824 | 0.0006197 |
| CERS2 | SNs | 1.987582768 | 0.01029004 |
| NRBP1 | SNs | 3.698491105 | 0.00020022 |
| SLC39A3 | SNs | 2.108068292 | 0.00779707 |
| PILRA | SNs | 5.146089384 | 7.14E-06 |
| TAX1BP3 | SNs | 5.172165286 | 6.73E-06 |
| CHST11 | SNs | 2.826863194 | 0.00148983 |
| DEF6 | SNs | 2.632592459 | 0.00233028 |
| ZFTRAF1 | SNs | 3.441315538 | 0.00036198 |
| COPS7A | SNs | 2.806820724 | 0.0015602 |
| F11R | SNs | 2.214120232 | 0.00610773 |
| ZBTB7B | SNs | 4.24960404 | 5.63E-05 |
| JPT1 | SNs | 6.523538498 | 3.00E-07 |
| EGFL7 | SNs | 2.599474886 | 0.00251493 |
| PLEKHO1 | SNs | 3.018021124 | 0.00095935 |
| RAPGEFL1 | SNs | 2.373159113 | 0.00423488 |
| GP6 | SNs | 2.990013497 | 0.00102326 |
| VRK3 | SNs | 2.768951609 | 0.00170235 |
| SHISA5 | SNs | 4.452894286 | 3.52E-05 |
| MARCHF2 | SNs | 2.68621201 | 0.00205962 |
| UBAP1 | SNs | 3.770246792 | 0.00016973 |
| C1RL | SNs | 2.943526253 | 0.00113887 |
| SCAND1 | SNs | 2.058670906 | 0.00873633 |
| GMIP | SNs | 5.455315346 | 3.50E-06 |
| SLC15A3 | SNs | 4.324946469 | 4.73E-05 |
| FAM53C | SNs | 3.112754923 | 0.00077134 |
| PHF21A | SNs | 3.348584689 | 0.00044814 |
| SPG21 | SNs | 3.353762397 | 0.00044283 |
| FZR1 | SNs | 2.503141785 | 0.00313948 |
| CHST15 | SNs | 2.148790932 | 0.00709919 |
| WDR83OS | SNs | 3.28731507 | 0.00051604 |

| | | | |
|---|---|---|---|
| RTF2 | SNs | 2.391237892 | 0.00406221 |
| SIRT7 | SNs | 2.381479892 | 0.00415451 |
| SIRT6 | SNs | 2.438112135 | 0.0036466 |
| HDAC7 | SNs | 2.823438213 | 0.00150163 |
| PIGT | SNs | 2.085896321 | 0.00820547 |
| CYB5R1 | SNs | 2.510392731 | 0.0030875 |
| MSRB1 | SNs | 5.663082681 | 2.17E-06 |
| INPP5K | SNs | 3.538035683 | 0.00028971 |
| TUBA8 | SNs | 5.660808355 | 2.18E-06 |
| PTOV1 | SNs | 3.359320074 | 0.0004372 |
| FXYD6 | SNs | 3.595019665 | 0.00025409 |
| RAB4B | SNs | 3.448758212 | 0.00035583 |
| RAB24 | SNs | 3.232738735 | 0.00058514 |
| TREM1 | SNs | 3.884960718 | 0.00013033 |
| YIPF1 | SNs | 2.354608225 | 0.00441969 |
| PRR13 | SNs | 4.627558594 | 2.36E-05 |
| TOLLIP | SNs | 3.566157352 | 0.00027155 |
| CHPF2 | SNs | 4.915403298 | 1.22E-05 |
| SMOX | SNs | 2.73792627 | 0.00182841 |
| ADAMTSL4 | SNs | 5.05116349 | 8.89E-06 |
| APBB1IP | SNs | 4.643693631 | 2.27E-05 |
| FAM193B | SNs | 2.402928883 | 0.00395431 |
| NECAB2 | SNs | 3.166937153 | 0.00068087 |
| PAF1 | SNs | 1.989605935 | 0.01024222 |
| PARP14 | SNs | 2.138594179 | 0.00726785 |
| GTPBP2 | SNs | 2.450351312 | 0.00354526 |
| MANSC1 | SNs | 2.356962713 | 0.00439579 |
| XAF1 | SNs | 2.169325878 | 0.00677133 |
| PPP1R12C | SNs | 2.943599567 | 0.00113868 |
| DNAJB12 | SNs | 3.840341505 | 0.00014443 |
| TRPM4 | SNs | 2.043846762 | 0.00903968 |
| GATAD2A | SNs | 5.226214484 | 5.94E-06 |
| NDE1 | SNs | 2.73829572 | 0.00182686 |
| DEF8 | SNs | 3.601775835 | 0.00025016 |
| TOR4A | SNs | 2.314872242 | 0.00484315 |
| ALKBH5 | SNs | 2.128764535 | 0.00743422 |
| DUSP23 | SNs | 2.091716168 | 0.00809625 |
| ADPRS | SNs | 2.926452921 | 0.00118453 |
| SSH3 | SNs | 3.467508889 | 0.00034079 |
| BANP | SNs | 2.079069215 | 0.00833548 |
| ADISSP | SNs | 2.229425703 | 0.00589623 |
| SLC35F6 | SNs | 2.965890604 | 0.00108171 |
| CLN6 | SNs | 3.4001137 | 0.000398 |
| AURKAIP1 | SNs | 2.204470751 | 0.00624495 |
| LAMTOR1 | SNs | 5.995769512 | 1.01E-06 |
| VPS37C | SNs | 4.98846697 | 1.03E-05 |
| WIPI1 | SNs | 2.857105574 | 0.00138961 |
| RNF31 | SNs | 3.650614849 | 0.00022356 |
| TAPBPL | SNs | 2.238726284 | 0.0057713 |
| SAMD4B | SNs | 4.607292286 | 2.47E-05 |
| XKR8 | SNs | 3.369996045 | 0.00042658 |
| RBM23 | SNs | 2.818725422 | 0.00151801 |
| NADSYN1 | SNs | 2.890995816 | 0.0012853 |
| MAP1S | SNs | 2.599799566 | 0.00251305 |
| TRAPPC14 | SNs | 4.865555823 | 1.36E-05 |
| TMEM140 | SNs | 2.758654581 | 0.00174319 |
| TBC1D2 | SNs | 3.03077475 | 0.00093159 |
| PI4K2A | SNs | 2.13510811 | 0.00732642 |
| TMCO6 | SNs | 2.58990332 | 0.00257097 |
| NAGK | SNs | 2.362469509 | 0.00434041 |
| OTUB1 | SNs | 2.889778778 | 0.00128891 |
| ZDHHC7 | SNs | 2.882815305 | 0.00130974 |
| RAB20 | SNs | 2.724584974 | 0.00188545 |
| TMEM127 | SNs | 6.019207256 | 9.57E-07 |
| NPLOC4 | SNs | 3.14692572 | 0.00071297 |
| PACS1 | SNs | 4.64893738 | 2.24E-05 |
| MAP7D1 | SNs | 3.825609906 | 0.00014941 |
| ARHGEF40 | SNs | 5.106857438 | 7.82E-06 |
| EDEM2 | SNs | 3.655742105 | 0.00022093 |
| MINDY1 | SNs | 5.364253837 | 4.32E-06 |
| ADAP2 | SNs | 2.276909325 | 0.00528556 |

| | | | |
|---|---|---|---|
| PSENEN | SNs | 3.911157942 | 0.0001227 |
| ACSS2 | SNs | 4.040543357 | 9.11E-05 |
| BIN3 | SNs | 3.602780975 | 0.00024959 |
| APOBR | SNs | 5.337500312 | 4.60E-06 |
| INKA2 | SNs | 2.530605826 | 0.0029471 |
| ZMAT5 | SNs | 2.507396179 | 0.00310888 |
| LIN37 | SNs | 3.114545727 | 0.00076816 |
| NSFL1C | SNs | 4.791074535 | 1.62E-05 |
| TMEM234 | SNs | 2.227551257 | 0.00592173 |
| MEPCE | SNs | 2.170196259 | 0.00675778 |
| PANX2 | SNs | 3.423918684 | 0.00037677 |
| RETN | SNs | 2.573877066 | 0.00266761 |
| GPR137 | SNs | 3.190504632 | 0.0006449 |
| C15orf39 | SNs | 6.267953735 | 5.40E-07 |
| NCLN | SNs | 2.402426041 | 0.00395889 |
| GPR108 | SNs | 3.952585723 | 0.00011154 |
| ATXN7L3 | SNs | 2.631721008 | 0.00233496 |
| SLC12A9 | SNs | 4.618708381 | 2.41E-05 |
| COQ8A | SNs | 2.55540904 | 0.0027835 |
| CHRNA10 | SNs | 2.356098964 | 0.00440454 |
| AGTRAP | SNs | 4.408374462 | 3.91E-05 |
| PNPLA2 | SNs | 2.374853038 | 0.00421839 |
| APMAP | SNs | 2.266946293 | 0.00540821 |
| RGL3 | SNs | 2.007849587 | 0.00982088 |
| RNPEPL1 | SNs | 2.480690241 | 0.00330605 |
| SLC44A2 | SNs | 5.208065464 | 6.19E-06 |
| ZNFX1 | SNs | 5.493236564 | 3.21E-06 |
| SELENON | SNs | 2.717872488 | 0.00191482 |
| SLC45A4 | SNs | 2.694665723 | 0.00201992 |
| TTC7A | SNs | 2.236531624 | 0.00580054 |
| ERGIC1 | SNs | 3.662504779 | 0.00021752 |
| PBXIP1 | SNs | 2.684113904 | 0.0020696 |
| RCN3 | SNs | 4.441188256 | 3.62E-05 |
| SCYL1 | SNs | 5.497174226 | 3.18E-06 |
| RHBDD2 | SNs | 3.773561031 | 0.00016844 |
| SLC24A3 | SNs | 3.458320802 | 0.00034808 |
| ELAPOR1 | SNs | 3.025789449 | 0.00094235 |
| PREX1 | SNs | 3.269845464 | 0.00053722 |
| MRTFA | SNs | 4.020390715 | 9.54E-05 |
| ZNF687 | SNs | 5.684387896 | 2.07E-06 |
| VPS18 | SNs | 3.965389812 | 0.0001083 |
| LRFN1 | SNs | 3.46128352 | 0.00034571 |
| PHF12 | SNs | 3.29044352 | 0.00051234 |
| GRAMD1A | SNs | 4.162303572 | 6.88E-05 |
| CALCOCO1 | SNs | 3.971127436 | 0.00010687 |
| FBRSL1 | SNs | 2.138830025 | 0.0072639 |
| NCKAP5L | SNs | 2.484265765 | 0.00327895 |
| GBA2 | SNs | 2.908264615 | 0.00123519 |
| WDFY4 | SNs | 2.55839284 | 0.00276444 |
| DENND1A | SNs | 2.974373197 | 0.00106078 |
| RAB40C | SNs | 3.080118432 | 0.00083154 |
| NFE4 | SNs | 2.575089685 | 0.00266018 |
| CTDSP1 | SNs | 4.830332961 | 1.48E-05 |
| CXCL16 | SNs | 4.468785411 | 3.40E-05 |
| TP53INP2 | SNs | 2.731970236 | 0.00185366 |
| TRAPPC1 | SNs | 2.916849226 | 0.00121102 |
| SCAF1 | SNs | 2.88824025 | 0.00129348 |
| EPS15L1 | SNs | 5.378042702 | 4.19E-06 |
| PGAP6 | SNs | 3.544098359 | 0.00028569 |
| PPCDC | SNs | 2.86597869 | 0.00136151 |
| RIC8A | SNs | 2.829859237 | 0.00147959 |
| MIIP | SNs | 3.091235385 | 0.00081052 |
| ZFAND3 | SNs | 5.134768052 | 7.33E-06 |
| ABHD4 | SNs | 5.2132231 | 6.12E-06 |
| PCIF1 | SNs | 2.865272929 | 0.00136373 |
| GPSM3 | SNs | 4.570133796 | 2.69E-05 |
| PARVG | SNs | 5.135904896 | 7.31E-06 |
| LRRC4 | SNs | 2.392065301 | 0.00405448 |
| TMBIM1 | SNs | 2.486455662 | 0.00326245 |
| VSIR | SNs | 6.338104484 | 4.59E-07 |
| DPEP2 | SNs | 4.299564157 | 5.02E-05 |

| | | | |
|---|---|---|---|
| DPEP3 | SNs | 3.873023858 | 0.00013396 |
| SEMA4A | SNs | 4.990177986 | 1.02E-05 |
| PDLIM2 | SNs | 2.616044451 | 0.00242078 |
| FBRS | SNs | 4.821482672 | 1.51E-05 |
| ARHGAP9 | SNs | 3.428609028 | 0.00037273 |
| HS1BP3 | SNs | 4.19861999 | 6.33E-05 |
| SIL1 | SNs | 3.056536342 | 0.00087794 |
| MMP25 | SNs | 3.455511092 | 0.00035034 |
| ARAP3 | SNs | 3.787954338 | 0.00016295 |
| MTMR14 | SNs | 3.668397191 | 0.00021459 |
| MOSPD3 | SNs | 2.798581764 | 0.00159008 |
| GIGYF1 | SNs | 2.165004964 | 0.00683904 |
| CSRNP1 | SNs | 5.001072965 | 9.98E-06 |
| GORASP1 | SNs | 2.80659545 | 0.00156101 |
| SMAP2 | SNs | 3.598627696 | 0.00025198 |
| PLPPR2 | SNs | 4.302455791 | 4.98E-05 |
| MICAL1 | SNs | 2.871458144 | 0.00134444 |
| ELOVL1 | SNs | 2.391342507 | 0.00406123 |
| ISL2 | SNs | 3.039118397 | 0.00091386 |
| MARCKSL1 | SNs | 3.041987919 | 0.00090785 |
| INTS3 | SNs | 2.522444587 | 0.003003 |
| NADK | SNs | 4.944977779 | 1.14E-05 |
| YIPF2 | SNs | 2.129645863 | 0.00741915 |
| PRR14 | SNs | 5.940482351 | 1.15E-06 |
| CUEDC2 | SNs | 2.241429014 | 0.0057355 |
| KXD1 | SNs | 2.235940784 | 0.00580844 |
| TSEN34 | SNs | 5.509868075 | 3.09E-06 |
| ATG9A | SNs | 2.980735287 | 0.00104536 |
| TMUB2 | SNs | 5.558869103 | 2.76E-06 |
| BBLN | SNs | 2.576814242 | 0.00264963 |
| PHF23 | SNs | 2.875258042 | 0.00133273 |
| MBOAT7 | SNs | 6.602617195 | 2.50E-07 |
| LILRA6 | SNs | 3.18568445 | 0.0006521 |
| RBM42 | SNs | 2.500308028 | 0.00316004 |
| EFHD2 | SNs | 5.399826312 | 3.98E-06 |
| CYBC1 | SNs | 4.424118174 | 3.77E-05 |
| WDR25 | SNs | 2.196436535 | 0.00636156 |
| RIPOR1 | SNs | 3.251230305 | 0.00056075 |
| ABHD8 | SNs | 1.993864209 | 0.01014228 |
| CORO7 | SNs | 4.238114113 | 5.78E-05 |
| CERS4 | SNs | 3.073340182 | 0.00084462 |
| CCNJL | SNs | 4.306000181 | 4.94E-05 |
| GALNT14 | SNs | 2.798182564 | 0.00159154 |
| ARMC7 | SNs | 2.731358221 | 0.00185627 |
| USB1 | SNs | 5.174910225 | 6.68E-06 |
| RHBDF2 | SNs | 2.275730913 | 0.00529992 |
| NLRX1 | SNs | 6.828936173 | 1.48E-07 |
| LRRK1 | SNs | 4.991958669 | 1.02E-05 |
| COLGALT1 | SNs | 4.895065246 | 1.27E-05 |
| IGFLR1 | SNs | 2.616425461 | 0.00241866 |
| NPEPL1 | SNs | 3.766077552 | 0.00017137 |
| VPS37B | SNs | 3.948572531 | 0.00011257 |
| ZNF768 | SNs | 3.31029313 | 0.00048945 |
| NSUN7 | SNs | 3.648477516 | 0.00022466 |
| TBC1D17 | SNs | 2.272187491 | 0.00534334 |
| DHRS12 | SNs | 2.124418067 | 0.007509 |
| MICALL2 | SNs | 2.26920494 | 0.00538016 |
| TREML2 | SNs | 3.174118024 | 0.0006697 |
| RIN3 | SNs | 4.505665409 | 3.12E-05 |
| NAA60 | SNs | 2.733130506 | 0.00184871 |
| DOK3 | SNs | 6.509771802 | 3.09E-07 |
| KIAA0319L | SNs | 4.684838766 | 2.07E-05 |
| COQ8B | SNs | 3.81205535 | 0.00015415 |
| SLC35E1 | SNs | 2.854070215 | 0.00139936 |
| PAQR6 | SNs | 2.014615454 | 0.00966907 |
| DGLUCY | SNs | 2.735048489 | 0.00184057 |
| UBTD1 | SNs | 4.8996628 | 1.26E-05 |
| MYO15B | SNs | 2.814161126 | 0.00153405 |
| SLC8B1 | SNs | 2.756866282 | 0.00175039 |
| SLC66A2 | SNs | 2.112526662 | 0.00771744 |
| PGGHG | SNs | 2.628746766 | 0.002351 |

| | | | |
|---|---|---|---|
| FUZ | SNs | 2.095229503 | 0.00803102 |
| ORAI2 | SNs | 2.320329168 | 0.00478267 |
| RUFY1 | SNs | 2.085350161 | 0.0082158 |
| ATOSB | SNs | 4.564618493 | 2.73E-05 |
| PLEKHO2 | SNs | 6.748012245 | 1.79E-07 |
| ABTB1 | SNs | 5.275375509 | 5.30E-06 |
| DNAJC5 | SNs | 4.459837775 | 3.47E-05 |
| REEP4 | SNs | 2.179036852 | 0.0066216 |
| AKNA | SNs | 3.060507604 | 0.00086995 |
| MPIG6B | SNs | 4.100508776 | 7.93E-05 |
| LY6G5C | SNs | 2.498235781 | 0.00317515 |
| CPTP | SNs | 2.394988262 | 0.00402728 |
| LIMD2 | SNs | 2.262486685 | 0.00546403 |
| B9D2 | SNs | 3.962829089 | 0.00010894 |
| CMIP | SNs | 3.346838336 | 0.00044995 |
| EEPD1 | SNs | 3.232558613 | 0.00058538 |
| SH3BP5L | SNs | 5.712149825 | 1.94E-06 |
| ZBP1 | SNs | 2.241892895 | 0.00572937 |
| HM13 | SNs | 3.999128687 | 0.0001002 |
| TRIM11 | SNs | 2.686253979 | 0.00205943 |
| NDEL1 | SNs | 3.479019415 | 0.00033188 |
| TRIM8 | SNs | 5.575597866 | 2.66E-06 |
| TSC22D4 | SNs | 4.048669219 | 8.94E-05 |
| NUAK2 | SNs | 6.853359409 | 1.40E-07 |
| MED25 | SNs | 5.377499602 | 4.19E-06 |
| ISG20L2 | SNs | 2.097000993 | 0.00799832 |
| RAB1B | SNs | 5.45073495 | 3.54E-06 |
| SH3BGRL3 | SNs | 3.890010711 | 0.00012882 |
| FAM110A | SNs | 3.342073507 | 0.00045491 |
| RASSF5 | SNs | 2.69654947 | 0.00201118 |
| BCL2L12 | SNs | 2.257014275 | 0.00553332 |
| CCM2 | SNs | 2.603123618 | 0.00249388 |
| C11orf68 | SNs | 4.605002522 | 2.48E-05 |
| DYNLRB1 | SNs | 2.19751212 | 0.00634582 |
| SESN2 | SNs | 2.630294787 | 0.00234264 |
| FERMT3 | SNs | 4.602232016 | 2.50E-05 |
| YPEL3 | SNs | 4.513451719 | 3.07E-05 |
| FRMD8 | SNs | 2.192238509 | 0.00642335 |
| KLF16 | SNs | 3.052972596 | 0.00088517 |
| TMEM120A | SNs | 5.340056849 | 4.57E-06 |
| RAB34 | SNs | 3.142373019 | 0.00072049 |
| STK40 | SNs | 5.744000543 | 1.80E-06 |
| SPNS1 | SNs | 2.00550955 | 0.00987394 |
| FAM234A | SNs | 2.009435602 | 0.00978508 |
| EMILIN2 | SNs | 3.041866858 | 0.0009081 |
| TMEM222 | SNs | 3.156910654 | 0.00069677 |
| FHIP1B | SNs | 2.983953571 | 0.00103764 |
| PRAM1 | SNs | 4.915562469 | 1.21E-05 |
| KAT8 | SNs | 3.157207275 | 0.00069629 |
| DRC7 | SNs | 2.260848578 | 0.00548468 |
| TRAF7 | SNs | 3.590038888 | 0.00025702 |
| ZDHHC18 | SNs | 3.882364794 | 0.00013111 |
| CAMKK1 | SNs | 2.694154192 | 0.0020223 |
| CARD19 | SNs | 3.207661068 | 0.00061992 |
| POLDIP3 | SNs | 2.611153751 | 0.0024482 |
| NUDT22 | SNs | 2.163693361 | 0.00685972 |
| VPS25 | SNs | 2.339145953 | 0.00457988 |
| ARFGAP2 | SNs | 2.733883286 | 0.00184551 |
| CYSTM1 | SNs | 2.805587201 | 0.00156463 |
| RAB11FIP4 | SNs | 2.377908493 | 0.00418882 |
| ACRBP | SNs | 3.732174716 | 0.00018528 |
| MAP1LC3A | SNs | 2.56069663 | 0.00274981 |
| NTNG2 | SNs | 2.870182477 | 0.0013484 |
| TNRC18 | SNs | 2.463856944 | 0.00343671 |
| DGAT2 | SNs | 6.037579999 | 9.17E-07 |
| GLYR1 | SNs | 2.399618614 | 0.00398457 |
| GLIS2 | SNs | 3.227051902 | 0.00059285 |
| PPP1R9B | SNs | 5.386676593 | 4.11E-06 |
| HDGFL2 | SNs | 2.096701628 | 0.00800384 |
| PSRC1 | SNs | 3.820194202 | 0.00015129 |
| TUBA1C | SNs | 5.209905375 | 6.17E-06 |

| | | | |
|---|---|---|---|
| NFKBID | SNs | 3.364461661 | 0.00043205 |
| ORAI1 | SNs | 2.440669513 | 0.00362519 |
| ZDHHC12 | SNs | 3.535673565 | 0.00029129 |
| ZNF341 | SNs | 2.308297244 | 0.00491703 |
| SPRYD3 | SNs | 3.409063928 | 0.00038988 |
| HSH2D | SNs | 4.525839358 | 2.98E-05 |
| RELT | SNs | 2.490417352 | 0.00323283 |
| UBL7 | SNs | 2.020000379 | 0.00954992 |
| SMIM3 | SNs | 2.162802822 | 0.0068738 |
| H2BC12 | SNs | 2.428837943 | 0.00372531 |
| LGALS12 | SNs | 3.751113374 | 0.00017737 |
| STON2 | SNs | 2.326920862 | 0.00471063 |
| ITPRIP | SNs | 3.636831117 | 0.00023076 |
| FGD3 | SNs | 4.397846115 | 4.00E-05 |
| FCHSD1 | SNs | 2.849079056 | 0.00141554 |
| ATG16L2 | SNs | 4.345899355 | 4.51E-05 |
| AQP10 | SNs | 3.195915137 | 0.00063692 |
| ZNF628 | SNs | 1.990361564 | 0.01022441 |
| MIDN | SNs | 5.079792043 | 8.32E-06 |
| TMEM250 | SNs | 2.403193998 | 0.0039519 |
| KIAA2013 | SNs | 3.768246157 | 0.00017051 |
| BMF | SNs | 2.479326241 | 0.00331645 |
| YIF1B | SNs | 2.945652499 | 0.00113331 |
| LRSAM1 | SNs | 2.546968448 | 0.00283813 |
| CFAP119 | SNs | 4.128537876 | 7.44E-05 |
| SPOCD1 | SNs | 3.385473013 | 0.00041165 |
| ESAM | SNs | 4.876331853 | 1.33E-05 |
| AP5B1 | SNs | 5.570711922 | 2.69E-06 |
| R3HDM4 | SNs | 4.176116406 | 6.67E-05 |
| RNF185 | SNs | 2.731090409 | 0.00185742 |
| UBXN11 | SNs | 2.629394119 | 0.0023475 |
| NLRP12 | SNs | 6.423240922 | 3.77E-07 |
| MYADM | SNs | 3.367162724 | 0.00042938 |
| TMEM88 | SNs | 4.521941545 | 3.01E-05 |
| STRADA | SNs | 3.81266023 | 0.00015394 |
| G6PC3 | SNs | 3.655916528 | 0.00022084 |
| ARRDC1 | SNs | 2.741651757 | 0.00181279 |
| SHKBP1 | SNs | 5.329923495 | 4.68E-06 |
| ZBTB47 | SNs | 2.95065038 | 0.00112034 |
| ORAI3 | SNs | 3.806688838 | 0.00015607 |
| MVB12A | SNs | 2.303787118 | 0.00496836 |
| PIGS | SNs | 2.045989668 | 0.00899519 |
| SFXN5 | SNs | 3.971240534 | 0.00010685 |
| SYTL3 | SNs | 2.046181598 | 0.00899122 |
| EPSTI1 | SNs | 2.124509027 | 0.00750742 |
| CMTM7 | SNs | 3.282928365 | 0.00052128 |
| NACC1 | SNs | 3.019363201 | 0.00095639 |
| RPUSD1 | SNs | 2.054339061 | 0.00882391 |
| SCAMP4 | SNs | 3.020121632 | 0.00095473 |
| MISP3 | SNs | 2.13954425 | 0.00725197 |
| DTX2 | SNs | 3.741142801 | 0.00018149 |
| NLRP3 | SNs | 2.298777637 | 0.005026 |
| MBD6 | SNs | 5.660013327 | 2.19E-06 |
| STK11IP | SNs | 2.33098892 | 0.00466671 |
| RASGRP4 | SNs | 7.480826347 | 3.31E-08 |
| DHRS1 | SNs | 2.169350454 | 0.00677095 |
| BATF2 | SNs | 2.312287726 | 0.00487206 |
| DNTTIP1 | SNs | 3.226576622 | 0.0005935 |
| KLHDC3 | SNs | 2.761832512 | 0.00173048 |
| CMTM5 | SNs | 3.483326928 | 0.0003286 |
| SLC26A8 | SNs | 2.84811008 | 0.0014187 |
| LRG1 | SNs | 5.68817449 | 2.05E-06 |
| ARAP1 | SNs | 4.917241282 | 1.21E-05 |
| JAML | SNs | 2.474684093 | 0.00335209 |
| TDRD9 | SNs | 2.495234538 | 0.00319717 |
| JDP2 | SNs | 2.023002503 | 0.00948413 |
| SPATA2L | SNs | 3.482923966 | 0.00032891 |
| ZC3H18 | SNs | 2.306683799 | 0.00493533 |
| UBALD1 | SNs | 2.22532389 | 0.00595218 |
| SNX20 | SNs | 2.577385306 | 0.00264615 |
| CANT1 | SNs | 4.327945879 | 4.70E-05 |

| | | | |
|---|---|---|---|
| CD300LB | SNs | 4.137335721 | 7.29E-05 |
| HEXIM2 | SNs | 3.46969253 | 0.00033908 |
| SLC43A2 | SNs | 6.164512776 | 6.85E-07 |
| SPNS2 | SNs | 2.12913557 | 0.00742787 |
| LOXHD1 | SNs | 2.292685327 | 0.005097 |
| RAVER1 | SNs | 4.242785494 | 5.72E-05 |
| OSCAR | SNs | 4.896373949 | 1.27E-05 |
| CCDC159 | SNs | 3.175756962 | 0.00066718 |
| ZNF787 | SNs | 2.034036249 | 0.00924621 |
| MOB3A | SNs | 5.420310044 | 3.80E-06 |
| LRRC25 | SNs | 5.168586809 | 6.78E-06 |
| TPRG1L | SNs | 3.392987633 | 0.00040459 |
| RNF19B | SNs | 4.286669238 | 5.17E-05 |
| ARL8A | SNs | 7.133167583 | 7.36E-08 |
| CHMP4B | SNs | 3.006318281 | 0.00098556 |
| TANGO2 | SNs | 3.239592466 | 0.00057598 |
| CMPK2 | SNs | 2.250228821 | 0.00562045 |
| ZFAND2B | SNs | 2.457759068 | 0.00348531 |
| ZDHHC19 | SNs | 2.431650222 | 0.00370126 |
| PPM1M | SNs | 3.753207657 | 0.00017652 |
| SIRPA | SNs | 4.94945163 | 1.12E-05 |
| IFITM3P2 | SNs | 2.250206785 | 0.00562074 |
| GLT1D1 | SNs | 4.280798188 | 5.24E-05 |
| TTC7B | SNs | 3.228557599 | 0.0005908 |
| B3GNTL1 | SNs | 3.959406713 | 0.0001098 |
| CD300LF | SNs | 3.551653982 | 0.00028077 |
| PIK3R6 | SNs | 2.266378394 | 0.00541529 |
| DHRS13 | SNs | 4.032896 | 9.27E-05 |
| CCDC17 | SNs | 2.487528892 | 0.0032544 |
| NFAM1 | SNs | 6.597908717 | 2.52E-07 |
| PLB1 | SNs | 4.675531801 | 2.11E-05 |
| GPBAR1 | SNs | 2.052249006 | 0.00886648 |
| ASPRV1 | SNs | 2.169160762 | 0.00677391 |
| GLIPR2 | SNs | 6.377322859 | 4.19E-07 |
| CIDECP1 | SNs | 2.353549638 | 0.00443048 |
| ZNF746 | SNs | 3.865744813 | 0.00013622 |
| PHOSPHO1 | SNs | 2.24515412 | 0.00568651 |
| DEDD2 | SNs | 5.663892651 | 2.17E-06 |
| GBP6 | SNs | 2.116798105 | 0.00764191 |
| OXER1 | SNs | 3.525025192 | 0.00029852 |
| ZNF467 | SNs | 3.752441526 | 0.00017683 |
| PWWP2B | SNs | 2.057844761 | 0.00875297 |
| PPP1R18 | SNs | 5.441520869 | 3.62E-06 |
| NLRP6 | SNs | 3.697810326 | 0.00020053 |
| CYP2T1P | SNs | 5.18911391 | 6.47E-06 |
| RILPL2 | SNs | 2.164324918 | 0.00684976 |
| ADCY4 | SNs | 2.111030131 | 0.00774408 |
| FAAP20 | SNs | 2.253591311 | 0.0055771 |
| SLC5A9 | SNs | 3.128255079 | 0.00074429 |
| CRTC2 | SNs | 6.484339855 | 3.28E-07 |
| LINC00528 | SNs | 2.344781838 | 0.00452083 |
| APOBEC3A | SNs | 3.239536025 | 0.00057606 |
| KLHDC8B | SNs | 3.336828126 | 0.00046044 |
| ARHGAP27 | SNs | 3.089802143 | 0.0008132 |
| SAMD14 | SNs | 2.185438754 | 0.00652471 |
| UNC13D | SNs | 4.540725523 | 2.88E-05 |
| TPCN2 | SNs | 2.890660191 | 0.00128629 |
| SAXO4 | SNs | 2.020859813 | 0.00953104 |
| OAF | SNs | 4.00352316 | 9.92E-05 |
| TIGD3 | SNs | 3.147702617 | 0.0007117 |
| CPNE2 | SNs | 4.0309116 | 9.31E-05 |
| C6orf89 | SNs | 2.811332336 | 0.00154407 |
| SMIM29 | SNs | 2.180783645 | 0.00659502 |
| ADGRG3 | SNs | 3.588880946 | 0.0002577 |
| PELI3 | SNs | 2.512817009 | 0.00307032 |
| KCTD13 | SNs | 3.478824046 | 0.00033203 |
| EHBP1L1 | SNs | 2.807117658 | 0.00155913 |
| INAFM1 | SNs | 2.783062699 | 0.00164792 |
| C19orf38 | SNs | 7.09711005 | 8.00E-08 |
| NATD1 | SNs | 3.788200406 | 0.00016285 |
| ARHGAP30 | SNs | 5.007884326 | 9.82E-06 |

| | | | |
|---|---|---|---|
| LY6G6F | SNs | 3.016767906 | 0.00096213 |
| PRSS33 | SNs | 2.038456324 | 0.00915258 |
| PDCD4-AS1 | SNs | 2.272501436 | 0.00533948 |
| KCTD21 | SNs | 2.024496631 | 0.00945156 |
| CCDC88B | SNs | 2.79453505 | 0.00160496 |
| C16orf54 | SNs | 2.951698092 | 0.00111764 |
| UBALD2 | SNs | 4.311850378 | 4.88E-05 |
| TMEM150B | SNs | 2.448756036 | 0.00355831 |
| MIR23AHG | SNs | 2.17476544 | 0.00668705 |
| THEM5 | SNs | 2.499418957 | 0.00316651 |
| SIRPB2 | SNs | 3.017719804 | 0.00096002 |
| NT5DC4 | SNs | 2.696881711 | 0.00200964 |
| RAB37 | SNs | 3.517044825 | 0.00030406 |
| ANKRD13D | SNs | 4.338176351 | 4.59E-05 |
| RAB43 | SNs | 2.960332099 | 0.00109564 |
| CLEC4G | SNs | 2.486882362 | 0.00325925 |
| TREML1 | SNs | 2.731680431 | 0.0018549 |
| TSPAN33 | SNs | 4.380534353 | 4.16E-05 |
| GARIN1A | SNs | 2.482910894 | 0.00328919 |
| RILPL1 | SNs | 2.192774605 | 0.00641542 |
| LILRA5 | SNs | 3.715208494 | 0.00019266 |
| TMEM179B | SNs | 2.202012833 | 0.0062804 |
| ZNF710 | SNs | 4.75585773 | 1.75E-05 |
| PEAK3 | SNs | 4.16519937 | 6.84E-05 |
| TMEM205 | SNs | 2.274893968 | 0.00531014 |
| B3GNT8 | SNs | 5.163820343 | 6.86E-06 |
| PEAR1 | SNs | 2.921360121 | 0.00119851 |
| SLC27A1 | SNs | 3.442518401 | 0.00036098 |
| TPTEP1 | SNs | 2.456211517 | 0.00349775 |
| LAMTOR4 | SNs | 2.102922949 | 0.00789 |
| LINC01002 | SNs | 4.778589177 | 1.66E-05 |
| GSEC | SNs | 3.726738317 | 0.00018761 |
| FAM174B | SNs | 2.55382481 | 0.00279367 |
| LINC01061 | SNs | 3.233911793 | 0.00058356 |
| LINC02908 | SNs | 3.635276208 | 0.00023159 |
| LINC01000 | SNs | 2.874794177 | 0.00133415 |
| IFITM10 | SNs | 3.098474044 | 0.00079712 |
| CUEDC1 | SNs | 3.257178937 | 0.00055312 |
| RAB43P1 | SNs | 2.255197896 | 0.00555651 |
| RNASEK | SNs | 3.5600432 | 0.0002754 |
| PLIN5 | SNs | 2.696359618 | 0.00201206 |
| NUDT4B | SNs | 2.351419481 | 0.00445226 |
| C8orf58 | SNs | 3.409144139 | 0.00038981 |
| PEF1 | SNs | 2.815203537 | 0.00153037 |
| TMEM91 | SNs | 3.593807076 | 0.0002548 |
| MROH6 | SNs | 2.126564768 | 0.00747197 |
| PGA4 | SNs | 4.411417925 | 3.88E-05 |
| PIK3CD-AS1 | SNs | 3.183994103 | 0.00065465 |
| NCF1 | SNs | 3.995975221 | 0.00010093 |
| FAM157A | SNs | 3.526754615 | 0.00029733 |
| CDK11A | SNs | 2.157794184 | 0.00695354 |
| PLIN4 | SNs | 3.465334005 | 0.0003425 |
| CASTOR2 | SNs | 2.079375831 | 0.0083296 |
| BORCS8 | SNs | 3.495972607 | 0.00031917 |
| SIGLEC14 | SNs | 2.142296809 | 0.00720615 |
| LINC01001 | SNs | 2.296184889 | 0.00505609 |
| PPM1F-AS1 | SNs | 2.450568829 | 0.00354349 |
| LINC00173 | SNs | 3.560397719 | 0.00027517 |
| DDX11L9 | SNs | 2.121788325 | 0.0075546 |
| SAP25 | SNs | 2.484764333 | 0.00327518 |
| PELATON | SNs | 6.227237838 | 5.93E-07 |
| LINC01503 | SNs | 3.688646136 | 0.00020481 |
| LINC00963 | SNs | 4.130328347 | 7.41E-05 |
| PTPRN2-AS1 | SNs | 3.087627098 | 0.00081728 |
| ZNF865 | SNs | 3.211653366 | 0.00061425 |
| SENCR | SNs | 5.437687923 | 3.65E-06 |
| ARPC4-TTLL3 | SNs | 2.046142788 | 0.00899202 |
| MMP24OS | SNs | 2.607743782 | 0.00246749 |
| C5orf67 | SNs | 2.706466296 | 0.00196577 |
| C3orf86P | SNs | 4.759977707 | 1.74E-05 |
| LINC02887 | SNs | 2.419666779 | 0.00380481 |

| | | | |
|---|---|---|---|
| LINC02723 | SNs | 2.502772942 | 0.00314215 |
| CORO1A-AS1 | SNs | 2.003499833 | 0.00991974 |

**Suppl Table 8.** Enriched pathway analysis for each ALS subtype based on multi-class LDA prediction in blood.

| Reference ALS-Subtype | Database | Term ID | Enriched Pathway | Gene Ratio | Adj P value | Count |
|---|---|---|---|---|---|---|
| ALS-SNs | BP | GO:0006909 | BP phagocytosis | 0.046 | 1.10E-09 | 57 |
| ALS-SNs | BP | GO:0007015 | BP actin filament organization | 0.063 | 2.73E-08 | 78 |
| ALS-SNs | BP | GO:0002275 | BP myeloid cell activation involved in immune response | 0.024 | 1.76E-07 | 30 |
| ALS-SNs | BP | GO:0050900 | BP leukocyte migration | 0.054 | 7.95E-07 | 67 |
| ALS-SNs | BP | GO:0002274 | BP myeloid leukocyte activation | 0.040 | 7.95E-07 | 50 |
| ALS-SNs | BP | GO:0060627 | BP regulation of vesicle-mediated transport | 0.066 | 7.95E-07 | 82 |
| ALS-SNs | BP | GO:0071706 | BP tumor necrosis factor superfamily cytokine production | 0.032 | 3.06E-06 | 40 |
| ALS-SNs | BP | GO:1903555 | BP regulation of tumor necrosis factor superfamily cytokine production | 0.032 | 3.06E-06 | 40 |
| ALS-SNs | BP | GO:0006935 | BP chemotaxis | 0.052 | 4.21E-06 | 65 |
| ALS-SNs | BP | GO:0007033 | BP vacuole organization | 0.041 | 4.21E-06 | 51 |
| ALS-SNs | BP | GO:0045088 | BP regulation of innate immune response | 0.061 | 9.51E-06 | 76 |
| ALS-SNs | BP | GO:0016032 | BP viral process | 0.059 | 1.49E-05 | 74 |
| ALS-SNs | BP | GO:0019221 | BP cytokine-mediated signaling pathway | 0.058 | 1.96E-05 | 72 |
| ALS-SNs | BP | GO:0009611 | BP response to wounding | 0.060 | 2.75E-05 | 75 |
| ALS-SNs | BP | GO:0032103 | BP positive regulation of response to external stimulus | 0.072 | 2.75E-05 | 90 |
| ALS-SNs | BP | GO:0006898 | BP receptor-mediated endocytosis | 0.038 | 3.00E-05 | 47 |
| ALS-SNs | BP | GO:0050727 | BP regulation of inflammatory response | 0.048 | 3.21E-05 | 60 |
| ALS-SNs | BP | GO:0008360 | BP regulation of cell shape | 0.025 | 3.51E-05 | 31 |
| ALS-SNs | BP | GO:0032418 | BP lysosome localization | 0.018 | 4.55E-05 | 23 |
| ALS-SNs | BP | GO:0007186 | BP G protein-coupled receptor signaling pathway | 0.063 | 4.61E-05 | 79 |
| ALS-SNs | BP | GO:0050878 | BP regulation of body fluid levels | 0.042 | 4.61E-05 | 53 |
| ALS-SNs | BP | GO:1990849 | BP vacuolar localization | 0.018 | 5.38E-05 | 23 |
| ALS-SNs | BP | GO:0002764 | BP immune response-regulating signaling pathway | 0.071 | 5.47E-05 | 89 |
| ALS-SNs | BP | GO:0032102 | BP negative regulation of response to external stimulus | 0.047 | 6.40E-05 | 59 |
| ALS-SNs | BP | GO:0043254 | BP regulation of protein-containing complex assembly | 0.054 | 7.01E-05 | 67 |
| ALS-SNs | BP | GO:0030595 | BP leukocyte chemotaxis | 0.032 | 0.0001 | 40 |
| ALS-SNs | BP | GO:0002697 | BP regulation of immune effector process | 0.050 | 0.0002 | 63 |
| ALS-SNs | BP | GO:0002831 | BP regulation of response to biotic stimulus | 0.065 | 0.0002 | 81 |
| ALS-SNs | BP | GO:0043299 | BP leukocyte degranulation | 0.017 | 0.0002 | 21 |
| ALS-SNs | BP | GO:0042554 | BP superoxide anion generation | 0.010 | 0.0003 | 13 |
| ALS-SNs | BP | GO:0032970 | BP regulation of actin filament-based process | 0.043 | 0.0004 | 54 |
| ALS-SNs | BP | GO:0045453 | BP bone resorption | 0.014 | 0.0004 | 17 |
| ALS-SNs | BP | GO:0043085 | BP positive regulation of catalytic activity | 0.067 | 0.0004 | 84 |
| ALS-SNs | BP | GO:0002683 | BP negative regulation of immune system process | 0.058 | 0.0005 | 73 |
| ALS-SNs | BP | GO:0046718 | BP symbiont entry into host cell | 0.024 | 0.0006 | 30 |
| ALS-SNs | BP | GO:0031623 | BP receptor internalization | 0.021 | 0.0007 | 26 |
| ALS-SNs | BP | GO:0051258 | BP protein polymerization | 0.038 | 0.0008 | 47 |
| ALS-SNs | BP | GO:0045785 | BP positive regulation of cell adhesion | 0.053 | 0.0008 | 66 |
| ALS-SNs | BP | GO:0009617 | BP response to bacterium | 0.065 | 0.0008 | 81 |
| ALS-SNs | BP | GO:0032956 | BP regulation of actin cytoskeleton organization | 0.039 | 0.0009 | 49 |
| ALS-SNs | BP | GO:0007596 | BP blood coagulation | 0.029 | 0.0009 | 36 |
| ALS-SNs | BP | GO:0051336 | BP regulation of hydrolase activity | 0.058 | 0.0010 | 72 |
| ALS-SNs | BP | GO:0002444 | BP myeloid leukocyte mediated immunity | 0.020 | 0.0010 | 25 |
| ALS-SNs | BP | GO:0016236 | BP macroautophagy | 0.048 | 0.0011 | 60 |
| ALS-SNs | BP | GO:0050817 | BP coagulation | 0.029 | 0.0012 | 36 |
| ALS-SNs | BP | GO:0052547 | BP regulation of peptidase activity | 0.034 | 0.0012 | 43 |
| ALS-SNs | BP | GO:1903131 | BP mononuclear cell differentiation | 0.059 | 0.0013 | 74 |
| ALS-SNs | BP | GO:0030833 | BP regulation of actin filament polymerization | 0.021 | 0.0014 | 26 |
| ALS-SNs | BP | GO:0002685 | BP regulation of leukocyte migration | 0.030 | 0.0014 | 37 |
| ALS-SNs | BP | GO:0050777 | BP negative regulation of immune response | 0.029 | 0.0014 | 36 |
| ALS-SNs | BP | GO:0051452 | BP intracellular pH reduction | 0.012 | 0.0016 | 15 |

| | | | | | | |
|---|---|---|---|---|---|---|
| ALS-SNs | BP | GO:0043312 | BP neutrophil degranulation | 0.006 | 0.0018 | 7 |
| ALS-SNs | BP | GO:0032535 | BP regulation of cellular component size | 0.039 | 0.0018 | 49 |
| ALS-SNs | BP | GO:0045862 | BP positive regulation of proteolysis | 0.039 | 0.0018 | 49 |
| ALS-SNs | BP | GO:0043304 | BP regulation of mast cell degranulation | 0.008 | 0.0018 | 10 |
| ALS-SNs | BP | GO:0016050 | BP vesicle organization | 0.046 | 0.0018 | 57 |
| ALS-SNs | BP | GO:0031329 | BP regulation of cellular catabolic process | 0.053 | 0.0021 | 66 |
| ALS-SNs | BP | GO:0010876 | BP lipid localization | 0.046 | 0.0023 | 57 |
| ALS-SNs | BP | GO:0002532 | BP production of molecular mediator involved in inflammatory response | 0.015 | 0.0025 | 19 |
| ALS-SNs | BP | GO:0042110 | BP T cell activation | 0.064 | 0.0027 | 80 |
| ALS-SNs | BP | GO:0007159 | BP leukocyte cell-cell adhesion | 0.048 | 0.0028 | 60 |
| ALS-SNs | BP | GO:0040017 | BP positive regulation of locomotion | 0.054 | 0.0032 | 67 |
| ALS-SNs | BP | GO:0010256 | BP endomembrane system organization | 0.064 | 0.0033 | 80 |
| ALS-SNs | BP | GO:0022411 | BP cellular component disassembly | 0.050 | 0.0034 | 63 |
| ALS-SNs | BP | GO:0046849 | BP bone remodeling | 0.014 | 0.0035 | 17 |
| ALS-SNs | BP | GO:0016052 | BP carbohydrate catabolic process | 0.024 | 0.0035 | 30 |
| ALS-SNs | BP | GO:0043542 | BP endothelial cell migration | 0.026 | 0.0036 | 32 |
| ALS-SNs | BP | GO:0001932 | BP regulation of protein phosphorylation | 0.064 | 0.0036 | 80 |
| ALS-SNs | BP | GO:0010324 | BP membrane invagination | 0.013 | 0.0036 | 16 |
| ALS-SNs | BP | GO:0141084 | BP inflammasome-mediated signaling pathway | 0.010 | 0.0036 | 12 |
| ALS-SNs | BP | GO:0036344 | BP platelet morphogenesis | 0.008 | 0.0036 | 10 |
| ALS-SNs | BP | GO:1902903 | BP regulation of supramolecular fiber organization | 0.042 | 0.0037 | 53 |
| ALS-SNs | BP | GO:0051051 | BP negative regulation of transport | 0.041 | 0.0037 | 51 |
| ALS-SNs | BP | GO:0060249 | BP anatomical structure homeostasis | 0.027 | 0.0041 | 34 |
| ALS-SNs | BP | GO:0090066 | BP regulation of anatomical structure size | 0.046 | 0.0042 | 57 |
| ALS-SNs | BP | GO:0050867 | BP positive regulation of cell activation | 0.044 | 0.0047 | 55 |
| ALS-SNs | BP | GO:0031589 | BP cell-substrate adhesion | 0.035 | 0.0050 | 44 |
| ALS-SNs | BP | GO:0140632 | BP canonical inflammasome complex assembly | 0.009 | 0.0053 | 11 |
| ALS-SNs | BP | GO:0032930 | BP positive regulation of superoxide anion generation | 0.006 | 0.0053 | 8 |
| ALS-SNs | BP | GO:1902107 | BP positive regulation of leukocyte differentiation | 0.026 | 0.0053 | 32 |
| ALS-SNs | BP | GO:0071222 | BP cellular response to lipopolysaccharide | 0.027 | 0.0057 | 34 |
| ALS-SNs | BP | GO:0007034 | BP vacuolar transport | 0.025 | 0.0062 | 31 |
| ALS-SNs | BP | GO:0097300 | BP programmed necrotic cell death | 0.011 | 0.0066 | 14 |
| ALS-SNs | BP | GO:0034341 | BP response to type II interferon | 0.018 | 0.0068 | 23 |
| ALS-SNs | BP | GO:0033280 | BP response to vitamin D | 0.007 | 0.0070 | 9 |
| ALS-SNs | BP | GO:0001909 | BP leukocyte mediated cytotoxicity | 0.022 | 0.0075 | 27 |
| ALS-SNs | BP | GO:0001906 | BP cell killing | 0.027 | 0.0081 | 34 |
| ALS-SNs | BP | GO:0006911 | BP phagocytosis, engulfment | 0.010 | 0.0087 | 12 |
| ALS-SNs | BP | GO:0034109 | BP homotypic cell-cell adhesion | 0.014 | 0.0088 | 18 |
| ALS-SNs | BP | GO:0006885 | BP regulation of pH | 0.014 | 0.0101 | 18 |
| ALS-SNs | BP | GO:0043303 | BP mast cell degranulation | 0.010 | 0.0106 | 12 |
| ALS-SNs | BP | GO:0034103 | BP regulation of tissue remodeling | 0.011 | 0.0115 | 14 |
| ALS-SNs | BP | GO:0051014 | BP actin filament severing | 0.006 | 0.0115 | 7 |
| ALS-SNs | BP | GO:0090208 | BP positive regulation of triglyceride metabolic process | 0.006 | 0.0115 | 7 |
| ALS-SNs | BP | GO:0006869 | BP lipid transport | 0.038 | 0.0117 | 48 |
| ALS-SNs | BP | GO:0045017 | BP glycerolipid biosynthetic process | 0.029 | 0.0117 | 36 |
| ALS-SNs | BP | GO:0097191 | BP extrinsic apoptotic signaling pathway | 0.027 | 0.0125 | 34 |
| ALS-SNs | BP | GO:0071375 | BP cellular response to peptide hormone stimulus | 0.032 | 0.0128 | 40 |
| ALS-SNs | BP | GO:0045730 | BP respiratory burst | 0.009 | 0.0136 | 11 |
| ALS-SNs | BP | GO:0002090 | BP regulation of receptor internalization | 0.011 | 0.0136 | 14 |
| ALS-SNs | BP | GO:0034329 | BP cell junction assembly | 0.038 | 0.0141 | 48 |
| ALS-SNs | BP | GO:1900015 | BP regulation of cytokine production involved in inflammatory response | 0.010 | 0.0147 | 13 |
| ALS-SNs | BP | GO:0090594 | BP inflammatory response to wounding | 0.006 | 0.0147 | 8 |
| ALS-SNs | BP | GO:0140888 | BP interferon-mediated signaling pathway | 0.015 | 0.0151 | 19 |
| ALS-SNs | BP | GO:0019882 | BP antigen processing and presentation | 0.018 | 0.0156 | 22 |
| ALS-SNs | BP | GO:0042327 | BP positive regulation of phosphorylation | 0.042 | 0.0160 | 53 |
| ALS-SNs | BP | GO:0033238 | BP regulation of amine metabolic process | 0.006 | 0.0164 | 7 |
| ALS-SNs | BP | GO:0010867 | BP positive regulation of triglyceride biosynthetic process | 0.005 | 0.0164 | 6 |
| ALS-SNs | BP | GO:0072659 | BP protein localization to plasma membrane | 0.032 | 0.0164 | 40 |
| ALS-SNs | BP | GO:1903900 | BP regulation of viral life cycle | 0.020 | 0.0164 | 25 |
| ALS-SNs | BP | GO:0097242 | BP amyloid-beta clearance | 0.008 | 0.0164 | 10 |
| ALS-SNs | BP | GO:0051209 | BP release of sequestered calcium ion into cytosol | 0.015 | 0.0166 | 19 |

| | | | | | | |
|---|---|---|---|---|---|---|
| ALS-SNs | BP | GO:0002228 | BP natural killer cell mediated immunity | 0.014 | 0.0168 | 17 |
| ALS-SNs | BP | GO:0034446 | BP substrate adhesion-dependent cell spreading | 0.014 | 0.0193 | 17 |
| ALS-SNs | BP | GO:0010720 | BP positive regulation of cell development | 0.041 | 0.0194 | 51 |
| ALS-SNs | BP | GO:0007565 | BP female pregnancy | 0.017 | 0.0196 | 21 |
| ALS-SNs | BP | GO:0097006 | BP regulation of plasma lipoprotein particle levels | 0.010 | 0.0200 | 13 |
| ALS-SNs | BP | GO:0071396 | BP cellular response to lipid | 0.054 | 0.0202 | 67 |
| ALS-SNs | BP | GO:0031669 | BP cellular response to nutrient levels | 0.029 | 0.0204 | 36 |
| ALS-SNs | BP | GO:0034330 | BP cell junction organization | 0.058 | 0.0215 | 72 |
| ALS-SNs | BP | GO:0071216 | BP cellular response to biotic stimulus | 0.029 | 0.0218 | 36 |
| ALS-SNs | BP | GO:0070661 | BP leukocyte proliferation | 0.036 | 0.0232 | 45 |
| ALS-SNs | BP | GO:0031579 | BP membrane raft organization | 0.006 | 0.0244 | 8 |
| ALS-SNs | BP | GO:1903035 | BP negative regulation of response to wounding | 0.012 | 0.0244 | 15 |
| ALS-SNs | BP | GO:0034113 | BP heterotypic cell-cell adhesion | 0.010 | 0.0254 | 12 |
| ALS-SNs | BP | GO:0090148 | BP membrane fission | 0.010 | 0.0254 | 12 |
| ALS-SNs | BP | GO:0002040 | BP sprouting angiogenesis | 0.014 | 0.0266 | 18 |
| ALS-SNs | BP | GO:0046847 | BP filopodium assembly | 0.010 | 0.0269 | 13 |
| ALS-SNs | BP | GO:0018212 | BP peptidyl-tyrosine modification | 0.022 | 0.0289 | 27 |
| ALS-SNs | BP | GO:0014044 | BP Schwann cell development | 0.008 | 0.0299 | 10 |
| ALS-SNs | BP | GO:0030193 | BP regulation of blood coagulation | 0.010 | 0.0299 | 12 |
| ALS-SNs | BP | GO:0007212 | BP G protein-coupled dopamine receptor signaling pathway | 0.006 | 0.0311 | 8 |
| ALS-SNs | BP | GO:0042593 | BP glucose homeostasis | 0.025 | 0.0313 | 31 |
| ALS-SNs | BP | GO:0050870 | BP positive regulation of T cell activation | 0.030 | 0.0316 | 37 |
| ALS-SNs | BP | GO:0051384 | BP response to glucocorticoid | 0.014 | 0.0337 | 17 |
| ALS-SNs | BP | GO:0006067 | BP ethanol metabolic process | 0.004 | 0.0337 | 5 |
| ALS-SNs | BP | GO:0006068 | BP ethanol catabolic process | 0.004 | 0.0337 | 5 |
| ALS-SNs | BP | GO:0006497 | BP protein lipidation | 0.010 | 0.0344 | 12 |
| ALS-SNs | BP | GO:0007163 | BP establishment or maintenance of cell polarity | 0.025 | 0.0358 | 31 |
| ALS-SNs | BP | GO:0044706 | BP multi-multicellular organism process | 0.018 | 0.0358 | 22 |
| ALS-SNs | BP | GO:0043434 | BP response to peptide hormone | 0.038 | 0.0374 | 47 |
| ALS-SNs | BP | GO:0051338 | BP regulation of transferase activity | 0.046 | 0.0377 | 58 |
| ALS-SNs | BP | GO:1901699 | BP cellular response to nitrogen compound | 0.054 | 0.0377 | 67 |
| ALS-SNs | BP | GO:0050920 | BP regulation of chemotaxis | 0.021 | 0.0382 | 26 |
| ALS-SNs | BP | GO:0042832 | BP defense response to protozoan | 0.006 | 0.0382 | 8 |
| ALS-SNs | BP | GO:0060759 | BP regulation of response to cytokine stimulus | 0.022 | 0.0382 | 27 |
| ALS-SNs | BP | GO:0051235 | BP maintenance of location | 0.033 | 0.0387 | 41 |
| ALS-SNs | BP | GO:0046634 | BP regulation of alpha-beta T cell activation | 0.017 | 0.0408 | 21 |
| ALS-SNs | BP | GO:0006816 | BP calcium ion transport | 0.034 | 0.0408 | 43 |
| ALS-SNs | BP | GO:0043549 | BP regulation of kinase activity | 0.040 | 0.0412 | 50 |
| ALS-SNs | BP | GO:0055082 | BP intracellular chemical homeostasis | 0.056 | 0.0427 | 70 |
| ALS-SNs | BP | GO:0043328 | BP protein transport to vacuole involved in ubiquitin-dependent protein catabolic process via the multivesicular body sorting pathway | 0.005 | 0.0434 | 6 |
| ALS-SNs | BP | GO:0071709 | BP membrane assembly | 0.010 | 0.0445 | 12 |
| ALS-SNs | BP | GO:0044091 | BP membrane biogenesis | 0.010 | 0.0445 | 13 |
| ALS-SNs | BP | GO:1905952 | BP regulation of lipid localization | 0.018 | 0.0445 | 22 |
| ALS-SNs | BP | GO:0045936 | BP negative regulation of phosphate metabolic process | 0.030 | 0.0445 | 38 |
| ALS-SNs | BP | GO:0031667 | BP response to nutrient levels | 0.043 | 0.0445 | 54 |
| ALS-SNs | BP | GO:0061615 | BP glycolytic process through fructose-6-phosphate | 0.006 | 0.0451 | 7 |
| ALS-SNs | BP | GO:0002287 | BP alpha-beta T cell activation involved in immune response | 0.013 | 0.0458 | 16 |
| ALS-SNs | BP | GO:0072376 | BP protein activation cascade | 0.004 | 0.0460 | 5 |
| ALS-SNs | BP | GO:0141086 | BP negative regulation of inflammasome-mediated signaling pathway | 0.004 | 0.0460 | 5 |
| ALS-SNs | BP | GO:1903238 | BP positive regulation of leukocyte tethering or rolling | 0.004 | 0.0460 | 5 |
| ALS-SNs | BP | GO:0019320 | BP hexose catabolic process | 0.008 | 0.0460 | 10 |
| ALS-SNs | BP | GO:0009896 | BP positive regulation of catabolic process | 0.053 | 0.0464 | 66 |
| ALS-SNs | BP | GO:0050818 | BP regulation of coagulation | 0.010 | 0.0491 | 12 |
| ALS-SNs | BP | GO:0044089 | BP positive regulation of cellular component biogenesis | 0.046 | 0.0495 | 57 |
| ALS-SNs | CC | GO:0030667 | CC secretory granule membrane | 0.066 | 0.0000 | 85 |
| ALS-SNs | CC | GO:0101002 | CC ficolin-1-rich granule | 0.044 | 0.0000 | 57 |
| ALS-SNs | CC | GO:0015629 | CC actin cytoskeleton | 0.065 | 0.0000 | 84 |
| ALS-SNs | CC | GO:0005774 | CC vacuolar membrane | 0.069 | 0.0000 | 90 |

| | | | | | | |
|---|---|---|---|---|---|---|
| ALS-SNs | CC | GO:0030139 | CC endocytic vesicle | 0.049 | 0.0000 | 64 |
| ALS-SNs | CC | GO:0005766 | CC primary lysosome | 0.032 | 0.0000 | 41 |
| ALS-SNs | CC | GO:0031252 | CC cell leading edge | 0.052 | 0.0000 | 68 |
| ALS-SNs | CC | GO:0010008 | CC endosome membrane | 0.068 | 0.0000 | 88 |
| ALS-SNs | CC | GO:0030027 | CC lamellipodium | 0.031 | 0.0000 | 40 |
| ALS-SNs | CC | GO:0030055 | CC cell-substrate junction | 0.052 | 0.0000 | 68 |
| ALS-SNs | CC | GO:0005911 | CC cell-cell junction | 0.047 | 0.0001 | 61 |
| ALS-SNs | CC | GO:0005938 | CC cell cortex | 0.037 | 0.0002 | 48 |
| ALS-SNs | CC | GO:0005811 | CC lipid droplet | 0.017 | 0.0003 | 22 |
| ALS-SNs | CC | GO:0034045 | CC phagophore assembly site membrane | 0.007 | 0.0015 | 9 |
| ALS-SNs | CC | GO:0030863 | CC cortical cytoskeleton | 0.015 | 0.0018 | 20 |
| ALS-SNs | CC | GO:0001931 | CC uropod | 0.005 | 0.0020 | 7 |
| ALS-SNs | CC | GO:0005775 | CC vacuolar lumen | 0.023 | 0.0025 | 30 |
| ALS-SNs | CC | GO:0043020 | CC NADPH oxidase complex | 0.005 | 0.0027 | 6 |
| ALS-SNs | CC | GO:0098552 | CC side of membrane | 0.061 | 0.0033 | 79 |
| ALS-SNs | CC | GO:0031254 | CC cell trailing edge | 0.005 | 0.0033 | 7 |
| ALS-SNs | CC | GO:0098978 | CC glutamatergic synapse | 0.047 | 0.0045 | 61 |
| ALS-SNs | CC | GO:0032587 | CC ruffle membrane | 0.015 | 0.0052 | 20 |
| ALS-SNs | CC | GO:0098793 | CC presynapse | 0.049 | 0.0080 | 64 |
| ALS-SNs | CC | GO:0030496 | CC midbody | 0.025 | 0.0109 | 33 |
| ALS-SNs | CC | GO:0030117 | CC membrane coat | 0.015 | 0.0145 | 20 |
| ALS-SNs | CC | GO:0048475 | CC coated membrane | 0.015 | 0.0145 | 20 |
| ALS-SNs | CC | GO:0098857 | CC membrane microdomain | 0.028 | 0.0185 | 36 |
| ALS-SNs | CC | GO:0008305 | CC integrin complex | 0.006 | 0.0224 | 8 |
| ALS-SNs | CC | GO:0031143 | CC pseudopodium | 0.004 | 0.0305 | 5 |
| ALS-SNs | MF | GO:0003779 | MF actin binding | 0.051 | 0.0000 | 66 |
| ALS-SNs | MF | GO:0050839 | MF cell adhesion molecule binding | 0.061 | 0.0001 | 78 |
| ALS-SNs | MF | GO:0019902 | MF phosphatase binding | 0.028 | 0.0038 | 36 |
| ALS-SNs | MF | GO:0005178 | MF integrin binding | 0.020 | 0.0038 | 26 |
| ALS-SNs | MF | GO:0005543 | MF phospholipid binding | 0.049 | 0.0066 | 63 |
| ALS-SNs | MF | GO:0031267 | MF small GTPase binding | 0.036 | 0.0090 | 46 |
| ALS-SNs | MF | GO:0043130 | MF ubiquitin binding | 0.018 | 0.0121 | 23 |
| ALS-SNs | MF | GO:0004683 | MF calcium/calmodulin-dependent protein kinase activity | 0.006 | 0.0150 | 8 |
| ALS-SNs | MF | GO:0003924 | MF GTPase activity | 0.036 | 0.0150 | 47 |
| ALS-SNs | MF | GO:0048029 | MF monosaccharide binding | 0.012 | 0.0150 | 16 |
| ALS-SNs | MF | GO:0038187 | MF pattern recognition receptor activity | 0.009 | 0.0218 | 12 |
| ALS-SNs | MF | GO:0019904 | MF protein domain specific binding | 0.061 | 0.0262 | 78 |
| ALS-SNs | MF | GO:0017022 | MF myosin binding | 0.010 | 0.0441 | 13 |
| ALS-SNs | MF | GO:0140313 | MF molecular sequestering activity | 0.009 | 0.0452 | 12 |
| ALS-SNs | MF | GO:0030695 | MF GTPase regulator activity | 0.049 | 0.0452 | 63 |
| ALS-SNs | MF | GO:0042826 | MF histone deacetylase binding | 0.016 | 0.0452 | 21 |
| ALS-SNs | MF | GO:0033691 | MF sialic acid binding | 0.005 | 0.0467 | 7 |
| ALS-SNs | MF | GO:0001784 | MF phosphotyrosine residue binding | 0.009 | 0.0467 | 12 |
| ALS-SNs | React | R-HSA-6798695 | React Neutrophil degranulation | 0.148 | 0.0000 | 132 |
| ALS-SNs | React | R-HSA-76002 | React Platelet activation, signaling and aggregation | 0.072 | 0.0000 | 64 |
| ALS-SNs | React | R-HSA-109582 | React Hemostasis | 0.121 | 0.0000 | 108 |
| ALS-SNs | React | R-HSA-76005 | React Response to elevated platelet cytosolic Ca2+ | 0.037 | 0.0000 | 33 |
| ALS-SNs | React | R-HSA-114608 | React Platelet degranulation | 0.036 | 0.0000 | 32 |
| ALS-SNs | React | R-HSA-9658195 | React Leishmania infection | 0.044 | 0.0000 | 39 |
| ALS-SNs | React | R-HSA-9824443 | React Parasitic Infection Pathways | 0.044 | 0.0000 | 39 |
| ALS-SNs | React | R-HSA-9006934 | React Signaling by Receptor Tyrosine Kinases | 0.090 | 0.0001 | 80 |
| ALS-SNs | React | R-HSA-373760 | React L1CAM interactions | 0.028 | 0.0002 | 25 |
| ALS-SNs | React | R-HSA-6802946 | React Signaling by moderate kinase activity BRAF mutants | 0.018 | 0.0002 | 16 |
| ALS-SNs | React | R-HSA-6802949 | React Signaling by RAS mutants | 0.018 | 0.0002 | 16 |
| ALS-SNs | React | R-HSA-6802955 | React Paradoxical activation of RAF signaling by kinase inactive BRAF | 0.018 | 0.0002 | 16 |
| ALS-SNs | React | R-HSA-9649948 | React Signaling downstream of RAS mutants | 0.018 | 0.0002 | 16 |
| ALS-SNs | React | R-HSA-9656223 | React Signaling by RAF1 mutants | 0.017 | 0.0002 | 15 |

| | | | | | | |
|---|---|---|---|---|---|---|
| ALS-SNs | React | R-HSA-449147 | React Signaling by Interleukins | 0.080 | 0.0005 | 71 |
| ALS-SNs | React | R-HSA-6802948 | React Signaling by high-kinase activity BRAF mutants | 0.015 | 0.0007 | 13 |
| ALS-SNs | React | R-HSA-6802952 | React Signaling by BRAF and RAF1 fusions | 0.021 | 0.0009 | 19 |
| ALS-SNs | React | R-HSA-373755 | React Semaphorin interactions | 0.019 | 0.0013 | 17 |
| ALS-SNs | React | R-HSA-372790 | React Signaling by GPCR | 0.075 | 0.0018 | 67 |
| ALS-SNs | React | R-HSA-114604 | React GPVI-mediated activation cascade | 0.015 | 0.0021 | 13 |
| ALS-SNs | React | R-HSA-5674135 | React MAP2K and MAPK activation | 0.015 | 0.0021 | 13 |
| ALS-SNs | React | R-HSA-190828 | React Gap junction trafficking | 0.012 | 0.0021 | 11 |
| ALS-SNs | React | R-HSA-9012999 | React RHO GTPase cycle | 0.076 | 0.0030 | 68 |
| ALS-SNs | React | R-HSA-157858 | React Gap junction trafficking and regulation | 0.012 | 0.0030 | 11 |
| ALS-SNs | React | R-HSA-9662851 | React Anti-inflammatory response favouring Leishmania parasite infection | 0.021 | 0.0030 | 19 |
| ALS-SNs | React | R-HSA-9664433 | React Leishmania parasite growth and survival | 0.021 | 0.0030 | 19 |
| ALS-SNs | React | R-HSA-388396 | React GPCR downstream signalling | 0.068 | 0.0030 | 61 |
| ALS-SNs | React | R-HSA-437239 | React Recycling pathway of L1 | 0.015 | 0.0046 | 13 |
| ALS-SNs | React | R-HSA-2029480 | React Fcgamma receptor (FCGR) dependent phagocytosis | 0.025 | 0.0046 | 22 |
| ALS-SNs | React | R-HSA-198933 | React Immunoregulatory interactions between a Lymphoid and a non-Lymphoid cell | 0.030 | 0.0047 | 27 |
| ALS-SNs | React | R-HSA-112316 | React Neuronal System | 0.048 | 0.0047 | 43 |
| ALS-SNs | React | R-HSA-1474244 | React Extracellular matrix organization | 0.040 | 0.0048 | 36 |
| ALS-SNs | React | R-HSA-112314 | React Neurotransmitter receptors and postsynaptic signal transmission | 0.030 | 0.0048 | 27 |
| ALS-SNs | React | R-HSA-1222556 | React ROS and RNS production in phagocytes | 0.012 | 0.0048 | 11 |
| ALS-SNs | React | R-HSA-164952 | React The role of Nef in HIV-1 replication and disease pathogenesis | 0.012 | 0.0048 | 11 |
| ALS-SNs | React | R-HSA-76009 | React Platelet Aggregation (Plug Formation) | 0.012 | 0.0048 | 11 |
| ALS-SNs | React | R-HSA-445144 | React Signal transduction by L1 | 0.010 | 0.0048 | 9 |
| ALS-SNs | React | R-HSA-6802957 | React Oncogenic MAPK signaling | 0.022 | 0.0050 | 20 |
| ALS-SNs | React | R-HSA-446353 | React Cell-extracellular matrix interactions | 0.008 | 0.0050 | 7 |
| ALS-SNs | React | R-HSA-166520 | React Signaling by NTRKs | 0.029 | 0.0055 | 26 |
| ALS-SNs | React | R-HSA-9612973 | React Autophagy | 0.035 | 0.0068 | 31 |
| ALS-SNs | React | R-HSA-438064 | React Post NMDA receptor activation events | 0.018 | 0.0087 | 16 |
| ALS-SNs | React | R-HSA-397795 | React G-protein beta:gamma signalling | 0.011 | 0.0095 | 10 |
| ALS-SNs | React | R-HSA-5626467 | React RHO GTPases activate IQGAPs | 0.011 | 0.0095 | 10 |
| ALS-SNs | React | R-HSA-354192 | React Integrin signaling | 0.010 | 0.0096 | 9 |
| ALS-SNs | React | R-HSA-9664407 | React Parasite infection | 0.018 | 0.0096 | 16 |
| ALS-SNs | React | R-HSA-9664417 | React Leishmania phagocytosis | 0.018 | 0.0096 | 16 |
| ALS-SNs | React | R-HSA-9664422 | React FCGR3A-mediated phagocytosis | 0.018 | 0.0096 | 16 |
| ALS-SNs | React | R-HSA-442755 | React Activation of NMDA receptors and postsynaptic events | 0.019 | 0.0110 | 17 |
| ALS-SNs | React | R-HSA-195258 | React RHO GTPase Effectors | 0.054 | 0.0110 | 48 |

| | | | | | | |
|---|---|---|---|---|---|---|
| ALS-SNs | React | R-HSA-2682334 | React EPH-Ephrin signaling | 0.021 | 0.0132 | 19 |
| ALS-SNs | React | R-HSA-4420097 | React VEGFA-VEGFR2 Pathway | 0.025 | 0.0132 | 22 |
| ALS-SNs | React | R-HSA-1483257 | React Phospholipid metabolism | 0.039 | 0.0132 | 35 |
| ALS-SNs | React | R-HSA-2029482 | React Regulation of actin dynamics for phagocytic cup formation | 0.018 | 0.0132 | 16 |
| ALS-SNs | React | R-HSA-194138 | React Signaling by VEGF | 0.026 | 0.0132 | 23 |
| ALS-SNs | React | R-HSA-877300 | React Interferon gamma signaling | 0.024 | 0.0142 | 21 |
| ALS-SNs | React | R-HSA-418594 | React G alpha (i) signalling events | 0.037 | 0.0159 | 33 |
| ALS-SNs | React | R-HSA-187037 | React Signaling by NTRK1 (TRKA) | 0.025 | 0.0159 | 22 |
| ALS-SNs | React | R-HSA-354194 | React GRB2:SOS provides linkage to MAPK signaling for Integrins | 0.007 | 0.0159 | 6 |
| ALS-SNs | React | R-HSA-399955 | React SEMA3A-Plexin repulsion signaling by inhibiting Integrin adhesion | 0.007 | 0.0159 | 6 |
| ALS-SNs | React | R-HSA-9651496 | React Defects of contact activation system (CAS) and kallikrein/kinin system (KKS) | 0.007 | 0.0159 | 6 |
| ALS-SNs | React | R-HSA-202733 | React Cell surface interactions at the vascular wall | 0.026 | 0.0159 | 23 |
| ALS-SNs | React | R-HSA-909733 | React Interferon alpha/beta signaling | 0.019 | 0.0159 | 17 |
| ALS-SNs | React | R-HSA-8854214 | React TBC/RABGAPs | 0.015 | 0.0159 | 13 |
| ALS-SNs | React | R-HSA-112315 | React Transmission across Chemical Synapses | 0.035 | 0.0159 | 31 |
| ALS-SNs | React | R-HSA-9013149 | React RAC1 GTPase cycle | 0.035 | 0.0177 | 31 |
| ALS-SNs | React | R-HSA-1500931 | React Cell-Cell communication | 0.024 | 0.0194 | 21 |
| ALS-SNs | React | R-HSA-2172127 | React DAP12 interactions | 0.013 | 0.0197 | 12 |
| ALS-SNs | React | R-HSA-9668328 | React Sealing of the nuclear envelope (NE) by ESCRT-III | 0.010 | 0.0210 | 9 |
| ALS-SNs | React | R-HSA-1632852 | React Macroautophagy | 0.030 | 0.0212 | 27 |
| ALS-SNs | React | R-HSA-6807878 | React COPI-mediated anterograde transport | 0.024 | 0.0214 | 21 |
| ALS-SNs | React | R-HSA-392451 | React G beta:gamma signalling through PI3Kgamma | 0.009 | 0.0225 | 8 |
| ALS-SNs | React | R-HSA-5668599 | React RHO GTPases Activate NADPH Oxidases | 0.009 | 0.0225 | 8 |
| ALS-SNs | React | R-HSA-9013148 | React CDC42 GTPase cycle | 0.029 | 0.0225 | 26 |
| ALS-SNs | React | R-HSA-451927 | React Interleukin-2 family signaling | 0.012 | 0.0225 | 11 |
| ALS-SNs | React | R-HSA-190840 | React Microtubule-dependent trafficking of connexons from Golgi to the plasma membrane | 0.007 | 0.0225 | 6 |
| ALS-SNs | React | R-HSA-190872 | React Transport of connexons to the plasma membrane | 0.007 | 0.0225 | 6 |
| ALS-SNs | React | R-HSA-9671793 | React Diseases of hemostasis | 0.007 | 0.0225 | 6 |
| ALS-SNs | React | R-HSA-216083 | React Integrin cell surface interactions | 0.016 | 0.0247 | 14 |
| ALS-SNs | React | R-HSA-8873719 | React RAB geranylgeranylation | 0.016 | 0.0247 | 14 |
| ALS-SNs | React | R-HSA-2132295 | React MHC class II antigen presentation | 0.026 | 0.0255 | 23 |
| ALS-SNs | React | R-HSA-9619483 | React Activation of AMPK downstream of NMDARs | 0.009 | 0.0297 | 8 |
| ALS-SNs | React | R-HSA-111885 | React Opioid Signalling | 0.019 | 0.0308 | 17 |
| ALS-SNs | React | R-HSA-111933 | React Calmodulin induced events | 0.010 | 0.0321 | 9 |
| ALS-SNs | React | R-HSA-111997 | React CaM pathway | 0.010 | 0.0321 | 9 |
| ALS-SNs | React | R-HSA-112040 | React G-protein mediated events | 0.013 | 0.0321 | 12 |

| | | | | | | |
|---|---|---|---|---|---|---|
| ALS-SNs | React | R-HSA-3928662 | React EPHB-mediated forward signaling | 0.012 | 0.0321 | 11 |
| ALS-SNs | React | R-HSA-2871796 | React FCERI mediated MAPK activation | 0.011 | 0.0321 | 10 |
| ALS-SNs | React | R-HSA-190861 | React Gap junction assembly | 0.007 | 0.0321 | 6 |
| ALS-SNs | React | R-HSA-2142691 | React Synthesis of Leukotrienes (LT) and Eoxins (EX) | 0.007 | 0.0321 | 6 |
| ALS-SNs | React | R-HSA-399954 | React Sema3A PAK dependent Axon repulsion | 0.007 | 0.0321 | 6 |
| ALS-SNs | React | R-HSA-199977 | React ER to Golgi Anterograde Transport | 0.031 | 0.0321 | 28 |
| ALS-SNs | React | R-HSA-381038 | React XBP1(S) activates chaperone genes | 0.015 | 0.0349 | 13 |
| ALS-SNs | React | R-HSA-1445148 | React Translocation of SLC2A4 (GLUT4) to the plasma membrane | 0.018 | 0.0432 | 16 |
| ALS-SNs | React | R-HSA-196025 | React Formation of annular gap junctions | 0.006 | 0.0471 | 5 |
| ALS-SNs | React | R-HSA-372708 | React p130Cas linkage to MAPK signaling for integrins | 0.006 | 0.0471 | 5 |
| ALS-SNs | React | R-HSA-198323 | React AKT phosphorylates targets in the cytosol | 0.007 | 0.0471 | 6 |
| ALS-SNs | React | R-HSA-391160 | React Signal regulatory protein family interactions | 0.007 | 0.0471 | 6 |
| ALS-SNs | React | R-HSA-112043 | React PLC beta mediated events | 0.012 | 0.0471 | 11 |
| ALS-SNs | React | R-HSA-948021 | React Transport to the Golgi and subsequent modification | 0.035 | 0.0471 | 31 |
| ALS-SNs | React | R-HSA-381070 | React IRE1alpha activates chaperones | 0.015 | 0.0471 | 13 |
| ALS-SNs | React | R-HSA-9013404 | React RAC2 GTPase cycle | 0.019 | 0.0471 | 17 |
| ALS-SNs | React | R-HSA-8856828 | React Clathrin-mediated endocytosis | 0.027 | 0.0471 | 24 |
| ALS-SNs | React | R-HSA-8980692 | React RHOA GTPase cycle | 0.027 | 0.0471 | 24 |
| ALS-SNs | React | R-HSA-111996 | React Ca-dependent events | 0.010 | 0.0471 | 9 |
| ALS-OxA | BP | GO:0022613 | BP Ribonucleoprotein complex biogenesis | 0.103 | 0.0000 | 73 |
| ALS-OxA | BP | GO:0008380 | BP Rna splicing | 0.083 | 0.0000 | 59 |
| ALS-OxA | BP | GO:0006364 | BP Rrna processing | 0.048 | 0.0001 | 34 |
| ALS-OxA | BP | GO:0022618 | BP Protein-rna complex assembly | 0.042 | 0.0003 | 30 |
| ALS-OxA | BP | GO:0016072 | BP Rrna metabolic process | 0.052 | 0.0003 | 37 |
| ALS-OxA | BP | GO:0071826 | BP Protein-rna complex organization | 0.042 | 0.0006 | 30 |
| ALS-OxA | BP | GO:0001510 | BP Rna methylation | 0.021 | 0.0126 | 15 |
| ALS-OxA | BP | GO:0002181 | BP Cytoplasmic translation | 0.031 | 0.0183 | 22 |
| ALS-OxA | BP | GO:0006119 | BP Oxidative phosphorylation | 0.025 | 0.0413 | 18 |
| ALS-OxA | CC | GO:0005681 | CC Spliceosomal complex | 0.048 | 0.0000 | 35 |
| ALS-OxA | CC | GO:0019866 | CC Organelle inner membrane | 0.078 | 0.0001 | 57 |
| ALS-OxA | CC | GO:0044391 | CC Ribosomal subunit | 0.039 | 0.0001 | 29 |
| ALS-OxA | CC | GO:0098687 | CC Chromosomal region | 0.056 | 0.0009 | 41 |
| ALS-OxA | CC | GO:0098798 | CC Mitochondrial protein-containing complex | 0.034 | 0.0013 | 25 |
| ALS-OxA | CC | GO:0034709 | CC Methylosome | 0.008 | 0.0013 | 6 |
| ALS-OxA | CC | GO:0005763 | CC Mitochondrial small ribosomal subunit | 0.012 | 0.0025 | 9 |
| ALS-OxA | CC | GO:0120114 | CC Sm-like protein family complex | 0.020 | 0.0057 | 15 |
| ALS-OxA | CC | GO:0030684 | CC Preribosome | 0.022 | 0.0082 | 16 |
| ALS-OxA | CC | GO:0098803 | CC Respiratory chain complex | 0.019 | 0.0082 | 14 |
| ALS-OxA | CC | GO:0035098 | CC Esc/e(z) complex | 0.007 | 0.0179 | 5 |
| ALS-OxA | CC | GO:0000792 | CC Heterochromatin | 0.015 | 0.0243 | 11 |
| ALS-OxA | CC | GO:0034719 | CC Smn-sm protein complex | 0.007 | 0.0302 | 5 |
| ALS-OxA | CC | GO:0015030 | CC Cajal body | 0.014 | 0.0343 | 10 |
| ALS-OxA | CC | GO:0045259 | CC Proton-transporting atp synthase complex | 0.007 | 0.0487 | 5 |
| ALS-OxA | MF | GO:0003735 | MF Structural constituent of ribosome | 0.039 | 0.0003 | 28 |
| ALS-OxA | React | R-HSA-72163 | React mRNA Splicing - Major Pathway | 0.073 | 0.0000 | 37 |
| ALS-OxA | React | R-HSA-72203 | React Processing of Capped Intron-Containing Pre-mRNA | 0.089 | 0.0000 | 45 |
| ALS-OxA | React | R-HSA-72172 | React mRNA Splicing | 0.073 | 0.0000 | 37 |
| ALS-OxA | React | R-HSA-72766 | React Translation | 0.087 | 0.0000 | 44 |
| ALS-OxA | React | R-HSA-72706 | React GTP hydrolysis and joining of the 60S ribosomal subunit | 0.043 | 0.0003 | 22 |

| | | | | | | |
|---|---|---|---|---|---|---|
| ALS-OxA | React | R-HSA-72613 | React Eukaryotic Translation Initiation | 0.043 | 0.0006 | 22 |
| ALS-OxA | React | R-HSA-72737 | React Cap-dependent Translation Initiation | 0.043 | 0.0006 | 22 |
| ALS-OxA | React | R-HSA-8868773 | React rRNA processing in the nucleus and cytosol | 0.059 | 0.0006 | 30 |
| ALS-OxA | React | R-HSA-6791226 | React Major pathway of rRNA processing in the nucleolus and cytosol | 0.057 | 0.0006 | 29 |
| ALS-OxA | React | R-HSA-72312 | React rRNA processing | 0.059 | 0.0014 | 30 |
| ALS-OxA | React | R-HSA-156827 | React L13a-mediated translational silencing of Ceruloplasmin expression | 0.039 | 0.0014 | 20 |
| ALS-OxA | React | R-HSA-168273 | React Influenza Viral RNA Transcription and Replication | 0.043 | 0.0037 | 22 |
| ALS-OxA | React | R-HSA-72689 | React Formation of a pool of free 40S subunits | 0.035 | 0.0037 | 18 |
| ALS-OxA | React | R-HSA-168255 | React Influenza Infection | 0.047 | 0.0037 | 24 |
| ALS-OxA | React | R-HSA-73856 | React RNA Polymerase II Transcription Termination | 0.028 | 0.0043 | 14 |
| ALS-OxA | React | R-HSA-192823 | React Viral mRNA Translation | 0.031 | 0.0075 | 16 |
| ALS-OxA | React | R-HSA-5368287 | React Mitochondrial translation | 0.033 | 0.0086 | 17 |
| ALS-OxA | React | R-HSA-9010553 | React Regulation of expression of SLITs and ROBOs | 0.045 | 0.0088 | 23 |
| ALS-OxA | React | R-HSA-9633012 | React Response of EIF2AK4 (GCN2) to amino acid deficiency | 0.033 | 0.0089 | 17 |
| ALS-OxA | React | R-HSA-2408557 | React Selenocysteine synthesis | 0.031 | 0.0097 | 16 |
| ALS-OxA | React | R-HSA-5389840 | React Mitochondrial translation elongation | 0.031 | 0.0097 | 16 |
| ALS-OxA | React | R-HSA-5419276 | React Mitochondrial translation termination | 0.031 | 0.0097 | 16 |
| ALS-OxA | React | R-HSA-376176 | React Signaling by ROBO receptors | 0.051 | 0.0102 | 26 |
| ALS-OxA | React | R-HSA-975956 | React Nonsense Mediated Decay (NMD) independent of the Exon Junction Complex (EJC) | 0.031 | 0.0102 | 16 |
| ALS-OxA | React | R-HSA-111367 | React SLBP independent Processing of Histone Pre-mRNAs | 0.010 | 0.0114 | 5 |
| ALS-OxA | React | R-HSA-2467813 | React Separation of Sister Chromatids | 0.047 | 0.0119 | 24 |
| ALS-OxA | React | R-HSA-69618 | React Mitotic Spindle Checkpoint | 0.033 | 0.0119 | 17 |
| ALS-OxA | React | R-HSA-72165 | React mRNA Splicing - Minor Pathway | 0.022 | 0.0128 | 11 |
| ALS-OxA | React | R-HSA-156902 | React Peptide chain elongation | 0.030 | 0.0141 | 15 |
| ALS-OxA | React | R-HSA-77588 | React SLBP Dependent Processing of Replication-Dependent Histone Pre-mRNAs | 0.010 | 0.0165 | 5 |
| ALS-OxA | React | R-HSA-156842 | React Eukaryotic Translation Elongation | 0.030 | 0.0182 | 15 |
| ALS-OxA | React | R-HSA-72764 | React Eukaryotic Translation Termination | 0.030 | 0.0182 | 15 |
| ALS-OxA | React | R-HSA-191859 | React snRNP Assembly | 0.022 | 0.0182 | 11 |
| ALS-OxA | React | R-HSA-194441 | React Metabolism of non-coding RNA | 0.022 | 0.0182 | 11 |
| ALS-OxA | React | R-HSA-1799339 | React SRP-dependent cotranslational protein targeting to membrane | 0.033 | 0.0183 | 17 |
| ALS-OxA | React | R-HSA-159236 | React Transport of Mature mRNA derived from an Intron-Containing Transcript | 0.026 | 0.0183 | 13 |
| ALS-OxA | React | R-HSA-5368286 | React Mitochondrial translation initiation | 0.030 | 0.0187 | 15 |
| ALS-OxA | React | R-HSA-68882 | React Mitotic Anaphase | 0.053 | 0.0191 | 27 |
| ALS-OxA | React | R-HSA-2555396 | React Mitotic Metaphase and Anaphase | 0.053 | 0.0191 | 27 |
| ALS-OxA | React | R-HSA-611105 | React Respiratory electron transport | 0.039 | 0.0191 | 20 |
| ALS-OxA | React | R-HSA-2408522 | React Selenoamino acid metabolism | 0.033 | 0.0191 | 17 |

| | | | | | | |
|---|---|---|---|---|---|---|
| ALS-OxA | React | R-HSA-927802 | React Nonsense-Mediated Decay (NMD) | 0.033 | 0.0191 | 17 |
| ALS-OxA | React | R-HSA-975957 | React Nonsense Mediated Decay (NMD) enhanced by the Exon Junction Complex (EJC) | 0.033 | 0.0191 | 17 |
| ALS-OxA | React | R-HSA-68886 | React M Phase | 0.075 | 0.0471 | 38 |
| ALS-OxA | React | R-HSA-72202 | React Transport of Mature Transcript to Cytoplasm | 0.026 | 0.0479 | 13 |
| ALS-OxA | React | R-HSA-69620 | React Cell Cycle Checkpoints | 0.057 | 0.0484 | 29 |
| ALS-OxA | React | R-HSA-1428517 | React Aerobic respiration and respiratory electron transport | 0.053 | 0.0484 | 27 |

**Suppl Table 9.** Beta regression coefficients and odds ratios for disease progression-subtype probability models.

| Collection Point (Continous, Range 0 - 1) | | | | | |
|---|---|---|---|---|---|
| **Coefficients** | **Estimate** | **Odds ratio** | **Standard Error** | **z value** | **p value** |
| Cluster: OxA | -1.02 | 0.36 | 0.79 | -1.29 | 0.195 |
| Cluster: SNs | -1.46 | 0.23 | 0.69 | -2.11 | 0.035 |
| Sex Male | -0.17 | 0.84 | 0.21 | -0.81 | 0.415 |
| Age at collection | 0.01 | 1.01 | 0.01 | 1.63 | 0.103 |
| *Collection point:ClusterNeu* | *-2.63* | *0.07* | *0.90* | *-2.94* | *0.003* |
| Collection point:ClusterOxA | 1.86 | 6.39 | 1.24 | 1.50 | 0.133 |
| *Collection point:ClusterSNs* | *3.16* | *23.52* | *1.10* | *2.88* | *0.004* |
| **Collection Point (Categorical, Early vs Late)** | | | | | |
| Cluster: OxA | -0.17 | 0.84 | 0.51 | -0.34 | 0.73 |
| Cluster: SNs | -0.34 | 0.71 | 0.45 | -0.76 | 0.45 |
| Sex Male | -0.19 | 0.83 | 0.27 | -0.69 | 0.49 |
| Age at collection | 0.01 | 1.01 | 0.01 | 0.98 | 0.33 |
| *Collection paperlate:ClusterNeu* | *-1.10* | *0.33* | *0.51* | *-2.16* | *0.03* |
| Collection paperlate:ClusterOxA | 0.60 | 1.82 | 0.70 | 0.85 | 0.39 |
| *Collection paperlate:ClusterSNs* | *1.44* | *4.22* | *0.63* | *2.30* | *0.02* |

### Supplemental NYGC ALS Consortium authors

Hemali Phatnani, PhD[1], Justin Kwan, MD[2], Dhruv Sareen, PhD[3], James R. Broach, PhD[4], Zachary Simmons, MD[5], Ximena Arcila-Londono, MD[6], Edward B. Lee, MD, PhD[7], Vivianna M. Van Deerlin, MD, PhD[8], Neil A. Shneider, MD, PhD[9], Ernest Fraenkel, PhD[10], Lyle W. Ostrow, MD, PhD[11], Frank Baas, MD, PhD[12], Noah Zaitlen, PhD[13], James D. Berry, MD, MPH[14], Andrea Malaspina, MD, PhD[15], Pietro Fratta, MD, PhD[16], Gregory A. Cox, PhD[17], Leslie M. Thompson, PhD[18], Steve Finkbeiner, MD, PhD[19], Efthimios Dardiotis, MD, PhD[20], Timothy M. Miller, MD, PhD[21], Siddharthan Chandran, PhD[22], Suvankar Pal, MD[23], Eran Hornstein, MD, PhD[24], Daniel J. MacGowan, MD[25], Terry Heiman-Patterson, MD[26], Molly G. Hammell, PhD[27], Nikolaos. A. Patsopoulos, MD, PhD[28], Oleg Butovsky, PhD[29], Joshua Dubnau, PhD[30], Avindra Nath, MD[31], Robert Bowser, PhD[32], Matthew Harms, MD[33], Eleonora Aronica, MD, PhD[34], Mary Poss, DVM, PhD[35], Jennifer Phillips-Cremins, PhD[36], John Crary, MD, PhD[37], Nazem Atassi, MD[38], Dale J. Lange, MD[39], Darius J. Adams, MD[40], Leonidas Stefanis, MD, PhD[41], Marc Gotkine, MBBS[42], Robert H. Baloh, MD, PhD[43], Suma Babu, MBBS, MPH[44], Towfique Raj, PhD[45], Sabrina Paganoni, MD, PhD[46], Ophir Shalem, PhD[47], Colin Smith, MD[48], Bin Zhang, PhD[49], Justin Kwan MD, Thomas Blanchard PhD[50], Brent Harris, MD, PhD[51], Iris Broce, PhD[52], Vivian Drory, MD[53], John Ravits, MD[54], Corey McMillan, PhD[55], Vilas Menon, PhD[56], Lani Wu, PhD[57], Steven Altschuler, PhD[58], Yossef Lerner, MD[59], Rita Sattler, PhD[60], Kendall Van Keuren-

Jensen, PhD[61], Orit Rozenblatt-Rosen, PhD[62], Kerstin Lindblad-Toh, PhD[63], Katharine Nicholson, MD[64], Peter Gregersen, MD[65], Jeong-Ho Lee M.D, PhD[66], Matt Brauer VP, Data Sciences, Tara Nickerson CBO[67], Shameek Biswas Director, Kimberly A Wilson Executive Director, Translational Epidemiology[68], Sulev Koks, PhD[69], Stephen Muljo, PhD[70], Bryan J. Traynor, MD, PhD[71], Robert Moccia PhD, Seng Cheng SVP, Chief Scientific Officer, Rare Disease Research Unit [72], Andrew Deubler VP, Giovanni Coppola PhD, Mickey Atwal PhD, Michael Cantor PhD, William Salerno PhD, Eli Stahl PhD, Matt Anderson PhD, David Frendewey PhD.[73], Daphne Koller PhD, Mary Rozenman CBO/CFO[74]

[1] Center for Genomics of Neurodegenerative Disease (CGND), New York Genome Center, New York, NY

[2] Department of Neurology, Lewis Katz School of Medicine, Temple University, Philadelphia, PA

[3] Cedars-Sinai Biomanufacturing Center, Department of Biomedical Sciences, Board of Governors Regenerative Medicine Institute and Brain Program, Cedars-Sinai Medical Center

[4] Department of Biochemistry and Molecular Biology, Penn State Institute for Personalized Medicine, The Pennsylvania State University, Hershey, PA

[5] Department of Neurology, The Pennsylvania State University, Hershey, PA

[6] Department of Neurology, Henry Ford Health System, Detroit, MI

[7] Department of Pathology and Laboratory Medicine, Perelman School of Medicine, University of Pennsylvania, Philadelphia, PA

[8] Department of Pathology and Laboratory Medicine, Perelman School of Medicine, University of Pennsylvania, Philadelphia, PA

[9] Department of Neurology, Center for Motor Neuron Biology and Disease, Institute for Genomic Medicine, Columbia University, New York, NY

[10] Department of Biological Engineering, Massachusetts Institute of Technology, Cambridge, MA

[11] Department of Neurology, Johns Hopkins School of Medicine, Baltimore, MD

[12] Department of Neurogenetics, Academic Medical Centre, Amsterdam and Leiden University Medical Center, Leiden, The Netherlands

[13] Department of Medicine, Lung Biology Center, University of California, San Francisco, CA

[14] ALS Multidisciplinary Clinic, Neuromuscular Division, Department of Neurology, Harvard Medical School, and Neurological Clinical Research Institute, Massachusetts General Hospital, Boston, MA

[15] Centre for Neuroscience and Trauma, Blizard Institute, Barts and The London School of Medicine and Dentistry, Queen Mary University of London, London, and Department of Neurology, Basildon University Hospital, Basildon, United Kingdom

[16] Institute of Neurology, National Hospital for Neurology and Neurosurgery, University College London, London, United Kingdom

[17] The Jackson Laboratory, Bar Harbor, ME

[18] Department of Psychiatry & Human Behavior, Department of Biological Chemistry, School of Medicine, and Department of Neurobiology and Behavior, School of Biological Sciences, University California, Irvine, CA

[19] Taube/Koret Center for Neurodegenerative Disease Research, Roddenberry Center for Stem Cell Biology and Medicine, Gladstone Institute

[20] Department of Neurology & Sensory Organs, University of Thessaly, Thessaly, Greece

[21] Department of Neurology ,Washington University in St. Louis, St. Louis, MO

[22] Centre for Clinical Brain Sciences, Anne Rowling Regenerative Neurology Clinic, Euan MacDonald Centre for Motor Neurone Disease Research, University of Edinburgh, Edinburgh, United Kingdom
[23] Centre for Clinical Brain Sciences, Anne Rowling Regenerative Neurology Clinic, Euan MacDonald Centre for Motor Neurone Disease Research, University of Edinburgh, Edinburgh, United Kingdom
[24] Department of Molecular Genetics, Weizmann Institute of Science, Rehovot, Israel

[25] Department of Neurology, Icahn School of Medicine at Mount Sinai, New York, NY

[26] Center for Neurodegenerative Disorders, Department of Neurology, the Lewis Katz School of Medicine, Temple University, Philadelphia, PA

[27] Cold Spring Harbor Laboratory, Cold Spring Harbor, NY

[28] Computer Science and Systems Biology Program, Ann Romney Center for Neurological Diseases, Department of Neurology and Division of Genetics in Department of Medicine, Brigham and Women's Hospital, Boston, MA, Harvard Medical School, Boston, MA, and Program in Medical and Population Genetics, Broad Institute, Cambridge, MA

[29] Ann Romney Center for Neurologic Diseases, Brigham and Women's Hospital, Harvard Medical School, Boston, MA

[30] Department of Anesthesiology, Stony Brook University, Stony Brook, NY

[31] Section of Infections of the Nervous System, National Institute of Neurological Disorders and Stroke, NIH, Bethesda, MD

[32] Department of Neurology, Barrow Neurological Institute, St. Joseph's Hospital and Medical Center, Department of Neurobiology, Barrow Neurological Institute, St. Joseph's Hospital and Medical Center, Phoenix, AZ

[33] Department of Neurology, Division of Neuromuscular Medicine, Columbia University, New York, NY

[34] Department of Neuropathology, Academic Medical Center, University of Amsterdam, Amsterdam, The Netherlands

[35] Department of Biology and Veterinary and Biomedical Sciences, The Pennsylvania State University, University Park, PA

[36] New York Stem Cell Foundation, Department of Bioengineering, School of Engineering and Applied Sciences, University of Pennsylvania, Philadelphia, PA

[37] Department of Pathology, Fishberg Department of Neuroscience, Friedman Brain Institute, Ronald M. Loeb Center for Alzheimer's Disease, Icahn School of Medicine at Mount Sinai, New York, NY

[38] Department of Neurology, Harvard Medical School, Neurological Clinical Research Institute, Massachusetts General Hospital, Boston, MA

[39] Department of Neurology, Hospital for Special Surgery and Weill Cornell Medical Center, New York, NY

[40] Medical Genetics, Atlantic Health System, Morristown Medical Center, Morristown, NJ, and Overlook Medical Center, Summit, NJ

[41] Center of Clinical Research, Experimental Surgery and Translational Research, Biomedical Research Foundation of the Academy of Athens (BRFAA), 4 Soranou Efesiou Street, 11527, Athens, Greece; 1st Department of Neurology, Eginition Hospital, Medical School, National and Kapodistrian University of Athens, Athens, Greece

[42] Neuromuscular/EMG service and ALS/Motor Neuron Disease Clinic , Hebrew University-Hadassah Medical Center, Jerusalem, Israel

[43] Board of Governors Regenerative Medicine Institute, Los Angeles, CA; Department of Neurology, Cedars-Sinai Medical Center, Los Angeles, CA

[44] Neurological Clinical Research Institute, Massachusetts General Hospital, Boston, MA

[45] Departments of Neuroscience, and Genetics and Genomic Sciences, Ronald M. Loeb Center for Alzheimer's disease, Icahn School of Medicine at Mount Sinai, New York, NY

[46] Harvard Medical School, Department of Physical Medicine & Rehabilitation, Spaulding Rehabilitation Hospital, Boston, MA

[47] Center for Cellular and Molecular Therapeutics, Children's Hospital of Philadelphia, Philadelphia, PA; Department of Genetics, Perelman School of Medicine, University of Pennsylvania, Philadelphia, PA

[48] Centre for Clinical Brain Sciences, University of Edinburgh, Edinburgh, UK; Euan MacDonald Centre for Motor Neurone Disease Research, University of Edinburgh, Edinburgh, UK

[49] Department of Genetics and Genomic Sciences, Icahn Institute of Data Science and Genomic Technology, Icahn School of Medicine at Mount Sinai, New York, NY

[50] University of Maryland Brain and Tissue Bank and NIH NeuroBioBank

[51] Department of Neuropathology, Georgetown Brain Bank, Georgetown Lombardi Comprehensive Cancer Center, Georgetown University Medical Center, Washington DC

[52] Neuroradiology Section, Department of Radiology and Biomedical Imaging, University of California, San Francisco, San Francisco, CA

[53] Neuromuscular Diseases Unit, Department of Neurology, Tel Aviv Sourasky Medical Center, Sackler Faculty of Medicine, Tel-Aviv University, Tel-Aviv, Israel

[54] Department of Neuroscience, University of California San Diego, La Jolla, CA

[55] Department of Neurology, University of Pennsylvania Perelman School of Medicine, Philadelphia, PA

[56] Department of Neurology, Columbia University Medical Center, New York, NY

[57] Department of Pharmaceutical Chemistry, University of California San Francisco, San Francisco, CA

[58] Department of Pharmaceutical Chemistry, University of California San Francisco, San Francisco, CA

[59] Hadassah Hebrew University

[60] Department of Translational Neuroscience, Barrow Neurological Institute, Phoenix, Arizona

[61] The Translational Genomics Research Institute (TGen), Phoenix, Arizona

[62] Broad Institute, Cambridge, Massachusetts

[63] Broad Institute, Cambridge, Massachusetts

[64] Massachusetts General Hospital, Boston, Massachusetts

[65] Institute of Molecular Medicine, Feinstein Institutes for Medical Research, Northwell Health, Manhasset, New York

[66] Korea Advanced Institute of Science and Technology (KAIST), Daejeon, South Korea

[67] Maze Therapeutics

[68] Bristol-Myers Squibb

[69] Perron Institute for Neurological and Translational Science

[70] Integrative Immunobiology Section, National Institute of Allergy and Infectious Disease, NIH

[71] Neuromuscular Disease Research Section, National Institute of Aging

[72] Pfizer

[73] Regeneron

[74] Insitro
